# *In silico* analyses of maleidride biosynthetic gene clusters

**DOI:** 10.1101/2021.10.26.465875

**Authors:** Katherine Williams, Kate M. J. de Mattos-Shipley, Christine L. Willis, Andrew M. Bailey

## Abstract

Maleidrides are a family of structurally related fungal natural products, many of which possess diverse, potent bioactivities. Previous identification of several maleidride biosynthetic gene clusters, and subsequent experimental work, has determined the ‘core’ set of genes required to construct the characteristic medium-sized alicyclic ring with maleic anhydride moieties. Through genome mining, this work has used these core genes to discover ten entirely novel maleidride biosynthetic gene clusters, amongst both publicly available genomes, and encoded within the genome of the previously un-sequenced epiheveadride producer *Wicklowia aquatica* CBS125634. We have undertaken phylogenetic analyses and comparative bioinformatics on all known and putative maleidride biosynthetic gene clusters to gain further insights regarding these unique biosynthetic pathways.

## 1. Introduction

Maleidrides are an important bioactive family of polyketide-derived secondary metabolites, produced by diverse filamentous fungi and are characterised by a medium-sized alicyclic ring with one or two fused maleic anhydride moieties. The majority of reported maleidrides are nonadrides, assembled on a 9-membered ring core, such as byssochlamic acid **1** and the glauconic **2** and glaucanic **3** acids.^1^ More recently, octadrides (with an 8-membered central ring) such as zopfiellin **4**^2^ and viburspiran **5**,^3^ and heptadrides (7-membered central ring) such as agnestadride A **6**^4^ have been isolated (Figure 1).

**Figure 1:**
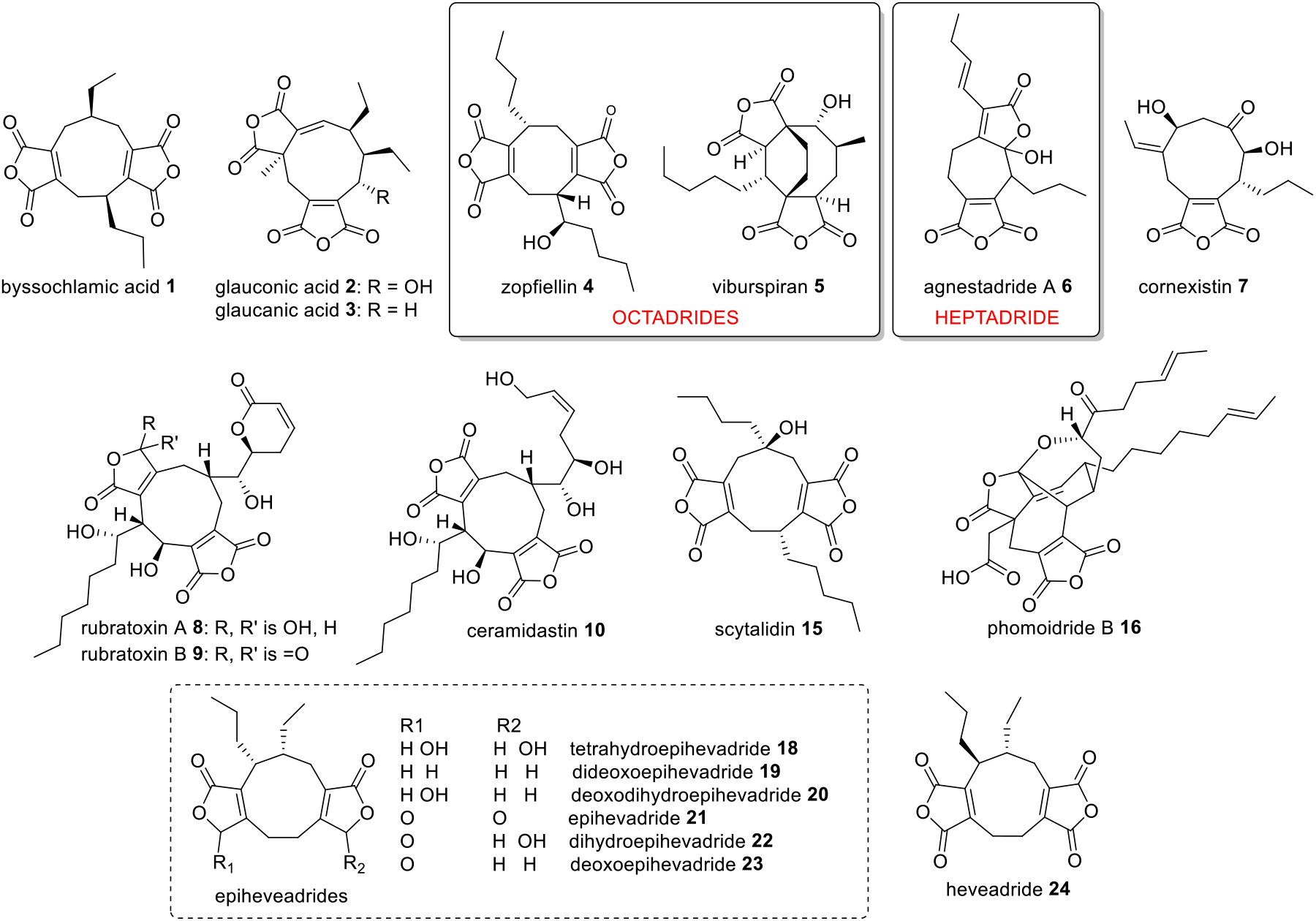
Examples of maleidride compounds. All are nonadrides except where otherwise noted. Numbering system described by de Mattos-Shipley *et al*.^12^

Most known maleidrides possess bioactivities which make them attractive pharmaceutical or agrochemical lead compounds, such as the selective herbicide cornexistin **7**,^5^ the antifungal zopfiellin **4**,^6^ or the protein phosphatase 2A (PP2A) inhibitor, rubratoxin A **8**.^7^ It is interesting to note that minor structural changes can lead to significant shifts in potency or target, for example, the γ-hydroxybutenolide motif present in the PP2A inhibitor rubratoxin A **8** makes it ∼100 times more potent than its anhydride analogue rubratoxin B **9**.^7^ Another rubratoxin analogue, ceramidastin **10**, which lacks the α,β-unsaturated lactone moiety of the rubratoxins, is a novel inhibitor of bacterial ceramidase,^8^ an enzyme thought to contribute to skin infections of patients with atopic dermatitis.^9^

In recent years, studies^10-15^ have begun to reveal the genetic basis and complexity of the biosynthetic pathways to these fascinating molecules. Isotopic labelling,^16-18^ heterologous expression experiments, ^10,13,15^ gene knock-outs^11,12,14^ and *in vitro* work^19^ and have led to the proposal that the biosynthetic pathway begins with precursors (e.g. **11**) assembled by an iterative highly reducing polyketide synthase (hrPKS), where the chain length and level of saturation varies according to the structure of the mature natural product. Coupling of **11** with oxaloacetate is catalysed by an alkylcitrate synthase (ACS), followed by dehydration catalysed by an alkylcitrate dehydratase (ACDH) to generate the maleidride type A monomer **12** (Scheme 1).^10,13^ Decarboxylation then gives the tautomeric structures **13** and **14** (types B and C). The next step involves coupling of the monomers in different modes and cyclisation to form the various carbocyclic rings. This involves maleidride dimerising cyclases (MDCs – see section 2.2.2) and phosphatidylethanolamine binding protein-like (PEBP) enzymes (Scheme 1).^10^ The mode of cyclisation determines the carbon framework, with tailoring modifications leading to the observed diversity of natural products (Figure 1). These experiments have defined the core genes required for maleidride biosynthesis, which enables the rapid discovery of additional maleidride biosynthetic gene clusters (BGCs). Based on this knowledge this work set out to identify maleidride BGCs from both in-house and publicly available genome sequences; linking them to compounds where possible and performing phylogenetic analyses and comparative bioinformatics to gain further insights regarding these unique biosynthetic pathways.

**Scheme 1:**
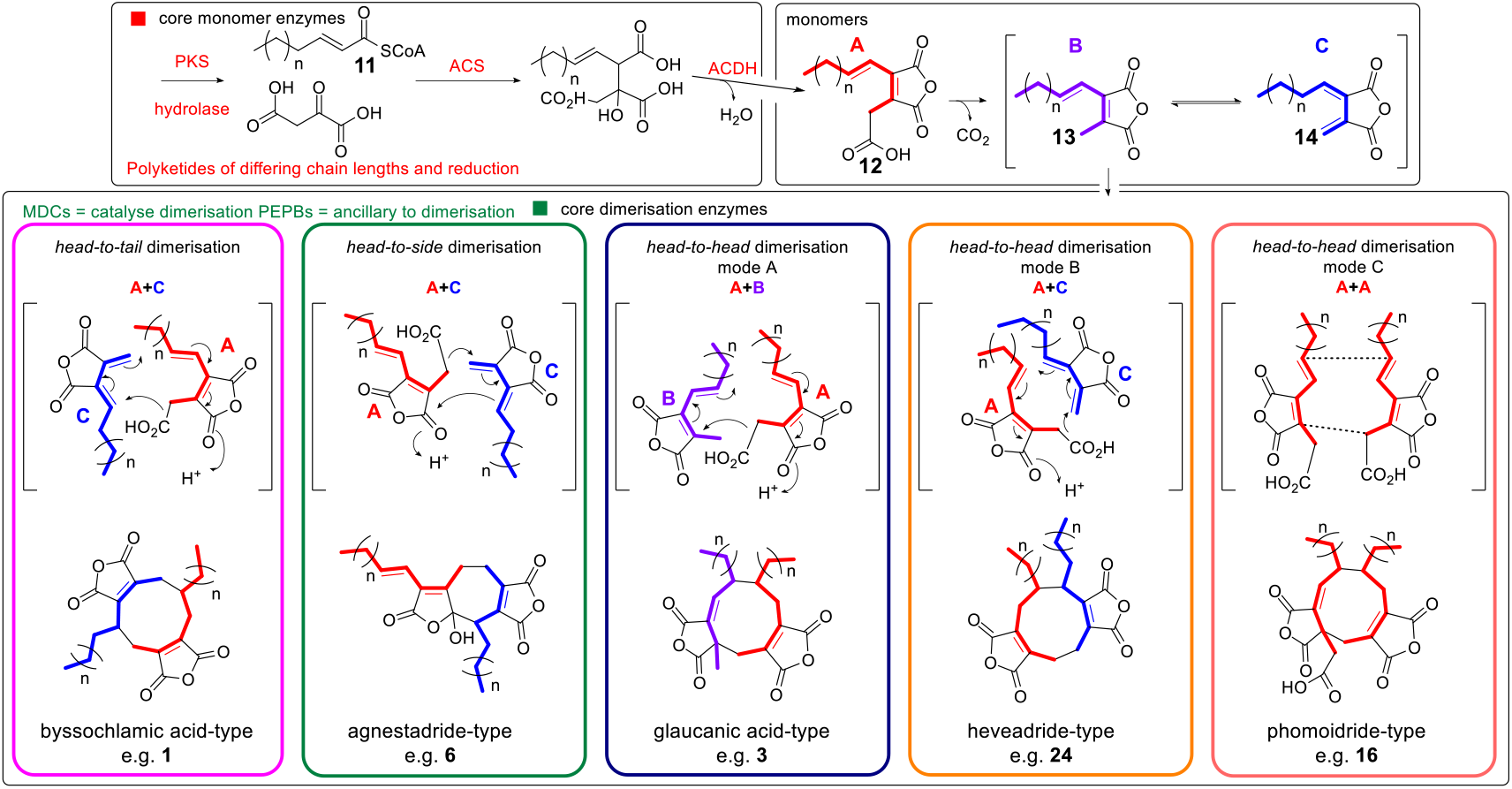
Proposed biosynthesis of maleidride monomers and dimerisations required to construct the diverse structures of known maleidrides.

## 2. Results and Discussion

### 2.1. Identification of maleidride BGCs

#### 2.1.1. Previously identified maleidride BGCs

In fungi the genes required for the biosynthesis, regulation and transport of a specific natural product are generally co-located as a single biosynthetic gene cluster (BGC).^20,21^ Six BGCs have been identified and definitively linked to the biosynthesis of specific maleidrides through experimental approaches including gene disruptions and heterologous expression: the byssochlamic acid **1** / agnestadrides (e.g. **6**) BGC,^10^ the rubratoxin (e.g. **8**) BGC,^14^ the cornexistin **7** BGC,^11^ two zopfiellin **4** BGCs,^12,15^ and the scytalidin **15** BGC.^12^ Oikawa and colleagues identified a single likely maleidride BGC in the genome of the known phomoidride (e.g. **16**) producer, the unidentified fungus ATCC 74256.^13^ Heterologous expression of the hrPKS, the ACS and the ACDH in *Aspergillus oryzae* resulted in the production of a new metabolite which the authors proposed was a maleidride monomer e.g. **12**, but yields of this compound were too low for purification and structural elucidation. Expression of homologous genes from an orphan maleidride BGC identified in the genome of *Talaromyces stipitatus* (referred to as *T. stipitatus* cluster 1 in this work), led to the isolation and characterisation of **17** (Figure 2), a hexaketide-derived type B maleidride monomer (see Scheme 1, compound **13**).^13^ Although the *T. stipitatus* BGC is not necessarily a phomoidride cluster, based on the chain length and saturation pattern it was proposed that **17** is the decarboxylated phomoidride monomer.^13^ *T. stipitatus* has not been confirmed as a producer of maleidrides, but many *Talaromyces* species produce glauconic acid **2** and glaucanic acid **3**, as well as the more complex rubratoxins (e.g. **8**).^22^

**Figure 2:**
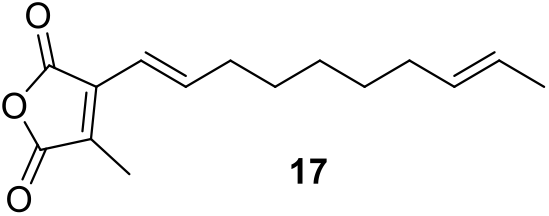
Hexaketide maleidride type B monomer isolated from *T. stipitatus* BGC 1 genes *tstA,I,J* expressed in *A. oryzae*.

#### 2.1.2. Newly identified maleidride BGCs

Due to our ongoing interest in maleidrides, we have sequenced the genome of *Wicklowia aquatica* strain CBS125634, a known producer of epiheveadrides **18** - **23**.^23^ One candidate BGC has been identified (see Figure S40 and Table S3 for details, GenBank accession OK490366).

To identify additional maleidride BGCs in publicly available genomes, a cblaster^24^ analysis was carried out. This analysis searches for homologues to a number of ‘query’ protein sequences that are co-located within a genome. All of the confirmed maleidride BGCs contain a ‘core’ set of genes, which have previously been shown through heterologous expression to be capable of producing maleidrides in high yields,^10^ namely: a polyketide synthase (PKS), an AlnB-like hydrolase, an alkylcitrate synthase (ACS), an alkylcitrate dehydratase (ACDH), an MDC and a PEBP-like protein (See Table S1). These core protein sequences from the byssochlamic acid pathway were used as queries to search the NCBI database, which highlighted a number of known BGCs as well as identifying twelve orphan BGCs (Figure 3). Three of these orphan BGCs (from *Oidiodendron maius Zn, Aspergillus glaucus* and *Imshaugia aleurites*) lack any genes encoding PEBP-like proteins (Figure 3). However, as previous experimental work has demonstrated that these enzymes are not an absolute requirement for maleidride biosynthesis (see Section 2.2.2 for a more detailed discussion), these three clusters were still considered likely maleidride BGCs.

**Figure 3:**
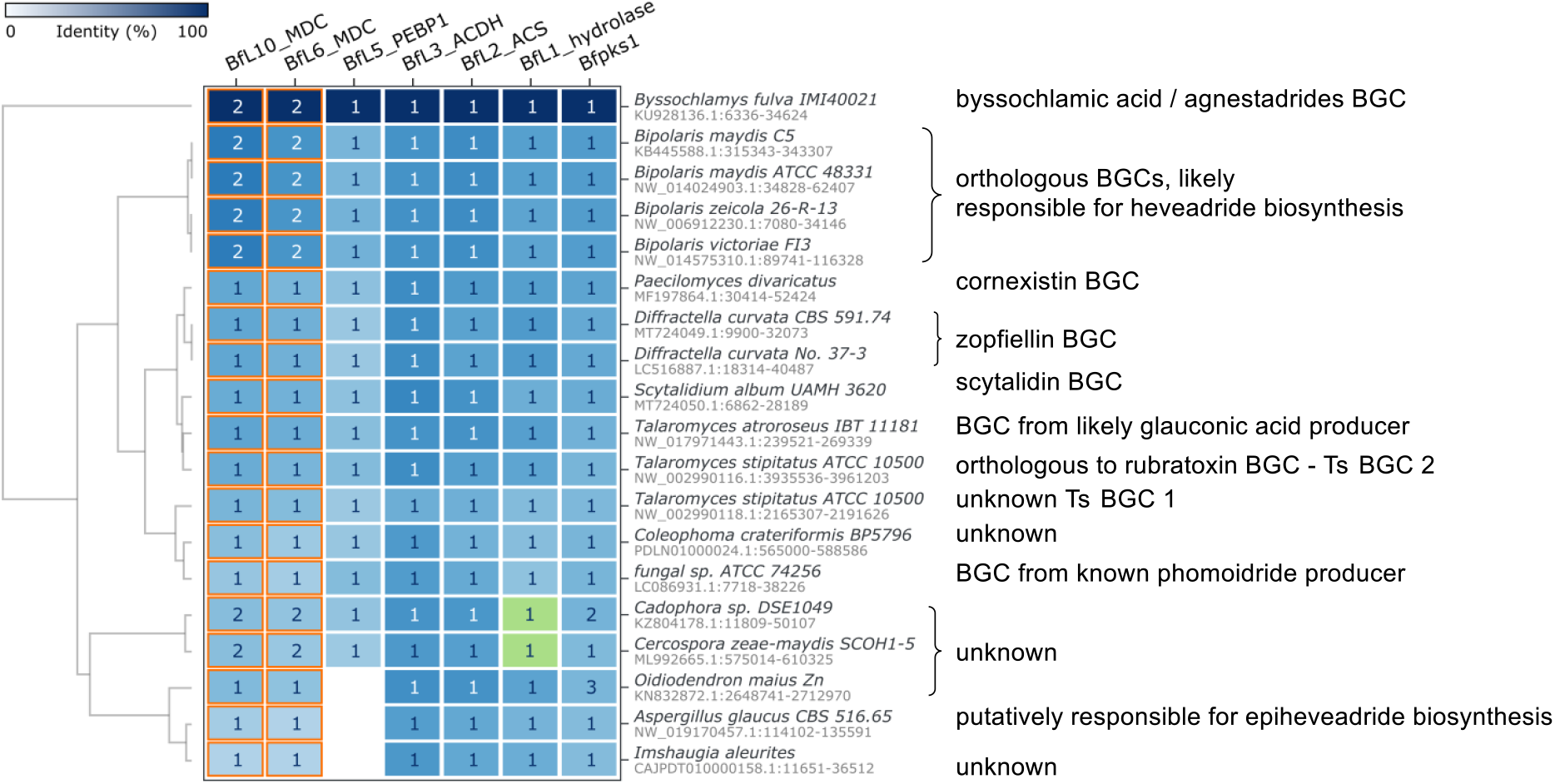
cblaster^24^ analysis searching for co-localised proteins (defined as within 50 kb) in annotated fungal genomes within the NBCI database. BfPKS1 (PKS), BfL1 (hydrolase), BfL2 (ACS), BfL3 (ACDH), BfL5 (PEBP1), BfL6 (MDC) and BfL10 (MDC) were used as queries. BfPKS1, BfL2, BfL3 and BfL6 were set as required hits. The number within each box denotes the number of hits to a specific query sequence within the co-localised region. The orange cell border denotes proteins which generated hits with multiple query sequences. The green boxes denote instances where cblaster failed to identify an annotated homologue, but manual interrogation of the DNA sequence has identified a previously unannotated homologue. cblaster uses the BLASTp algorithm.

Four of the newly identified BGCs are from closely related *Bipolaris* sp. and are likely to be orthologous BGCs based on their identical gene organisation and the very high protein identity of encoded proteins (See Figure S40). It is probable that these BGCs are responsible for the biosynthesis of the nonadride, heveadride **24**, as *Bipolaris* species are known to produce this compound.^25-27^ Furthermore, these orthologous BGCs all demonstrate significant similarity to the BGC we have now identified from *Wicklowia aquatica*, the confirmed producer of the epiheveadrides **18** - **23** (See Figure S40 and Table S3). A BGC identified from *Aspergillus glaucus* (synonym *Eurotium herbariorum*) may also be responsible for the biosynthesis of epiheveadride **21**, as this compound is known to be produced by certain strains of this species.^28^ It is noteworthy however that this BGC differs quite significantly from both the *Bipolaris* and *W. aquatica* BGCs, lacking homologues to certain encoded proteins such as the type 1 PEBP, AMP dependant CoA ligase and isochorismatase and with additional uniquely encoded proteins such as two P450s and a methyltransferase (See Figure S40 and Table S3).

Samson and co-workers^29^ detected a compound with the same chromophore and retention time as glauconic acid **2** from extracts of *Talaromyces atroroseus* IBT 11181. This suggests that the maleidride BGC identified within the *T. atroroseus* IBT 11181 genome may encode the production of **2**, and likely glaucanic acid **3**, a predicted precursor to **2**. ^18,29^ Two maleidride BGCs were identified within the genome of *T. stipitatus* ATCC 10550. *T. stipitatus* cluster 1 is the same BGC that was utilised in heterologous expression experiments by Oikawa and colleagues when investigating phomoidride (e.g. **16**) biosynthesis (see Section 2.1.2), although it has not been unequivocally identified as a phomoidride BGC. *T. stipitatus* cluster 2 is highly likely to encode rubratoxin (e.g. **8**) biosynthesis. The sequence for the confirmed rubratoxin BGC of *Penicillium dangeardii* Pitt is not publicly available,^14^ but *T. stipitatus* cluster 2 demonstrates complete synteny in terms of the encoded activities and order of the genes in the cluster, therefore we have named the genes within this cluster according to the *P. dangeardii* Pitt BGC (see Figure S37). The five other orphan maleidride BGCs identified through cblaster^24^ cannot be linked to the production of any known maleidrides, so are likely to encode novel maleidrides.

MultiGeneBlast^30^ was utilised to search unannotated genomes using the tBLASTn algorithm, where hits on NCBI had suggested the presence of a possible maleidride BGC. This process identified two further putative maleidride BGCs, from the species *Talaromyces borbonicus* CBS 141340 and *Talaromyces funiculosus* X33. There have been no reports of either of these species producing maleidrides, so they cannot easily be linked to specific compounds, but the BGC from *Talaromyces funiculosus* is clearly closely related to the potential rubratoxin BGC identified from the *T. stipitatus* genome (Figure S38), referred to here as *T. stipitatus* cluster 2. Almost complete synteny exists between the *T. funiculosus* BGC and *T. stipitatus* cluster 2 from *TsrbtI* to *TsrbtT*, but the *T. funiculosus* BGC lacks the genes required for most of the tailoring steps to produce mature rubratoxins (RbtA, RbtH, RbtB, RbtU and RbtG). This suggests that the *T. funiculosus* BGC might encode the biosynthesis of a compound similar to the rubratoxin core structure, but with fewer oxidative modifications.

clinker^31^ comparisons of all of the above BGCs (Figure 4 and Figure 5) highlight the many genes which are core to, or common within, maleidride clusters, as well as identifying uniqueness, particularly in clusters known to encode heavily derivatised maleidrides. All clusters linked to specific compounds – through experimental verification, identification from known producers, or clear synteny with a confirmed maleidride BGC – are summarised in Figure 4 and Table S1. All orphan maleidride BGCs – which can not yet be confidently linked to specific compounds – are summarised in Figure 5 and Table S2.

**Figure 4:**
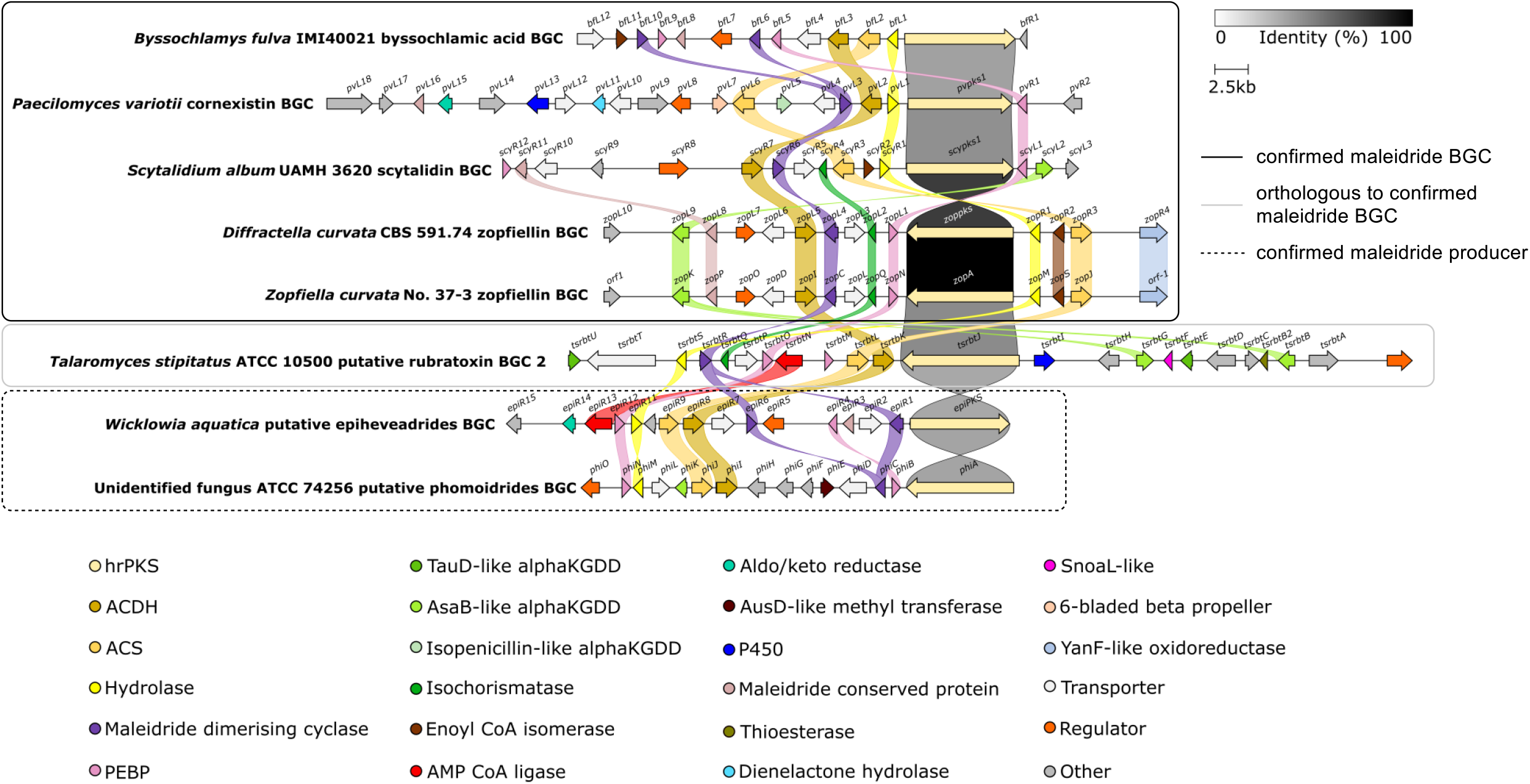
clinker^31^ comparison between definitively linked maleidride BGCs (through gene knockout or heterologous expression), as well as those identified from the genomes of confirmed maleidride producing strains. The *T. stipitatus* cluster 2 is included as there is clear evidence that it is an orthologous rubratoxin BGC (see Figure S37). Links between homologous genes are shown using their specific colour, except for the PKSs where the links are shown according to the percentage identity (see identity scale bar). Links between transport and regulatory genes have been removed for clarity. See Table S1 for putative functions for uniquely encoded proteins and accession numbers for all proteins.

**Figure 5:**
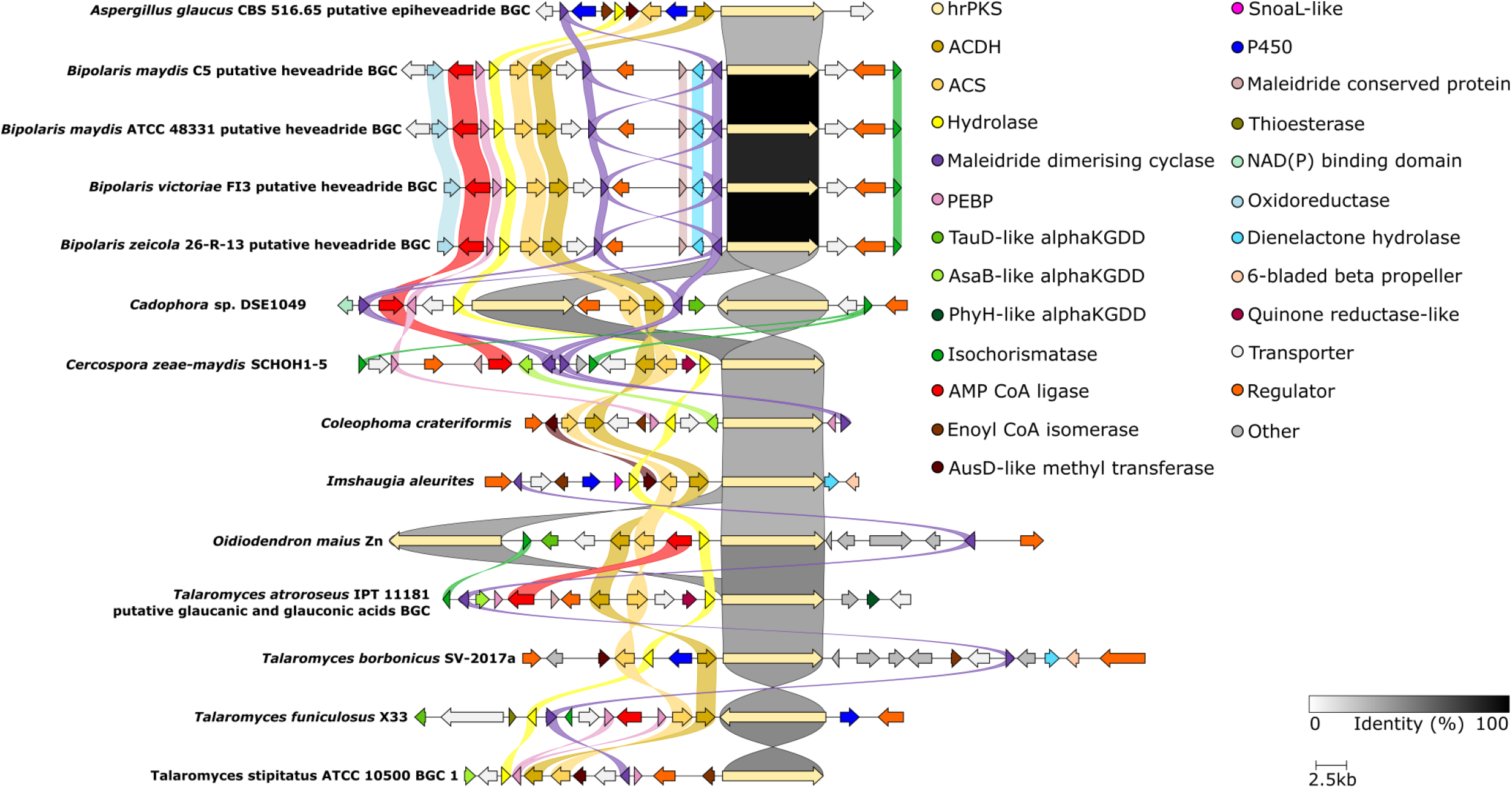
clinker^31^ comparison between orphan maleidride BGCs. Links between homologous genes are shown using their specific colour, except for the PKSs where the links are shown according to the percentage identity (see identity scale bar). Links between transport and regulatory genes have been removed for clarity. See Table S2 for accession numbers and putative functions for uniquely encoded proteins.

### 2.2. Core maleidride enzymes

As discussed previously, and seen clearly in Table S1, the core maleidride enzymes can be defined as the PKS, the hydrolase, ACS, the ACDH, the MDC and the PEBP-like type 1 proteins. This has been experimentally verified in the heterologous host *Aspergillus oryzae*, where expression of this core set of genes resulted in the production of the simple maleidride byssochlamic acid **1** in good yields.^10^

#### 2.2.1. Maleidride monomer forming enzymes

##### 2.2.1.1. PKS

All known and putative maleidride PKSs contain the domains normally present in a fungal hrPKS: a ketosynthase (KS - IPR020841); an acyltransferase (AT - IPR014043); a dehydratase (DH - IPR020807); a C-methyltransferase (CMet - IPR029063); an enoyl reductase (ER - IPR020843); a ketoreductase (KR - IPR013968); and an acyl-carrier protein (ACP - IPR009081) (Figure 6, A). Although all maleidride PKSs contain a CMet domain, none of the known maleidride structures suggest a requirement for methylation of the polyketide chain. Similarly, in cases where the maleidride monomer has been isolated, there is no evidence of methylation. ^10,11,13,15^ Conserved CMet motifs^32^ are present in several maleidride PKSs, but most have critical mutations in at least one motif (Figure S42). However, these conserved motifs are also found in other PKSs which synthesise non-methylated PKSs,^33,34^ which suggests that control of methylation by fungal hrPKSs is still not understood.^32^ The maleidrides PKSs range from 2416 – 2620 amino acids in length and demonstrate at least 42 % protein identity, (See Table S4 for % identity matrix).

**Figure 6:**
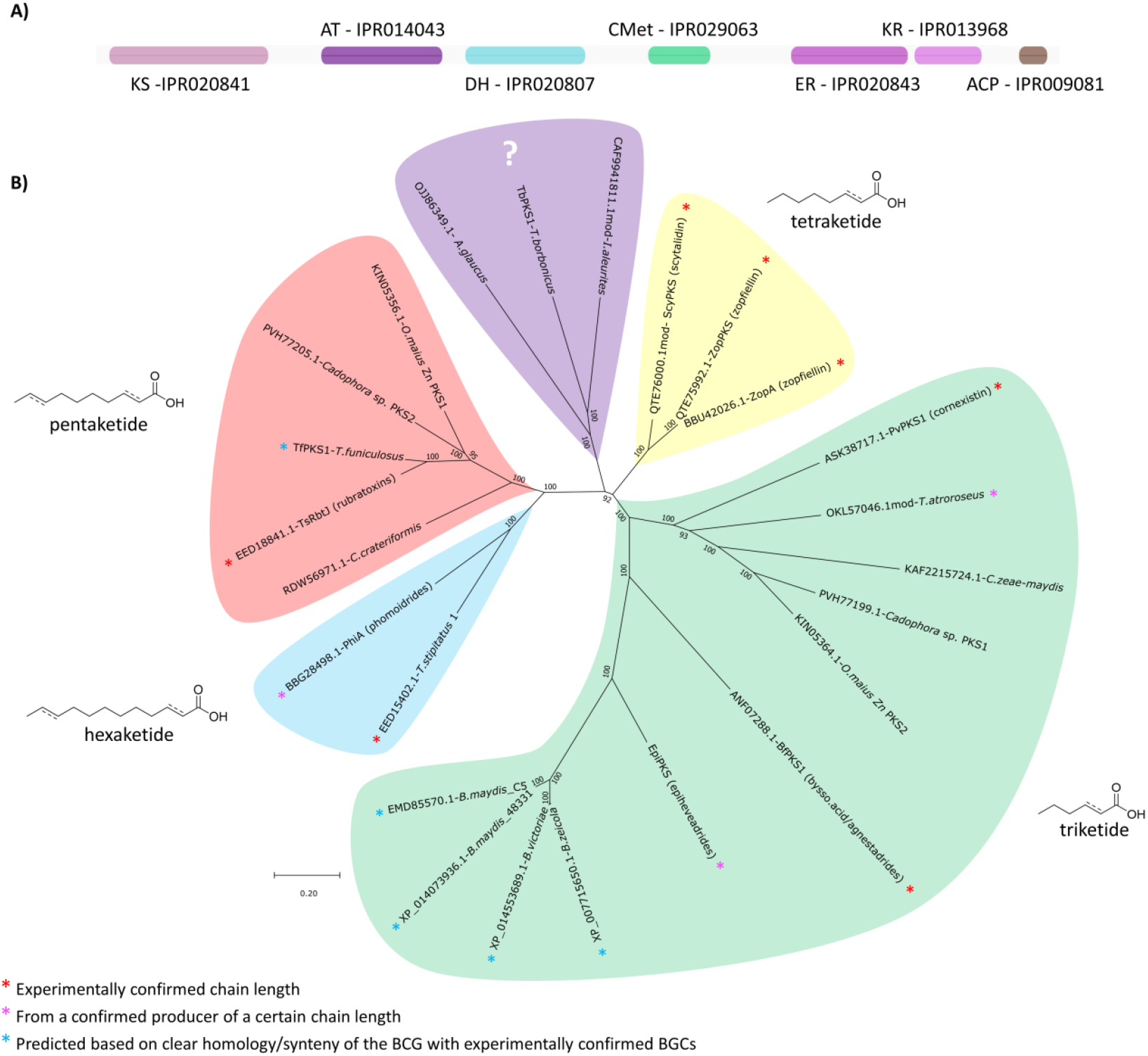
A) Diagram depicting the organisation of domains within a maleidride PKS. B) An unrooted phylogenetic analysis of PKSs from known and predicted maleidride BGCs. The evolutionary history was inferred by using the Maximum Likelihood method and Le_Gascuel_2008 model.^36^ The tree with the highest log likelihood (−62209.96) is shown. A discrete Gamma distribution was used to model evolutionary rate differences among sites (5 categories (+G, parameter = 1.1913)). The rate variation model allowed for some sites to be evolutionarily invariable ([+I], 8.34% sites). The tree is drawn to scale, with branch lengths measured in the number of substitutions per site. This analysis involved 24 amino acid sequences. There were a total of 2339 positions in the final dataset. Evolutionary analyses were conducted in MEGA X.^37,38^

Maleidride PKSs are known to produce polyketides of varying chain lengths. Triketide-derived monomers go on to produce many maleidrides including cornexistin **7**, byssochlamic acid **1**, the agnestadrides (e.g. **6**), the (epi)heveadrides **18** to **24**, and the glauconic and glaucanic acids **2** and **3**; tetraketides form the backbone of both zopfiellin **4** and scytalidin **15**; the rubratoxins (e.g. **8**) are biosynthesised from pentaketides; and currently the longest known chain-lengths are the hexaketides seen in phomoidride (e.g. **16**) biosynthesis. A phylogenetic analysis shows that the polyketide synthases fall into distinct clades based on the chain length they produce (Figure 6, B). This relationship allows the products of new uncharacterised maleidride PKSs to be predicted. Regarding the newly identified and uncharacterised PKSs from single PKS BGCs, the PKS from *Coleophoma crateriformis* (KAF2215724.1) falls within the likely pentaketide producing clade. Sitting in a separate clade which does not contain any previously characterised maleidride PKSs, and therefore difficult to predict, are those of *Talaromyces borbonicus* (TbPKS1), *Aspergillus glaucus* (OJJ86349.1) and *Imshaugia aleurites* (CAF9941811.1mod). Focussing on the clusters which contain two distinct PKS encoding genes, namely those of *O. maius Zn* and *Cadophora* sp., the individual PKSs clearly fall into different clades (Figure 6). CadPKS2 (PVH77205.1) and OmPKS1 (KIN05356.1) are similar to each other (75 % protein identity) and both cluster with the pentaketide producing PKS (EED18841.1) from the *T. stipitatus* rubratoxin cluster (showing 70.5 % and 72.9 % protein identity respectively). CadPKS1 (PVH77199.1) and OmPKS2 (KIN05364.1) are again most similar to each other (72.2 % protein identity) and both cluster with the known triketide producing cornexistin PKS - ASK38717.1 (56.3 % and 55.5 % protein identity respectively).

In relation to the clusters containing two PKSs, it is unknown whether both encoded enzymes are involved in maleidride biosynthesis, but it is intriguing to consider whether these pathways could involve the heterodimerisation of distinct monomers biosynthesised by different polyketide synthases. Another possibility is that one PKS is involved in the biosynthesis of the maleidride monomer, and the other is involved in downstream decorations, as is seen in some fungal secondary metabolites, such as the cryptosporioptide pathway.^35^

##### 2.2.1.2. Maleidride hydrolase

All known and putative maleidride BGCs contain a hydrolase homologue, strongly suggesting it is important for the biosynthesis of maleidride compounds. Interpro^39^ analysis shows all the identified putative maleidride hydrolases contain an α/β hydrolase fold (IPR029058), which is common to a number of hydrolytic enzymes of differing catalytic function.^40^ They all contain the conserved active site of thiolesterase enzymes (Figure S43), consisting of a catalytic triad: a nucleophile (usually serine), an acidic residue (usually aspartic acid), and a histidine,^40,41^ categorising them within the serine hydrolase superfamily (IPR005645).^42,43^ Additionally, Interpro^39^ analysis shows that all the identified putative maleidride hydrolases fall within the subfamily PTHR48070:SF4, the AlnB-like esterases. AlnB is involved in the biosynthesis of the polyketide-based compound, asperlin.^44^ Deletion of *alnB* from the asperlin producer, *Aspergillus nidulans*, completely halted asperlin biosynthesis, demonstrating that it is an essential early step in this pathway, and it was proposed that AlnB is responsible for the hydrolytic release of the polyketide from the PKS.^44^ *In vitro* work with the *B. fulva* hydrolase (BfL1) is consistent with this, showing that BfL1 rapidly hydrolyses acyl-ACP species. How *in vivo* selectivity is controlled is currently unknown, as the *B. fulva* hydrolase also rapidly hydrolysed other thiolesters.^19^

The absolute requirement for the action of the hydrolase in maleidride biosynthesis is difficult to ascertain, given contradictory results from different pathways. Heterologous expression experiments by Oikawa and co-workers^13,15^ to investigate maleidride monomer formation from the phomoidride (e.g. **16**), zopfiellin **4** and *T. stipitatus* 1 BGCs did not include hydrolases, despite each cluster containing such a gene. Low monomer yields were seen in the cases of the *phi* and *zop* BGCs, which may indicate that the hydrolase is required for the pathway to function efficiently. Work by Cox and co-workers^10,11^ has strongly suggested that in the pathway for byssochlamic acid **1** and the agnestadrides (e.g. **6**), as well as cornexistin **7**, the hydrolase is essential. One possible explanation for the apparent contradictions is the action of an endogenous enzyme in the heterologous host, *A. oryzae*. A homologue of the maleidride hydrolases was identified from the genome of *A. oryzae*, which shows 40.9 - 57.7 % sequence identity to maleidride hydrolases (Table S5). A phylogenetic analysis, (Figure S51) clearly shows that OOO04461.1 clades with other maleidride hydrolases. Interestingly, this hydrolase is situated next to an hrPKS (OOO04460.1), although no other maleidride related genes are co-located.

It is noteworthy that the clusters with two PKSs (see section 2.2.1.1) do not have two hydrolases, highlighting that one hydrolase may catalyse the release of different polyketide chains from different synthases. This is consistent with the previous observation that the hydrolase from the *B. fulva* byssochlamic acid pathway has relaxed substrate specificity.

##### 2.2.1.3. Alkylcitrate synthase (ACS) and Alkylcitrate dehydratase (ACDH)

A key step in the biosynthesis of maleidrides - as well as other related fungal natural products such as hexylaconitic anhydride^45^ and the cordyanhydrides^46^ - is the formation of the maleic anhydride moiety. Barton and Sutherland first proposed in 1965 that maleidride monomers are modified products of the citric acid cycle,^47^ this hypothesis informed the identification of the first maleidride BGCs, ^10,13^ which each contain a citrate synthase (CS)-like enzyme and a 2-methylcitrate dehydratase (2MCDH)-like enzyme. Based on the roles of CS and 2MCDH enzymes in primary metabolism, these enzymes were predicted to catalyse the condensation of oxaloacetate with a polyketide chain to give an alkyl citrate, followed by dehydration to produce the maleic acid and its equilibrating anhydride (see Scheme 1). This has been borne out by heterologous expression experiments, and more recently, by detailed *in vitro* experiments. The *in vitro* work in particular^13,15,19^ has confirmed that the CS-like enzymes involved in maleidride biosynthesis require an CoA thiolester substrate, rather than directly reacting with polyketide produced ACP-acyl substrates. This is analogous to both the citrate synthases and 2-methylcitrate synthases from primary metabolism, which require acetyl-CoA and propionyl-CoA respectively as their substrates, and which catalyse both the condensation with oxaloacetate and hydrolytic release of CoA. Consistent with this functional homology is the presence of a citrate synthase domain (PTHR11739:SF4) and conserved acyl CoA-binding residues within all of the maleidride CS-like enzymes, including those newly identified in this work (Figure S44).

The work by Cox and colleagues^19^ also demonstrated that both the CS-like and 2MCDH-like enzymes from the byssochlamic acid pathway follow exactly the same stereochemical course as their primary metabolism counterparts. In order to further clarify the evolutionary relationship of these enzymes, a phylogenetic analysis was conducted, which included known primary metabolism enzymes, as well as the maleidride CS-like enzymes and CS-like enzymes from other secondary metabolite pathways (alkylcitric acids,^45^ the oryzines,^48^ the sporothriolides^49^ and squalestatin;^50-52^ a comparison of these BGCs with the byssochlamic acid BGC can be seen in Figure S41). As can be seen in Figure 7, primary metabolism citrate synthases from a range of different species form a tight clade. Known methylcitrate synthases fall within two clear clades, consisting of those from fungi and those from bacteria, and interestingly, the enzymes from fungal secondary metabolite pathways appear to be more closely related to the bacterial methylcitrate synthases. CS-like enzymes from secondary metabolism, which are known to function as alkylcitrate synthases (ACSs), all fall within a monophyletic group, with the maleidride CS-like enzymes branching off later than those involved in other secondary metabolite pathways. The clear functional and evolutionary relationship of these enzymes emphasises that it is reasonable to refer to them as alkylcitrate synthases, rather than the previously used and more tentative title of ‘citrate synthase (CS)-like’ enzymes.

**Figure 7:**
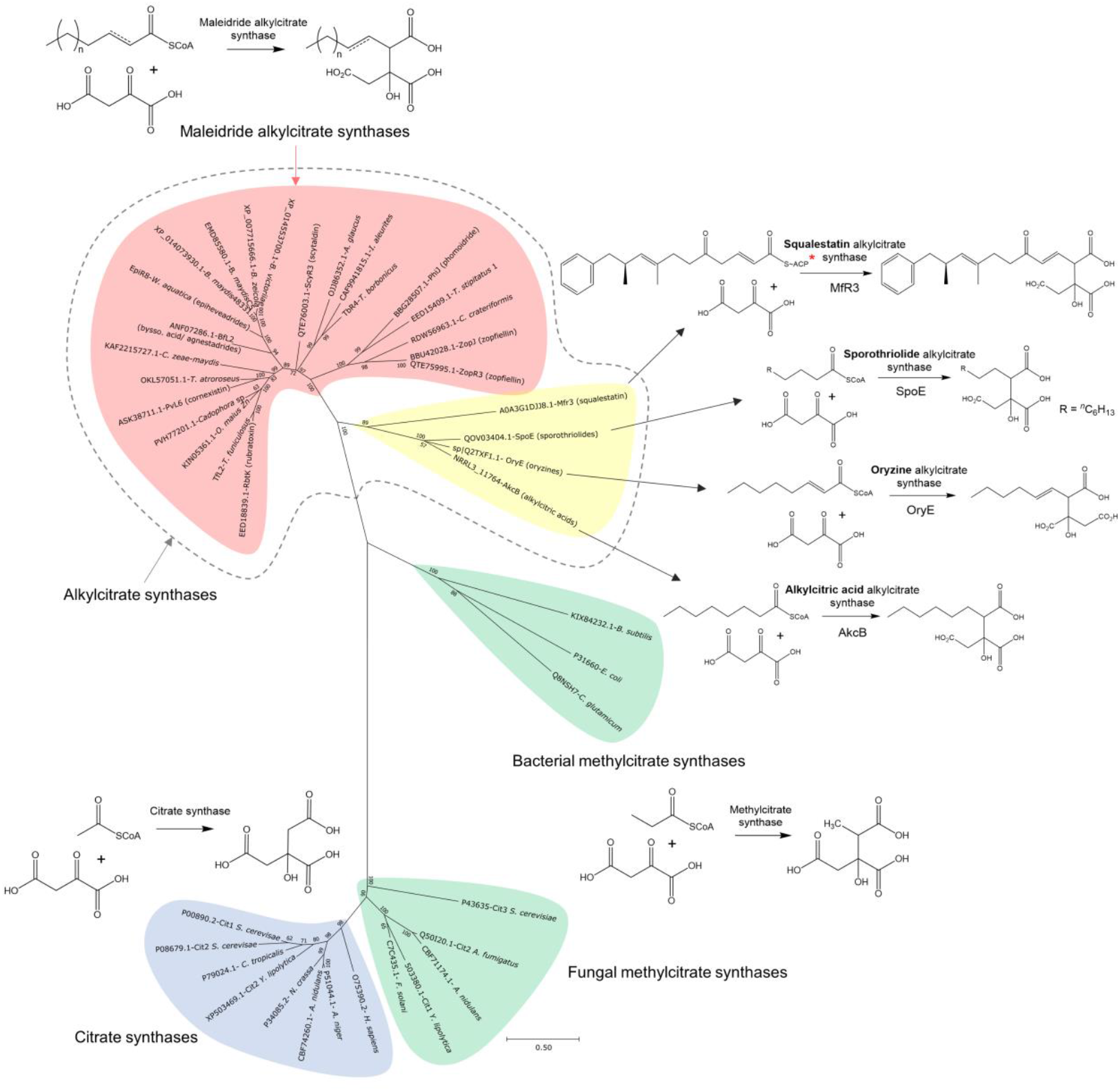
An unrooted phylogenetic analysis of various citrate synthases, methylcitrate synthases and alkylcitrate synthases (ACSs), including all from known and putative maleidride BGCs. The evolutionary history was inferred by using the Maximum Likelihood method and Le_Gascuel_2008 model.^36^ The tree with the highest log likelihood (−17034.49) is shown. The percentage of trees in which the associated taxa clustered together is shown next to the branches. A discrete Gamma distribution was used to model evolutionary rate differences among sites (5 categories (+G, parameter = 1.8246)). The rate variation model allowed for some sites to be evolutionarily invariable ([+I], 2.56% sites). The tree is drawn to scale, with branch lengths measured in the number of substitutions per site. This analysis involved 43 amino acid sequences. There were a total of 360 positions in the final dataset. Evolutionary analyses were conducted in MEGA X.^37^ **\*** It has been proposed that the ACS from the squalestatin pathway directly accepts the ACP bound acyl chain as a substrate (as shown here) but this has not been conclusively demonstrated.

**Figure 8:**
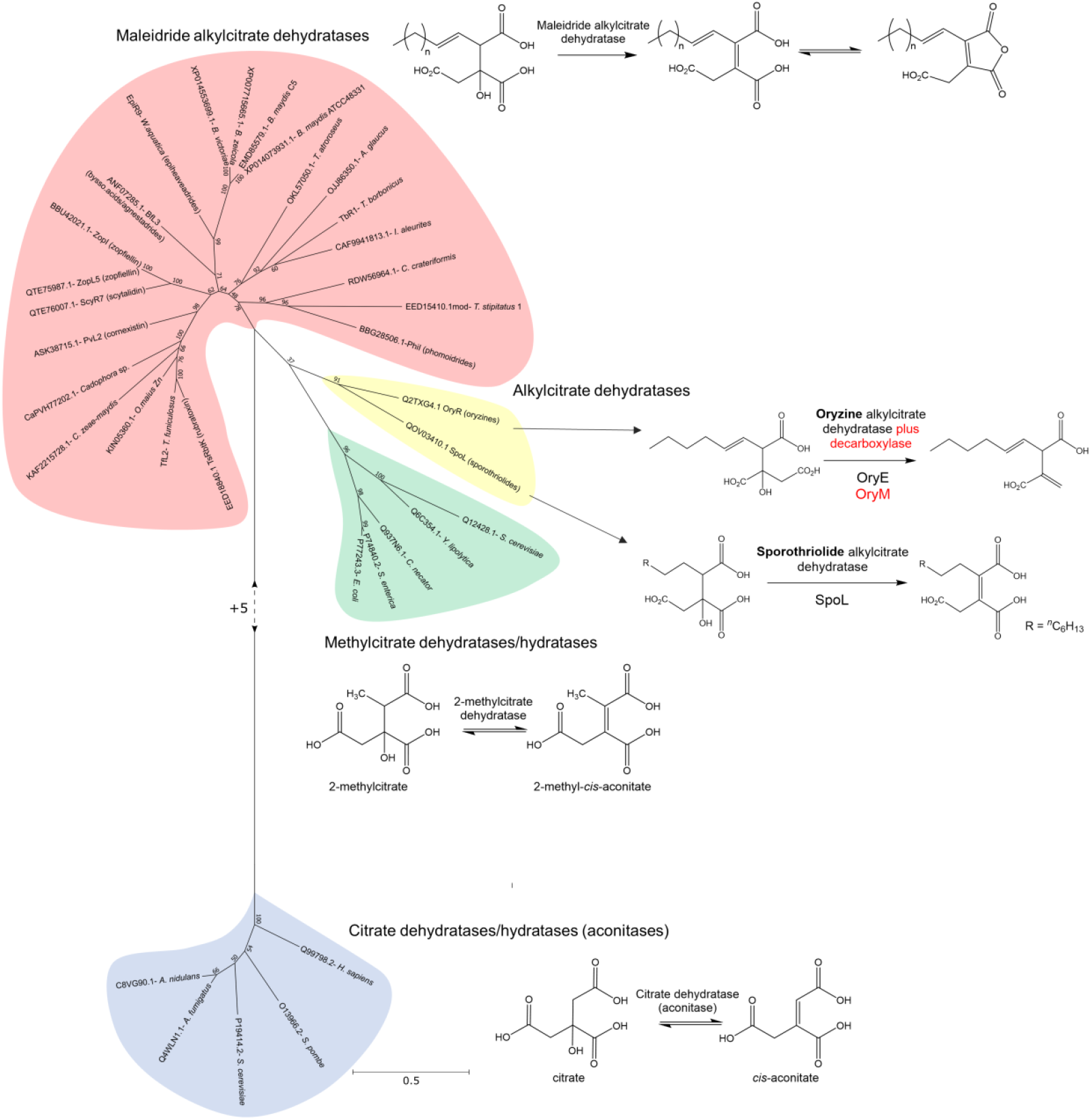
An unrooted phylogenetic analysis of various citrate dehydratases, 2-methylcitrate dehydratases and alkylcitrate dehydratases. The evolutionary history was inferred by using the Maximum Likelihood method and Le_Gascuel_2008 model.^36^ The tree with the highest log likelihood (−17005.47) is shown. The percentage of trees in which the associated taxa clustered together is shown next to the branches. A discrete Gamma distribution was used to model evolutionary rate differences among sites (5 categories (+G, parameter = 1.1622)). The rate variation model allowed for some sites to be evolutionarily invariable ([+I], 0.00% sites). The tree is drawn to scale, with branch lengths measured in the number of substitutions per site. This analysis involved 34 amino acid sequences. There were a total of 466 positions in the final dataset. Evolutionary analyses were conducted in MEGA X.^37^

The phylogenetic analysis (Figure 7) clearly supports the previously proposed scenario,^19^ whereby citrate synthases evolved to accept larger substrates, starting with methylcitrate, then expanding to longer chain fatty acids produced by typical fungal fatty acid synthases (as is the case for the alkylcitric acids,^45^ sporothriolides^49^ and likely the oryzines^48^), and finally by pairing up with polyketide synthases in the biosynthesis of squalestatin^50-52^ and the maleidrides.^10-15^ All of the ACSs in this analysis catalyse a condensation with oxaloacetate, but there are fungal natural products in the literature (e.g. the cordyanhydrides^46,13^) which are likely to come about *via* the condensation of an acyl-CoA with another keto diacid such as α-ketoglutarate (αKG). As none of these enzymes have been identified, it is not possible to infer their evolutionary relationship to the alkylcitrate synthases, although it is likely that they are closely related, and may have evolved more relaxed substrate specificity in regards the keto diacid as well as the acyl-CoA substrate.

A similar evolutionary relationship can be seen with the 2-methylcitrate dehydratase-like enzymes. The dehydratases involved in fungal secondary metabolism are closely related to microbial 2-methylcitrate dehydratases involved in propionate metabolism, but are relatively distantly related to citrate dehydratases. The dehydratases from the PKS-based maleidride pathways sit within a closely related monophyletic group, and the dehydratases from the sporothriolide and oryzine pathways, which are FAS-based biosynthetic pathways, sit within a separate clade. We propose that enzymes within both clades can be referred to as alkylcitrate dehydratases (ACDHs).

#### 2.2.2. Dimerisation enzymes

One of the most unusual aspects of maleidride biosynthesis is the dimerisation of the monomer (Scheme 1), which is controlled by an enzyme with some similarities to ketosteroid isomerases (KSI-like). *In vivo* experiments have demonstrated that this enzyme is solely responsible for the dimerisation reaction,^10,11^ with some evidence suggesting that the PEBP protein assists in this process.^10,11^ Experiments by Cox and co-workers^19^ using cell free yeast extracts show that either of the pair of KSI-like enzymes from the byssochlamic acid **1** BGC (BfL6 and BfL10) can catalyse the dimerisation of the maleidride monomer, although these experiments were very low yielding. Introduction of the two PEBPs from the byssochlamic acid BGC (BfL5 and BfL9) to the cell free extract experiments does not appear to significantly increase titre, but it is hard to determine their impact due to the general low yield of these experiments, and questions as to whether the PEBP is active under the conditions tested.^19^

All the identified known and putative maleidride BGCs encode at least one (sometimes two) KSI-like proteins. These enzymes contain an NTF2 domain (nuclear transport factor 2 - IPR032710), which categorises them within the NTF2-like superfamily. This is a large group of proteins which share a common fold, which consists of a cone-like shape with a cavity inside that can be adapted to serve a range of functions.^53^ Various enzymatic activities are catalysed by this family, including as a polyketide cyclase, dehydratase, epoxide hydrolase and ketosteroid isomerase.^53^

All the current evidence demonstrates that the KSI-like enzymes are sufficient to catalyse dimerisation of maleic anhydride monomers to produce the dimerised maleidride core structures.^10,11,15,19^ Thus we propose that this new family of enzymes be renamed as ‘maleidride dimerising cyclases’ (MDCs). All identified MDCs are categorised in the currently unnamed subfamily PTHR31779:SF6. To date, this subfamily contains 133 proteins, all of which are fungal. According to Interpro, none of these sequences have been linked to specific BGCs or to a specific function. It is interesting to note that a phylogenetic analysis of the MDCs (Figure S52) does not clade according to mode of dimerisation. Although the MDCs appear to catalyse the regiochemical dimerisation reaction, at present how the specific dimerisation mode is controlled remains cryptic.

Interpro analysis shows that the known and putative maleidride PEBP sequences all contain a phosphatidylethanolamine-binding protein domain (IPR008914), however phylogenetic analysis shows they cluster into two distinct groups (Figure S53). All verified and most putative maleidride clusters that contain PEBPs have at least one PEBP for which we propose the name: ‘maleidride PEBP type 1’. Type 1 PEBPs appear to cluster most closely to other diverse characterised eukaryotic PEBPs (Figure S53), including a PEBP identified from the bovine brain^54^ and various PEBPs from plants involved in the promotion and suppression of flowering and control of plant architecture.^55,56^ Several other clusters contain a second different PEBP, which form a separate clade that are distantly related to prokaryotic, eukaryotic and maleidride type 1 PEBPs, for which we propose the name: ‘maleidride PEBP type 2’ (Figure S53), the function of these proteins is currently unknown.

### 2.3. Common maleidride enzymes

Many enzymes are common across maleidride BGCs, including those that could function early in the biosynthetic pathways, such as the AMP-CoA ligases and enoyl CoA isomerases, as well as those that are likely to encode tailoring steps. Tailoring of the core alicyclic ring increases the diversity of maleidride structures beyond their mode of dimerisation, and is important for the bioactivities of the mature compounds.

#### 2.3.1. AMP-CoA ligase

Interpro analysis of the AMP CoA ligases from putative and known maleidride BGCs shows that they all contain the AMP-dependent synthetase/ligase domain IPR000873, as well as the C-terminal domain IPR025110 which is found associated with IPR000873. This places these enzymes in the ANL family, which are known to catalyse the activation of a carboxylate substrate with ATP to form an acyl adenylate intermediate, which can then be used in subsequent diverse partial reactions, most commonly the formation of a thiolester.^57^ Interpro analysis also shows that all the maleidride AMP CoA ligases fall within the family PTHR24096 (long-chain fatty acid CoA ligase), and in the currently unnamed subgroup PTHR24096:SF312. Characterised enzymes belonging to subgroup SF312 are mostly involved in the biosynthesis of cyclic or linear lipopeptides, activating the acyl moiety to form an acyl-AMP prior to transfer to the thiolation domain of the NRPS.^58^

Of the confirmed maleidride BGCs, only the rubratoxin (e.g. **8**) cluster contains an AMP CoA ligase (Table S1), and to date no molecular studies have been conducted for this gene. Of the newly identified BGCs, the likely epiheveadride **21** BGC of *W. aquatica*, the orthologous clusters from *Bipolaris* species, and the uncharacterised BGCs from *Cadophora* sp., *C. zeae-maydis, T. atroroseus* and *O. maius Zn* are all predicted to encode AMP CoA ligases, which collectively demonstrate 47 – 75 % protein identity (Table S10). Cox and co-workers^19^ postulated that AMP-CoA ligases would be required to activate the polyketide released by the PKS as a carboxylic acid, to form a CoA thiolester, since their *in vitro* assays with the byssochlamic acid **1** alkylcitrate synthase (ACS) show it will only accept a CoA substrate.^19^ As discussed in section 2.2.3.1, this is consistent with their relationship to primary metabolism enzymes which require CoA thiolesters as substrates. Cox and colleagues proposed that clusters without an AMP-CoA ligase could utilise an endogenous enzyme from primary metabolism.^19^

*In vitro* assays could be used to conclusively determine whether the AMP CoA ligases which are common within the maleidride clusters do indeed activate the products of the maleidride polyketide synthases prior to the action of the maleic acid forming ACS.

#### 2.3.2. Isochorismatase

Genes encoding isochorismatase-like (ICM-like) enzymes are present in the confirmed rubratoxin (e.g. **8**), scytalidin **15** and zopfiellin **4** BGCs (Table S1), and can now be identified in the likely maleidride BGCs of *T. atroroseus, T. funiculosus, O. maius Zn, Cadophora* sp. (Table S2) as well as *C. zeae-maydis*, which contains two predicted ICM-like enzymes. Although the level of sequence similarity varies significantly, with some demonstrating less than 25 % protein identity with each other (see Table S11), Interpro analyses shows that they all fall into the isochorismatase-like superfamily (IPR036380), many members of which are hydrolases. At present the role of the isochorismatases is enigmatic, although interestingly an I-Tasser analysis conducted for a range of the isochorismatases, identified maleic acid as a potential ligand (Table S19).

#### 2.3.3. Enoyl CoA Isomerase

Putative enoyl CoA isomerases are present in 50 % of maleidride BGCs (excluding orthologous BGCs) including those for byssochlamic acid **1**, scytalidin **15** and the zopfiellin **4**. Annotation of the *Z. curvata* zopfiellin BGC by Shiina *et al*.^15^ did not initially identify an enoyl CoA isomerase, but work on the orthologous BGC from *D. curvata* by de Mattos-Shipley *et al*.^12^ confirmed the presence of such a gene *via* RNAseq analysis. Re-interrogating the *Z. curvata* sequence can now identify a homologous enoyl CoA isomerase at the same locus, which we name ZopS (see ESI).

Of the newly identified BGCs, those from *A. glaucus, C. crateriformis, I. aleurites, T. borbonicus, T. stipitatus* BGC1 and the epiheveadride (e.g. **21**) BGC from *W. aquatica* contain an enoyl CoA isomerase (Tables S1 and S2). Percentage identity between these enzymes ranges from 15.91 - 62.06 % (excluding orthologues) (Table S12), and Interpro analysis shows that they all fall into the ClpP/crotonase-like domain superfamily (IPR029045), with the exception of Ts1R1. Ts1R1 is a newly identified protein (see Table S2), which contains the CATH / Gene3D domain for 2-enoyl-CoA hydratases (3.90.226.10), a domain which is also present in all other maleidride enoyl CoA isomerases, therefore we conclude that it is a related enzyme. The common theme uniting the ClpP/crotonase-like superfamily is the need to stabilize an enolate anion intermediate derived from an acyl-CoA substrate.^59^ A top hit on the Swissprot database for most of the maleidride enoyl CoA isomerases is the characterised Δ^3,5^-Δ^2,4^-dienoyl-CoA isomerase (At5g43280) from *Arabidopsis thaliana* (Scheme 2).^60^

**Scheme 2:**
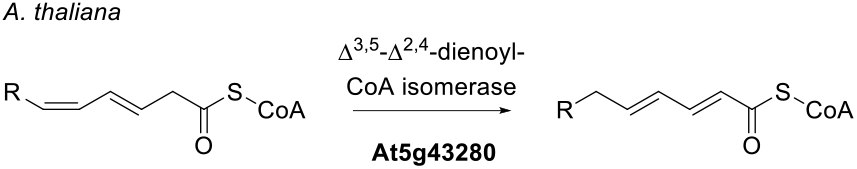
Reaction catalysed by the *A. thaliana* Δ^3,5^ -Δ^2,4^ -dienoyl-CoA isomerase.

At present, no molecular studies have been conducted on the maleidride enoyl CoA isomerases. Despite the presence of an enoyl CoA isomerase in the byssochlamic acid **1** BGC (BfL11), it was not considered in the original heterologous expression experiment utilising *A. oryzae* as a host organism.^10^ Both byssochlamic acid **1** and the agnestadrides (e.g. **6**) were produced under heterologous induction conditions, which could suggest that the enoyl CoA isomerase is not necessary for biosynthesis,^10^ although a homologue to the enoyl CoA isomerases is present within the *A. oryzae* genome (BAE65732.1 which ranges from 16.44 - 47.08 % sequence identity to maleidride enoyl CoA isomerases).

A likely scenario is that the enoyl CoA isomerases act on the acyl-CoA intermediates prior to the action of the alkylcitrate synthases (ACSs; see Section 2.2.1.3). *In vitro* studies by Cox and colleagues^19^ showed that the ACS from the byssochlamic acid **1** BGC produced exclusively 3,4-*anti* products (e.g. **25**), with evidence suggesting that **25** is the 3*S*,4*R* stereoisomer. In some other citrate based natural products, e.g. squalestatin, the product of the citrate synthase is the 3*S*,4*S* stereoisomer. The authors suggested that differences in the configuration at the 4-position of **25** must be controlled by the geometry of the enoyl CoA substrate of the ACS.^19^ This could suggest a role for the maleidride enoyl CoA isomerases in controlling the geometry of the various acyl-CoA substrates, providing the relevant ACS with the correct substrate to set the downstream stereochemistry.

**Scheme 3:**
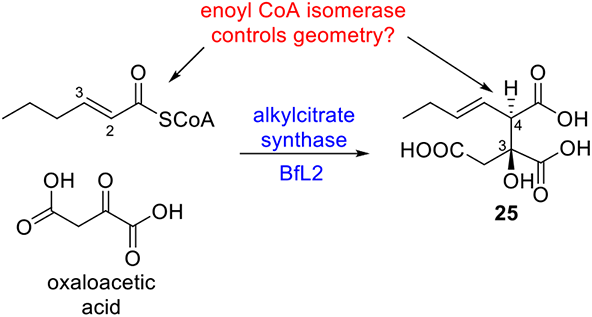
BfL2 produces a 3,4-*anti* product from enoyl CoA and oxaloacetate. The enoyl CoA isomerase may control the geometry of the enoyl CoA, and therefore ultimately control differences in the configuration at the 4-position of **25**.

#### 2.3.4. Maleidride conserved protein

Amongst the identified maleidride BGCs, 44 % (excluding orthologous BGCs) contain a protein which is conserved across these BGCs, for which we suggest the name ‘maleidride conserved protein’. Identity between the maleidride conserved proteins ranges from 22.99 - 55.91% (excluding orthologues) (Table S13). BLASTp analysis of these sequences has not revealed hits to any characterised proteins. However, PSI-BLAST has been demonstrated to be useful in detecting remote homologues *via* sequence searches.^61^ PSI-BLAST analysis of the maleidride conserved proteins provided a hit to AlnI, a protein identified from the asperlin BGC (between 13.71 - 23.04% identity – Table S13), which has been shown to be essential to biosynthesis.^44^ At present, no molecular studies have been conducted on the maleidride conserved proteins, however, there is a maleidride conserved protein (BfL8, ANF07280.1) within the byssochlamic acid BGC, which was not considered in the *A. oryzae* heterologous production study.^10^ Both byssochlamic acid **1** and the agnestadrides (e.g. **6**) were synthesised by the host organism, which could suggest that the maleidride conserved protein is not essential for biosynthesis, although a homologue which might complement the activity of BfL8 is present within the *A. oryzae* genome (BAE60519.1), and has 36% identity to BfL8.

#### 2.3.5. AusD-like methyl transferase

Amongst the maleidride BGCs linked to the biosynthesis of specific compounds (Figure 4 and Table S1), only the phomoidride (e.g. **16**) BGC contains an AusD-like methyl transferase (PTHR35897), although 5 other homologues are found within the orphan maleidride BGCs (Figure 5 and Table S2). All 6 maleidride methyl transferases contain the S-adenosyl-L-methionine-dependent methyltransferase domain IPR029063. AusD and other known homologues are involved in the biosynthesis of meroterpenoids.^62-66^ Those AusD-like enzymes which have been characterised perform a methyl esterification reaction.^62,66^ A prediction for the role of these enzymes within maleidride biosynthetic pathways is difficult to determine, as no methyl esterification appears to be required for phomoidride biosynthesis and the remaining maleidride AusD-like enzymes are from orphan BGCs.

#### 2.3.6. Aldo/keto reductase

Both the cornexistin **7** and epiheveadrides (e.g. **21**) BGCs contain putative aldo/keto reductases with the domain PTHR11732:SF461 (PvL15 - ASK38702.1 and EpiR14). This family of enzymes are known to catalyse the reversible NAD(P)H-dependent reduction of a carbonyl-containing compound to the corresponding alcohol.^67^ *W. aquatica* is known to produce a range of epiheveadrides (**18**-**23**), including those where the carbonyls of the anhydrides are reduced. Therefore we speculate that EpiR14 is involved in catalysing this reduction. There is no evidence for reduction of the single remaining maleic anhydride moiety present in cornexistin **7**, however it is possible that PvL15 may be involved in the removal of the second maleic anhydride ring. Interestingly, one of the maleic anhydride moieties in the rubratoxin pathway is also partially reduced to form rubratoxin A **8**. In this case, an unusual multi-domain ferric reductase (RbtH) catalyses the reduction of the C-X carbonyl.^14^ No other identified maleidride BGCs contain a ferric reductase. A reduced form of byssochlamic acid **1**, named dihydrobyssochlamic acid, has been isolated as a major product of *B. fulva*, but no enzymes which could catalyse such a reduction have been identified.^4^

#### 2.3.7. Quinone reductase

The BGC from *T. atroroseus* likely to encode glauconic acid **2** biosynthesis and the orphan BGC from *Cadophora* sp. both contain putative quinone reductases with the domain PTHR32332:SF28 (Table S2). An alignment with the known NADH dependent quinone reductase PA1024 (Q9I4V0.1)^68^ shows high homology, with 51.99 - 52.91 % sequence identity (Table S14). Six motifs which define this class of enzyme are almost completely conserved within the maleidride sequences (Figure S45).^68^ Inspection of the structure of glauconic acid **2** does not reveal an obvious activity for a quinone reductase within this pathway.

#### 2.3.8. Cytochrome P450

Cytochrome P450s (CYPs) are very common in fungal natural product pathways,^69,70^ however, only 33 % of maleidride BGCs contain a CYP (excluding orthologous BGCs). In general CYP enzymes have low sequence identity, but show structural similarities, and have four characteristic conserved motifs, involved in heme and oxygen-binding.^69^ Two P450 enzymes from maleidride clusters have been investigated – RbtI from the rubratoxin (e.g. **8**) BGC, and PvL13 from the cornexistin **7** BGC. Some evidence points towards RbtI being involved in terminal hydroxylation of one of the rubratoxin monomers prior to dimerisation.^14^ Phylogenetic analysis supports this role, as TsRbtI (EED18842.1) clades with other CYPs that perform similar functions (Figure S54).^71,72^ TfR1, the CYP from the semi-orthologous *T. funiculosus* BGC also falls within this clade.

No other maleidride P450 falls within the same clade as the CYP PvL13 (Figure S54) which is involved in late-stage oxidation of cornexistin **7**, however the structure of cornexistin **7** differs significantly from all other known maleidrides, in that it only contains one maleic anhydride ring, therefore it is unsurprising that late-stage enzymes in this pathway are distinct from other common maleidride enzymes.

The remaining maleidride CYPs are all from orphan BGCs and cluster as a single clade (Figure S54). Characterised similar CYPs have varied roles in sesquiterpenoid, sesterterpenoid, cyclohexadepsipeptide and indole-tetramic acid biosynthesis, therefore the function of the orphan maleidride CYPs are hard to predict.

#### 2.3.9. α-ketoglutarate dependent dioxygenases

The α-ketoglutarate dependent dioxygenases (αKGDDs) are versatile enzymes that catalyse various C-H bond activation reactions.^73^ αKGDDs are common in maleidride BGCs, with many contributing to unique features of specific maleidrides. Most prevalent are the AsaB-like αKGDDs (IPR044053), with 44 % of known and putative maleidride BGCs containing at least one (excluding orthologous BGCs). TauD-like αKGDDs (IPR042098) are found in 22 % (excluding orthologous BGCs). A single IPNS-like αKGDD (IPR027443) is found within the cornexistin cluster, which is responsible for a late-stage central ring oxidation in the biosynthesis of cornexistin **7**.^11^ A single PhyH-like αKGDD (IPR008775) is found within the BGC from *T. atroroseus*, a species which has putatively been linked to glauconic acid **2** biosynthesis, although the encoded gene is located towards the end of the predicted BGC, so whether it is part of the BGC or not is unknown.^29^

The two AsaB-like αKGDDs from the rubratoxin BGC, RbtB and RbtG, perform hydroxylations of an alkyl group attached to the central ring structure of the rubratoxins (e.g. **8**).^14^ Additionally, there are two TauD-like αKGDDs, RbtE and RbtU, which also undertake oxidative modifications involved in the production of the mature rubratoxins (e.g. **8**) including hydroxylation of the alicyclic ring.^14^ ScyL2 is an AsaB-like αKGDD which hydroxylates the central ring structure to produce the mature nonadride scytalidin **15**.^12^ ZopK/ZopL9 is a multifunctional AsaB-like αKGDD that both hydroxylates the nonadride precursor to zopfiellin **4**, and performs the oxidative ring contraction to produce the 8-membered central ring structure of the octadride, zopfiellin **4**. ^12,15^

Phylogenetic analysis of maleidride αKGDDs with other characterised fungal αKGDDs places each type of αKGDD into a separate clade (Figure 9). Within the orphan maleidride BGCs, *C. zeae-maydis, C. crateriformis, T. atroroseus* and the *T. stipitatus* BGC 1 all contain AsaB-like αKGDDs. TauD-like αKGDDs have been identified within the orphan BGCs from *Cadophora* sp., *O. maius Zn* and *T. funiculosus*. The TauD-like αKGDD from *T. funiculosus* has 77.05 % identity with TsRbtE, suggesting it could perform the same central ring hydroxylation (Table S16).^14^ Due to the versatile nature of these enzymes, and the range of reactions catalysed even within the characterised maleidride αKGDDs, it is difficult to predict the catalytic function for the remainder of the orphan and uncharacterised αKGDDs.

**Figure 9:**
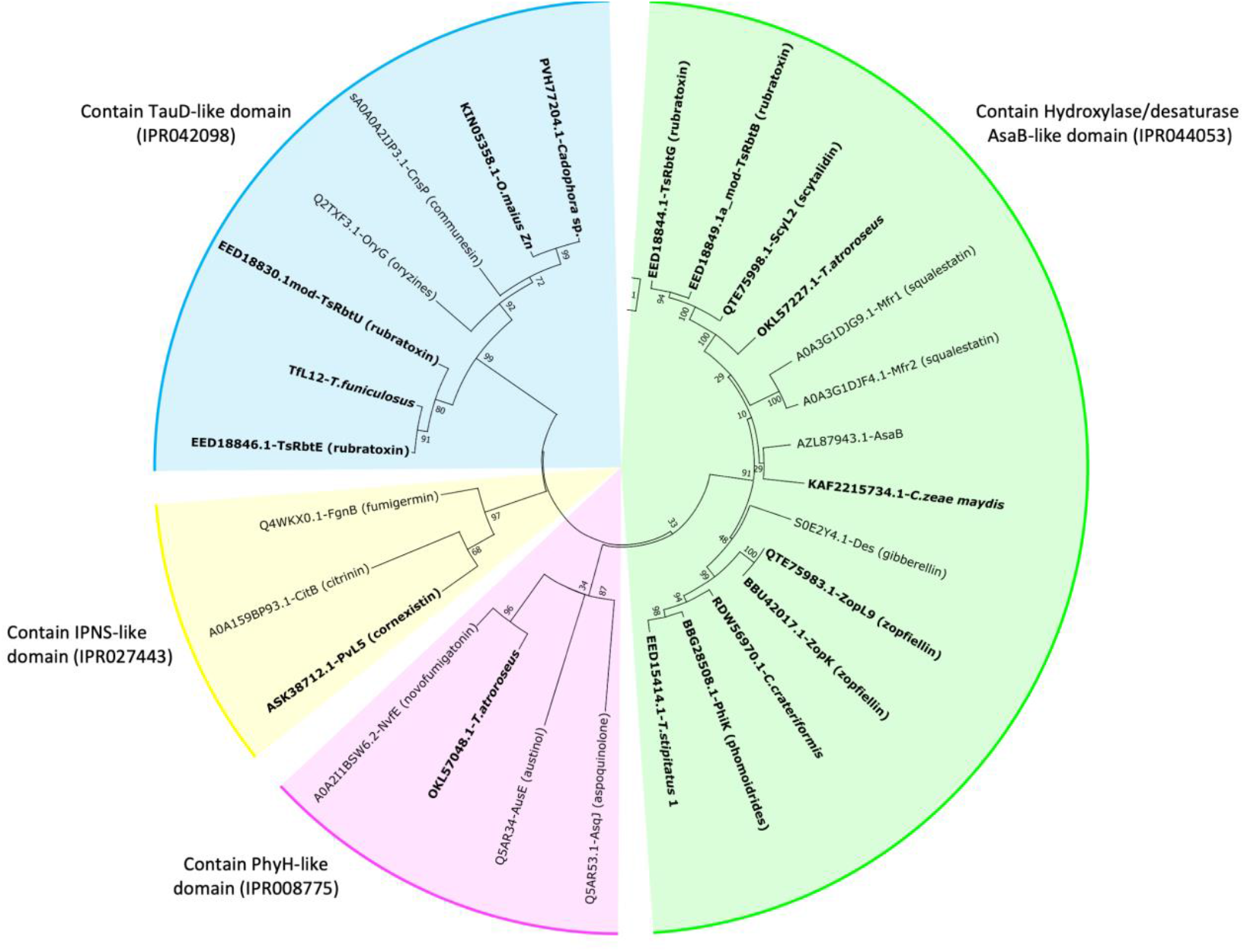
An unrooted phylogenetic analysis of α-ketoglutarate dependent dioxygenases (αKGDDs), present within maleidride BGCs (highlighted in bold). A number of known αKGDDs from other fungal natural product pathways were included in this analysis. The evolutionary history was inferred by using the Maximum Likelihood method and Le_Gascuel_2008 model.^36^ The tree with the highest log likelihood (−12257.60) is shown. A discrete Gamma distribution was used to model evolutionary rate differences among sites (5 categories (+*G*, parameter = 10.2669)). The rate variation model allowed for some sites to be evolutionarily invariable ([+I], 0.00% sites). The tree is drawn to scale, with branch lengths measured in the number of substitutions per site. This analysis involved 28 amino acid sequences. There were a total of 206 positions in the final dataset. Evolutionary analyses were conducted in MEGA X.^37^

All of the proteins included in this analysis were confirmed as containing the conserved “facial triad” motif comprised of two His and one Asp or Glu residue (HXD/EXnH) (see Figures S46 – S49). This triad is responsible for chelating the free iron required for the function of this superfamily of enzymes.^74^ It is worth noting that there are multiple different annotations of AsaB from *Aspergillus flavus* available on the NCBI database. The protein used in this analysis (AZL87943.1) contains the aforementioned conserved motif. However, the AsaB protein from the SwissProt database (B8N0E7.1) is shorter and is missing the conserved motif (see Figure S46). It is therefore likely that the B8N0E7.1 annotation is incorrect, and should be treated with caution.

All of the proteins in this analysis also contain the invariant arginine residue responsible for binding α-ketoglutarate (Figures S46-S49).^75^ PhyH-like αKGDDs are known to contain a conserved glutamine residue, which is also thought to be involved in binding αKG.^76^ As can be seen in Figure S49, this residue is present in all PhyH-like proteins analysed during this study, with the exception of NvfE from the novofumigatonin pathway, which is known to be an unusual, cofactor free, isomerase.^62^

#### 2.3.10. Other enzymes

The cornexistin **7** BGC encodes enzymes which are predicted to be involved in maleic anhydride ring removal, including the dienelactone hydrolase (PvL11) and 6-bladed β-propeller-like protein (PvL7).^11^ Interestingly the orphan maleidride BGCs identified from the lichen fungus, *I. aleurites*, and from *T. borbonicus* both contain homologues to PvL7 and PvL11, which could suggest that these BGCs may encode for similar single maleic anhydride maleidrides (Figure S39). Recent evidence suggests that a homologue to PvL7, Asr5 (A0A2U8U2M1.1) has hetero Diels–Alderase activity (Figure S50).^77^

The rubratoxins are heavily oxidized maleidrides which undergo a complex series of redox reactions to produce the mature compound, rubratoxin A **8**.^14^ The roles of several of the encoded enzymes within the rubratoxin BGC currently remain obscure, including RbtF which encodes a SnoaL-like domain and TsRbtB2, a thioesterase not identified in the original annotation. SnoaL itself is a polyketide cyclase, which catalyses ring closure *via* aldol condensation.^78^ Homologues to RbtF and TsRbtB2 are present in the *I. aleurites* and *T. funiculosus* BGCs respectively, suggesting that they could have a conserved function in different maleidride biosynthetic pathways.

## 3. Conclusions

Maleidride BGCs are distributed across a diverse range of fungi, all within the Pezizomycotina, but across five different classes within this subphylum. There appears to be evidence of horizontal gene transfer, for example, the closely related zopfiellin and scytalidin BGCs are found encoded within fungi from different classes. Many of the known maleidride producers, as well as fungi containing orphan maleidride BGCs, are associated with plants, either as pathogens,^79-82^ endophytes,^83-86^ or involved in fruit spoilage.^87,88^ This might provide a hint towards an evolutionary advantage for the acquisition of a maleidride BGC – several maleidrides are known to be phytotoxic, ^5,79,89^ and therefore might be virulence factors; whilst others are antifungal, ^2,3,90,91^ perhaps providing a defence for plant-fungal symbionts. Maleidrides form part of a biosynthetic continuum of alkyl citrate-based compounds isolated from fungi. These compounds range from those that are likely or known to be formed from an alkylcitrate synthase mediated condensation between a fatty acid and oxaloacetate, such as the alkylcitric acids,^45^ the oryzines, ^48^ and the sporothriolides;^49^ to those where a polyketide is condensed with oxaloacetate, such as squalestatin^50-52^ and the maleidrides e.g. byssochlamic acid **1**.^10^ Where characterised,^19^ the early steps towards alkyl citrate-based compounds appear to mirror those from primary metabolism, and suggests that these pathways probably evolved by the simple diversification of methylcitrate synthases to accept longer fatty acid/polyketide substrates, followed by the addition of various enzymes to diversify. The phylogenetic analysis of alkylcitrate synthases conducted here corroborates this hypothesis further (Figure 7).

An interesting result from our phylogenetic analyses concerns the potential prediction of chain length synthesised by maleidride PKSs (Figure 6, B). Those PKSs that have been linked to specific maleidride compounds, and therefore can be expected to produce specific polyketides, appear to clade together according to chain length. This allows for a tentative prediction of the chain length of polyketides synthesised by uncharacterised maleidride PKSs. To the best of our knowledge, this is unprecedented amongst the hrPKSs, where in general prediction of chain length remains cryptic.^92^ This might suggest that from an early prototype maleidride BGC, the hrPKS has diversified by changing its capacity to synthesise specific polyketide chain lengths, in a manner that is reflected by the sequence relatedness. This might be due to the very simple programming required for the production of all known maleidride PKSs – in most cases, only chain length differs, although the level of saturation is programmed in several, e.g. phomoidride B **16**. Linkage and characterisation of further maleidride BGCs could confirm this hypothesis, especially from the clade which currently contains no characterised PKSs.

One of the most intriguing aspects of maleidride biosynthesis – the key dimerisation reaction leading to diverse carbon frameworks - appears to be solely controlled by the maleidride dimerising cyclases. Phylogenetic analysis of these enzymes (Figure S52) does not appear to demonstrate any evolutionary relationship regarding mode of dimerisation, therefore, at present how dimerisation is controlled remains cryptic. Detailed *in vitro* and modelling studies of MDCs as well as the linkage of known or novel maleidrides to specific BGCs (therefore specifying mode of dimerisation) would provide further insights into these intriguing enzymes.

Despite increasing knowledge of characterised maleidride biosynthetic pathways, and our *in silico* analyses of both characterised and orphan BGCs, many questions remain to be answered regarding the biosynthesis of this important family of compounds. Currently, in common with most fungal natural product pathways, sequence data alone is not sufficient to allow for accurate predictions of chemical structure, however, any further orphan maleidride BGCs discovered could be subjected to similar analyses undertaken in this work, allowing for certain predictions to be made regarding the output encoded within the BGC.

## 4. Methods

### 4.1. Identifying maleidride BGCs

#### 4.1.1. cblaster

cblaster^24^ is a tool to rapidly detect co-located genes in local and remote databases. A set of query protein sequences are provided to cblaster, which can then search the NCBI database (or a local database) using the BLASTp algorithm. The genomic context of each hit is identified, and those sequences which are co-located (default is 20 kb) are returned. The user can define various filters, for example specific sequences can be set as required, minimum identity and minimum coverage can be altered (default is 30% and 70% respectively), furthermore the user can specify a taxonomic group of interest using an Entrez search query. To detect maleidride BGCs we provided cblaster with the core protein sequences from the *B. fulva* byssochlamic acid / agnestadrides BGC (Bfpks, BfL1 – hydrolase, BfL2 – ACS, BfL3 – ACDH, BfL5 – PEBP type 1, BfL6 – MDC, BfL10 – MDC) with a maximum distance between any two hits set at 50 kb. The PKS, ACS, ACDH and MDC were all set as required hits. Although the evidence suggests that the hydrolase is important for maleidride biosynthesis (see section 2.2.1.2 for reasoning), our experience has shown that it can often be missed by automatic annotations and was therefore not set as required. This was confirmed by the identification of two orphan BGCs which had no hits to the BfL1 hydrolase, although subsequent manual annotation of the sequences did identify homologues within these clusters. The type 1 PEBP, BfL5, was also not set as required, as current evidence suggests it holds an ancillary role, and is not absolutely required for maleidride biosynthesis. Three orphan BGCs were identified with no type 1 PEBP apparent within the cluster.

#### 4.1.2. MultiGeneBlast

The core *B. fulva* sequences used for the cblaster analysis were also used to search unannotated genomes from the subphylum Pezizomycotina in the NCBI database using tBLASTn. Where multiple hits suggested a possible maleidride BGC (without knowledge of genomic context) the unannotated genomes were downloaded.

MultiGeneBlast is a tool which performs homology searches on multigene modules.^30^ Raw nucleotide sequences can be used to create databases that can be queried with protein sequences using the tBLASTn algorithm. The downloaded genomes were used to create a tBLASTn databases using MultiGeneBlast, which were subsequently queried using the same core *B. fulva* sequences, specifying a maximum distance between genes to be 50 kb. Novel putative maleidride BGCs were identified from the resultant output. The newly assembled *W. aquatica* genome (see section 4.2), was also included in this analysis to identify the location of a likely epiheveadride BGC.

### 4.2. Genome sequencing

Genomic DNA was prepared from cultures of *Wicklowia aquatica* strain CBS125634, grown in static MEB (17 g/L malt extract and 3 g/L mycological peptone) cultures for 14 days. Fungal material was lyophilized and ground under liquid nitrogen before using the GenElute Plant Genomic DNA Miniprep kit (Sigma), according to the manufacturers instructions. Sequencing was performed using the Illumina MiSeq system with a 600-cycle (2 × 300 bp) kit, and assembled using Newbler v2.9. This resulted in an estimated genome size of 43.1 Mbp, a total of 540 scaffolds with an N50 of 147,393 bp, and a total combined scaffold length of 42.04 Mbp.

### 4.3. Annotation of maleidride BGCs

#### 4.3.1. Gene prediction

Putative maleidride BGCs were identified from the unannotated genomes of *T. borbonicus, T. funiculosus* and *W. aquatica*. The scaffold which contained the putative maleidride BGC, as identified by MultiGeneBlast (see section 4.1.2), was submitted to AUGUSTUS^93^ using their pre-trained species parameters for *Fusarium graminearum*. The gene structure parameters were set at ‘predict only complete genes’. The .gff output was combined with the specific scaffold .fasta file to create a .gbk file with gene structure predictions.

#### 4.3.2. Functional annotation

BLAST and Interpro were used to predict functions for each gene within a cluster (even where predictions had previously been made), as well as those genes up and downstream where possible, to determine the boundaries of the cluster. Homologues of core and common protein sequences were aligned using the multiple sequence alignment algorithm, MUSCLE^94^. Where obvious errors in intron/exon boundaries were identified, the boundaries were either manually edited, or submitted to FGENESH^95^ using an appropriate model species for gene finding parameters to obtain the likely correct sequence. Artemis Comparison Tool,^96^ which can be used to view tBLASTx pairwise comparisons between DNA sequences, was used to identify unannotated homologous genes within BGCs.

### 4.4. clinker

clinker^31^ is a tool to perform global alignments between protein sequences within multiple gene clusters, which can then be visualised as an interactive image using clustermap.js.^31^ All maleidride BGCs were compared using clinker,^31^ with minimum alignment sequence identity set to 25% except where otherwise stated.

### 4.5. Phylogenetics

Mega X^37^ was used for all phylogenetic analysis.^97^ Multiple sequence alignments between homologues were generated using MUSCLE^94^ except where otherwise stated. A best model for a maximum likelihood tree for the alignment was estimated using Mega X.^37^ The phylogenetic tree was constructed using the appropriate model with the recommended rates among sites. Gaps/missing data were treated with the partial deletion option: all positions with less than 95% site coverage were eliminated, i.e., fewer than 5% alignment gaps, missing data, and ambiguous bases were allowed at any position. Initial tree(s) for the heuristic search were obtained automatically by applying Neighbour-Join and BioNJ algorithms to a matrix of pairwise distances estimated using the JTT model, and then selecting the topology with superior log likelihood value. An estimation of the reliability of each tree was conducted using the bootstrap method with at least 250 replicates.

## Supporting information

ESI - In silico analyses of maleidride biosynthetic gene clusters

