## Supplementary material for "*In silico* analyses of maleidride biosynthetic gene clusters": ESI - In silico analyses of maleidride biosynthetic gene clusters

### Reannotation of sequences

#### Polyketide Synthases

Based on multiple sequence alignments, certain of the known and putative maleidride PKSs appear to have been misannotated. An alignment of the original and reannotated sequences with other maleidride PKSs is shown in Figure S1. The newly predicted protein sequences are shown in Figure S2 - Figure S4.

```
CLUSTAL multiple sequence alignment by MUSCLE (3.8)

KAF2215724.1(Czm)           ------------MPSLLSNGNTAQLSPSASAEG------LEADKMMPIAVVGLGFRGPKD
OJJ86349.1(Ag)              ---------------------------------------------MPIAVVGLGFRGPGD
PhiA_BBG28498.1             -----------MSPFM-----------------------EPQDEIMPIAVVGMSFRGPGD
EED15402.1(Ts1)             -----------MSPLIHDDQP------------------YSWNAAMPIAVVGIGFRGPGD
RDW56971.1(Cc)              -----------MAFSQGTDTNGDPPSP------------LACGDAMPIAVIGMGFVGPGD
TsRbtJ_EED18841.1(Ts2)      -----------MAVQE-----------------------LTEDASMPIAIVGIGFRGPGD
TfPKS                       -----------MTRQE-----------------------LTEDASMPIAIVGMGFRGPGD
PVH77205.1(CadPKS2)         -----------MAKLRNGD--------------------PADDATMPIAIVGMGFRGPGD
KIN05356.1(OmPKS1)          ---------------------------------------------MPIAIVGIGFRGPGD
TbPKS1                      -----------MGSYALSNTDG-----------------VVKNEVMPIAVVGIGFRGPGD
CAF9941811.1(Ia)            ------------------------------------------------------------
CAF9941811.1mod(Ia)         ------------MPNEHQSREG--------------------GEIMPIAVIGMGFRGPGD
ZopPKS_QTE75992.1           -----------MAPYLNGHHDEVLSLPSGSPTSNPA-LTISEDRSMPIAVVGMGMRGPAD
ZopA_BBU42026.1             -----------MAPYLNGHHDEVLSLPSGSPTSNPA-LTISEDRSMPIAVVGMGMRGPAD
ScyPKS_QTE76000.1           -----------MAPFLNGQHNGTGDAAHGSSVTNGI-SILSEDKSMPIAVVGMSMRGPAD
ScyPKS_QTE76000.1mod        -----------MAPFLNGQHNGTGDAAHGSSVTNGI-SILSEDKSMPIAVVGMSMRGPAD
PvPKS1_ASK38717.1           -----------MVENVSSPSSPRTSSPSGSCTPTSATSVGSDDKSMPIAVVGMSFRGPRD
OKL57046.1(Ta)              MTPLILNHADDDVPPLDKTFNGYSTLPNEYHHGCMNGNPLKDDKTMPIAIVGMSFRGPAD
OKL57046.1mod(Ta)           MTPLILNHADDDVPPLDKTFNGYSTLPNEYHHGCMNGNPLKDDKTMPIAIVGMSFRGPAD
PVH77199.1(CadPKS1)         ---------------------------------------MAKDKSMPIAVIGMSFRGPKD
KIN05352.1(OmPKS2)          -----------MTPSLDNIPEGSFSRPTSN---------MARDKNMPIAVVGMSFRGPKD
BfPKS1_ANF07288.1           --------MTCKDKKMEDCLENSGRMTHGVAD-------AMGDKTMPIAIVGMSFRGPGD
EpiPKS                      -----------MAPFLSQDVEGSNKSTSG----------TLEDKSMPIAIVGMSFRGPGT
XP_014073936.1(Bm_ATTC)     -----------MAPFLDDNLGASSGINHN----------GKEDGSMPIAIVGMSFRGPGS
EMD85570.1(Bm_C5)           -----------MAPFLDDNLGASSGINHN----------GKEDGSMPIAIVGMSFRGPGS
XP_007715650.1(BZ)          -----------MAPFLNGNLGTSNGIDQN----------GKADGSMPIAIVGMSFRGPGS
XP_014553689.1(Bv)          -----------MAPFLNGNLGTSNGIDQN----------GKADGSMPIAIVGMSFRGPGS

KAF2215724.1(Czm)           ATTPEGLWRMIERRREAWTTIPKERWNNDAFYHPDNNRHGT-------------------
OJJ86349.1(Ag)              ATDVESLYRMILEAREAWSPVPKSKWNNDAFYHPEANHNGT-------------------
PhiA_BBG28498.1             ATNVENLLKMVAEGRESRSKIPKHKWNHEAFYHPDPSRYGS-------------------
EED15402.1(Ts1)             ATNVENLFRMIAEGRESRIDIPKEKWNHEAFYHPDPSRFGT-------------------
RDW56971.1(Cc)              ATNVEKLWTMISEGRESWTEIPKERWNHEAFYHPDPNRYGT-------------------
TsRbtJ_EED18841.1(Ts2)      ATNVESLWKMISEGRESWSKIPKDRWNHDAFYHPDANRHGT-------------------
TfPKS                       ATNVENLWKMISEGRESWSKIPKNRWNHDAFYHPDANRHGT-------------------
PVH77205.1(CadPKS2)         ATNVENLWKMISEGRESWSEIPKERWNHKAFYHPDANRHGT-------------------
KIN05356.1(OmPKS1)          ATNVENLWKMISEGRESWSEIPKQRWNHEAFYHPDASRHGT-------------------
TbPKS1                      ATDVEKFWNMICEAREARTVVPKEKWNNEAFYHPDSNRNGT-------------------
CAF9941811.1(Ia)            ------------------------------------------------------------
CAF9941811.1mod(Ia)         ATNVENLWNMMLEAREAWSLVPKQKWNHGAFYHPDANRNGT-------------------
ZopPKS_QTE75992.1           ATNVEGLWKLVSEAREGWSKIPKERWNNDAFYHPDSNRNGT-------------------
ZopA_BBU42026.1             ATNVEGLWKLVSEAREGWSKIPKERWNNDAFYHPDSNRNGT-------------------
ScyPKS_QTE76000.1           ATSVEGLWKLVSEAREGRTSIPKERWNNDAFYHPDSNRHGT-------------------
ScyPKS_QTE76000.1mod        ATSVEGLWKLVSEAREGRTSIPKERWNNDAFYHPDSNRHGT-------------------
PvPKS1_ASK38717.1           AISVESLWRMISEGREGWSKIPKSRWNNDAFYHPDHSRHGT-------------------
OKL57046.1(Ta)              ATNVESLWEMICERREGWSEVPKERWNNDAFYHPDNTRHGTVSVEHFDPTKMKCNHSNTP
OKL57046.1mod(Ta)           ATNVESLWEMICERREGWSEVPKERWNNDAFYHPDNTRHGT-------------------
PVH77199.1(CadPKS1)         ATSVENLWKMISEQREGWSKIPKERWNNDAFYHVDNARHGT-------------------
KIN05352.1(OmPKS2)          ATSVENLWKMISERREGWSKIPMEKWNNGAFYHPDNARHGT-------------------
BfPKS1_ANF07288.1           ATNVENLWKMISEARESRSPIPQERWNNDAFYHPDHKHHGT-------------------
EpiPKS                      ASSVENLWKMICEKREARTTIPKSRWNNDAFYHPNFERHGT-------------------
XP_014073936.1(Bm_ATTC)     ANNIENLFKMICEKRESRTAIPKSRWNNKAFYHPNFQRHGS-------------------
EMD85570.1(Bm_C5)           ANNIENLFKMICEKRESRTAIPKSRWNNKAFYHPNFQRHGS-------------------
XP_007715650.1(BZ)          ANNIENLFKMICAKKESRTAIPKSRWNNKAFYHPNFQRHGS-------------------
XP_014553689.1(Bv)          ANNIENLFKMICAKKESRTAIPKSRWNNKAFYHPNFQRHGS-------------------
```

```
KAF2215724.1(Czm)           ---INVVGGHFLEEDLAHFDAPFFNLTSAEASVLDPQQRLLLEVTYEAFENAGIPLEKVF
OJJ86349.1(Ag)              ---SNVQAGHYFTGDLSRFDAPFFNMTHTEAEALDPQQRLLLECCYEGLENAGIPLGKAA
PhiA_BBG28498.1             ---HNVEYGHWFQQDVTRFDAPFFNMTAAEAAALDPQQRMLLECTYEAMENSGTHMPNFV
EED15402.1(Ts1)             ---HNVTGGHYFQQDVSRFDAPFFNMTAAEAAALDPQQRMLLECTYEAMENSGTKMHDFV
RDW56971.1(Cc)              ---YNVRGGHFFQQDLSRFDAPFFHMTAAEAAALDPQQRLLLECTYEAMENAGISMDQFC
TsRbtJ_EED18841.1(Ts2)      ---YNVTGGHFLQQDVSQWDAPFFQMSAAEAAALDPQQRLLLECTYEAMENSGTTMEQFC
TfPKS                       ---YNVTGGHFLQQDVSQWDAPFFQMSAAEAAALDPQQRFLLESTYEAMENSGITMEQFC
PVH77205.1(CadPKS2)         ---YNVTGGHFIKQDLSQWDAPFFQMSAAEAAALDPQQRLLLECTYEAMENSGTTMKDFC
KIN05356.1(OmPKS1)          ---YNVTGGHFLQQDLSKWDAPFFQMSAAEAAALDPQQRLLLECTYEAMENSGTTMEQFC
TbPKS1                      ---SNVLAGHYFKDDLAKFDAPFFNMTNAEAESLDPQQRLLLECTYEALENAGVPMDKAT
CAF9941811.1(Ia)            ------------------------------------------------------------
CAF9941811.1mod(Ia)         ---SNVKGLHTFTDDLSKFDAPFFSMTRAEAEALDPQQRLLLECCYEALENSGTPLERAM
ZopPKS_QTE75992.1           ---TNVTAGHFMTQDLACFDAPFFNMTNAEAAALDPQQRLLLECTYEALENAGTPLAEIQ
ZopA_BBU42026.1             ---TNVTAGHFMTQDLACFDAPFFNMTNAEAAALDPQQRLLLECTYEALENAGTPLAEIQ
ScyPKS_QTE76000.1           ---HNVTAGHFMTEDLSRFDAPFFNMTNAEAAALDPQQRLLLECTYEALENSGTPLSEVQ
ScyPKS_QTE76000.1mod        ---HNVTAGHFMTEDLSRFDAPFFNMTNAEAAALDPQQRLLLECTYEALENSGTPLSEVQ
PvPKS1_ASK38717.1           ---INVEGGHFLEEDLARFDAPFFNMTNAEAAALDPQQRLLLESTFEAVENAGIPLDKML
OKL57046.1(Ta)              PFKINVKGGHFFSEDLAHFDAPFFNMTNAEAAALDPQQRLLLEGTFEALENGGITLEKIM
OKL57046.1mod(Ta)           ---INVKGGHFFSEDLAHFDAPFFNMTNAEAAALDPQQRLLLEGTFEALENGGITLEKIM
PVH77199.1(CadPKS1)         ---INVEGGHFFEENLAHFDAPFFNMTNSEAAALDPQQRLLLEGAFEAFENGGIPLERIM
KIN05352.1(OmPKS2)          ---INVEGGHFLEEDLARFDAPFFNMTDSEAAALDPQQRLLLEGAFEALENAGIPLEKIM
BfPKS1_ANF07288.1           ---HNVKAGHFFKEDLSRFDAPFFSMTSTEAAALDPSQRLLLECTYEALENGGIPMEKIV
EpiPKS                      ---HNVEYGHFFEEDISKFDAPFFNMTSAEAAALDPAQRLLLECTYEALENGGIPLEKIV
XP_014073936.1(Bm_ATTC)     ---HNVEYGHFFHDDISKFDAPFFNMTKEEAAALDPAQRLLLESTYEALENGGIPLEKIV
EMD85570.1(Bm_C5)           ---HNVEYGHFFHDDISKFDAPFFNMTKEEAAALDPAQRLLLESTYEALENGGIPLEKIV
XP_007715650.1(BZ)          ---HNVEYGHFFQDDISKFDAPFFNMTREEAAALDPAQRLLLESTYEALENGGIPLEKIV
XP_014553689.1(Bv)          ---HNVEYGHFFQDDISKFDAPFFNMTREEAAALDPAQRLLLESTYEALENGGIPLEKIV

KAF2215724.1(Czm)           GSKTSCFVGSFSGDYTDMLVRDPDCVPMYQCTNAGQSRAMTANRVSYFFDLKGQSTTVDT
OJJ86349.1(Ag)              GSNTAVFVGSFCGDYTDVLLRDPDAMPLYQATSSGHSRAIISNRLSYFFDFRGPSATIDT
PhiA_BBG28498.1             GTETSVFVGSFCTDYADVLWRDPETVPMYQCTNAGHSRANTANRISYSYDLKGPSVTVDT
EED15402.1(Ts1)             GSNTSVFVGSFCADYADVLWRDPETVPMYQCTNAGHSRANTANRVSYIYDLKGPSVTVDT
RDW56971.1(Cc)              GTKTSVFAGAFCTDYTDILWRDPESTPMYQCTNAGNSRANLANRLSYFFDLRGASVSVDT
TsRbtJ_EED18841.1(Ts2)      GSDTSVFAGAFTTDYTDILWRDPESTPMYQCTNSGQCRSNIANRLSYFFDLHGQSVDVDT
TfPKS                       GSDTSVFAGAFTTDYTDILWRDPESTPMYQCTNSGQCRSNFANRLSYFFDLQGQSVDVDT
PVH77205.1(CadPKS2)         GSNTSVFAGAFCTDYTDILWRDPESTPMYQCTNASNTRSNLANRISYFFDLRGQSVAVDT
KIN05356.1(OmPKS1)          GSNTSVFAGAFCTDYTDILWRDPESTPMYQCTNSAHARSNLANRISYFFDLRGQSVAVDT
TbPKS1                      GSKTSVFVGSFCGDYTDIIMRDPETVPLYQATSSGHSRAIISNRLSYFFDFSGPSVTIDT
CAF9941811.1(Ia)            ------------------------------------------------------------
CAF9941811.1mod(Ia)         GTDTSVYVGSFCGDYTDLLLRDTENIPLYQATSSGHSRAIIANRLSYFFDFKGPSVTIDT
ZopPKS_QTE75992.1           GSKTSCFVGSFCGDYTDMLMRDPETVPMYQCTNSGHSRAILANRLSYFFNLQGPSVTIDT
ZopA_BBU42026.1             GSKTSCFVGSFCGDYTDMLMRDPETVPMYQCTNSGHSRAILANRLSYFFNLQGPSVTIDT
ScyPKS_QTE76000.1           GSKTSCFVGSFCGDYTDLLMRDPETVPI--------------------------------
ScyPKS_QTE76000.1mod        GSKTSCFVGSFCGDYTDLLMRDPETVPMYQCTNSGHSRAILANRVSYFFNLQGPSVTIDT
PvPKS1_ASK38717.1           GSKTSCFVGSFCGDYTDMLVRDPEAIPMYQCTNAGQSRAITANRVSYFFDLRGPSVTVDT
OKL57046.1(Ta)              GSKTSCFVGSFSGDYTDMLLRDPDSVPMYQCTNAGQSRAITANRISYFFDLKGPSVTVDT
OKL57046.1mod(Ta)           GSKTSCFVGSFSGDYTDMLLRDPDSVPMYQCTNAGQSRAITANRISYFFDLKGPSVTVDT
PVH77199.1(CadPKS1)         GTKTSCFVGSFSGDYTDMLLRDPDCVPMYQCTNAGQSRAMTANRVSYFFDLKGPSVTVDT
KIN05352.1(OmPKS2)          GSKTACYVGSFSGDYTDMLLRDPDCVPMYQCTNAGQSRAITANRVSYFFDLKGPSVTVDT
BfPKS1_ANF07288.1           GSKTSVFVGSFATDYTDLLLRDPETVPMYQCTNASQSRAMISNRLSYFFDLHGPCVTVDT
EpiPKS                      GTKTSVFVGSFATDYTDLLTRDPETVPMYQCTNSGQSRAMVANRLSYFFDLHGPSVTVDT
XP_014073936.1(Bm_ATTC)     GTKTSVFVGSFATDYTDLLTRDPESVPMYQCTNSGQSRAMISNRLSYFFDLHGPSVTVDT
EMD85570.1(Bm_C5)           GTKTSVFVGSFATDYTDLLTRDPESVPMYQCTNSGQSRAMISNRLSYFFDLHGPSVTVDT
XP_007715650.1(BZ)          GTKTSVFVGSFATDYTDLLTRDPESVPMYQCTNSGQSRAMVSNRLSYFFDLHGPSVTVDT
XP_014553689.1(Bv)          GTKTSVFVGSFATDYTDLLTRDPESVPMYQCTNSGQSRAMVSNRLSYFFDLHGPSVTVDT
```

```
KAF2215724.1(Czm)            ACSGSLVALHLACQSLRTGDAKVALAAGVNTVLSHEFASTMSMMRFLSPDGRCH-TFDEK
OJJ86349.1(Ag)               ACSASLVALHMACQSLRTGESEQAVVAGANVILSHEITIGMSRMRFLSPDGRCY-AFDER
PhiA_BBG28498.1              ACSASLVALHLGCQSLLTGDAKQAIVAGSSAILSHEGMVTMSMMRLLSHEGRCY-TFDER
EED15402.1(Ts1)              ACSASLVALHLGCQSLRTGDAKQALVAGCSAILSHEGMVTMSMMRLLSPEGRCY-TFDER
RDW56971.1(Cc)               ACSTSLVGLHLGCQSLRTGDAKMALVAGASVILSHEAWVTMSMMRFLSPDGRCY-TFDER
TsRbtJ_EED18841.1(Ts2)       ACSTSLVGLHLGCQSLRTGEAKLSVVAGVNVILSHEAWVTMSMMRFLSPDGRCY-SFDER
TfPKS1                       ACSTSLIGLHLGCQTLRTGEAKMSIVAGVNVILSHEAWVTMSMMRFLSPDGRCY-SFDER
PVH77205.1(CadPKS2)          ACSTSLVGLHLGCQSLRAGESKLSIVAGANVILSHELWVTMSMMRFLSPEGRCY-TFDER
KIN05356.1(OmPKS1)           ACSTSLVGLHLGCQSLRTGESTLSIVAGANVILSHEVMVTMSMMRFLSPDGRCY-TFDER
TbPKS1                       ACSSSLVALHLACQSLRTGESEQAVVAGANVILSHEMTISMSMMRFLSPDGRCY-TFDDR
CAF9941811.1(Ia)             -------------------------------------------MMRFLSPDGRCY-TFDDR
CAF9941811.1mod(Ia)          ACSASLVALHLACQSLRTGESHQAVVAGANVILSHEITISMSMMRFLSPDGRCYFTFDDR
ZopPKS_QTE75992.1            ACSASLVALHLGCQSLRTGDATRAVVAGANVILSHEIMITMSMMRFMSPDGRCY-TFDDR
ZopA_BBU42026.1              ACSASLVALHLGCQSLRTGDATRAVVAGANVILSHEIMITMSMMRFMSPDGRCY-TFDDR
ScyPKS_QTE76000.1            --------------------------------------------FMSPDGRCY-TFDDR
ScyPKS_QTE76000.1mod         ACSASLVALHLGCQSLRTGDAKRAVVAGANVILSHEIMITMSMMRFMSPDGRCY-TFDDR
PvPKS1_ASK38717.1            ACSGSLVALHLACQSLRTGDAKMAIVSGVNTILSHEFMSTMSMMRFLSPDGRCY-TFDER
OKL57046.1(Ta)               ACSGSLVALHLACQSIRTGDAKTAIAAGVNTVLSHEFMSTMSMMRFLSPDGRCY-TFDER
OKL57046.1mod(Ta)            ACSGSLVALHLACQSIRTGDAKTAIAAGVNTVLSHEFMSTMSMMRFLSPDGRCY-TFDER
PVH77199.1(CadPKS1)          ACSGSLIALHLACQSLRTGEVKLAFAAGVNTILSHEFMTTMSMMKFLSPDGRCY-TFDER
KIN05352.1(OmPKS2)           ACSGSLVALHLACQSLRTGDAKLAFAAGVNTILSHEFMSTMSMMRFLSPDGRCY-TFDER
BfPKS1_ANF07288.1            ACSGSLVALHLGCRSLQTGDAKCSIVAGVNVILNHEFMITMSMMKFLSPDGRCY-TFDDR
EpiPKS                       ACSGSLVALHMACQSLRTGEAKCAIAAGVNVVLNHEFMITMSMMKFLSPDGRCY-TFDER
XP_014073936.1(Bm_ATTC)      ACSGSLVALHLACQSLRAGEAKSAIAAGVNVVLNHEFMTTMSMMKFLSPDGRCY-AFDER
EMD85570.1(Bm_C5)            ACSGSLVALHLACQSLRAGEAKSAIAAGVNVVLNHEFMTTMSMMKFLSPDGRCY-AFDER
XP_007715650.1(Bz)           ACSGSLVALHLACQSLRTGEANSAIAAGVNVVLNHEFMTTMSMMKFLSPDGRCY-AFDER
XP_014553689.1(Bv)           ACSGSLVALHLACQSLRTGEANSAIAAGVNVVLNHEFMTTMSMMKFLSPDGRCY-AFDER
                                                                        ::* :***: :**:.
```

Figure S1: Multiple sequence alignment using MUSCLE[1] of various known and predicted maleidride PKSs. An asterisk (*) marks completely conserved amino acid residues, whereas a colon (:) identifies highly conserved residues and a period (.) indicates moderately conserved residues. The region in red denotes a putative unidentified intron in sequence OKL57046.1 (*Talaromyces atroroseus*), which when removed (new sequence named OKL57046.1mod), aligns better with other maleidride PKSs. The region in blue highlights a region of the ScyPKS annotation of *Scytalidium album* (accession number QTE76000.1) which appears to have incorrectly identified the boundaries of an intron. When this intron was shortened, the newly predicted protein sequence (named QTE76000.1mod) aligns well with other maleidride PKSs. The regions in pink compare the published protein sequence for *Imshaugia aleurites* CAF9941811.1, with the newly annotated CAF9941811.1mod (annotated using FGENESH[2]). The published annotation appears to have missed multiple 5' exons, significantly truncating the predicted protein at the N terminus. See Figure S2Figure S4for the protein sequences of the newly annotated PKSs.

```
>OKL57046.1mod
MTPLILNHADDDVPPLDKTFNGYSTLPNEYHHGCMNGNPLKDDKTMPIAIVGMSFRGPADATNVESLWEMICERREGWSEVPKERWNNDAFYHPDNTRH
GTINVKGGHFFSEDLAHFDAPFFNMTNAEAAALDPQQRLLLEGTFEALENGGITLEKIMGSKTSCFVGSFSGDYTDMLLRDPDSVPMYQCTNAGQSRAI
TANRISYFFDLKGPSVTVDTACSGSLVALHLACQSIRTGDAKTAIAAGVNTVLSHEFMSTMSMMRFLSPDGRCYTFDERANGYARGEGVGCLILKPLKD
ALRDNDTIRAVIRGSGSNQDGKTSGITLPSGAAQEELVRNVYEAAGLDPLETEYVEAHGTGTQAGDPQETGALSRVFCPGRSSDKPLRVGSIKTNVGHL
EGASGIAGVIKATMMLENRMFLPNRNFETLNPRIPLDLWKLKVQLDIEPWETDGPHRVSVNSFGYGGSNAHVILEDAAGYLKSHEMKGYYRNPKAIAPR
DSKEHSNSNHTTDESSHSKRFTNAHSDNILNGEKKEENGHSQRKQLLVLSSFDEASGKRQSDRLQKYLLARKHLANEEFMTNLAFTLNNRRTSFMWKVA
ISGSNVQEVAQTLSKGVKFSRAMKKPTLGFVFTGQGAQWCGMGKELLAAYPVFSDSINKIGAYLKSLGAPFDVKEEITRDPSGSQINLALYSQPMCSAV
QIAIVDLLYSWGIKPASVTGHSSGEIAAAYTAGVLSMEDAMAVAYYRGVASTLMLTISQTQGAMMAVGLSKEDAEPYLSKLQFGKAVIACVNSPSSITI
SGDVLAIDELKGILDDQKIFARKLAVEVAYHSHHMQLVSEEYYQLISKINTKIAADEITESVEFFSSVTGSKAVASELGPNYWVKNMLGQVKFADSVRQ
LCLETTDPTNRTKGAGRKTRKRAGAATKASVDHIIEIGPHSALAGPIKQIIKANETLSSASIAYSSALVRKVNAVSSILDLAGNLVMAGWPLDLIALNS
PHGVGTNYERQALVDLPPYPWNHSNTYWAEPRLSKVYRNRKFPRTDLLGVLDRYSSPLEPRWRNHIRTTEIPWVNDHKIQSNIVYPAAGYIVMAIEAMH
QWMPEHYPEGTISGYTLREVNIGAALVISEQSAVEVMISLRPYRVSARGVSKVWREFSILSVTEENKWTEHSSGLICAHFEADTSNNLKQDAEVLHQTL
AEIYAKCDMQVNVNEFYKHLRQLGLEYGETFANMTEAYSATDSCVAEITISDTAATMPMNFQYPFVIHPSTLDSMFHPIFVALSAERGLIQDPAVPIAV
DEVYVSNSLKTTPGHRFSVYVCTEKKDESNIVASIVAVDKEEDVSSHMEHVGLSIKGLTCRVLPREVGNGTDEERERTAYNIKWNADPELLSAAGIASM
CTSSQPSEDDLQQLRLYEKCSSVYVADAVSQLDSKDIPAEKPHIQKLWKLLKASNIKISVSDDEKAWITEQTKRSGPEGELLCTLGDNLLSILRDGIDP
WNIMMTENRLDAYLNDTSRLVRNYKIAAKYVRLLGNKNPQLSLLEIGAGTGEASLPILQALSEGNTSVWLKRYTFTDLNTDMFEVAQKKLSDWADLIKF
KEFDVAGSLEEQGFKSHSYDVVILGHGIHLAKSTDRMLKNIRNLLKDEGKFIFVDEVYQNESIERSLVSGTFSSLWEDDVEGKYRVNAYTEDDWHQALL
DARFSGMEVCLRDAASHGSSVMISKAVGDGFPSPALDILLITEDGDCGVSESSLLDHLKTAGARVETCRFNEAHPDGRTCIVLSDLSFPVLAKSDATAF
EIVKSLFLHSTGVLWVTRGGSGLTINPNASLITGFARTARAEADGANIVTLDLDGQTPLSPERAAETIASLFRHRFVTRANTTEQDVEYAERQGIISIP
RVTESIGLNRDLESILGRASPTTQALYQPGRKLRPIVATATDMNSMHFVDEPMQQLPDDYVQIEVRASGINKSDSLLARGQRPLGAECSGIVCAIGKSV
MDLSVGDRVLCLGSGTITNYHQDKESAFQKVPEDMSFEHAAALPAAYCTAYYVVYNLARIVKTDSVLIHNIAEPTGQAILELCNLIGARVFGTVSEASQ
KKYVVKQLPIPEENILYYCHTTFAKEIIRMTNKKGVDVLVNCLDGDTEMRRFSWSCVASYGRFIDLGSRGIADNSRLEMGNFAKNSLFASFDLLSLLKE
KTSVAQKVWADVMCLFRTKAIRGFSSLLVHDVSDIGKALTEMESGGRLGKTVIVAKPGSVVKALPRDKSGELLRNDMSYLLVGGLGGIGRATASWMIDR
GAKFIIFANRSGLSREESKDTIRQLEAKGAKVAVYSCDIIDESDVIGMVKSASREMPPIKGVIQAAMVLRDTLIENMSFEEFSASLKPKFNGTWNLHNH
LPKNMDFFVMLSSISGVIGNASQAAYAAGNTFMDAFAGFRNSLGLPAVALDLGVITGVGYLSQNTELLAAMERQGFQGTDERTLMALIQTAISQPHRQD
SDAQIVTGLGSWKDGKSLGNFDQAIFSHFRRQFSGGENSSEEGTSADQLKENLRACKTLEEAASVICAALIEHIAARLETPAENINSSKSLSDYSIDSL
VAVEIRTWIAKEMSSTIPILELLASSSLLQLSEKIATRSTLVKVPEMSS
```

Figure S2: Modified sequence for the PKS from the *T. atroroseus* putative maleidride BGC, OKL57046.1mod.

```
>QTE76000.1mod
MAPFLNGQHNGTGDAAHGSSVTNGISILSEDKSMPIAVVGMSMRGPADATSVEGLWKLVSEAREGRTSIPKERWNNDAFYHPDSNRHGTHNVTAGHFMT
EDLSRFDAPFFNMTNAEAAALDPQQRLLLECTYEALENSGTPLSEVQGSKTSCFVGSFCGDYTDLLMRDPETVPMYQCTNSGHSRAILANRVSYFFNLQ
GPSVTIDTACSASLVALHLGCQSLRTGDAKRAVVAGANVILSHEIMITMSMMRFMSPDGRCYTFDDRANGYARGEGVGCVILKPLEDALRDGDTIRAVI
RGTGSNQDGKTSGITLPSGSAQESLIRTVYKTAGLDPLETSYVEAHGTGTPAGDPLETGALSRVFCPDRSPDEPLRIGSIKTNVGHLEGASGIAGVIKT
ILMLENKTLLPNRNFKNPNPRIPFYDWKLKVNTLVEPWIVSGPRRASVNSFGYGGANAHAILESAEDYLSSRKLNGSFRTTKSVLSLELNPSANGHSNG
STNGVTNGHAAESANGHSNGSAKKPVQTERTRLFIVSGFDEATSKRQAQTLSNYLESRQDLDDDNYLDNLAYTLGERRTEFIWKAAVPASSKSDLSKAL
SGDIKFSKSNKKPTLGFVFTGQGAQWCGMGKELLEAYPVFRKTINKIGSYLTSLGAPFDVADELTKDPKISQIGLALYSQPLCSAVQIALVDLLASWGI
KPASVTGHSSGEIAAAYTMGALNLEDAMAVAYYRGLASSNMQKTGTVSGSMMAVGMSKEDALPYISGLTKGKATVACVNSPSSITVSGDVTAIDELHVI
LEEKKLFARKLAVEVAYHSHHMELVADEYNTSISNIKIQETGDVEFYSSVYGKRIDASELGASYWVANMLREVKFADSVRLLCLETSSGKKTRKRASTP
VVNIIVEIGPHSALAGPIKQILQADSKLKEAGISYVSPLIRKVDAVKTTLDLASKLLVGGYPVDLSAVNRPIGTESHSVLADLPPYPWNHANSYWAEPR
LSKVFRQRTSPRTDLLGALDRNANPLEPRWRNHIRESEIPWVKDHKIQSNVVYPAAGYIVMAIEAAFQRATEKSLTINGYKLREVSIGSALVIPEQSEG
VETLVTFKPHTDSIRAPSDLWDEFCVYSVTDDNRWTEHCRGLIMVQTPQRTVNIIDGDVQAVAEKKSYIEIIAETEDKCKIDVDVKEFYEQLAQLGIEY
GETFANMTKARAAHNSCIGTISIADTAAVMPMHFQFPFVIHPSTLDAIFHPIFVALAAEVGPLKDPAVPVFIEEIYVSHQISSKPGDELVVYAGTKKKD
DKYLMASMIVVDGDHPDGEPVLTISDLTCTTLAREVAVESGDEIKRVTYNFEWRADIDLLSINDASKLCAYPTPPTQERHQIRSLEQTAYYYMEWALSK
ISTDELPSMEPHFQKLYACMEKFVKDVQEEKLGVPTALWVAADQAERANLRHNIRDSGPEGHLICLMGRNIPAIMKKEIDPWTLLTEKQSLEAYFRDTP
RIARTYDAVAKYFYLLGHKNPHLSILQIGTGTGGATFPILKSLGGADGEIPRFQKYDFTDTSNILSEELKQKLAPWKDLIAFKELDLNNDPIAQGYTVE
SYDVILAAHTLRTSKSLHTALGNARRLLKPGGKLVILDVTRERMAPSLVFGTLPNWWAAEEEDRQASPILSEDAWQTALLTSHFSGLELVVWDTPDEPE
HQSSMMVATALHKETIKTPDVLVVAEEDDCGVSVTHLLERLADLKINFEVIPFAQAKPTGKVCIVLSELAKSILSDPSKEEFETTKDIFIESGGVLWVT
RGGILSPTDPNSNLVTGFARTSRAETGGTIITTLDLDSQKPLSPTSSAETIFSLFKSLFTLDHPSTNEIDMEYAERDGQLLIPRLIENGKLSKRILASS
RRPVPEPQPFHQPGRPLVMHVGQPGLLDTIHFIDDERMAQPLADDEVELKVKATGLNFKDVMMGLGQIESETLGIECAGVVTAIGKKVQGFAIGDRVST
YGFGTFSNFYREKWTALQKIPDDMSFEMGAALPITYGTAYYSVYHLARVEKSDSILVHAATGGLGQAIIELCQLIGAEIYATVGTKEKKELLMDLFKIP
EDHIFYSRDGSFAKGIMRMTGGKGADVIFNSLAGEALRITWDCIAPYGRFIELGARDLTVNTRLEMKNFIKNPMFAAFNFIYLVRAKREVADKVCADVM
DLFRNKLIKGPSPLHVHRISKVEEALRIMQTGKHMGKLVAVSEPDEIVQTIPRDTSKNLLRADASYLLVGGLGGIGRATALWMIEHGAKNLIFANRSGL
ASQEARDTIDALKSKGASAAVYSCDVSKSEALAELVAESSKSMPPIRGVIQGAMVLRDGLLEKMPLSDYTSVIRPKVQGTWNLHNLLPKDMDFFLMLSS
ISGIIGNAAQAAYAAGSTFMDAFATYRNSLDLPAITIDLGVITEVGYLAETNKELAAGMQRQGFEGTNKEKLLALIQSAIADPKREGRLSQVVTGLGMW
KEGESLATFDLPVFNHFRRQALKKENGAADAGARVRDTLKVAKTLEEAADKICAALIDKISSRSNIPVDNISQDNPMSDYGIDSLVAVEMRNWIVREMD
STMPILELLANQSLQQLSAKIAQRSRLVDLKVVEAEA
```

Figure S3: Modified sequence for the maleidride PKS from the *S. album* scytalidin BGC, QTE76000.1mod.

```
>CAF9941811.1mod
MPNEHQSREGGEIMPIAVIGMGFRGPGDATNVENLWNMMLEAREAWSLVPKQKWNHGAFYHPDANRNGTSNVKGLHTFTDDLSKFDAPFFSMTRAEAEA
LDPQQRLLLECCYEALENSGTPLERAMGTDTSVYVGSFCGDYTDLLLRDTENIPLYQATSSGHSRAIIANRLSYFFDFKGPSVTIDTACSASLVALHLA
CQSLRTGESHQAVVAGANVILSHEITISMSMMRFLSPDGRCYTFDDRANGYARGEGVGCILLKPLHDALRDGDTVRGVIRNTGVNQDGRTSGITLPSRQ
AQEDLIQTVYEKAGLDPLDTSYVECHGTGTPAGDPLEAAAISRVFGPGRSHEQPLHIGSVKTNIGHLEGASGVAGIIKSILMLENQTILPNRNFQNANK
QIPLHDWKLFVPTRPEKWKSQGPLRASINSFGYGGTNAHAVLEDARGYLISQDMPESTSRTSRVRRSQSKAEVNGNCPSTTNTNSVGDVHPACLVNGNS
DCQPTDLTSPKSMVNGYTMEEKDIRTTSRVRLFLLSSFDQNAGRSQAKLLRQYLVDRLLHTAEDQFLNDLAYTLGERRSQFAYKNILAAKSTPQLIERL
DDENLKFTESSSGKKALAYIFTGQGAQWYAMGRELMQTYPIFHNSLFRASVWLNILGAPWNLIDELSKDAETSQVGSVHLSQPLCTALQLALVELLASW
NIRPTAVTGHSSGEIAAAYTVGALTFEDAIAVAYHRGVVSNLVKKTRKVRGAMMAVGMTPEDAAPLIAGLTQGKASVACINSPSSITVSGDFPAITELE
DILKAKGTFARKLDVEVAYHSHHMESVAQEYLAALSEVRAQAVVADIEFYSSVTGQRAESAELGPSYWVENMVGQVKFAESLRQLCLATGQKNTKSRQR
GRDSTVNILVEIGPHSALAGPIKQILQADSKINHAKAIYQTALVRKLDAVETCLALASKILTAGYAVNFAACNRPASAQKPRVLVDLPPYAWNHSTSYW
AESRVSKAYRFRPYPRTDLLGVPEVSRDSPRPSWRNYVRASEIPWVKDHKVQDSAVYPAAGYLVMAIEAASQRASAKDTNVAQYRLREVTFGQALVIPE
QSGEVETLVQLRPHQEGTRLLSDTWDEFCILSVTDENRWTEHCRGMISVAKATGRNDIVQSGLQIAGHLQKITHMEALCQTSVDTTQFYQQLDSIGLHY
GPSFTNMKSARSAPNMCIGKIEIPDTAATMPMGFQYPFVIHPATADSIFHGLFAALSSSPGSLKDPLLPVFVKEMSVSSQISHDPGHELIAYSSTERKD
IRQITASMIVLDGESHNSEPVITIDGLTCTMLANDSQVQASSRSRQIAYCLDWRPDVDFISSDDVSILCNYLRPPPTELEAVRSLERAGFYFMENALKT
LDPNKIQNMLPYHKLLWACFESFVSTVREGRLGFSTAAWITADEAERAEHIKEVRASGAEGALLCQVGQNLPNIIARKVEALPIMVEQGRLDAYYRENA
RFDRNYRAATKYIDLLAHKNPYIKVLEIGAGTGGATLPIIEAIGGGDAELPRLAEWHFTDISSSFFDAAKEKLERWNHLVSYAKLDIETDPVKQGFENE
SYDVVVAANVLHATKSLHQTLINVRRLLKRGGRLVLIELTRERMTTSTIFGTLSGWWAGAEDNRQMGPTLTEEEWDRKLKQIGFSGLDAAVWDSLTEPE
HQGSMMVSRVIDNDPIAENLGVLLICDDSLPWNIRKHMTTRISDECATTVDVQTLTAAIPAGKLCIVLCEATRPFLSDPSPEEFEATKRIFASASGVMW
VTKGAKMSSQDPASNLITGLTRTVRSEFGSINIVLFDLDPDQPFPSTAAAESVLSLFLSHFGANAQLSDDTEFEFVHRNGRIMIPRMIPNAAVNRKIES
ANSELKPEPQTFQQPGRHLVMEIRTPGLLDTIYFVEDDRINDALLEDQVEIEVKATGFNFKDVMMAMGQVEVERLGLECAGIITGCGKSTKHVSVGDRV
SCFAFGAFSTRYRTDAISVQKLPDDMSFEKGACLPVIYCTAYHSVYNVANVQKGETVLIHAASGGLGQAAIELCQLVGAEIFATVGTREKKTYLMNRFG
IPEDHIFFSRDGSFAEGIMAMTKGVGMDVILNSVAGEMLRLTWECIAPFGRFIELGARDYTINTRLEMHKFARNVTFAVVNLVSLVRERPQAAAQVWSK
AMDLFRSKKVDGPSPITVYGISEIEKALRIMQSGKHMGKLVAVARPDEMVMAIPHMKTGNLLRPSASYLLVGGLGGLGRATALWMADQGAKNLIFASRS
GLAKQEARDLVKALKERGVTVAVQSCDVGDSSQLRNALAQTSYMPPIRGVIQGAMVLQDSLLEKMTLSDYAAAIKPKVDGTWNLHELLPKDMDFFVMLS
STSGIIGNASQANYAAGSTFLDAFCDYRRGLGLPAVTIDLGVILGVGYVAENQELAGKLDRQGFEGTTKEELMQLIEIAITSPPNKSQSGQIVTGIGTW
SESSHGAFASPMFSHFRRMALDSGHSANESNHQTGHAIRDQLRKARSLDDATQQICESMIAKVSSLSMTPVEDISETKPMSEYGMDSLVAVEMRNWLFK
EMDTTIPILELLANQSLLSLAAKIVKGCKLVDPAIVRGVGEE
```

Figure S4: Modified sequence for the PKS from the *I. aleurites* putative maleidride BGC, CAF9941811.1mod.

#### Hydrolases

PhiM (BBG28510.1 from the phomoidride BGC), when aligned with BfL1, is missing approximately 15 amino acids (Figure S5). Interrogation of phomoidride BGC sequence shows that the intron identified in the sequence can be removed without introducing any stop codons. The sequence of BfL1 has been verified by RNAseq, whereas no evidence for similar analyses was apparent for the phomoidride cluster,[3] therefore it is likely that the alternative sequence (PhiM_BBG28510.1mod) identified in Figure S6 is correct.

```
BfL1_ANF07287.1            MPQVKLLCLHGAGTSIQILQSQLAPLIRELQKDSTASFHFIEGEVECGPGPGIEGIFEGP
PhiM_ BBG28510.1mod        MPGLRILCLHGQGSNTEIMKAQLGPITKLLKQNGVATFEWLEGTIPTEAGPGVSGVFEGP
PhiM_BBG28510.1            MPGLRILCLHGQGSNTEIMKAQLGPITKLLKQNGVATFEWLEGTIPTEAGPGVSGVFE--
                           ** :.:***** *:. :*:::**.*: . *:::..*:* ::** :   .***:.*:**

BfL1_ANF07287.1            YFSFYKWPRTL-EDDDESLYDAYGYLSDVIEEEGPFDGVIGFSHGGTLAFGYLAHLAKTR
PhiM_ BBG28510.1mod        YYQYFSWPPTACESDEQSIREACEDLEDYLEENGPYDGILGFSQGSTLLAEFLCDFARRN
PhiM_BBG28510.1            --------------DEQSIREACEDLEDYLEENGPYDGILGFSQGSTLLAEFLCDFARRN
                                         *::*: :*   *.* :**:**:**::***:*.**   :*. :*. .

BfL1_ANF07287.1            PNDALPFRCAVFMNAPPPFRMDTYGNLIVEEG-LQDLVRIPTLHVVGKKDFIYNFSLKLH
PhiM_ BBG28510.1mod        PGNEPPCRCAIFMNGIPPYRMGDDEKPIIDYGLLEHFPSIPTLHVVGKKDFVYEYSTILQ
PhiM_BBG28510.1            PGNEPPCRCAIFMNGIPPYRMGDDEKPIIDYGLLEHFPSIPTLHVVGKKDFVYEYSTILQ
                           *.:  * ***:***. **:**.   : *:: * *: :  ************:*::*  *:

BfL1_ANF07287.1            ELCDSANSTLVLHEKGHEIPSDTKNVNRITKALRELCQVSAFL-
PhiM_BBG28510.1mod         ASTASAWATLIAHEKGHEISNDTRIVQKINSAFEQMSIRIALGC
PhiM_BBG28510.1            ASTASAWATLIAHEKGHEISNDTRIVQKINSAFEQMSIRIALGC
                               **.:**: *******..**. *:.*..*: ::.   *:
```

Figure S5: Multiple sequence alignment of BfL1, PhiM (BBG28510.1) and the deduced alternative PhiM sequence (BBG28510.1mod) using MUSCLE.[1,4] An asterisk (*) marks completely conserved amino acid residues, whereas a colon (:) identifies highly conserved residues and a period (.) indicates moderately conserved residues.

```
> PhiM_BBG28510.1mod
MPGLRILCLHGQGSNTEIMKAQLGPITKLLKQNGVATFEWLEGTIPTEAGPGVSGVFEGPYYQYFSWPPTACESDEQSIREACEDLEDYLEENGPYDGI
LGFSQGSTLLAEFLCDFARRNPGNEPPCRCAIFMNGIPPYRMGDDEKPIIDYGLLEHFPSIPTLHVVGKKDFVYEYSTILQASTASAWATLIAHEKGHE
ISNDTRIVQKINSAFEQMSIRIALGC
```

Figure S6: Modified sequence for the maleidride hydrolase PhiM from the phomoidride BGC (BBG28510.1mod ).

An alignment of all maleidride hydrolases suggests that the annotation of EED15412.1 - the hydrolase from the *Talaromyces stipitatus* cluster 1 - may have identified the start codon incorrectly (Figure SX).

```
Cc_RDW56968.1              ---------------------MPSLKILCLHGTGCNAKPN-IERQQALLNTHLQRKSIAS
TbR3                       ---------------------MPTFRFLCLHGSGTNSDVCLLS---RPIIRKLEGDGIAS
Ia_CAF9941818.1_b          ---------------------MPSLKFLCLHGAGTREDI--MESQMRNLTRDLGKDHSAS
Ag_OJJ86354.1              ---------------------MPQLRFLCLHGAGTNTDI--LRSQLGALSRELSNDQTAD
EpiR11                     ---------------------MPALKVLCLHGAGTSVKI--LQSQLGPLMRDLQSDHSAT
Bm48331_XP_014073929.1     ---------------------MPPLRVLCLHGAGTNTEI--LQFQLRPLIRELESDHTAT
BmC5_EMD85581.1            ---------------------MPPLRVLCLHGAGTNTEI--LQFQLRPLIRELESDHTAT
Bv_XP_014553701.1          ---------------------MPPLRVLCLHGAGTNTKI--LQFQLRSLIRDLESDHTAA
Bz_XP_007715667.1          ---------------------MPPLRVLCLHGAGTNTKI--LQFQLRSLIRDLESDHTAT
BfL1_ANF07287.1            ---------------------MPQVKLLCLHGAGTSIQI--LQSQLAPLIRELQKDSTAS
CzmL1                      ---------------------MP-LKFLCLHGWGTNSSI--LRSQLDTVRKNLLVDHTVE
CaL1                       ---------------------MP-LKFLCLHGWGTNSKVSTLLDGVGALMRELKRDNTAE
Om_KIN05363.1              ---------------------MT-LRFLCLHGWGTSVKI--LESQLDALMRELKRDNMAD
TsRbtS_EED18832.1          ---------------------MSALKILCLHGAGTNAQI--LDSQLAPLVRALQKDQIAT
TfL9                       ---------------------MPGLKILCLHGAGTNAEI--LDTQLAPLVRELRKDSTAT
ScyR1_QTE76001.1           ---------------------MPAMRFLCLHGAGTNVQI--LNSQLGPIMRELQKDNSAT
Dc_ZopR1_QTE75993.1        ---------------------MPTLKFLCLHGAGTNAQI--LGSQLGPLMRELGKDNSAI
Zc_ZopM_BBU42027.1         ---------------------MPTLKFLCLHGAGTNAQI--LGSQLGPLMRELGKDNSAI
PvL1_ASK38716.1            ---------------------MP-VRFLCLHGWGTNIQI--LQSQLGPLMRELQKDNSAE
Ta_OKL57045.1              ---------------------MP-MRLLCLHGWGTNVKI--LQSQLAPLMKDLQRDNSAV
PhiM_BBG28510.1mod         ---------------------MPGLRILCLHGQGSNTEI--MKAQLGPITKLLKQNGVAT
Ts1_EED15412.1mod          ---------------------MPALKLLCLHGAGMNSEI--MKSHLSSLAKTLEYRNIAQ
Ts1_EED15412.1             MSRAHPKQPVTGRNIPILAITMPALKLLCLHGAGMNSEI--MKSHLSSLAKTLEYRNIAQ
                                                *. ...***** *   .   :      :   *     .
```

Figure S7: Section from a multiple sequence alignment using MUSCLE[1] of various maleidride hydrolase enzymes. An asterisk (*) marks completely conserved amino acid residues, whereas a colon (:) identifies highly conserved residues and a period (.) indicates moderately conserved residues. The deposited sequence from the *T. stipitatus* 1 BGC is shown (Ts1_EED15412.1) with the extra sequence depicted in red, as well as the modified sequence (Ts1_EED15412.1mod) in which the sequence begins at a downstream methionine.

```
>Ts1_EED15412.1mod
MPALKLLCLHGAGMNSEIMKSHLSSLAKTLEYRNIAQFAYAEGSVETEPGPGITPGLYEGPYYSFHIWPPKAGNLQDEESIQNAYEELLEIIDDEGPFD
GLLGFSHGGSFLAELLARYARDNPATDVERLARCAVFINSFPPFRNDPDQNPIIDYELLKHFPKIPTLHVVGTSDFVHEYSTILYEKLHQKAPTSTGLV
THSKGHEIPRDPKVLDKVVAGFEKLNFAVSFSH
```

Figure S8: Modified sequence for the maleidride hydrolase from the *T. stipitatus* 1 BGC, EED15412.1mod

CAF9941818.1 from the *Imshaugia aleurites* BGC appears to be incorrectly annotated. Comparisons using ACT[5] determined that this sequence has been incorrectly fused, and in fact encodes two genes: a maleidride hydrolase, and a gene containing a SnoaL-like domain, with homology to TsRbtF. The *I. aleurites* genomic sequence (CAJPDT010000158.1:26000-27600 – the 3′ end of the fused gene) was submitted to FGENESH[6] using the *Oidiodendron maius* gene-finding parameters (both *I. aleurites* and *O. maius* are in the Leotiomyceta clade). The subsequent sequence was aligned to other maleidride hydrolases (Figure S9), and showed high homology.

```
Cc_RDW56968.1              MPSLKILCLHGTGCNAKPN-IERQQALLNTHLQRKSIASLSFLEGDVET-EPGPGIG-DF
TbR3                       MPTFRFLCLHGSGTNSDVCLLS---RPIIRKLEGDGIASFEFIDGELESTTPGPGIQ-GF
Ia_CAF9941818.1_b_mod      MPSLKFLCLHGAGTREDI--MESQMRNLTRDLGKDHSASFIYVGGEISC-DPGPGIE-HI
Ag_OJJ86354.1              MPQLRFLCLHGAGTNTDI--LRSQLGALSRELSNDQTADLHFLEGGVDS-PPGPGVL-GY
EpiR11                     MPALKVLCLHGAGTSVKI--LQSQLGPLMRDLQSDHSATFHFTEGEVDS-DPGPGVA-TF
Bm48331_XP_014073929.1     MPPLRVLCLHGAGTNTEI--LQFQLRPLIRELESDHTATFHFTEGVVDS-GPGPNVE-NF
BmC5_EMD85581.1            MPPLRVLCLHGAGTNTEI--LQFQLRPLIRELESDHTATFHFTEGVVDS-GPGPNVE-NF
Bv_XP_014553701.1          MPPLRVLCLHGAGTNTKI--LQFQLRSLIRDLESDHTAAFHFTEGNIDS-GPGPNVE-NF
Bz_XP_007715667.1          MPPLRVLCLHGAGTNTKI--LQFQLRSLIRDLESDHTATFHFTEGNIDS-GPGPNVE-NF
BfL1_ANF07287.1            MPQVKLLCLHGAGTSIQI--LQSQLAPLIRELQKDSTASFHFIEGEVEC-GPGPGIE-GI
CzmL1                      MP-LKFLCLHGWGTNSSI--LRSQLDTVRKNLLVDHTVEFHFHEGDIES-EPGPGIE-GF
CaL1                       MP-LKFLCLHGWGTNSKVSTLLDGVGALMRELKRDNTAEFYFFEGDLDS-EPGPGIE-GY
Om_KIN05363.1              -MTLRFLCLHGWGTSVKI--LESQLDALMRELKRDNMADFYLVEGDLAS-EPGPGIE-GF
TsRbtS_EED18832.1          MSALKILCLHGAGTNAQI--LDSQLAPLVRALQKDQIATFHSVEGEVED-SPGPGIE-GF
TfL9                       MPGLKILCLHGAGTNAEI--LDTQLAPLVRELRKDSTATFHSVEGEVED-APGPGVE-GF
ScyR1_QTE76001.1           MPAMRFLCLHGAGTNVQI--LNSQLGPIMRELQKDNSATFHFVEGEIES-LPGPGVE-GF
ZopR1_QTE75993.1           MPTLKFLCLHGAGTNAQI--LGSQLGPLMRELGKDNSAIFHLVEGDVGS-PPGPGIE-GF
ZopM_BBU42027.1            MPTLKFLCLHGAGTNAQI--LGSQLGPLMRELGKDNSAIFHLVEGDVGS-PPGPGIE-GF
PvL1_ASK38716.1            MP-VRFLCLHGWGTNIQI--LQSQLGPLMRELQKDNSAEFHFIQGDVEA-DPGPGIE-GF
Ta_OKL57045.1              MP-MRLLCLHGWGTNVKI--LQSQLAPLMKDLQRDNSAVFHFIEGEIEA-EPGPGID-GY
PhiM_BBG28510.1mod         MPGLRILCLHGQGSNTEI--MKAQLGPITKLLKQNGVATFEWLEGTIPT-EAGPGVS-GV
Ts1_EED15412.1mod          MPALKLLCLHGAGMNSEI--MKSHLSSLAKTLEYRNIAQFAYAEGSVET-EPGPGITPGL
                              ...***** *   .   :      :   *     . :    * :    .**.:
```

Figure S9: Section of a multiple sequence alignment of maleidride hydrolases using MUSCLE.[1] An asterisk (*) marks completely conserved amino acid residues, whereas a colon (:) identifies highly conserved residues and a period (.) indicates moderately conserved residues. The modified *Imshaugia aleurites* sequence (CAF9941818.1_b_mod) is highlighted in red.

```
>Ia_CAF9941818.1_b_mod
MPSLKFLCLHGAGTREDIMESQMRNLTRDLGKDHSASFIYVGGEISCDPGPGIEHIYEGPYYSYYNWPQRLETDDQESVQSAYDLLDDIISTEGPFDGI
LGFSHGATLAFAFLVQHARKHPYEPPPVRCAVFVCAMPPFRLGGNDEWIYDQAFLEPGALAIPSVHVVGKSDFVLEHSLKLYAVCDAARARLVVHGKGH
EIPGDKENVALMALAVRELAHRAMFG
```

Figure S10: Modified sequence for the maleidride hydrolase from the *I. aleurites* BGC, CAF9941818.1_b_mod.

#### PEBPs

The deposited sequence for the maleidride PEBP type 2 from the *T. stipitatus* 1 BGC (EED15405.1) appears incorrect. An alignment with the other type 2 PEBPs, along with an N-terminal truncated version of EED15405.1, shows that the modified version is likely to be correct (Figure S11).

```
ScyR12_PEBP2_QTE76012.1             ------------------------------------MGIPNSVEYCLGRLLINQKGRDKG
Cc_PEBP2_RDW56972.1                 --------------------------------MGLLSTILKYVEYALGKLLISRRGYDDG
PhiB_PEBP2_BBG28499.1               -----------------------------------MSAILKLLEYCLGRLFYRRRGYDAQ
Ts1_PEBP2_EED15405.1mod             -----------------------------------MWFILRCLEWTLAKLLYRRRGYDHD
Ts1_EED15405.1                      MPDGIVASEAKDNTNNNLVHSFLGPKFPAASESSIMWFILRCLEWTLAKLLYRRRGYDHD
BfL9_PEBP2_ANF07279.1               ------------------------------------MSILAYVQYGLGKAFSPIRGHDSK
EpiR4_PEBP2                         ------------------------------------MAVIDYIEYALGTLLSCIRGHDSK
                                                                          :   ::: *.  :   .* *

ScyR12_PEBP2_QTE76012.1             LFFKTPAFTSMAEPTFTVTSPDCGPSTSHMKEEYTGV---GKDRFPELAWQKPSP-DVVE
Cc_PEBP2_RDW56972.1                 LFSRTPAFRKCPEPTIQLSSLDCGPSYSRLSHDYSMF---GKGLMPTLTWPEANE-NIRE
PhiB_PEBP2_BBG28499.1               LFYKSAAFAKHSSPTIPITSPDCGKTGAILTTEYSKF---GSGKIPQFTWPAAAP-DVKE
Ts1_PEBP2_EED15405.1mod             LFYKGLAFGKYPNPTFTVTSPDCGPTGAKLGVEYSQW---GSGKVPQLTWPVSGI-EVKE
Ts1_EED15405.1                      LFYKGLAFGKYPNPTFTVTSPDCGPTGAKLGVEYSQW---GSGKVPQLTWPVSGI-EVKE
BfL9_PEBP2_ANF07279.1               AITKTPAFKDIPQPNMTLEAPECGPSGSKLLDHHTCLAKDGKGDFPELRWSAPGLGDVKE
EpiR4_PEBP2                         LFTKGPAFSDFPAPNINLECPECGPTGSAMHYHHTSF---AAEDFPHLTWKDTPP-GTVE
                                     : .  ** . . *.: : . :** : : :   ::     .   .* : *  .      *

ScyR12_PEBP2_QTE76012.1             YVLIVEDPDAPLPM-PIT-HGLFYAIPGNKTQITNDDISV--EKITGKEKHLKGGFRLGK
Cc_PEBP2_RDW56972.1                 YLVIIEDVDAPFGGKPNV-HGIYALIPPSTTSLSPHDLETVDTDVKG-QHRLKSGFRVGK
PhiB_PEBP2_BBG28499.1               FLMLCEDPDAPMGH-PNV-HGIYCFIPPTVTSFGPTDLELI-KEVDG-VKVLESGYRVGK
Ts1_PEBP2_EED15405.1mod             YLIISEDPDAPLGH-SNV-HGIYCFVPGNKTGFGPDDLELLGEDKNG-LKQISSGYLVGK
Ts1_EED15405.1                      YLIISEDPDAPLGH-SNV-HGIYCFVPGNKTGFGPDDLELLGEDKNG-LKQISSGYLVGK
BfL9_PEBP2_ANF07279.1               YVLICEDLDLPIPG-LVMHHGIFYGIPPSTTSATNADVQHNGKDAK--DYVTTAGWKYIP
EpiR4_PEBP2                         YLLVVEDADSPIPS-PVC-HGIYYGIPASRNQLTAEEFHVINDGGN--EKKLSGGFYYGD
                                    :::: ** * *:       **::  :* . .     :.               .*:
```

```
ScyR12_PEBP2_QTE76012.1             NILGSVYGGPKPPLGHGVHRYYYTLVALKEPLDASKMSPLATKNEIAEAIEGKMIGWGQW
Cc_PEBP2_RDW56972.1                 NRRNVVYIPCRPPLGHGPHRYFCELVALSEKLDPDTLSPVPTKEELAELVKGKVVAWGEW
PhiB_PEBP2_BBG28499.1               NRRNVVYIAPRPPLGHGPHRYLFELVALSEKLDPEGISKVPDKGEIEKAIEGKVASWGLW
Ts1_PEBP2_EED15405.1mod             NRRNTVYIAPRPPLGHGPHRYFFEIVALSQPLDPEKLSPVPTKQELSDMIIGKVCGWGLW
Ts1_EED15405.1                      NRRNTVYIAPRPPLGHGPHRYFFEIVALSQPLDPEKLSPVPTKQELSDMIIGKVCGWGLW
BfL9_PEBP2_ANF07279.1               NMMGSPYLGPAPPLGHGSHRYVFYIVALKEPLDPEQPEKL-NRQTLAEAMTGKVIGWGQW
EpiR4_PEBP2                         IVKGKHYIGARAVLGHGPHRYVYQLVALREKLDVSPMGGKVAKDVLGRAIEGKVGGWGVW
                                       .  *    . **** ***   :*** : ** .       .  :   : **: .** *

ScyR12_PEBP2_QTE76012.1             VGLFERKWE--
Cc_PEBP2_RDW56972.1                 VGVYESDWNER
PhiB_PEBP2_BBG28499.1               EATYESTWDRK
Ts1_PEBP2_EED15405.1mod             TATFEQKWSM-
Ts1_EED15405.1                      TATFEQKWSM-
BfL9_PEBP2_ANF07279.1               IGTWERPWPR-
EpiR4_PEBP2                         IGTWERVWG--
                                     . :*  *
```

Figure S11: Multiple sequence alignment using MUSCLE[1] of various maleidride type 2 PEBPs, including the deposited sequence for the *T. stipitatus* 1 BGC (EED15405.1), as well as a modified version (EED15405.1mod). An asterisk (*) marks completely conserved amino acid residues, whereas a colon (:) identifies highly conserved residues and a period (.) indicates moderately conserved residues.

```
>Ts1_EED15405.1mod
MWFILRCLEWTLAKLLYRRRGYDHDLFYKGLAFGKYPNPTFTVTSPDCGPTGAKLGVEYSQWGSGKVPQLTWPVSGIEVKEYLIISEDPDAPLGHSNVH
GIYCFVPGNKTGFGPDDLELLGEDKNGLKQISSGYLVGKNRRNTVYIAPRPPLGHGPHRYFFEIVALSQPLDPEKLSPVPTKQELSDMIIGKVCGWGLW
TATFEQKWSM
```

Figure S12: Modified sequence for the maleidride type 2 PEBP, Ts1R3, from the *T. stipitatus* 1 BGC, EED15405.1mod.

The annotation of PhiN, the type 1 PEBP from the published phomoidride BGC also appears incorrect. The published protein sequence (BBG28511.1) aligns poorly with known type 1 PEBPs (Figure S13), and the intron structure is immediately apparent as unlikely (Figure S14). Re-annotation of this genomic locus generates a predicted protein (BBG28511.1mod) which aligns much better with known type 1 PEBPs (Figure S15 and Figure S16).

```
phiN_BBG28511.1_PEBP1      MFSSLLDLRVHRVYHISTWFGQVRESYRKPPTSTLVNGFGNHVGPRTSERELTYGVFGED
BfL5_ANF07283.1_PEBP1      MIGYIVQFSILLTIHVLA---------QTPP------GYEPS-SNHELYLEYPGEISVFP
PvR1_ASK38718.1_PEBP1      MRPDLWLLLALVCLPLIA--------AQTPP------GYWPK-TPRGLDVIFRDQQLIQP
                           *   :  :       : :         ..**      *:      .

phiN_BBG28511.1_PEBP1      GKLTKLCIGRYVEFAPRLDLKSLSSCSPSAYLAFMIDIDIIRDGRTVHLLHWYQPDLVLA
BfL5_ANF07283.1_PEBP1      GISLELSDTKCIPTLSTLGLSRIQT-----YIVFLIDIDVILEPNATTILHWYQPDLKAQ
PvR1_ASK38718.1_PEBP1      GQLVLPDDAIDPPVFGRQGLSVFQS-----YLAVMIDVEVNHEGTATPLVHWLQPDLKVR
                           *                 .*. :.:     *:..:**:::  :  :. ::** ****

phiN_BBG28511.1_PEBP1      H-KTHELVQTCSDRKGALYGAPAPPGGSSHRYVELVFQQPLNFTFPESFEHYLEPTIPAR
BfL5_ANF07283.1_PEBP1      D-LGSHLVNTTNN--GAAYLGPHPPAGNTHRYVFLLFQQPQGYTLPSCFSTIFPETVEAR
PvR1_ASK38718.1_PEBP1      DPFTGRLARLSDE--DVPYVGPRPVPGPRHTYVLLLFEQPTTYRFPECFSSTRPISVDTR
                                 *..  .:  .. * .* *  *  * ** *:*:**  : :*..*.     :: :*

phiN_BBG28511.1_PEBP1      LFFNITEFAAAAELGNPVAANYFTVLGTRTSTEEQQIIEL-------------------
BfL5_ANF07283.1_PEBP1      AGFNIDEFIEVAGLGDVVAANYVNVTNPATPSTTLT-ATTTSLSTAPCPITMAKFLSST
PvR1_ASK38718.1_PEBP1      SGFNLAQFMHVAGLQEPIAASYFTARNEETPSAPPPRVTTTSLSTAPCA-TPTRFVWC-
                             **: :*  .* * : :**.*... .  *.:
```

Figure S13: Multiple sequence alignment using MUSCLE[1] comparing two sequences of verified type 1 PEBPs to the original PhiN sequence (BBG28511.1). The original PhiN has 33.33% identity with BfL5 and 28.93% identify with PvR1. An asterisk (*) marks completely conserved amino acid residues, whereas a colon (:) identifies highly conserved residues and a period (.) indicates moderately conserved residues.

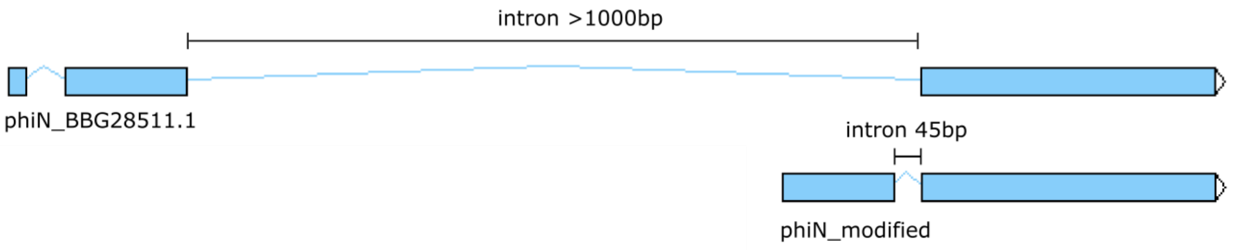

Figure S14: Comparison of the original phiN sequence, and intron structure, compared to the modified phiN sequence.

```
PhiN_BBG28511.1mod          MTLSVVASILSSFVLLTRP-ISGQTPPGFSPSTPNNLGVTFSNETEVAFPGAQLSGNDVE
BfL5_ANF07283.1_PEBP1       ----MIGYIVQFSILLTIH-VLAQTPPGYEPSSNHELYLEYPGEISV-FPGISLELSDTK
PvR1_ASK38718.1_PEBP1       ----MRPDLWLLLALVCLPLIAAQTPPGYWPKTPRGLDVIFRDQQLI-QPGQLVLPDDAI
                                :   :     *:    : .*****: *.: . * : : .:  :  **  :   .*.

PhiN_BBG28511.1mod          FAPRLDLKSLSSCSPSAYLAFMIDIDIIRDGRTVHLLHWYQPDLVLAH-KTHELVQTCSD
BfL5_ANF07283.1_PEBP1       CIPTLSTLGLSRI--QTYIVFLIDIDVILEPNATTILHWYQPDLKAQD-LGSHLVNTTNN
PvR1_ASK38718.1_PEBP1       DPPVFGRQGLSVF--QSYLAVMIDVEVNHEGTATPLVHWLQPDLKVRDPFTGRLARLSDE
                              * :.  .**    .:*:..:**:::  :  :. ::** ****         *..  .:

PhiN_BBG28511.1mod          RKGALYGAPAPPGGSSHRYVELVFQQPLNFTFPESFEHYLEPTIPARLFFNITEFAAAAE
BfL5_ANF07283.1_PEBP1       --GAAYLGPHPPAGNTHRYVFLLFQQPQGYTLPSCFSTIFPETVEARAGFNIDEFIEVAG
PvR1_ASK38718.1_PEBP1       --DVPYVGPRPVPGPRHTYVLLLFEQPTTYRFPECFSSTRPISVDTRSGFNLAQFMHVAG
                              .. * .* *  *  * ** *:*:**  : :*..*.     :: :*  **: :*  .*

PhiN_BBG28511.1mod          LGNPVAANYFTVLGTRTSTEEQQIIEL-------------------
BfL5_ANF07283.1_PEBP1       LGDVVAANYVNVTNPATPSTTLT-ATTTSLSTAPCPITMAKFLSST
PvR1_ASK38718.1_PEBP1       LQEPIAASYFTARNEETPSAPPPRVTTTSLSTAPCA-TPTRFVWC-
                            * : :**.*... .  *.:
```

Figure S15: Multiple sequence alignment using MUSCLE[1] comparing two sequences of verified type 1 PEBPs to the modified PhiN sequence (BBG28511.1mod). An asterisk (*) marks completely conserved amino acid residues, whereas a colon (:) identifies highly conserved residues and a period (.) indicates moderately conserved residues. The modified PhiN has 40% identity with BfL5 and 32.65% identify with PvR1. The type 1 PEBPs contain modified versions of the known PEBP motifs: DPDXPX$_n$H and and GXHR, depicted in red.[7 8]

```
>PhiN_BBG28511.1mod
MTLSVVASILSSFVLLTRPISGQTPPGFSPSTPNNLGVTFSNETEVAFPGAQLSGNDVEFAPRLDLKSLSSCSPSAYLAFMIDIDIIRDGRTVHLLHWY
QPDLVLAHKTHELVQTCSDRKGALYGAPAPPGGSSHRYVELVFQQPLNFTFPESFEHYLEPTIPARLFFNITEFAAAAELGNPVAANYFTVLGTRTSTE
EQQIIEL
```

Figure S16: Modified sequence for the maleidride type 1 PEBP, PhiN, from the phomoidride BGC, BBG28511.1mod.

#### ACDHs

The sequence deposited for the ACDH enzyme from the *T. stipitatus* 1 BGC (EED15410.1) appears to be incorrect. An alignment with other maleidride ACDH enzymes, as well as a modified version of the *T. stipitatus* sequence (EED15410.1mod) where a read-through intron has been removed, shows that the modified version is likely to be correct (Figure S17).

```
Ts1_ACDH_EED15410.1mod          GCYQLQNAFNAYGIDHVILVKLAAASVCSWLMGMTETQTMSVISHVWMDGHPSRVYRSGA
Ts1_ACDH_EED15410.1             GCYQLQNAFNAYGIDHVILVKLAAASVCSWLMGMTETQTI-----------------SGA
Cc_ACDH_RDW56964.1              GCYQMRNAFNEHGLDHVILVKLASTAVVSWLLGLTEEQTMAAISHVWMDGHPSRVYRSGA
BBG28506.1_ACDH_PhiI            GCYQLLNAFNIYGIDHVVLVKLASTAAVSWLLGLTEEQTMAAISHVWMDGQPTRVYRSGE
Ta_ACDH_OKL57050.1              GCLLLRNAFNAVGLDHTILVKLASTAVVSWLLGLSEDQTLAAISHAWMDLAPLRVYRSGS
Wa_ACDH_EpiR8                   GCFLLRNAFNAHGLDHVILVKLASTAVVSWLLGLTEEQTMAAISHVWMDGHPLRTYRQGT
Bv_ACDH_XP_014553699.1          GCFLLQNAFNKHGLDHVILVKLASTAVVAWLLGLTEEETMAAISHVWMDGHPLRTYRHGT
Bz_ACDH_XP_007715665.1          GCFLLQNAFNKHGLDHVILVKLASTAVVAWLLGLTEEETMAAISHVWMDGHPLRTYRHGT
Bm_C5_ACDH_EMD85579.1           GCFLLRNAFNKQGLDHVILVKLASTAVVAWLLGLTEEETMAAISHVWMDGHPLRTYRHGT
Bm_48331_ACDH_XP_014073931.1    GCFLLRNAFNKQGLDHVILVKLASTAVVAWLLGLTEEETMAAISHVWMDGHPLRTYRHGT
Bf_ACDH_ANF07285.1              GCFLLHNAFNSHGLDHVILVKLASTALVCWLLGLTETHTMAAISHVWMDGHPLRVYRSGS
Dc_ACDH_QTE75987.1              GCFLLRNAFNAHGLDHVILVKLASTAVVSWLLGLTEQQCHAAISHVWMDGQPLRVYRSGV
Zc_ACDH_BBU42021.1              GCFLLRNAFNAHGLDHVILVKLASTAVVSWLLGLTEQQCHAAISHVWMDGQPLRVYRSGV
Sa_ACDH_QTE76007.1              GCFLLKNAFNAHGLDHVILVKLASTAVVAWLLGLTEHQCQAAISHVWMDGHPLRVYRSGV
Pv_ACDH_ASK38715.1              GCMLLQNAFHSHGLDHVVLVKLASTAVVAWLLGLSETQTMAAISQVWMDGHPLRVYRSGS
Czm_ACDH_KAF2215728.1           GCMLLRNAFNQYGLDHVILVKLASTAVTAWLMGLSIPQIRAAISQVWMDGHALRVYRQKG
Ca_ACDH_PVH77202.1              GCMLLRNAFNAHGLDHVILVKLASTAVVAWMVGLSEEQTMQAISQVWMDGHPLRVYRQKG
Om_ACDH_KIN05360.1              GCMLLRNAFNAYGLDHVILVKLASTAIISWLLGLTEDQTMAAISQVWMDGHALRVYRQKG
Ts2_ACDH_EED18840.1_RbtK        GCMLLRNAFNAYGLDHVILVKLASTAVVSWLLGLTEIQTMAAISQVWMDGQALRVYRQRG
TfL1_ACDH                       GCMLLRNAFNAYGLDHVILVKLASTAVVSWLMGSTEGQTMAAISQVWMDGQALRVYRQKG
Ag_ACDH_OJJ86350.1              GCFLIKNAFHKKGLDHVILVKLASTAVVSWLLGLSEEQTLAALSHVWMDGAALRVYRSGS
TbR1_ACDH                       GCFLIKNAFNALGFDHVILVKLASTAVVSWLMGLSELQTLSALSHVWMDGAALRIYRSGS
Ia_ACDH_CAF9941813.1            GVFLLSNPFNAHGLDHIILVKLASTAVVSWLLGLSEVQTLAAISHVWMDGAALRIYRSGS
                                *   : *.*:  *:** :*****:::  .*::* :
```

Figure S17: Section from a multiple sequence alignment using MUSCLE[1] of various maleidride ACDH enzymes. An asterisk (*) marks completely conserved amino acid residues, whereas a colon (:) identifies highly conserved residues and a period (.) indicates moderately conserved residues. The deposited sequence for the ACDH from the *T. stipitatus* 1 BGC is shown (Ts1_EED15410.1), as well as a modified sequence, where a read-through intron has been removed (Ts1_EED15410.1mod).

```
>Ts1_EED15410.1mod
MTDEIPFDKPIRDIASYVHNYTIPSSPTSFKHARGVILDSLGCAIETLHRSSEACTLLGPVIPGTTFPYGFHLPGTSYVLDPVKGTFDLGVLIRYLDHN
DAFGGAEWGHPSDTLSAIIAVMDWLCRAEQRSSTTSIRYPPLTIKTLLEAAIKAYEIQGCYQLQNAFNAYGIDHVILVKLAAASVCSWLMGMTETQTMS
VISHVWMDGHPSRVYRSGANTTSRKGWAAADAAMRAVHLCLLTHAGQLGSKQPLNDKRYGFLVHTFGLEAGFALPRAFGDWAIQNIFTKLMPCEGHGIS
AVEAALVQGRKLKSCGHTVSDIKHIDLRVTAAANLIISKIGRLYNAADRDHCIQYVIALAFLKGRFPDAEDYMDDSPYANSKEMDDLREKIMMKVDQDL
TQGYLDPERKSCGTGMTVYLNNGTVLDEVLVEYPAGHLKNPRTKELHQRKFEKNMRLAFTDAEIANIVKCIEDDEMPISSFVDLFTRDSKGMAKL
```

Figure S18: Modified sequence for the maleidride ACDH from the *T. stipitatus* 1 BGC, EED15410.1mod

#### ICMs

The sequence deposited for the isochorismatase (ICM)-like protein from the rubratoxin cluster of *T. stipitatus* appears to contain an incorrectly identified read-through intron. When this intron is removed, the ICM aligns very well will other confirmed maleidride ICMs (Figure S19).

```
QTE75990.1-ZopL2(zopfiellin)          MAKTALLVMDVQGAFVPRLAQSSDYLPRLAKTIAAARPSVVKVIYTRVAFRPGHPEISPS    60
BBU42024.1-ZopQ(zopfiellin)           MAKTALLVMDVQGAFVPRLAQSSDYLPRLAKTIAAARPSVVKVIYTRVAFRPGHPEISPS    60
QTE76004.1-ScyR4(scytalidin)          MDKTALLVMDVQGGMVSRLPTAQTFIPLLSKTVAAARPF-VKIIYATVSFRPGHPEIAPS    59
EED18834.1mod-TsRbtQ(rubratoxin)      MSKTALLVMDYQAGIISRLSLPENHLQLLANTIDTARPY-AKIIYVTVAFRPGHPEVSAS    59
EED18834.1                            MSKTALLVMDYQAGIISRLSLPENHLQLLANTIDTARPY-AKIIYVTVAFRPGHPEVSAS    59
                                      * ******** *..:: **   . .:  *::*: :***  .*:**. *:*******:: *

QTE75990.1-ZopL2(zopfiellin)          NPTFSAAVKSKSFVEGSPETLIDPSIAPQEGDVLVDKKRVSAFSGSGLDIILSSLGIETV    120
BBU42024.1-ZopQ(zopfiellin)           NPTFSAAVKSKSFVEGSPETLIDPSIAPQEGDVLVDKKRVSAFSGSGLDIILSSLGIETV    120
QTE76004.1-ScyR4(scytalidin)          NVIFSAALKSGAFVAGSPETVIDPSIAPQEGDILVEKKRVSAFAGSGLDVILRGLGIETL    119
EED18834.1mod-TsRbtQ(rubratoxin)      NATFSAAAKSNSFVSGSPETQIDPVIAPKEGDILIEKKRVSAFTGSGLDLVLKGLGVETL    119
EED18834.1                            NATFSAAAKSNSFVSGSPETQIDPVIAPKEGDILIEKKRVSAFTGSGLDLVLKGLGVETL    119
                                      *  **** ** :** ***** *** ***:***:*::*******:*****::* .**:**:

QTE75990.1-ZopL2(zopfiellin)          VLAGMSTGGVVLSTVLEASDKDFGVVVLKDLCVDADETLHNTLMDKIFTKRGQVVEAEKW    180
BBU42024.1-ZopQ(zopfiellin)           VLAGMSTGGVVLSTVLEASDKDFGVVVLKDLCVDADETLHNTLMDKIFTKRGQVVEAEKW    180
QTE76004.1-ScyR4(scytalidin)          VLTGISTGGVVLSTVCEAADKDFKLVILKDLCADPDQAVHNVLMENVFTKRGEVLGAEEW    179
EED18834.1mod-TsRbtQ(rubratoxin)      VLAGISTGGVVLSTVCEAADKDFKLVVLKDLCVDGDEKLHNGLMSKIFSKRGEVLGAEEW    179
EED18834.1                            VLAGISTGGVVLSTVCEAADKDFKLVVLKDF--------------KIFSKRGEVLGAEEW    165
                                      **:*:********** **:**** :*:***:              ::*:***:*: **:*

QTE75990.1-ZopL2(zopfiellin)          LETLKA      186
BBU42024.1-ZopQ(zopfiellin)           LETLKA      186
QTE76004.1-ScyR4(scytalidin)          LEKLKA      185
EED18834.1mod-TsRbtQ(rubratoxin)      LQKLKA      185
EED18834.1                            LQKLKA      171
                                      *:.***
```

Figure S19: A multiple sequence alignment produced using clustal omega[9] of various isochorismatase-like enzymes from confirmed maleidride clusters. An asterisk (*) marks completely conserved amino acid residues, whereas a colon (:) identifies highly conserved residues and a period (.) indicates moderately conserved residues. The deposited sequence for the ICM-like enzyme from the *T. stipitatus* 2 BGC (likely rubratoxin BGC) is shown (EED18834.1), as well as a modified sequence (EED18834.1mod), where a read-through intron has been removed.

```
>EED18834.1mod-TsRbtQ(rubratoxin)
MSKTALLVMDYQAGIISRLSLPENHLQLLANTIDTARPYAKIIYVTVAFRPGHPEVSASNATFSAAAKSNSFVSGSPETQIDPVIAPKEGDILIEKKRV
SAFTGSGLDLVLKGLGVETLVLAGISTGGVVLSTVCEAADKDFKLVVLKDLCVDGDEKLHNGLMSKIFSKRGEVLGAEEWLQKLKA
```

Figure S20: Modified sequence for the maleidride ICM-like enzyme from the *T. stipitatus* 2 BGC, EED18834.1mod.

#### αKGDDs

```
TsRbtU_EED18830.1mod        RSAWLKYGILRFRGYDLTDEHQLKLTKLI--GSFLKREEDGAPTTYKDDEKVTVMTNL-I
TsRbtU_EED18830.1           RSAWLKYGILRFRGYDLTDEHQLNFL---------KREEDGAPTTYKDDEKVTVMTNL-I
TfL12                       KQAWRDYGVLRFRGYDITTQQHVNFSNLF--GR--HVPVKSASIGHEEQEEITVISNVKV
TSrbtE_EED18846.1           RAAWRDYGVLRFRGYDITTQQHAKFSNLF--GH--YVPVKGTSIAHHDQKEITVISNAKV
Ta_OKL57041.1               ALLVTERGVVFFRDQDLTTEKQVELFEHYEEGILDKHPAQKHKDIYGNTDDHREIANF--
Om_KIN05358.1               ALLIAERSVVFFRDQDLSPQKQEELGKYW--GRIEYHP--HVPHVPGVPGASVVWDGLKP
Ca_PVH77204.1               ALLIAERCVVFFRDQDITPQQQEELGKYY--GRVEIHP--HVPHVPGAEGATVIWDALKS
                                 .  :: **. *:: ::: ::
```

Figure S21: A section from a multiple sequence alignment produced using MUSCLE[1] of various TauD-like αKGDDs from maleidride BGCs. An asterisk (*) marks completely conserved amino acid residues, whereas a colon (:) identifies highly conserved residues and a period (.) indicates moderately conserved residues. The deposited sequence from the Ts2 BGC, EED18830.1, has a read-though intron (highlighted in red), which when removed to produce EED18830.1mod aligns better to the other maleidride TauD-like αKGDDs.

```
>EED18830.1mod-TsRbtU(rubratoxin)
MSGDAERRDLTIVANKVGAGADVLGFDFDTMPSHQVGALRSAWLKYGILRFRGYDLTDEHQLKLTKLIGSFLKREEDGAPTTYKDDEKVTVMTNLINGV
PSGAGSNVELEWHTDSWFWEYPPVGEILRAMELPQTGGDTYWADMYAVYDALPEDLRSTIEGRLIQFDTVYNGHGNLRKGKEAPKTDDFRLWEHIRHPI
IRTHPESGRKAVFVGQSKHEKNWIVGLPLEESKEILAKILSYVEKPEFQLHQKWQPGDTVIWDNRCTMHRRETWPDDQTRIMHRTTCNTKGQPRPFYVY
```

Figure S22: Modified sequence for the maleidride TauD-like αKGDD TsRbtU from the *T. stipitatus* 2 BGC, EED18830.1mod.

Interpro analysis of EED18849.1, a deposited sequence from the Ts2 BGC, shows an unusual structure, in that it contains an AsaB-like αKGDD domain (IPR044053) in the N-terminal half of the sequence, as well as a thioesterase domain (IPR006683) towards the C-terminus. Comparison of the image of the rubratoxin BGC from *Penicillium dangeardii* Pitt (Figure S37) to the orthologous Ts2 BGC shows that EED18849.1

should encode the ortholog to RbtB, which is a confirmed αKGDD. Bai *et al*[10] make no mention of a thioesterase domain within RbtB. Removal of the final 3 exons from EED18849.1 to produce EED18849.1_a_mod aligns better to other maleidride AsaB-like αKGDDs (Figure S23).

```
Czm_KAF2215734.1_alphaKGDD            VAAN----EEHKWYWMSAQQPDEPLLLKIYDSK--KEGIARFCPHTAVQT----------
ZopL9_QTE75983.1_alphaKGDD            LKTG---KKDHDWYYASNQQPDEVLLFTQYSDFP-NRNTADRVPHVSVKLPGQE-DKPR-
Cc_RDW56970.1_alphaKGDD               VKAASPGKEEHSWYFAPEQRPDELLLFNQYSDKA-DRGIADRVAHAAFVLPGTE-DKPT-
PhiK_BBU42017.1_alphaKGDD             VKAG----EGHQWYYVPQQRTDEMILFTQYSDNP-NRGIADRVAHCAFILPGTE-DKPV-
Ts1_EED15414.1_alphaKGDD              VLYD----QRHKWYWVPMQKPNEMLLFNQYSDDP-NRTLADRVAHCGFTLPGAE-DKEI-
Ta_OKL57227.1_alphaKGDD               LEHP-----TLKWHYLSQQQPDEAIIFKCTDS---HEGVAKCVPHASIELPNTNSGTPA-
ScyL2_QTE75998.1_alphaKGDD            IQAP-----SMKWYYQSKMEDNTLLVFKSYES---QDGVAKYASHCSFPLPTAGPMTPP-
TsRbtG_EED18844.1_alphaKGDD           VHSP-----AMKWYYQSGMEDGTLLVLKNYDSHAEEGGVARYSAHCSFPLPTAGPDTPP-
Ts2_EED18849.1                        VHEP-----TLKWYYQSGMEDDTLLVLKNYDS---EDGVAKYVPHCSFSLPTATASTPPL
TsRbtB_EED18849.1_a_mod_alphaKGDD     VHEP-----TLKWYYQSGMEDDTLLVLKNYDS---EDGVAKYVPHCSFSLPTATASTPP-
                                      :          .*:: .    .  :::.  ..       *   .* ..

Czm_KAF2215734.1_alphaKGDD            ------------------------------------------------------------
ZopL9_QTE75983.1_alphaKGDD            ------------------------------------------------------------
Cc_RDW56970.1_alphaKGDD               ------------------------------------------------------------
PhiK_BBU42017.1_alphaKGDD             ------------------------------------------------------------
Ts1_EED15414.1_alphaKGDD              ------------------------------------------------------------
Ta_OKL57227.1_alphaKGDD               ------------------------------------------------------------
ScyL2_QTE75998.1_alphaKGDD            ------------------------------------------------------------
TsRbtG_EED18844.1_alphaKGDD           ------------------------------------------------------------
Ts2_EED18849.1                        AQQMEVPCQRMKVTETNGLILPGARGVQKLKSPAEMASSSIHSIALGDKRLQMDLTPGVF
TsRbtB_EED18849.1_a_mod_alphaKGDD     ------------------------------------------------------------
```

Figure S23: A section from a multiple sequence alignment produced using MUSCLE[1] of AsaB-like maleidride αKGDDs with the deposited sequence from the Ts2 BGC, EED18849.1. An asterisk (*) marks completely conserved amino acid residues, whereas a colon (:) identifies highly conserved residues and a period (.) indicates moderately conserved residues. This sequence contains a significant extra number of amino acids, when the final 3 exons are removed to produce EED18849.1_a_mod, the sequence aligns better.

```
>EED18849.1_a_mod-TsRbtB (rubratoxin)
MATTTTLATTSKNGGDVPAKLTYIEWHDHYETEEPHFVLNTPDDPPDAYAGNVTFKEGEEEIIHDIRGHEDKFTLDKQGFVFTKAPTSLSPSEFLDEEK
IKEKYLPECEKYYREYFKGIDEVVFIHYRARNSITADDHNSPTGPARVAHVDLSGPEINARIRKAFPDRADFILRGRVRLVNLWRPINGPLQNWPLCVA
DCNSIQEKHLVATKRIRKTHQAVTRLVVHEPTLKWYYQSGMEDDTLLVLKNYDSEDGVAKYVPHCSFSLPTATASTPPRESIEVRAFLFNYPRNEKSV
```

Figure S24: Modified sequence for TsRbtB from the *T. stipitatus* 2 BGC, EED18849.1_a_mod.

```
TsRbtU_EED18830.1mod_alphaKGD       LRSTIEG--------------------------RLIQFDTVYNGHGNLRKGKEAPKTDDF
TsRbtE_EED18846.1_alphaKGD          TRKIIEG--------------------------RLIQFNIVYDAVGRVRPGQEKPETDDF
TfL12_alphaKGD                      TRKIIEG--------------------------RLIQFDIVYDGYGRLRPGQEKPEEEDF
Ta_OKL57041.1mod_alphaKGD           YKRFLEGLH------------------------ALIIITIDDTIADLWGIAPNRPPIDTH
Ta_OKL57041.1_alphaKGD              YKRFLEGLHAVHTSRLQCMLCLSGSQNSDLIHDKLIIITIDDTIADLWGIAPNRPPIDTH
Om_KIN05358.1_alphaKGD              FREFIDG--------------------------KMAVFSSTHTYIDRNDPYAGPKFIQNI
Ca_PVH77204.1_alphaKGD              FRQFIDG--------------------------KKAVFRSTHSYVDRDDPHGARRYNENI
```

Figure S25: A section from a multiple sequence alignment produced using MUSCLE[1] of TauD-like maleidride αKGDDs with the deposited sequence from the *T. atroroseus* BGC, OKL57041.1. This sequence contains a read through intron, which when removed to produce OKL57041.1mod, sequence aligns better.

```
>Ta_OKL57041.1mod
MASLESRKHISPDSGFSVTLHPHFNARIASHVPSSPIREAFDPPKDRAHFADPEKKALFAVARRVDLTEGVGTLLENVQLSQLNAQQLDELALLVTERG
VVFFRDQDLTTEKQVELFEHYEEGILDKHPAQKHKDIYGNTDDHREIANFTPWPIAEWHADTSFEINPPSYSLLRMEEHPEIGGDTAWISQYGTYDALS
DAYKRFLEGLHALIIITIDDTIADLWGIAPNRPPIDTHHPAVRTHPVTGLKALNVNPGFVTGFAELKKLESDKVLDFLYTHIHSADDHYVRWKWNVGSV
AFWDNRCVAHRVIPGSYETPRRGVRTTVYGEKPFLDPASEGRLQRKLRKAREAKTNGTAVKQ
```

Figure S26: Modified sequence, OKL57041.1mod, for the TauD-like αKGDD.

#### Other

No thioesterase was identified in the *Penicillium dangeardii* Pitt rubratoxin BGC, however, the presence of a thioesterase domain (IPR006683) in EED18849.1 suggested that there was a separate gene at that locus. FGENESH[6] analysis of the sequence NW_002990116.1: 3979164 to 3980589, using the *Byssochlamys spectabilis* gene finding parameters (both *Talaromyces* and *B. spectabilis* are in the order Eurotiales) identified a single exon gene which contains the thioesterase domain, we have named this gene *tsrbtB2* (EED18849.1_b_mod, see Figure S27 for protein sequence). Further evidence for the presence of this gene comes from the comparison between the Ts2 BGC and the BGC from *T. funiculosus*, these semi-orthologous BGCs both contain the thioesterase, suggesting it is important for biosynthesis. (Figure S38).

>EED18849.1_b_mod-TsRbtB2 (rubratoxin)
MPRLPRQFVPTSIDLDARGHFREYSWCDKIFEDPSLQPVLTVNQHSWSDVPSTFMWQALGLPGAITAAQSFCKGSMSPPNHSFVEGRTELWTLYCFGRA
VESFLHVAHGGFLASLLDQQTGSIVITYPVSTNPRTLSSTIRYHKALITPGAVLCRAWISKVEGRKVWAKAVLEDGKGETIADMEALWIFLKPSL

Figure S27: Sequence for TsRbtB2 from the *T. stipitatus* 2 BGC, EED18849.1_b_mod.

TsRbtF from the Ts2 BGC encodes an unusual SnoaL-like domain. BLASTing the deposited sequence (EED18845.1) against the nr-database highlights that the deposited sequence appears to have and extra ~20 amino acids compared to other similar sequences (grey box in A, Figure S28). Moving the start codon to a downstream methionine to create EED18845.1mod aligns better to other similar sequences (B, Figure S28).

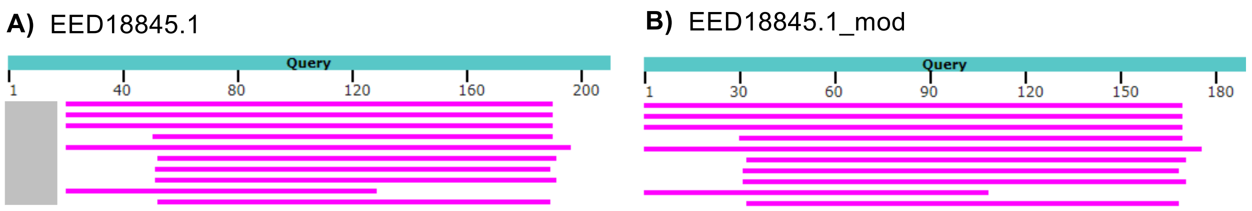

Figure S28: **A)** BLAST analysis of EED18845.1 against the nr-database shows that top hits are truncated in comparison to EED18845.1 which has an extra ~20 amino acids (grey box). **B)** BLAST analysis of EED18845.1mod (truncated at N-terminus) against the nr-database aligns better to other similar sequences.

>EED18845.1mod-TsRbtF (rubratoxin)
MRFLSILSAAVVVSVSRAQNLSALEKTLAITEIKNVQSLYGTIIDAKTMKDLSRVFTEDAVANYTVLGIGILTGLPLIIEGMTISQAHDVTQHAMTTSY
VDVLDENNANSTAYLTAETYGTGNNIGANNTGQLFTLWLKYEDQFVRTNGSWKIKNRNAVIMGTPLTGNFTPRVIPPPPSSLPIPTATAI

Figure S29: Modified sequence for TsRbtF from the *T. stipitatus* 2 BGC, EED18845.1mod.

TsRbtH from the Ts2 BGC encodes a ferric reductase. BLASTing the deposited sequence (EED18843.1) against the nr-database highlights that the deposited sequence appears to have and extra ~150 amino acids compared to other similar sequences (grey box, A, Figure S30). Removing the final 2 exons to create EED18845.1mod aligns better to other similar sequences (B, Figure S30).

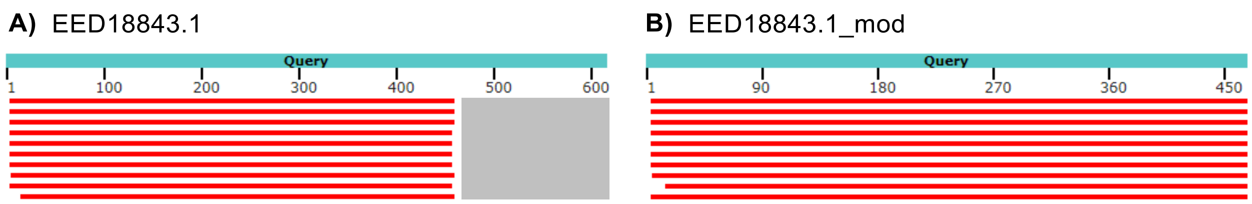

Figure S30: **A)** BLAST analysis of the deposited sequence (EED18843.1) against the nr-database highlights that the deposited sequence appears to have and extra ~150 amino acids compared to top hits (grey box). **B)** BLAST analysis against the nr-database of a C-terminus truncated sequence, EED18845.1mod, aligns better to other similar sequences.

>EED18845.1mod-TsRbtH (rubratoxin)
MSIATRNARRILLIPLPGVASVGQAVVIVLYAIVSIVLVFVDMPVSRGVDVYANRTGWLSFLNLSLITFLALRNSPLSFLTGYSYDKLILLHKVAGYST
IFWTILHIILFSIDGARKDLFSAEAKELKIVVGWVGGFAMLIILGTAVTRRWIPYEAFYTSHAALSIVILVTMCFHPDTWGCILVCIITAGFWSADLLI
RLIQLFWYSIGNRATLSPLPNGGVRVVMYKSPKTTPGSHCSLWIPSIRLIETHPVAVLSTDPLEFVIASYDGFTGELYKTSLNKKTVWASVHGSYGARI
DFTNLEKVVFIAGGSGASFTFALAMALMKSGGNSAKPGIEFIWVVRHQDPLNLYSGSLSWFSNELEELALSNRVNLTIHVSSLSDQEGDVEKYAWSGMV
TPHFSMEKIQAGRPDIDKIIDNVVGGMNLSDRLMVAACGPDGLMKTARRSVARNTRPGGPSIDFKCEEYSW

Figure S31: Modified sequence for TsRbtH from the *T. stipitatus* 2 BGC, EED18845.1mod.

```
A0A7H8R516|Epimerase      PFDMTALRPTNVYGYGSSFYGALFSFMEKGAAENDALKVFA-DPNNIMHSAHVDDCADAY
B8MDT8|Epimerase          PFDIVILRPTNVYGYGSSHTGSLFSWAEKSKKEGKKVLRAVADPNTILHYTHVDDCALAY
TSrbtC_EED18848.1mod      LFDTVVLRPTHVYGYMSSFYSFFFRFA-QEEKKKGEWVVEE-DPETILHSVHVDDCGDAY
TSrbtC_EED18848.1         LFDTVVLRPTH------------------EEKKKGEWVVEE-DPETILHSVHVDDCGDAY
Q5B221Epimerase           AFAPVLVRPTNVYGRSASYYRGFFEVAVLAKKQSLPLQIPV-SPASICHALHVDDCGDAY
A0A5N7CF08|Epimerase      VFSPVVVRPTNVYGRSASYYRGFFEAAAQSANTKQPLLIPV-PPSSICHALHVDDCGDAY
A0A5N7ASD9|Epimerase      VFSPIVVRPTNVYGRSASYYRGFFEAAAQSINTQLPLLIPV-LPSSICHALHVDDCGDAY
A0A5N7ELR8|Epimerase      VFSPVVVRPTNVYGRSASYYQGFFEAAAQSADAKQPLLIPV-PPSSICHALHVDDCGDAY
A0A7G5JVQ3|Epimerase      VFSPVVVRPTNVYGRSASYYRGFFEAAAQSVDTRQPLLIPV-PPNSICHALHVDDCGDAY
B8N4H7|Epimerase          VFSPVVVRPTNVYGRSASYYRGFFEAAAQSVDTRQPLLIPV-PPNSICHALHVDDCGDAY
I8ACV4|Epimerase          VFSPIVVRPTNVYGRSASYYRGFFEAAAQSVDTRKPLLIPV-PPNSICHALHVDDCGDAY
Q2UM63|Epimerase          VFSPVVVRPTNVYGRSASYYRGFFEAAAQSVDTRKPLLIPV-PPNSICHALHVDDCGDAY
                           *    :***:                                * .* *  *****. **
```

Figure S32: A section from a multiple sequence alignment produced using MUSCLE[1] of various protein sequences from the order Eurotiales (which contains *Talaromyces*) which have the Interpro domain PTHR43725:SF29. An asterisk (*) marks completely conserved amino acid residues, whereas a colon (:) identifies highly conserved residues and a period (.) indicates moderately conserved residues. EED18848.1 has a read through intron (highlighted in red) which when removed (EED18848.1mod), aligns better with other similar sequences.

```
>EED18848.1mod-TsRbtC (rubratoxin)
MASPSINVLVTGANGYIGNAVARAFVRAGYRTYGLVRQPKALPALASAEIIPILGSPEDVSFLHSVSAEGIVFSIIVSTTTTEGVSDYIPHYNAIISLF
RALATTSNAAGIRPLVFFTSGCKDYGRTALANSPGLVPHTELTPLNPQPSLVNRASYAVKTFENKDLFDTVVLRPTHVYGYMSSFYSFFFRFAQEEKKK
GEWVVEEDPETILHSVHVDDCGDAYVAIAQSKREVVANQCYNISAREYETLDQVLKALVKEYGIQGGLKYAKNEGKPGPRAKLFGWSQWVESEKLRRDT
GWTDKRMIFSEGIKQYRVAYESAVETGDEGLKKVLMKVAGRAASQK
```

Figure S33: Modified sequence for TsRbtC from the *T. stipitatus* 2 BGC, EED18848.1mod

The *I. aleurites* genomic sequence (CAJPDT010000158.1: 27086 to 28524, which includes the first half of the fused CAF9941818.1 sequence) was submitted to FGENESH[6] using the *Oidiodendron maius* gene-finding parameters (both *I. aleurites* and *O. maius* are in the Leotiomyceta clade). The resultant protein sequence (CAF9941818.1_a_mod, Figure S34) contains the SnoaL-like domain PF13577.

```
>Ia_CAF9941818.1_a_mod
MHNPLAVCSFLLLLPLSPSNAQQPLNRSLDLAIVEIQNTLNTFSIAVDTHNFTLLSDVFTPDATANFDDSHGELAGLVSISAVLQTGLEGLRSWHALST
QVVAFSGPREARTTSYLQGTFFGQGNLTGQIFTTYGRYLDTLELTGLGWRITRKMLARTGSIGNVKILG
```

Figure S34: Modified sequence CAF9941818.1_a_mod from the *I. aleurites* BGC.

The P450 from the *I. aleurites* BGC (CAF9941819.1) appears to be missing the C-terminus (Figure S35). FGENESH analysis of the sequence CAJPDT010000158.1: 28255 – 32114 from the *I. aleurites* genome, using the using the *Oidiodendron maius* gene-finding parameters (both *I. aleurites* and *O. maius* are in the Leotiomyceta clade) produces a protein sequence (CAF9941819.1mod, Figure S36) which aligns better with other maleidrides and characterised P450s.

```
                                          ExxR
TfR1_P450                     IKNMKYLQYTLKEVNRLCPIVPVNARAAVRDTTLPVGGGPDGKSPIFVKTGQTVNYQIYT
Ts2_EED18842.1_P450           IKNMKYLQYTLKEVNRLYPIVPFNARAAVRDTTLPVGGGSDGKSPILVKAGQAINYQIYT
Pv_ASK38704.1_P450            CNKL---DSFCKESQRFCPPGLIVMSRK-------IMSDIPMSNGTILPKGLFVATSNYD
Ag_OJJ86351.1_P450            TAKLRYQGAVLNESMRIKPAAPTSQPRL-------IPGKGEMIAGRWVPGGCRVAVPPLC
TbR2_P450                     CNAMPYMNAVIQEGLRVFPPAPCKFPRR-------TGHQGATIDGHYIPADTSVGVHQLA
Ag_OJJ86356.1_P450            AQQLPYLRAVIEEALRLYPPVPSRFPRR-------TGKGGAVIDGHFVPENVSVGTHQFA
Ia_CAF9941819.1mod_P450       VNKLPYLNACVKESLRLYPPVPARFPRR-------TASGHEVIDGHIIPVNTSIGVHQWA
Ia_CAF9941819.1               VNKLPYLNACVKESLRLYPPVPARFPRR-------TASGHEVIDGHIIPVNVSSMVD---
                                 :       :*  *. *                            :  .

                                                                          FxxGxRxCxG
TfR1_P450                     MHRRKDLYGEDALEFVPERWEHIRPTWQY-------------LPFNAGPRICIGQQFALT
Ts2_EED18842.1_P450           MHRRKDLYGEDALEFKPERWEHIRPTWQY-------------LPFNAGPRICIGQQFALT
Pv_ASK38704.1_P450            ATSDESVLGN-PDQFDAFRYERMRLQPQQRNLHQLVSTSTSELSFGFGTHACPGRFFAAF
Ag_OJJ86351.1_P450            MNHSPQFFWE-PLSFRPERWLD-NP--QFAS-----DNKAAFQPFSIGPRSCIGRTLALN
TbR2_P450                     AFHSPQNFHR-PLDFVPERWLE-NPPEEFRN-----DALDAVWPFSTGPRNCIGKPMAYM
Ag_OJJ86356.1_P450            TYQSSANFHN-PDQFIPERWLG-DAPKEYTG-----DVKASVQPFSMGPRNCLGKNFAYL
Ia_CAF9941819.1mod_P450       TYHSATNFHE-PDSYLPERWLD-DAPPEFKN-----DALNAVQPFSTGPRICIGKNMAWL
Ia_CAF9941819.1               ------------------------------------------------------------
```

```
TfR1_P450                    EASYTIIRLLQAFKSIRPREGEGSLSELL--TLTTAVRGGVHVGLTPA-------
Ts2_EED18842.1_P450          EASYTIIRLLQAFKSIRPREGEGPLTELL--ALTSSVRGGVNIGLTPA-------
Pv_ASK38704.1_P450           EIKMILIYLLLNYDLKFQEGVPPPRNEILVTAVMPSFQGKVMMKRRREKIGWHVD
Ag_OJJ86351.1_P450           ELKVVLARLLWNFDISLAVKSCDWADKLK--VYGLWEMKPFYIYLDPVKRG----
TbR2_P450                    QMRHTLSRLILYFDMELYPRDDNW-SDVP--ADLFWKMKPLMVRLSPRKQ-----
Ag_OJJ86356.1_P450           QIRVILCKLLWHFDLELAPQSAKW-NEQN--AHLLWEMKPLMVKLSHRKF-----
Ia_CAF9941819.1mod_P450      QMRSILCRLILNFEMELCKESESW-NDQK--SRLLWEMPPLNVKLSHRET-----
Ia_CAF9941819.1              -------------------------------------------------------
```

Figure S35: A section from a multiple sequence alignment produced using MUSCLE[1] of maleidride P450s. An asterisk (*) marks completely conserved amino acid residues, whereas a colon (:) identifies highly conserved residues and a period (.) indicates moderately conserved residues. The deposited sequence from the *I. aleurites* BGC (CAF9941819.1) appears truncated, and does not contain the FxxGxRxCxG motif which is involved in heme-binding.[11] [12] The modified sequence, CAF9941819.1mod, aligns better with other P450s.

```
>Ia_CAF9941819.1mod
MASPVYESQSVVVGGFLALVSRIQARRTPSLLMFVAIKVLIYQISRILYNIFLHPLARFPGPLLRSGFYIFNYWEELRGVQAKKALALHDRYGPVVRIG
PDSLSFNTANAWKDIYAVKPGKPEIPKDMGFFTQNTNKVPSILVCNHEDHVRIKRLVAHAFSDSALRNQEPLMTKYIELLINQLKKKVRAGQGTQDILS
WYMFTTFDIMSDLCFAEPMNALAIGAYQPWIVTILNAAKNGSYVRMERAYPLLAGVSQLYKMMFSGPSKLNTSRAEHMRYSIQKTESRMNNSMERNDIM
TPILQHNSEKGMSRAEIAQTLILFMTGGTEPTASGLSGVTYNVLHNRRVYDKLVKEVREAFSSDSEVNNVAVNKLPYLNACVKESLRLYPPVPARFPRR
TASGHEVIDGHIIPVNTSIGVHQWATYHSATNFHEPDSYLPERWLDDAPPEFKNDALNAVQPFSTGPRICIGKNMAWLQMRSILCRLILNFEMELCKES
ESWNDQKSRLLWEMPPLNVKLSHRET
```
Figure S36: Modified sequence CAF9941819.1mod from the *I. aleurites* BGC.

### Comparative bioinformatics

Table S1: Comparison of the proteins encoded within maleidride BGCs depicted in Figure 4, which identifies core (present in all confirmed maleidride BGCs), common (present across two or more BGCs) and unique proteins. Protein sequences which are not likely to encode catalytic activities (e.g: transporters and regulators) are not included. Domains identified for each set of homologues are shown, except for the PKSs which contain multiple domains. Compounds that have been definitively linked to a specific BGC are depicted in black. BGCs linked to specific compound production through either sequencing of a known producer, or through clear evidence of an orthologous BGC (Ts2) are depicted in blue. Bf: *Byssochlamys fulva*. Pv: *Paecilomyces variotii.* Ts2: *Talaromyces stipitatus* BGC 2. Sa: *Scytalidium album.* DC: *Diffractella curvata.* Zc: *Zopfiella curvata.* Wa: *Wicklowia aquatica.* Phi: Unidentified phomoidride producer ATCC 74256.

Proteins with 'mod' included after their accession number are instances where annotations have been updated during this work. See 'Reannotation of sequences' for details and justifications.

'a' denotes predicted proteins which have been newly identified and annotated during this work. Protein sequences can be acquired from the gbk files in the 'Genbank files' section. The accession number for epiheveadrides BGC is OK490366.

| | | | **bysso/ agnestadrides** | **cornexistin** | **Scytalidin** | **Zopfiellin** | **Zopfiellin** | **rubratoxins** | **Epiheveadrides** | **phomoidrides** |
|---|---|---|---|---|---|---|---|---|---|---|
| **Type** | **Predicted function or domain** | **Interpro domain** | **Bf** | **Pv** | **Sa** | **Dc** | **Zc** | **Ts2** | **Wa** | **Phi** |
| **Core** | PKS | | **Bfpks1** ANF07288.1 | **Pvpks1** ASK38717.1 | **ScyPKS** QTE76000.1mod | **ZopPKS** QTE75992.1 | **ZopA** BBU42026.1 | **TsRbtJ** EED18841.1 | **Epipks**[a] | **PhiA** BBG28498.1 |
| **Core** | Hydrolase | PTHR48070:SF4 | **BfL1** ANF07287.1 | **PvL1** ASK38716.1 | **ScyR1** QTE76001.1 | **ZopR1** QTE75993.1 | **ZopM** BBU42027.1 | **TsRbtS** EED18832.1 | **EpiR11**[a] | **PhiM** BBG28510.1mod |
| **Core** | ACS | PTHR11739:SF4 | **BfL2** ANF07286.1 | **PvL6** ASK38711.1 | **ScyR3** QTE76003.1 | **ZopR3** QTE75995.1 | **ZopJ** BBU42028.1 | **TsRbtL** EED18839.1 | **EpiR9**[a] | **PhiJ** BBG28507.1 |
| **Core** | ACDH | PTHR16943:SF22 | **BfL3** ANF07285.1 | **PvL2** ASK38715.1 | **ScyR7** QTE76007.1 | **ZopL5** QTE75987.1 | **ZopI** BBU42021.1 | **TsRbtK** EED18840.1 | **EpiR8**[a] | **PhiI** BBG28506.1 |
| **Core** | PEBP1 | PTHR11362 | **BfL5** ANF07283.1 | **PvR1** ASK38718.1 | **ScyL1** QTE75999.1 | **ZopL1** QTE75991.1 | **ZopN** BBU42025.1 | **TsRbtO** EED18836.1 & **TsRbtM** EED18838.1 | **EpiR12**[a] | **PhiN** BBG28511.1mod |
| **Core** | MDC | PTHR31779:SF6 | **BfL6** ANF07282.1 & **BfL10** ANF07278.1 | **PvL3** ASK38714.1 | **ScyR6** QTE76006.1 | **ZopL4** QTE75988.1 | **ZopC** BBU42022.1 | **TsRbtR** EED18833.1 | **EpiR1**[a] **EpiR6**[a] | **PhiC** BBG28500.1 |
| **Common** | Maleidride conserved protein | | **BfL8** ANF07280.1 | **PvL16** ASK38701.1 | **ScyR11** QTE76011.1 | **ZopL8** QTE75984.1 | **ZopP** BBU42018.1 | | **EpiR3**[a] | |
| **Common** | PEBP2 | PTHR30289:SF1 | **BfL9** ANF07279.1 | | **ScyR12** QTE76012.1 | | | | **EpiR4**[a] | **PhiB** BBG28499.1 |
| **Common** | Enoyl CoA isomerase-like | IPR029045 | **BfL11** ANF07277.1 | | **ScyR2** QTE76002.1 | **ZopR2** QTE75994.1 | **ZopS**[a] | | **EpiR10**[a] | |
| **Unique** | αKGDD-IPNS-like | IPR027443 | | PvL5 ASK38712.1 | | | | | | |
| **Common** | 6-bladed beta propeller | PTHR42060 | | **PvL7** ASK38710.1 | | | | | | |
| **Unique** | Carboxypeptidase | PTHR11802:SF189 | | **PvL9** ASK38708.1 | | | | | | |
| **Common** | Dienelactone hydrolase | PTHR47668 | | **PvL11** ASK38706.1 | | | | | | |
| **Unique** | P450 | PTHR46206 | | **PvL13** ASK38704.1 | | | | | | |
| **Unique** | Transketolase | PTHR43825 | | **PvL14** ASK38703.1 | | | | | | |
| **Common** | Aldo ketoreductase | PTHR11732:SF461 | | **PvL15** ASK38702.1 | | | | | **EpiR14**[a] | |

| | | | | | | | | | |
|---|---|---|---|---|---|---|---|---|---|
| **Unique** | Hypothetical | | **PvL17** ASK38700.1 | | | | | | |
| **Common** | αKGDD AsaB-like | IPR044053 | | **ScyL2** QTE75998.1 | **ZopL9** QTE75983.1 | **ZopK** BBU42017.1 | **TsRbtB** EED18849.1a_mod & **TsRbtG** EED18844.1 | | **PhiK** BBG28508.1 |
| **Unique** | Oxidoreductase | PTHR44229:SF6 | | **ScyL3** QTE75997.1 | | | | | |
| **Common** | Isochorismatase | IPR036380 | | **ScyR4** QTE76004.1 | **ZopL2** QTE75990.1 | **ZopQ** BBU42024.1 | **TsRbtQ** EED18834.1mod | | |
| **Unique** | Esterase/transferase | PTHR43283:SF3 | | | **ZopL10** | **Orf1** BBU42016.1 | | | |
| **Unique** | FAD oxidoreductase | PTHR42973:SF22 | | | **ZopR4** QTE75996.1 | **Orf-1** BBU42029.1 | | | |
| **Unique** | FAD binding domain | PTHR13878:SF70 | | | | | **TsRbtA** EED18850.1 & **TsRbtD** EED18847.1 | | |
| **Common** | Thioesterase | PTHR47260 | | | | | **TsRbtB2** EED18849.1_b_mod | | |
| **Unique** | NAD dependent Epimerase dehydratase | PTHR43725:SF29 | | | | | **TsRbtC** EED18848.1mod | | |
| **Common** | αKGDD TauD-like | IPR042098 | | | | | **TsRbtE** EED18846.1 & **TsRbtU** EED18830.1mod | | |
| **Common** | SnoaL-like domain | PF13577 | | | | | **TsRbtF** EED18845.1mod | | |
| **Unique** | Ferric reductase | PTHR32361 | | | | | **TsRbtH** EED18843.1mod | | |
| **Common** | P450 | PTHR24287 | | | | | **TsRbtI** EED18842.1 | | |
| **Common** | AMP dependent CoA ligase | PTHR24096:SF312 | | | | | **TsRbtN** EED18837.1 | **EpiR13** [a] | |
| **Unique** | α/β hydrolase | PTHR23024:SF210 | | | | | | **EpiR15** [a] | |
| **Common** | AusD-like methyltransferase | PTHR35897 | | | | | | | **PhiE** BBG28502.1 |
| **Unique** | DUF1115 | PTHR15955 | | | | | | | **PhiF** BBG28503.1 |
| **Unique** | α/β hydrolase | PTHR43194:SF4 | | | | | | | **PhiG** BBG28504.1 |
| **Unique** | Conserved Hypothetical | PTHR33048:SF134 | | | | | | | **PhiH** BBG28505.1 |

Table S2: Comparison of the proteins encoded within orphan maleidride BGCs depicted in Figure 5, which identifies core (present in all confirmed maleidride BGCs), common (present across two or more BGCs) and unique proteins. Protein sequences which are not likely to encode catalytic activities (e.g: transporters and regulators) are not included. Domains identified for each set of homologues are shown, except for the PKSs which contain multiple domains. Ag: *Aspergillus glaucus.* BmC5: *Bipolaris maydis* C5 (only one *Bipolaris* BGC has been included, as all 4 are almost completely orthologous. Sequences which are uniquely present within *Bipolaris* BGCs are categorised as 'Bipolaris'. Further information on the *Bipolaris* BGCs can be found in Figure S40 and Table S3.) Ca: *Cadophora* sp. Czm: *Cercospora zeae-maydis.* Cc: *Coleophoma crateriformis.* Ia: *Imshaugia aleurites.* Om: *Oidiodendron maius* Zn. Ta: *Talaromyces atroroseus.* Tb: *Talaromyces borbonicus.* Tf: *Talaromyces funiculosus.* Ts1: *Talaromyces stipitatus* BGC 1. * NAPEPLD: N-acyl-phosphatidylethanolamine-hydrolysing phospholipase D.

Proteins with 'mod' included after their accession number are instances where annotations have been updated during this work. See 'Reannotation of sequences' for details and justifications.

'a' denotes predicted proteins which have been newly identified and annotated during this work. Protein sequences can be acquired from the gbk files in the 'Genbank files' section. 'b' Ts1R1 does not contain IPR029045, however it does contain the related 3.90.226.10 domain, which places it with other maleidride enoyl CoA isomerases.

| Type | Predicted function or domain | Interpro domain | Ag | BmC5 | Ca | Czm | Cc | Ia | Om | Ta | Tb | Tf | Ts1 |
|---|---|---|---|---|---|---|---|---|---|---|---|---|---|
| **Core** | PKS | | OJJ86349.1 | EMD85570.1 | PVH77199.1 & PVH77205.1 | KAF2215724.1 | RDW56971.1 | CAF9941811.1mod | KIN05356.1 & KIN05364.1 | OKL57046.1 | TbPKS1 [a] | TfPKS1 [a] | EED15402.1 |
| **Core** | Hydrolase | PTHR48070:SF4 | OJJ86354.1 | EMD85581.1 | CaL1 [a] | CzmL1 [a] | RDW56968.1 | CAF9941818.1_b_mod | KIN05363.1 | OKL57045.1 | TbR3 [a] | TfL9 [a] | EED15412.1mod |
| **Core** | ACS | PTHR11739:SF4 | OJJ86352.1 | EMD85580.1 | PVH77201.1 | KAF2215727.1 | RDW56963.1 | CAF9941815.1 | KIN05361.1 | OKL57051.1 | TbR4 [a] | TfL2 [a] | EED15409.1 |
| **Core** | ACDH | PTHR16943:SF22 | OJJ86350.1 | EMD85579.1 | PVH77202.1 | KAF2215728.1 | RDW56964.1 | CAF9941813.1 | KIN05360.1 | OKL57050.1 | TbR1 [a] | TfL1 [a] | EED15410.1mod |
| **Core** | PEBP1 | PTHR11362 | | EMD85582.1 | PVH77197.1 | KAF2215738.1 | RDW56967.1 | | | OKL57043.1 | | TfL5 [a] & TfL3 [a] | EED15411.1 |
| **Core** | MDC | PTHR31779:SF6 | OJJ86357.1 | EMD85577.1 & EMD85571.1 | PVH77203.1 | KAF2215733.1 & KAF2215732.1 | RDW56973.1 | CAF9941823.1mod | KIN05369.1 | OKL57228.1 | TbL7 [a] | TfL8 [a] | EED15406.1 |
| **Common** | Maleidride conserved protein | | | EMD85573.1 | | KAF2215736.1 | | | | TaR7 [a] | | | |
| **Common** | PEBP2 | PTHR30289:SF1 | | | | | RDW56972.1 | | | | | | EED15405.1mod |
| **Common** | Enoyl CoA isomerase-like | IPR029045 | OJJ86355.1 | | | | RDW56966.1 | CAF9941821.1 | | | TbL5 [a] | | Ts1R1 [a,b] |
| **Common** | 6-bladed beta propeller | PTHR42060 | | | | | | CAF9941807.1 | | | TbL10 [a] | | |
| **Common** | Dienelactone hydrolase | PTHR47668 | | | | | | CAF9941809.1 | | | TbL9 [a] | | |
| **Common** | αKGDD AsaB-like | IPR044053 | | | | KAF2215734.1 | RDW56970.1 | | | OKL57227.1 | | | EED15414.1 |
| **Common** | Isochorismatase | IPR036380 | | EMD85567.1 | PVH77207.1 | KAF2215730.1 & KAF2215740.1 | | | KIN05357.1 | OKL57229.1 | | TfL7 [a] | |
| **Common** | Thioesterase | PTHR47260 | | | | | | | | | | TfL10 [a] | |
| **Common** | αKGDD TauD-like | IPR042098 | | | PVH77204.1 | | | | KIN05358.1 | | | TfL12 [a] | |
| **Common** | SnoaL-like domain | PF13577 | | | | | | CAF9941818.1_a_mod | | | | | |
| **Common** | P450 | PTHR24287 | | | | | | | | | | TfR1 [a] | |
| **Common** | AMP dependent CoA ligase | PTHR24096:SF312 | | EMD85583.1 | PVH77196.1 | KAF2215735.1 | | | KIN05362.1 | OKL57044.1 | | TfL4 [a] | |

| | | | | | | | | | | | | |
|---|---|---|---|---|---|---|---|---|---|---|---|---|
| **Common** | AusD-like methyltransferase | PTHR35897 | OJJ86353.1 | | | | RDW56962.1 | CAF9941817.1 | | | TbR5[a] | EED15408.1 |
| **Common** | P450 | PTHR24305:SF113 | OJJ86351.1 & OJJ86356.1 | | | | | CAF9941819.1 | | | TbR2[a] | |
| **Unique** | Oxidoreductase | PTHR24320:SF154 | OJJ86359.1 | | | | | | | | | |
| **Bipolaris** | Dienelactone hydrolase | PTHR17630:SF76 | | EMD85572.1 | | | | | | | | |
| **Bipolaris** | Reductase | PTHR43377:SF2 | | EMD85584.1 | | | | | | | | |
| **Unique** | CIPA-LIKE | PTHR47706:SF1 | | | PVH77194.1 | | | | | | | |
| **Common** | Quinone reductase | PTHR32332:SF28 | | | | KAF2215726.1 | | | | OKL57052.1 | | |
| **Unique** | Alcohol dehydrogenase | PTHR43669:SF3 | | | | KAF2215731.1 | | | | | | |
| **Unique** | HeLo domain | IPR038305 | | | | | | | KIN05366.1 | | | |
| **Unique** | ATP/GTP-binding protein-related | PTHR46082:SF6 | | | | | | | KIN05367.1 | | | |
| **Unique** | ATPase | PTHR46411 | | | | | | | KIN05365.1 | | | |
| **Unique** | Epoxide hydrolase | PTHR21661 | | | | | | | KIN05368.1 | | | |
| **Unique** | NAPEPLD* | IPR024884 | | | | | | | | OKL57047.1 | | |
| **Unique** | αKGDD PhyH-like | IPR008775 | | | | | | | | OKL57048.1 | | |
| **Unique** | 2-hydroxyacyl-CoA lyase | PTHR43710:SF2 | | | | | | | | | TbL2[a] | |
| **Unique** | Peroxisomal-CoA synthetase | PTHR43201:SF5 | | | | | | | | | TbL1[a] | |
| **Unique** | Phosphonomutase | PTHR42905:SF12 | | | | | | | | | TbL3[a] | |
| **Unique** | Acyltransferase | PTHR31625:SF26 | | | | | | | | | TbL8[a] | |
| **Unique** | Long-chain-fatty-acid-CoA ligase | PTHR43272:SF33 | | | | | | | | | TbR6[a] | |

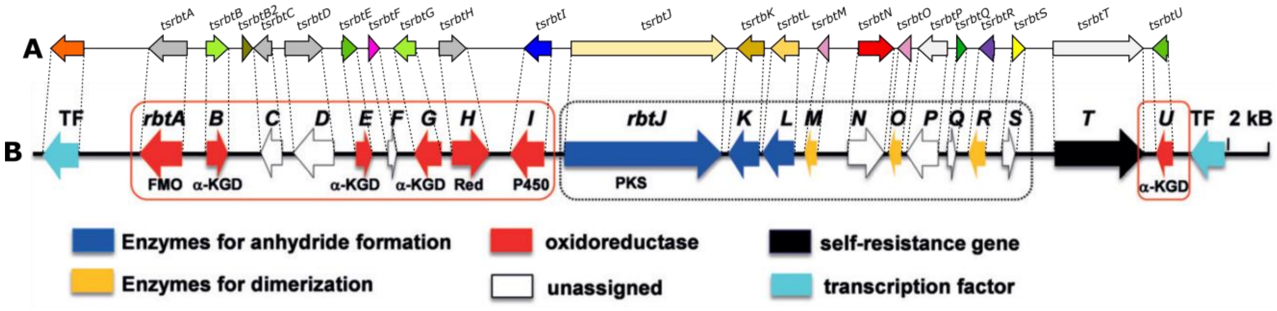

Figure S37: A likely rubratoxin BGC (**A**) was identified within the publicly available genome of *Talaromyces stipitatus*. Although not experimentally confirmed, *Talaromyces* fungi are known producers of rubratoxins, and the gene cluster demonstrates total synteny with the confirmed rubratoxin BGC from *Penicillium dangeardii* Pitt (**B**). The image of the cluster from *Penicillium dangeardii* Pitt has been reproduced with permission from reference Bai *et al*.[10]

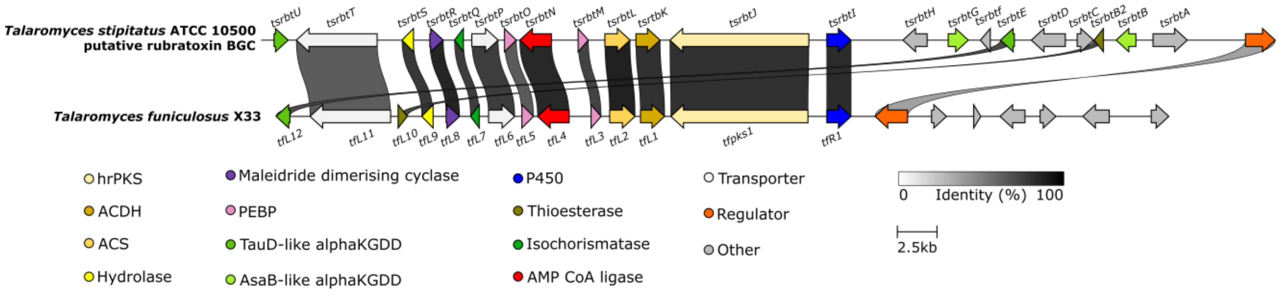

Figure S38: A clinker[13] comparison of the putative rubratoxin BGC with a semi-orthologous cluster from *Talaromyces funiculosus*. Only the best links between homologous genes are shown, according to their percentage identity (see identity scale bar).

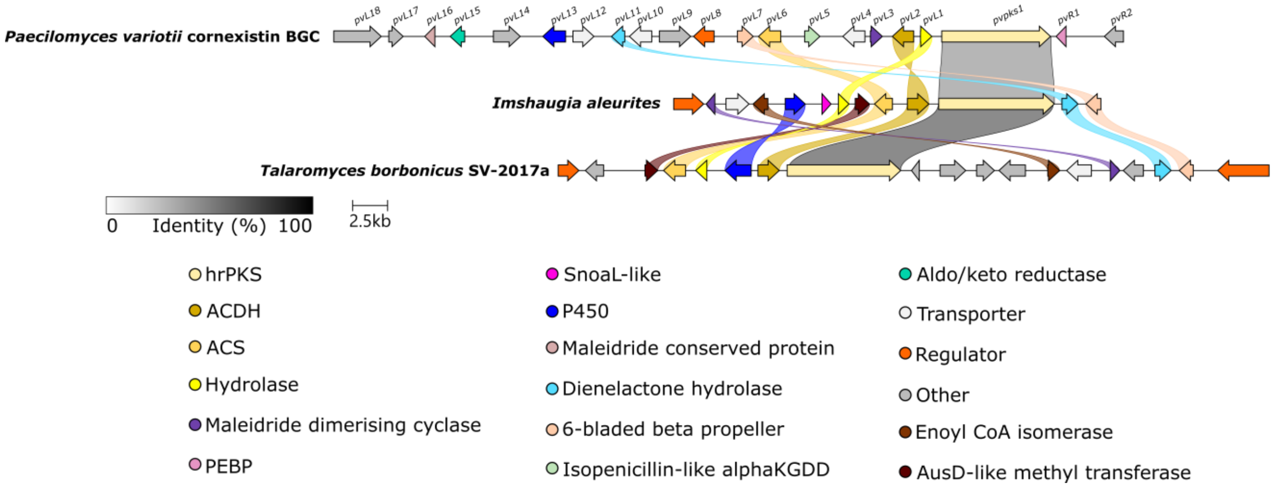

Figure S39: A clinker[13] comparison of the cornexistin BGC with orphan maleidride BGCs identified from the genomes of *I. aleurites* and *T. borbonicus.* The mininum identity was set at 25%. Links between homologous genes are shown using their specific colour, except for the PKSs where the links are shown according to the percentage identity (see identity scale bar). Links between transport and regulatory genes have been removed for clarity.

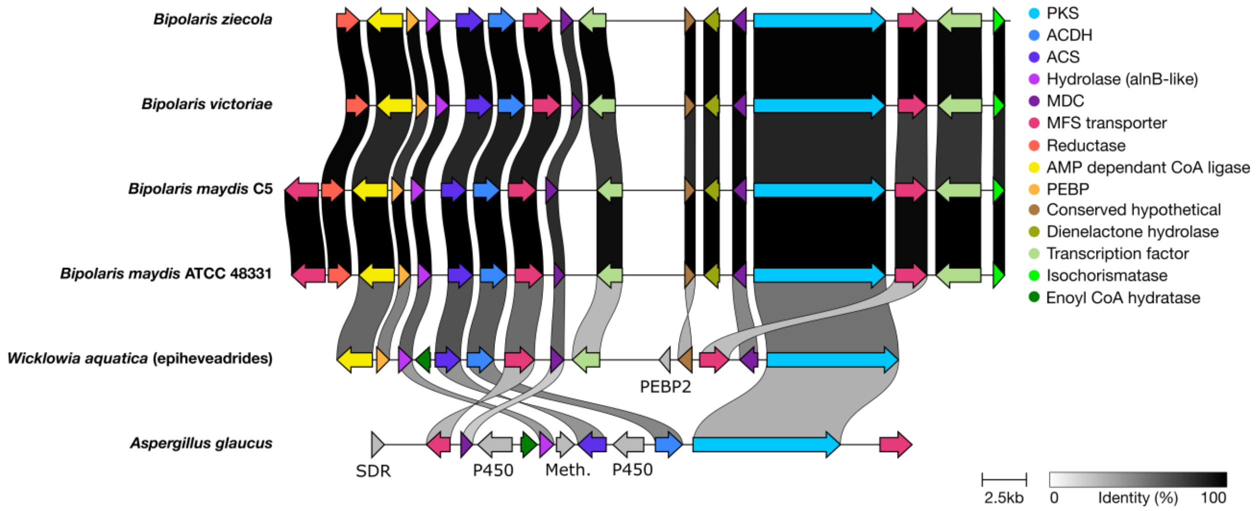

Figure S40: A clinker[13] comparison of the *Wicklowia aquatica* maleidride BCG (a confirmed epiheveadride producer) with putative maleidride BGCs from a range of *Bipolaris* species and *Aspergillus glaucus*. Links between proteins demonstrate % identity higher than 30 %, with darker links denoting higher % identity (see key at bottom right). See Table S3 for further details.

Table S3: Proteins encoded by the likely epiheveadride BGC of *Wicklowia aquatica*, compared to the newly identified maleidride BGCs of various *Bipolaris* species and *Aspergillus glaucus*. Homologues were identified by clinker analysis (See Figure S40), and percentage identity between *W. aquatica* proteins and their homologues are shown in brackets.

| Putative function | Domain | *W. aquatica* | *B. maydis* C5 | *B. maydis* ATCC 48331 | *B. victoriae* | *B. ziecola* | *A. glaucus* |
|---|---|---|---|---|---|---|---|
| Polyketide synthase | | EpiPKS | EMD85570.1 (63 %) | XP_014073936.1 (63 %) | XP_014553689.1 (63 %) | XP_007715650.1 (63 %) | OJJ86349.1 (41 %) |
| AlnB-like esterase | PTHR48070:SF4 | EpiR11 | EMD85581.1 (70 %) | XP_014073929.1 (70 %) | XP_014553701.1 (70 %) | XP_007715667.1 (71 %) | OJJ86354.1 (48 %) |
| Alkylcitrate synthase | PTHR11739:SF4 | EpiR9 | EMD85580.1 (74 %) | XP_014073930.1 (74 %) | XP_014553700.1 (70 %) | XP_007715666.1 (71 %) | OJJ86352.1 (51 %) |
| Alkylcitrate dehydratase | PTHR16943:SF22 | EpiR8 | EMD85579.1 (71 %) | XP_014073931.1 (71 %) | XP_014553699.1 (70 %) | XP_007715665.1 (71 %) | OJJ86350.1 (56 %) |
| PEBP1 | PTHR11362 | EpiR12 | EMD85582.1 (58 %) | XP_014073928.1 (58 %) | XP_014553702.1 (61 %) | XP_007715668.1 (59 %) | - |
| Maleidride dimerising cyclase | PTHR31779:SF6 | EpiR1<br>EpiR6 | EMD85571.1 (56 %)<br>EMD85577.1 (74 %) | XP_014073935.1 (56 %)<br>XP_014073932.1 (67 %) | XP_014553690.1 (58 %)<br>XP_014553697.1 (67 %) | XP_007715659.1 (58 %)<br>XP_007715663.1 (71 %) | OJJ86357.1 (31 %) |
| Conserved hypothetical | | EpiR3 | EMD85573.1 (41 %) | XP_014073934.1 (41 %) | XP_014553692.1 (42 %) | XP_007715652.1 (42 %) | - |
| MFS Efflux 1 | | EpiR7 | EMD85578.1 (65 %) | XP_014073913.1 (65 %) | XP_014553698.1 (65 %) | XP_007715664.1 (65 %) | OJJ86358.1 (39 %) |
| MFS Efflux 2 | | EpiR2 | EMD85569.1 (35 %) | XP_014073937.1 (36 %) | XP_014553688.1 (37 %) | XP_007715658.1 (37 %) | - |
| MFS transporter | | - | EMD85585.1 | XP_014073925.1 | - | - | - |
| TF 1 | | EpiR4 | EMD85574.1 (39 %) | XP_014073933.1 (38 %) | XP_014553696.1 (41 %) | XP_007715661.1 (41 %) | - |
| TF 2 | | - | EMD85568.1 | XP_014073910.1 | XP_014553688.1 | XP_007715657.1 | - |
| AMP dependant CoA ligase | PTHR24096:SF312 | EpiR13 | EMD85583.1 (67 %) | XP_014073927.1 (67 %) | XP_014553703.1 (67 %) | XP_007715669.1 (67 %) | - |
| Isochorismatase | IPR036380 | - | EMD85567.1 | XP_014073915.1 | XP_014553686.1 | XP_007715656.1 | - |
| Reductase | PTHR43377:SF2 | - | EMD85584.1 | XP_014073926.1 | XP_014553704.1 | XP_007715670.1 | - |
| Dienelactone hydrolase | PTHR17630:SF76: | - | EMD85572.1 | XP_014073908.1 | XP_014553691.1 | XP_007715660.1 | - |
| PEBP2 | PTHR30289:SF1 | EpiR4 | - | - | - | - | - |
| Enoyl CoA hydratase/isomerase | IPR029045 | EpiR10 | - | - | - | - | OJJ86355.1 (23 %) |
| P450 | PTHR24305:SF113 | - | - | - | - | - | OJJ86351.1 &<br>OJJ86356.1 |
| Methyltransferase | PTHR35897 | - | - | - | - | - | OJJ86353.1 |
| Oxidoreductase (SDR) | PTHR24320:SF154 | - | - | - | - | - | OJJ86359.1 |

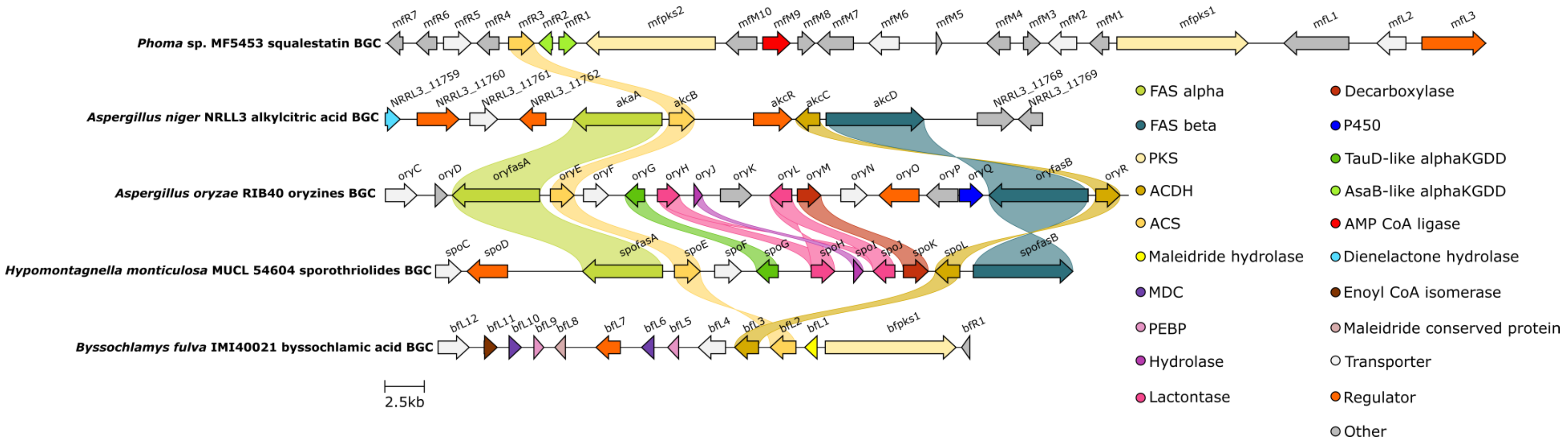

Figure S41: A clinker[13] comparison of the byssochlamic acid BGC with non-maleidride alkylcitrate based biosynthetic pathways. The mininum identity was set at 25%. Links between homologous genes are shown using their specific colour. Links between transport and regulatory genes have been removed for clarity.Only the alkylcitrate synthase is common between all BGCs.

### Identification of conserved active site or binding residues

```
                                                                                     MOTIF I
hrPKS_Fum1p(W7LKX1.1)         VKGELEGIELLQAEDGLTNYY-----NYVESRTDSIDFFATAGHTRPTLRVLEIGAGTGG
hrPKS-mlcB(Q8J0F5.1)          LRRETEPLELMMQDQLLSRYY-VNAIKWSRSNAQASELIRLCAHKNPRSRILEIGGGTGG
hrPKS-LDKS(Q9Y7D5.1)          LRREVDPLEVMMDGHLLSRYY-VDALKWSRSNAQASELVRLCCHKNPRARILEIGGGTGG
KAF2215724.1(Czm)             LLGQIDTRRHILDCGALELYRSSQTPHLSSLYGALARYLALLGHRDPHMSVIEINAGKGS
OJJ86349.1(Ag)                LRQEVEPLQLMVEEDRLEDYY-RDNTRFDRNYRQVARYLEALAHKNPAMSILEVGAGTGG
PhiA_BBG28498.1               VAGESSP-SDLVQGLDFNAFV-EDPHLFQ-NTRSAATYLDLVGHKNPNLSILTIGPQSGL
EED15402.1(Ts1)               VKGENSS-AEFIKELDLKVFV-DNTQLFQ-NTQSAARYFDLLQHKTPSLSVLAVGPGSGV
RDW56971.1(Cc)                LTSETAY-PEIVRSFDLGRLY-APSGIFL-NCKSAARYIDLLKHKSPNLSIMAVGDLAGV
TSrbtJ_EED18841.1(Ts2)        LRGELEP-SSLLSEIELSGFF-SSSDTFI-NSSTVAKYLELLAFKQPNVSILACGKLSGF
TfPKS                         LKGEIEA-SSLLSEVDLSGFY-SPSDMFT-DSTMVSKYLEMMAFKQPNLSILACGKLSGF
PVH77205.1(CadPKS2)           LRGELLP-ESLVNEIDIAQFF-GPSHIFA-NSTLVSKFVELLGFKHPNLSILACGRLSGI
KIN05356.1(OmPKS1)            LRGELSP-ASLVNEIDISQFF-SHSDMFI-NHALVAKFIEVLAFKDPSISILACGKLCGF
TbPKS1                        LSREVEALPLMIEDGRLDGYY-RDNHRFDRNYQAAARYINLLGHKNPHLNILEIGAGTGG
CAF9941811.1mod(Ia)           IARKVEALPIMVEQGRLDAYY-RENARFDRNYRAATKYIDLLAHKNPYIKVLEIGAGTGG
scyPKS_QTE76000.1mod          MKKEIDPWTLLTEKQSLEAYF-RDTPRIARTYDAVAKYFYLLGHKNPHLSILQIGTGTGG
ZopPKS_QTE75992.1             MKKEVDPWNLLIGEQRLEAYF-KDSPRIARTYLAAAEYFDLLGHKSPHLSILEIGVGTGG
ZopA_BBU42026.1               MKKEVDPWNLLIGEQRLEAYF-KDSPRIARTYLAAAEYFDLLGHKSPHLSILEIGVGTGG
PvPKS1_ASK38919.1             -----------------------------------ARYVKLLGHKNPHLAILEVGAGQGE
OKL57046.1mod(Ta)             LRDGIDPWNIMMTENRLDAYL-NDTSRLVRNYKIAAKYVRLLGNKNPQLSLLEIGAGTGE
PVH77199.1(CadPKS1)           FQNKVDIWSVITAENRLELYR-SKTPRLVRNYDAAGRYLSLLGHKNPHMSILEVSAGAGS
KIN05352.1(OmPKS2)            FRGKVDMWSVLADENRLESYR-SMTPRFVRNYDATARYLSLLGHKDPHISIIEVGAGAGS
BfPKS1_ANF07288.1             LRQRLDP-STLLERETLHRYI-RESWSANGIHDAAADYLSLLCHKHPDLSVLEIDGGIGS
epiPKS                        ----------------------------------AADYIRLLGHKNPNLSVFDATELQDI
XP_014073936.1(Bm_ATTC)       ----------------------------------AADYLRLLGFKNPEISVLDVSASSSS
EMD85570.1(Bm_C5)             ----------------------------------AADYLRLLGFKNPEISVLDVSASSSS
XP_007715650.1(BZ)            ----------------------------------AAEYIRLLGFKNPEISVLDISASSRS
XP_014553689.1(Bv)            ----------------------------------AAEYIRLLGFKNPEISVLDISASSRS
                                                                    .       *   ::

                                                    MOTIF II
hrPKS_Fum1p(W7LKX1.1)         GAQVILEGL---TNGKERLFSTYAYTDISAGFFV---AAQERFKAYKGL-DFKVLDITKD
hrPKS-mlcB(Q8J0F5.1)          CTKLIVNAL-----GNTKPIDRYDFTDVSAGFFE---SAREQFADWQDVMTFKKLDIESD
hrPKS-LDKS(Q9Y7D5.1)          CTQLVVDSL-----GPNPPVGRYDFTDVSAGFFE---AARKRFAGWQNVMDFRKLDIEDD
KAF2215724.1(Czm)             GTLRFLEAL----NSTAPAIRKYTVTDTSNEDFV---ATEELLQPWKSLVQFSELDIDRD
OJJ86349.1(Ag)                ATLPLLQAL-DRATHQAPCCSRYHFTDISSGFFD---AARERLDAWSGFLDFSKLDIEHD
PhiA_BBG28498.1               ASLGLLSLL-SELEGSTPRFTAFHHTDSELNITD---TVKEKFSAWANLIEFKDLAIHSD
EED15402.1(Ts1)               ASLGFLALL---NKKSSAPFERYEHNDVEFDIRD---VVKEKFPQWAQLIGTKQVDISRE
RDW56971.1(Cc)                TSIPIIAAL-CPSGNETLSFTKFEYVDPDFDLTE---MIKARVPALQNIVMRNELDMGKD
TSrbtJ_EED18841.1(Ts2)        LSIPVLAAL-DSKAGQLPYFSKFEYADDDFDLSD---LVKAKCSRWNGLISRKELDIAVD
TfPKS                         LSIPILAAL-ESKKQQLPYFSKFEYADDDFDLAN---LVKAKCAKWNDLVTRKELDIEVD
PVH77205.1(CadPKS2)           ASIPVLAAL-NSATGGVPCFSKFEYADIQFDLTD---SVKRKIPQWSELISRKELDIEVD
KIN05356.1(OmPKS1)            LVIPVLAAL-NSVTGELPCFSKFEYTDVEFDLSD---LIKGKCPAWSNSISRKELDIEVD
TbPKS1                        ATLPLLQAL-GGTEGDLPRFKNFHFTDISSGFFD---AAKEKLSVWSNLITYGKLDIEKD
CAF9941811.1mod(Ia)           ATLPIIEAI-GGGDAELPRLAEWHFTDISSSFFD---AAKEKLERWNHLVSYAKLDIETD
scyPKS_QTE76000.1mod          ATFPILKSL-GGADGEIPRFQKYDFTDTSNILSE---ELKQKLAPWKDLIAFKELDLNND
ZopPKS_QTE75992.1             ATFPILKSL-GGADGDLPRFQKYDCTDVNSTLFD---SLKERAAPWKDLVTFKELNIDND
ZopA_BBU42026.1               ATFPILKSL-GGADGDLPRFQKYDCTDVNSTLFD---SLKERAAPWKDLVTFKELNIDND
PvPKS1_ASK38919.1             LCIPVFRAL-AGEANSTPSFQSYTLADTEPGLSETIATIADQFDERADLIQYKELDISSD
OKL57046.1mod(Ta)             ASLPILQAL--SEGNTSVWLKRYTFTDLNTDMFE---VAQKKLSDWADLIKFKEFDVAGS
PVH77199.1(CadPKS1)           ATLRILQTI-GGHHGEAPSFEKYTITNPDASTFD---TTSLAFEPWKDCLEFKALDIEGD
KIN05352.1(OmPKS2)            ATFPILQTL-GGANGEVASIKNYTFTDTDTSGFD---SASKILEPWKDFLKFKQLDIEGD
BfPKS1_ANF07288.1             VASKAVPAMMGPEGSDMPNFVQYMLTSMQASRLE---VLKAENAPLINLISIKELNMAED
epiPKS                        -TISILGALKGGDSGTLPRFKTYTLTNSPKSSSS---VTNDALYAWHELISLKEFDVETD
XP_014073936.1(Bm_ATTC)       -ILSFLKALEIGNSAAKQRCRAYTVAHPQRNKAA---LVSDVPDLPYDLVTKKALDLEIG
EMD85570.1(Bm_C5)             -ILSFLKALEIGNSAAKQRCRAYTVAHPQRNKAA---LVSDVPDLPYDLVTKKALDLEIG
XP_007715650.1(BZ)            -TLSFLKALEVGNSAAKQRCRAYTLAHAQGNKMA---LVSNVPNLPYDLVTKKTLDLDLG
XP_014553689.1(Bv)            -TLSFLKALEVGNSAAKQRCRAYTLAHAQGNKMA---LVSDVPNLPYDLVTKKTLDLDLG
                                   .  :             :                                . :
```

```
                                        MOTIF III                                MOTIF IV
hrPKS_Fum1p(W7LKX1.1)        PSEQ-GFESGSFDLIIAGNVIHATPTLNETLANVRKLLAPEGYLFLQE--LSPKMRMVNL
hrPKS-mlcB(Q8J0F5.1)         PEQQ-GFECATYDVVVACQVLHATRCMKRTLSNVRKLLKPGGNLILVE--TTRDQLDLFF
hrPKS-LDKS(Q9Y7D5.1)         PEAQ-GFVCGSYDVVLACQVLHATSNMQRTLTNVRKLLKPGGKLILVE--TTRDELDLFF
KAF2215724.1(Czm)            IEEQ-NFEKHSYDVVIMAHSTLLVKNMVHALRRTRRLLKPNGKLVLVDIVIDEPSLVQTM
OJJ86349.1(Ag)               PALQ-GYEPHSYDLVIAANVLHATRSMQATMANVRKLLKPTGKLVLIE--LTRERLTTSA
PhiA_BBG28498.1              QSEE----NEVYDVVVAFHVLGSSNSLPETFSSSRRLLKPGGKLLLVG--RALKSLVATV
EED15402.1(Ts1)              IQGQEDIEPNSYDVLVAFHVLGDATGMNNVLAQSKQLLKPDGKVLFIG--RPLKSLVASV
RDW56971.1(Cc)               VALQ-NCNLKSYDVVIAFHVIGSAKSTRAIMANAQKLLKPGGKLLLIG--RVMRSLAAAT
TSrbtJ_EED18841.1(Ts2)       PSKQ-DIALGSYDALIVFQSLGSENLVRSTLENAHKLLKPGGKLLLVG--RPMKSLIVST
TfPKS                        PAKQ-DVSLESYDVLIVFQPLGCEEAVRRTLENAHKLLKPGGKLLLVG--RPMKSLAVST
PVH77205.1(CadPKS2)          PASQ-DCALESYHIVIAYCVLESAKSVRSTLQHAHKLLKPGGKLLLIG--RPMESLAEST
KIN05356.1(OmPKS1)           LANQ-DCALESYDVVIAFHALGSAKLVRRTLENAHKLLKSGGKLMLIG--RPMRSLVAST
TbPKS1                       PGSQ-GYELGTYDVVVAANVLHATKSMHNTMSNVRKLLKPGGKLILVE--LTRERMTTST
CAF9941811.1mod(Ia)          PVKQ-GFENESYDVVVAANVLHATKSLHQTLINVRRLLKRGGRLVLIE--LTRERMTTST
scyPKS_QTE76000.1mod         PIAQ-GYTVESYDVILAAHTLRTSKSLHTALGNARRLLKPGGKLVILD--VTRERMAPSL
ZopPKS_QTE75992.1            PVAQ-GFTAESYDVIFAAYALHTSKSLHTALGNARKLLKPGGKLVILD--VTREHMASSL
ZopA_BBU42026.1              PVAQ-GFTAESYDVIFAAYALHTSKSLHTALGNARKLLKPGGKLVILD--VTREHMASSL
PvPKS1_ASK38919.1            PLQQ-GFNAHSLDLILLPSRGVSATLRSKILKHAHQLLTPEGRLIVVD------------
OKL57046.1mod(Ta)            LEEQ-GFKSHSYDVVILGHGIHLAKSTDRMLKNIRNLLKDEGKFIFVDEVYQNESIERSL
PVH77199.1(CadPKS1)          LHDQ-GYDSNRFDIVILAHGTHMAKSRGAVLKNVHTLLNSNGRLILIDTVAESQSIAQKM
KIN05352.1(OmPKS2)           LDDQ-GYDSNSFDVVIMAHGMYMSMSRDAVLKNVHALLKPNGRLILIDIIAEGESMAGSM
BfPKS1_ANF07288.1            PIRQ-GFKEASFDVIIAPAQIAANASSVAVLRNVKKLLKNKGRLIILG--TACIQPSLLL
epiPKS                       LASE-GFQLNTYDVMVIATLHASSTSPRKLLAKARKLLKSDGRLIVLN------------
XP_014073936.1(Bm_ATTC)      PQTP-DFELKSFDVIIVSASLADFNNPHKALGHIKSLLRPDGRLIVLH----EGTMDQDP
EMD85570.1(Bm_C5)            PQTP-DFELKSFDVIIVSASLADFNNPHKALGHIKSLLRPDGRLIVLH----EGTMDQDP
XP_007715650.1(BZ)           PQTP-DFELKSFDVIIVSASLADSNNPHKALDHIKSLLKPDGRLIVLH----EVTMDQDP
XP_014553689.1(Bv)           PQTP-DFELKSFDVIIVSASLADSNNPHKALDHIKSLLKPDGRLIVLH----EVTMDQDP
                                         :.                :   . **    * ...
```

Figure S42: Section from a multiple sequence alignment of maleidride PKSs, along with several characterised hrPKSs, using MUSCLE.[1-4] An asterisk (*) marks completely conserved amino acid residues, whereas a colon (:) identifies highly conserved residues and a period (.) indicates moderately conserved residues. The *CMet* motifs identified by Ebizuka and co-workers[3] are highlighted. In several maleidride PKSs, all motifs are conserved, whilst others are missing at least one motif.

```
                                                                   S
AAG52991.1_RifR                  EVL----------RP---FGDRPLALFGHSMGAIIGYELALRM-----PEAG--L--PAP
AAF43096.2_TEII                  GEM----------AP---LADRPHAFFGHSMGALLAYELARELRRRALP--G--P-----
Q0C8M2.1_LOVG_ASPTN              DPIRRTIVDWQTQHPHIPIV----GAIGFSEGALVTTLLLWQQQMGRLPWLPRMSVAMLI
A0A0A2JY30.1_CNSH(communesin)    ETL-------HTYGP---FD----GAFGFSQGAALIVSYLLERRAAY-PDES--L--PFR
C8VJR6.1_ALNB(asperlin)          EMI-------QAAGP---FD----GIIGFSQGGSVALSYLLQRQIDGHP-----P--PFR
Cc_RDW56968.1                    AKI-------EEDGP---FD----GILGFSHGGTLVAEFLRDWASRN-PWSP--P--PVR
TbR3                             EII-------EEEGP---FD----GVIGFSHGATLAYGFLAQHARRN-PFDP--PDTLFR
Ia_CAF9941818.1_b_mod             DII-------STEGP---FD----GILGFSHGATLAFAFLVQHARKH-PYEP--P--PVR
Ag_OJJ86354.1                    EVI-------EEDGP---FD----GVIGFSHGGALAHGLMVRHELQR-PHDP--P--LFR
EpiR11                           DVI-------ETDGP---FD----AVIGFSHGGTLACGFIAHWTKQH-PYEN--P--PFR
Bm48331_ENH99967.1               DFI-------EIEGP---FD----AVIGFSHGGTLACGFLAQWSTRH-PYEA--P--PFK
BmC5_EMD85581.1                  DFI-------EIEGP---FD----AVIGFSHGGTLACGFLAQWSTRH-PYEA--P--PFK
Bv_XP_014553701.1                DVI-------ETEGP---FD----AVIGFSHGGTLACGFLAQWSTRH-PCEA--P--PFK
Bz_EUC30036.1                    DVI-------ETEGP---FD----AVIGFSHGGTLACGFLAQWSTRH-PYEA--P--PFK
BfL1_ANF07287.1                  DVI-------EEEGP---FD----GVIGFSHGGTLAFGYLAHLAKTR-PNDA--L--PFR
CzmL1                            DVI-------EDDGP---FD----GILGFSHGGTLASGFIMHHMAMR-PSEP--L--PFR
CaL1                             EIV-------EEEGP---FD----GVLGFSHGGTLASGWLIHHAAKY-PLEP--L--PVR
Om_KIN05363.1                    EII-------ETEGP---FD----GVLGFSHGGTLASGWLMHHAATR-PLEP--S--PVR
TsRbtS_EED18832.1                EII-------EEDGP---FD----CILGFSHGGAVAAGLMVHHITQN-PYDS--P--LFR
TfL9                             EII-------EEEGP---FD----CVLGFSHGGAVAAGLMVHHAEQN-PYDS--P--LFR
ScyR1_QTE76001.1                 ETI-------EEDGP---FD----GIIGFSHGGALATGFMAHHAAKN-PYDS--P--LFQ
Dc_ZopR1_QTE75993.1              EVI-------EEEGP---FD----GILGFSHGGTLATGFLAHHATLN-PYDP--P--LFR
Zc_ZopM_BBU42027.1               EVI-------EEEGP---FD----GILGFSHGGTLATGFLAHHATLN-PYDP--P--LFR
PvL1_ASK38716.1                  DII-------AEEGP---FD----GILGFSHGGTLASGFLIHHSKTS-PYTP--P--PFR
Ta_OKL57045.1                    DVI-------EEDGP---FD----GVMGFSHGGTLASGFLIHHAKTR-PYDP--P--PFR
PhiM_BBG28510.1mod               DYL-------EENGP---YD----GILGFSQGSTLLAEFLCDFARRN-PGNE--P--PCR
Ts1_EED15412.1mod                EII-------DDEGP---FD----GLLGFSHGGSFLAELLARYARDN-PATD--VERLAR
                                   :           *         :*.* *.  .            *
```

```
AAG52991.1_RifR                      VHLFASGRRAPSR-YRDDDVRGASDERLVAELR---------------------------
AAF43096.2_TEII                      CHLFLSGRFAPTP-QGSDSDRLDTDEKVIAMIR---------------------------
Q0C8M2.1_LOVG_ASPTN                  CPWYQD---EASQ-YMRNEVM---------------------------------------
A0A0A2JY30.1_CNSH(communesin)        FLILCS---PVVP-LAGNAEYCHRILGCLSRDNESRIRSCQDTQISDLPERARIAMTMLT
C8VJR6.1_ALNB(asperlin)              WAVFFS---TVIA-FAPNDTFGSNILANLTDHEIRLLDGYPATDLSSLHPLTRALCETTA
Cc_RDW56968.1                        CAIFIC---SFAP-F---------------------------------------------
TbR3                                 CGVFIG---AMPP-F---------------------------------------------
Ia_CAF9941818.1_b_mod                CAVFVC---AMPP-F---------------------------------------------
Ag_OJJ86354.1                        CAIFFN---SMPPLF---------------------------------------------
EpiR11                               CAVFFN---SLPP-F---------------------------------------------
Bm48331_ENH99967.1                   CAIFFN---SLPP-F---------------------------------------------
BmC5_EMD85581.1                      CAIFFN---SLPP-F---------------------------------------------
Bv_XP_014553701.1                    CAIFFN---SLPP-F---------------------------------------------
Bz_EUC30036.1                        CAIFFN---SLPP-F---------------------------------------------
BfL1_ANF07287.1                      CAVFMN---APPP-F---------------------------------------------
CzmL1                                CALFFN---ALPP-F---------------------------------------------
CaL1                                 CVVFLN---SLPP-F---------------------------------------------
Om_KIN05363.1                        CAVFLN---SLPP-F---------------------------------------------
TsRbtS_EED18832.1                    CAIFFN---SFPP-F---------------------------------------------
TfL9                                 CAIFFS---SFPP-F---------------------------------------------
ScyR1_QTE76001.1                     CAIFFN---SMPP-F---------------------------------------------
Dc_ZopR1_QTE75993.1                  CAVFFS---SLAP-F---------------------------------------------
Zc_ZopM_BBU42027.1                   CAVFFS---SLAP-F---------------------------------------------
PvL1_ASK38716.1                      CAVFFN---SLPP-F---------------------------------------------
Ta_OKL57045.1                        CAIFLN---SLPP-F---------------------------------------------
PhiM_BBG28510.1mod                   CAIFMN---GIPP-Y---------------------------------------------
Ts1_EED15412.1mod                    CAVFIN---SFPP-F---------------------------------------------

AAG52991.1_RifR                      -----------------------KLGGSDA-------AMLADPELLAMVLPAIRSDYRAV
AAF43096.2_TEII                      -----------------------RLGGTVG-------KVFDDPDVMEMVMPPLRADYRAV
Q0C8M2.1_LOVG_ASPTN                  -----------------------ENHDDDH-------DSKDTEWQEELVIRI--------
A0A0A2JY30.1_CNSH(communesin)        DILDASTTITQEPRRFYLD----RELPDVP-------CALHP-DLCLTRLPV--------
C8VJR6.1_ALNB(asperlin)              QTFYSAKTGGFISPNTPIAEFSKRDDPSQP-------RVFHP-ALLGDRIPI--------
Cc_RDW56968.1                        -----------------------RMDKNGE-------PVFQNMDSVTGGIGL--------
TbR3                                 -----------------------RINDTKS-------FVYDQ-DL-DNMVRI--------
Ia_CAF9941818.1_b_mod                -----------------------RLGGNDE-------WIYDQAFLEPGALAI--------
Ag_OJJ86354.1                        -----------------------RTDKDDANGVAGPCIIYDKLALQNSVCGV--------
EpiR11                               -----------------------VVGRDGE-------FAFEK-GL-KGCVNM--------
Bm48331_ENH99967.1                   -----------------------VVSGSGN-------FIFEK-HL-KGKITI--------
BmC5_EMD85581.1                      -----------------------VVSGSGN-------FIFEK-HL-KGKITI--------
Bv_XP_014553701.1                    -----------------------VVSGNGD-------FIFEK-HL-KGRLTI--------
Bz_EUC30036.1                        -----------------------VVSGNGD-------FIFEK-HL-KGRLTI--------
BfL1_ANF07287.1                      -----------------------RMDTYGN-------LIVEE-GL-QDLVRI--------
CzmL1                                -----------------------RKHPGQD-------PVVDE-GLEDGCIAI--------
CaL1                                 -----------------------RMKAGED-------PVVDE-GFKDGCIQI--------
Om_KIN05363.1                        -----------------------RMNPGED-------PVVDE-GLRDGCIQI--------
TsRbtS_EED18832.1                    -----------------------RMNDENE-------PILNE-GL-EGKLTV--------
TfL9                                 -----------------------RMNDNNE-------PILNE-GL-EGKITV--------
ScyR1_QTE76001.1                     -----------------------RMDDEQN-------PIFEK-GL-EGKIRI--------
Dc_ZopR1_QTE75993.1                  -----------------------RMNDDET-------PVFEE-GL-EGKIKI--------
Zc_ZopM_BBU42027.1                   -----------------------RMNDDET-------PVFEE-GL-EGKIKI--------
PvL1_ASK38716.1                      -----------------------RMDPGEE-------LVVDD-DL-ARHLTI--------
Ta_OKL57045.1                        -----------------------RMNPGES-------PVVDA-DL-EGYLTL--------
PhiM_BBG28510.1mod                   -----------------------RMGDDEK-------PIIDY-GLLEHFPSI--------
Ts1_EED15412.1mod                    -----------------------RNDPDQN-------PIIDY-ELLKHFPKI--------
```

```
                                                         D                               H
AAG52991.1_RifR                    ETYRHEPGRRVDCPVTVFTGDHD-PRVSVGEARAWE---EHTTGPADLRVLPGGHFFLVD
AAF43096.2_TEII                    GAYTWQPGPPLAVPVTVLVGDQD-PVVPVAAAAAWR---EHTTAGSDLRVLPGGHFYLDQ
Q0C8M2.1_LOVG_ASPTN                -------------PTLHLQGRDD--FALAGSKMLVAR--HFSPREAQVLEFAGQHQFPNR
A0A0A2JY30.1_CNSH(communesin)      -------------ATLHVRGTTDAKALWNCGFLIQS---FFDSPKLRVFEHKSGHDIPRS
C8VJR6.1_ALNB(asperlin)            -------------PTVHITGRKDNSLMVGLSVLVQG---LCDQRLIRSLTHSGGHNVPRS
Cc_RDW56968.1                      -------------PTLHVVGKQD--FAYKYGMGLYR---LFEGQGGILVTHEKGHELPND
TbR3                               -------------PTVHIAGKMD--FVYQHSLKLFA---LCSKSWTKLWTHDKGHEIPKD
Ia_CAF9941818.1_b_mod              -------------PSVHVVGKSD--FVLEHSLKLYA---VCDAARARLVVHGKGHEIPGD
Ag_OJJ86354.1                      -------------LSLHVVGQKD--FAYEHSMALYK---DWEPNSAMLIVHERGHVIPSD
EpiR11                             -------------PTLHVVGRED--FIYEHSLKLYR---VCEEKSATLILHEKGHEIPSE
Bm48331_ENH99967.1                 -------------PTLHVVGRKD--FIYEHSLKLHQ---ICGDKKATLMLHDNGHEIPRE
BmC5_EMD85581.1                    -------------PTLHVVGRKD--FIYEHSLKLHQ---ICGDKKATLMLHDNGHEIPRE
Bv_XP_014553701.1                  -------------PTLHVVGRRD--FIYEHSLKLHQ---ICGDKNATLMLHDRGHEIPQE
Bz_EUC30036.1                      -------------PTLHVVGRRD--FIYEHSLKLHQ---ICGDKNATLMLHDRGHEIPQE
BfL1_ANF07287.1                    -------------PTLHVVGKKD--FIYNFSLKLHE---LCDSANSTLVLHEKGHEIPSD
CzmL1                              -------------PSVHVVGEND--FVYEYSIRLYH---LCDTRAAQLVTHKRGHDIPRD
CaL1                               -------------PSVSVAGTKD--FVYDYSIKLHK---LCDPRKSQLVIHDKGHDIPSD
Om_KIN05363.1                      -------------PTVSVVGTKD--FAYNYSMKLHQ---LCDPRKSQLVVHNKGHDIPND
TsRbtS_EED18832.1                  -------------PTLHVAGRQD--FVYNYSLNLYK---ICNAETSTLLTHDGGHEIPTD
TfL9                               -------------PTLHVMGSKD--FVYKYSLDLYN---LCKSESSAILTHDKGHEIPTD
ScyR1_QTE76001.1                   -------------PTLHVMGTKD--FAYKYSVNLYN---LCDSKSSSIVLHDKGHEIPRD
Dc_ZopR1_QTE75993.1                -------------PTLHVMGTKD--FIYSYSLHLFN---LCDVKSSAILVHDRGHEIPSD
Zc_ZopM_BBU42027.1                 -------------PTLHVMGTKD--FIYSYSLHLFN---LCDVKSSAILVHDRGHEIPSD
PvL1_ASK38716.1                    -------------PTLSIAGTRD--FVYKQSLMLHQ---LCDEKSSQLILHGKGHEIPGD
Ta_OKL57045.1                      -------------PTLSIAGKKD--FVFESSVALYN---LCDPRKSQLVVHENGHDVPRD
PhiM_BBG28510.1mod                 -------------PTLHVVGKKD--FVYEYSTILQA---STASAWATLIAHEKGHEISND
Ts1_EED15412.1mod                  -------------PTLHVVGTSD--FVHEYSTILYEKLHQKAPTSTGLVTHSKGHEIPRD
                                                    . *  *                               *
```

Figure S43: Section from a multiple sequence alignment of putative maleidride hydrolases, with known hydrolytic enzymes using MUSCLE.[1, 4] An asterisk (*) marks completely conserved amino acid residues, whereas a colon (:) identifies highly conserved residues and a period (.) indicates moderately conserved residues. Conserved residues from the active site are highlighted in bold red. The conserved Sm-X-Nu-X-Sm-Sm (Sm = small residue, Nu = nucleophile) motif is also shown highlighted in red.[14]

```
BBU42028.1_ACS_ZC           --MAHPTNGTLFVRDSRTKQEYEIPIVDNAVLAVDFKNIRGYSDGSKM-----RGLLLYD
QTE75995.1_ACS_DC           --MAHPTNGTLFVRDSRTKQEYEIPIVDNAVLAVDFKNIRGYSDGSKM-----RGLLLYD
BBG28507.1_ACS_phiJ         MALSSQAQGKLFVRDSRTSREYEIPISNNTINAADFQKINLPTKGKSLTKAL--GLQLYD
EED15409.1_ACS_TS1          -----MSDGTLFIQDSRTSKQYTISVTSDTITAVDFQKITSPT----------GKLALYD
OJJ86352.1_ACS_AG           -----MSDGTLSIKDSRTGRDYEIPIRDNAILATSLKQIKGPAETANPADKVAGGLRCYD
ANF07286.1_ACS_BF           -----MSSGTLFVKDSRTSLNYEIPIHRNAIAATAFKKIKAPVSGSDPADKVDGGLRVHD
epiR8_ACS_WA                -----MSRGSLFVRDSRTSLDYEIPIERNYIPATAFKKIKAPNANANRADKVGGGLRVHD
ENH99968.1_ACS_BM48331      -----MSNGFLFVRDSRTTQEYRVPIQRNAILATAFKDIKAPSSSGNRADKLDSGLRVHD
EMD85580.1_ACS_BMC5         -----MSNGFLFVRDSRTTQEYRVPIQRNAILATAFKDIKAPSSSGNRADKLDSGLRVHD
EUC30035.1_ACS_BZ           -----MSNGFLLVKDSRTTLEYRVPIQRNSVLATAFKDIKAPSSSGNRADKVGSGLRVHD
XP_014553700.1_ACS_BV       -----MSNGFLLVKDSRTTLEYRVPIQRNSVLATAFKDIKAPSSSGNRADKVGSGLRVHD
QTE76003.1_ACS_SA           -----MPEGKLLVKDTRTSLEYEIPITRNAVSATDFNKIKGRATAANRADKISSGLRIYD
ASK38711.1_ACS_PV           -----MSHGALHIRDSRTSREYEIPIRRNTVLATDLKQIKASPVGADRADKVGDGLRVFD
OKL57051.1_ACS_TA           -----MSDGSLYIKDSRTSREYEIPIHRNTVLATAFKKIKASLNGANKADKVADGLRLYD
vibL3_ACS                   -----MSNGTLFIVDSRSGKNYEIPIRRNTVLATDLKKIKASDVGANRADKVADGLRLYD
EED18839.1_ACS_TS2          -----MSDGTLFVEDSRSGKKYEIPIRHNTVLATDLKRIKASSTAANRADKVADGLRLYD
ACS_TF                      -----MSDGTLFIQDSRSGKKYEIPIRHNTVLATDLKKIKASSVGANRADKVADGLRLYD
                                  . * * : *:*:  .* :.:  : : *. :: *                *  .*

BBU42028.1_ACS_ZC           PGLQNTAIKRSQISSSDA-RGLPMIRGYSVEQLYSLQSDFEDLFHLMVLGKYPTPEEKEI
QTE75995.1_ACS_DC           PGLQNTAIKRSQISSSDA-RGLPMIRGYSVEQLYSLQSDFEDLFHLMVLGKYPTPEEKEI
BBG28507.1_ACS_phiJ         PGMQNTAIKKTEIIGRDPSTGLPLLRGVTSQELWKRRCDFEELFSLMVFGNYPTIVEREA
EED15409.1_ACS_TS1          PGLQNTIIKKTQITGRDPVTGITLFRGLSAKEIWNRHADFEDHFHLLVFGKYPSPEESEA
OJJ86352.1_ACS_AG           PGLKNTAPIKSSLTWIDGEKGVLLFQGYSIEQLW--DCDFEDILHLMLRGDLPSLSQREG
ANF07286.1_ACS_BF           PGLQNTTVVETDISFSNSDSGLLLFRGYSLDQLW--DSDFEELFHLLVWGKYPTRVQKDD
epiR8_ACS_WA                PGLQNTTVVETGVSFAD--RGLLLFRGYSLEQLW--GSDFEDMLHLMVWAKYPKPLQRET
ENH99968.1_ACS_BM48331      PGLLNTTVVETGVSFADGERDLLLFRGYSLEQLW--QSDYEDMLHLLVWAKYPTPVQKES
EMD85580.1_ACS_BMC5         PGLLNTTVVETGVSFADGERDLLLFRGYSLEQLW--QSDYEDMLHLLVWAKYPTPVQKES
EUC30035.1_ACS_BZ           PGLLNTTVVETGVSFADGERDLLLFRGYSLEQLW--QSDYEDMLHLMVWAKYPTPVQKES
XP_014553700.1_ACS_BV       PGLLNTTVVETGVTYRDGERDLLLFRGYSLEQLW--QSDYEDMLHLMVWAKYPTPVQKES
QTE76003.1_ACS_SA           PGLQNTAVVETSTTFADSDNGLLLYRGYSLKQVW--ESDFEEILHLMVWGKYPTSSQKES
ASK38711.1_ACS_PV           PGLKNTCVVETNMTYTDGHRGLLLFRGYALEQLW--QAEFEDMLHLLVWGKYPTPSQREA
OKL57051.1_ACS_TA           PGLQNTTVVETGMTYTDSERGLLLFRGYALEQLW--DAEFEDMLHLMVWGKLPTLSQRES
vibL3_ACS                   PGLENTAVVETSMTFAD--RGLLILRGYSLEQLW--QSDFEDILHLLVWGKYPTPSQSES
EED18839.1_ACS_TS2          PGLENTTVVETSMTYADADRGLLMFRGYALEQLW--ESEFEDMLHLMVWGKYPTPSQSAS
ACS_TF                      PGLENTTVIETSMTYADSDRGLLMFRGYALEQLW--ESDFEDMLHLMVWGKYPTPSQSES
                            **: **    :     :   .: : .* : .:::   .::*: : *:: .. *.  :
```

```
BBU42028.1_ACS_ZC           LRHTFAQEMINVPDTVVAAIKAFP---PSSTGMSMMLAGLSAYLASYDHAIPAKQGGNLY
QTE75995.1_ACS_DC           LRHTFAQEMINVPDTVVAAIKAFP---PSSTGMSMMLAGLSAYLASYDHAIPAKQGGNLY
BBG28507.1_ACS_phiJ         LRYQFAEYMKEVPKVVGSVIRKFP---PSSPPLPMITAGLSAYLSSDPDFIPAIHGGNIY
EED15409.1_ACS_TS1          LRRRLAVQMTVVPETVIKAVQAFP---RTSHPLPMIIAGLAAFISADPSSLPAIRGGNIY
OJJ86352.1_ACS_AG           FRQRIANAMRDIPQSVVDVVRSFP---RSSPMMPVVMAGLSCYVGNQPDSIPANRGGNIY
ANF07286.1_ACS_BF           LSNTLANYMKNVPENVVKAIRSFP---RSTSPMPMVIAGLAAYIASDPESIPAANGGNIY
epiR8_ACS_WA                LRQALVTAMLDIPESVIAAVHALPAKRRTSPPMPMVFAGLAAYLGSNPHMIPASNGGNIY
ENH99968.1_ACS_BM48331      LRRALIAAMLEVPKTVFEVVSAFP---SASPPMPMVVAGLAAYLGSNPDMIPASNGGNIY
EMD85580.1_ACS_BMC5         LRRALIAAMLEVPKTVFEVVSAFP---SASPPMPMVVAGLAAYLGSNPDMIPASNGGNIY
EUC30035.1_ACS_BZ           LRRLLIAAMLEVPKNVFEIVSAFP---SSSPPMPMVVAGLAAYLGSNPAMIPASSGGNIY
XP_014553700.1_ACS_BV       LRRLLIAAMLEVPKNVFEIVSAFP---SSSPPMPMIVAGLAAYLGSNPAMIPASSGGNIY
QTE76003.1_ACS_SA           LRRALALTMSKVPDSVVKAVQSFS---TSSPPMPMILAGLAAYIGSDPESIPAFKGGNIY
ASK38711.1_ACS_PV           LRLTLATLMSTTPKHVAEVIETFP---PTSPPMPMIVAGLSAYLAHHADSIPACAGGNIY
OKL57051.1_ACS_TA           LRKTLATLMANIPEHVVEVITTFP---KAGPIMPMIVAGLSAYLADDVGSIPAYNGGNIY
vibL3_ACS                   LRKTLASLMSDVPQTVFEVIENFP---RECPPMPMIVAGLAAYLANDLDSIPAFNGGNIF
EED18839.1_ACS_TS2          LRKDLASLMGDIPKTVFEVIEKFP---RDCPPMPMLVAGLAAYLSDDLDSIPTFNGGNIF
ACS_TF                      LRKDLASLMSNIPSTVFEVIEDFP---RDCPPMPMLVAGLAAYLSNDLDSIPAFNGGNIY
                            :   :   *   *. *   :  :.        :.:: ***:.::.     :*:  ***::

BBU42028.1_ACS_ZC           HNNPRLANKAAIKTTAAYAVVLGLTISHRKGIPFAPAQANKSFYENLFNMMGMADPITKL
QTE75995.1_ACS_DC           HNNPRLANKAAIKTTAAYAVVLGLTISHRKGIPFAPAQANKSFYENLFNMMGMADPITKL
BBG28507.1_ACS_phiJ         HRNPRAADEAAIKTAAVYAVVIGLISCHKKGINFVPADTDDTFLENLFRMCGLVDPATRR
EED15409.1_ACS_TS1          HGNRALCDEGVIRATAAYAVVMGLINSHRKQLPYVPADPQKSFYENVFAMM-------RC
OJJ86352.1_ACS_AG           HGNGNQVDAGIIRTVAAFAVVVGLAASHLKGTSFCLPSRDRTFYENLFVMMGHVDPSTGE
ANF07286.1_ACS_BF           QGNTAQTDLGILKTVSAYAVVLGLVASHRKDIPFVPASSENSYYENLFIMMGMVDRVSGR
epiR8_ACS_WA                QGNIKKTDEAIIKTVAAYAVVIGLVSSHRRGIKFTPASMEYTYIENLFRMMGMVDSATGQ
ENH99968.1_ACS_BM48331      QGDIEKTDKAIVKTIAAYAVVVGMAACHRKGIEFTAPLLDYNFIENLFHMSGMVDDLTGR
EMD85580.1_ACS_BMC5         QGDIEKTDKAIVKTIAAYAVVVGMAACHRKGIEFTAPLLDYNFIENLFHMSGMVDDLTGR
EUC30035.1_ACS_BZ           QGNIEKTDEAIINTIAAYAVIVGMAACHRKGIEFTAPSLDYSFVENLFHMSGMVDDLTGR
XP_014553700.1_ACS_BV       QGNIEKTDKAIINTIAAYAVIVGMAACHRKGIEFTAPSLDYSFVENLFHMSGMVDDLTGR
QTE76003.1_ACS_SA           HGNLDRVDKAIVKTVAAYGVCIGLAASHRKGIKFVPASPDKTYFENIFTMMGHVDPATGH
ASK38711.1_ACS_PV           HGNPEQTDRAILRTAAAYAVVMGMAASHRRGIAFTPATTDNTYFENLFIMMGLVEPYTGR
OKL57051.1_ACS_TA           HGNIEKVDEAILKTVAAFATVVGLAASHRRGIKFKPATTDNTYHENLFIMMGLVEPFTER
vibL3_ACS                   QGNVEKTDEAILKTVAAYASVMGIVGSHRRGIKFTQPSEDKTYFENMLTMMGIVDPATGT
EED18839.1_ACS_TS2          HGNVEKTDEAILKTVAAFASVVGIAGSHRRGIAFTPPSLDKGYLDNLFKMMGIVDPTTNK
ACS_TF                      HGNVEKTDEAILKTVAAFASVVGIASSHRREIKFTPPSLDKGYLDNLFTMMGVVEPTTGS
                            : :    : . :.: :.:.  :*:  .* .   :  .  :  : :*:: *

BBU42028.1_ACS_ZC           PNPL-YIDVFRRAAILNADNGMTHSTFLTLVAGSALPDPLSCLISATSAAYGPLHYGAQE
QTE75995.1_ACS_DC           PNPL-YIDVFRRAAILNADNGMTHSTFLTLVAGSALPDPLSCLISATSAAYGPLHYGAQE
BBG28507.1_ACS_phiJ         PDPH-VLAVFRKGLVLNCDNGMTQSNLVLCATASSLCDPISCLISAISAAYGPLHYGAQE
EED15409.1_ACS_TS1          PVHHNYLVTFREGMVLNSDNGMTQSSVVLLSTASSLPDPISCLISAITAAYGPLHYGAQE
OJJ86352.1_ACS_AG           PNHT-QLSCFRHFGALSVDHGLSNSTLTLLVTASTLPDPTSALISALASAYGPLHFGASE
ANF07286.1_ACS_BF           PDTL-QLSCFRRFAALSCDNGMALSVFATLVCASSLADPISCLISALAAAYGPLHFGATE
epiR8_ACS_WA                PDPK-KLSCFRRFAALNMDHGMALAVFSTLVTASSLTDPISCLIASLAAAYGPLHFGATE
ENH99968.1_ACS_BM48331      PDAM-KVSCFRRFAALNMDHGMALAVFSTMVTASSLTDPISCLVASLAAAYGPLHFGATE
EMD85580.1_ACS_BMC5         PDAM-KVSCFRRFAALNMDHGMALAVFSTMVTASSLTDPISCLVASLAAAYGPLHFGATE
EUC30035.1_ACS_BZ           PDAM-KVSCFRRFAALNMDHGMALAVFSTMVTASSLTDPVSCLIASLAAAYGPLHFGATE
XP_014553700.1_ACS_BV       PDAM-KVSCFRRFAALNMDHGMALAVFSTMVTASSLTDPVSCLIASLAAAYGPLHFGATE
QTE76003.1_ACS_SA           PDPL-ILSCFRRFAVLNADHGMTQSTFVLLATASSLPDPISCLISALAAAYGPLHFGAPE
ASK38711.1_ACS_PV           PDPV-KLSCFRRFAALNSEHGMALSSFAMVVTASSLTDPISGLISSLVAAYGPLHFGAPE
OKL57051.1_ACS_TA           PDPL-KLSCFRRFAVLNADHGMALSSFSLLVTASSLTDPISGLIGSLAAAYGPLHFGAPE
vibL3_ACS                   PDTL-KLSCYRRFAILNSDHGMALSVFANLVTTSSLTDPISGVISSLAAAYGPLHFGAPE
EED18839.1_ACS_TS2          PSPE-KLDCFRRFTIINTDHGMALSAFAHLVATSALADPISGLIGSLVAAYGPLHFGAPE
ACS_TF                      PSPD-KLDCFRRFTIINTDHGMALSAFSHLVATSALADPISGLIGSLVAAYGPLHFGAPE
                            *     :  :*    :. ::*:: : .      *:* ** * ::.:  :******:** *

BBU42028.1_ACS_ZC           ATWNNLIEIETKENVPAFLEQVKRRERRLFGYGHRVHKTEDPRLAIIKGLL-KELNVSPA
QTE75995.1_ACS_DC           ATWNNLIEIETKENVPAFLEQVKRRERRLFGYGHRVHKTEDPRLAIIKGLL-KELNVSPA
BBG28507.1_ACS_phiJ         AGYRTLSEIGSADRVPHFLEQVKRRERRLFGYGHRTFATEDPRLNAVKGWL-QELDFDSK
EED15409.1_ACS_TS1          AGSTTLKSIGSLDKVPEFLEQVKRRERRLFGFGHRLHKREDPRLASVKRWL-KMMDYTPD
OJJ86352.1_ACS_AG           SAFRVMTELGSPDRVPALIKEVKQGKRRLFGYGHRSYKTVDPRIRPIQKLL-QMLDAQSN
ANF07286.1_ACS_BF           AAHRALQEIGSVERVPDFLEQVKRGERKLFGYGHRTYKGTDPRVIPIKKLL-EDSNATSN
epiR8_ACS_WA                AAHLTLQNIGGKANVPAFIEEIKQGKRKLFGYGHRTYKGIDPRVRPIKELV-EESGANED
ENH99968.1_ACS_BM48331      AAHLNLRSIGDKSKVPEFISEVKQGKRKLFGYGHRTYKGTDPRVRPIKELI-EDSGANSD
EMD85580.1_ACS_BMC5         AAHLNLRSIGDKSKVPEFISEVKQGKRKLFGYGHRTYKGTDPRVRPIKELI-EDSGANSD
EUC30035.1_ACS_BZ           AAHLSLRSIGDKSKVPEFISEVKQGKRKLFGYGHRTYKGTDPRVRPIKELI-EDSGANSD
XP_014553700.1_ACS_BV       AAHLSLRSIGDKSKVPEFISEVKQGKRKLFGYGHRTYKGTDPRVRPIKELI-EDSGANSD
QTE76003.1_ACS_SA           SAYRVIKKVGKPENVPALIDEVKRGERRLFGYGHRTYKTIDPRLAPIKELL-VELNAASN
ASK38711.1_ACS_PV           AEYKTIQAVGSVDQVPAFLEEVKSGKRRLFGYGHRTYTTIDPRVKPIKDMLFSELDGDSD
OKL57051.1_ACS_TA           AAYKTLQKVGSVENVPAFLDNVRTGKQRLFGYGHRTYKTVDPRVKPIKDML-VELDADSD
vibL3_ACS                   SAYKTISKIGGPENIPAFLDEVKSGKRRLFGYGHRTYKTVDPRLAPIKEML-QELNMESD
EED18839.1_ACS_TS2          AAYKTIKSIGGPQNVPSFLDEVKSGKRRLFGYGHRTYRTVDPRLAPIKSAL-QTLNVETD
ACS_TF                      AAYKTIRNIGGPENVPAFLEEVKSGKKRLFGYGHRTYKTVDPRLAPIKSAL-QTLDVPDD
                            :    :  :    .:* ::.::.  :..***:*** .   ***:  ::  :    .
```

```
BBU42028.1_ACS_ZC          TIPLMEIAYEIDRLAALDQYFTSRGLSANADFYFGFLIHAFGFDPDMITLANLAMRILGL
QTE75995.1_ACS_DC          TIPLMEIAYEIDRLAALDQYFTSRGLSANADFYFGFLIHAFGFDPDMITLANLAMRILGL
BBG28507.1_ACS_phiJ        REPLMKIAEEIDRLAAQDDYFTSRGLRANADFYTLFVFRAYGFDWDMIGAANFCMRIIGF
EED15409.1_ACS_TS1         QEPLLELAQEIDRLASSDEYFIKRNLRANADFYTHFLFKAWGFDWDMLCAANMFHRIIGL
OJJ86352.1_ACS_AG          --PLFAVAQEIDRVASQDEYFVSRGLKANADLYGQFFYTAMGWPPSFIPLMMIIHRLPGL
ANF07286.1_ACS_BF          --PLIEIAKSIEIHASTDDYFKSRGLSANADFYGNFVFSAIGFDPDFIPVAMLAQRIIGI
epiR8_ACS_WA               --PLIEVAKEIERLASTDEFFKSRGLHPNADFYGNFVFTAIGFESAFIPIAMLSQRLIGI
ENH99968.1_ACS_BM48331     --PLIEIAREVERLASNDDYFTSRGLHPNADFYGNFVFTAVGFQSDFIPIAMISQRLIGI
EMD85580.1_ACS_BMC5        --PLIEIAREVERLASNDDYFTSRGLHPNADFYGNFVFTAVGFQSDFIPIAMISQRLIGI
EUC30035.1_ACS_BZ          --RLIEIAREIERLASNDDYFTSRGLHPNADFYGNFVFTAVGFQSDFIPIAMISQRLIGI
XP_014553700.1_ACS_BV      --PLIEIAREIERLASNDDYFTSRGLHPNADFYGNFVFTAVGFHSDFIPIAMISQRLIGI
QTE76003.1_ACS_SA          --PLLKVAQEIDRIAANDDYFRKRGLNANADFYGVFFFIACGFEAEFVPILMLAQRIAGI
ASK38711.1_ACS_PV          --PLVKVAREIDRLSSTDEYFIKRKLHANADFYGTFFFNRLGFQPEEIPVAMVAQRLVGI
OKL57051.1_ACS_TA          --PLLQTAREIDRLSSTDEYFLRRGLHANADFYGTFFFVALGFTPEEIPIAMLAQRIVGI
vibL3_ACS                  --PLLKIAREIDLISATDEYFVKRGLHANADFYGVFFFIGIGFQPEEIPVAMFAQRLVGV
EED18839.1_ACS_TS2         --IPLETAYEIDRLASNDDYFLKRGLHANADFYTPYCFIKIGFHPEEFPIAMFAQRIIGI
ACS_TF                     --VPLKTAYEIDRLAANDEYFLKRGLHANADFYTPYCFIMIGFEPEEFPIAMFAQRIIGI
                               .  * .::  :: *::*  * * .***:*  :     *:    .    .  *: *.

BBU42028.1_ACS_ZC          MAHWREAMDQ-EIKLFRP-----------------------LHIYTGPKQRITA------
QTE75995.1_ACS_DC          MAHWREAMDQ-EIKLFRP-----------------------LHIYTGPKQRITA------
BBG28507.1_ACS_phiJ        MAHWREAMEQ-EIKIFRA-----------------------RDYYVGPSKKDPNRESSGT
EED15409.1_ACS_TS1         MAHWREAMDQ-PIKIFRA-----------------------TDLYVGPVVIQEDNRTVLE
OJJ86352.1_ACS_AG          MGHWRNAMLS-EIMVYRP-----------------------PHLYVGPVEKRAP------
ANF07286.1_ACS_BF          MAHWREYMLK-RGKLFRP-----------------------SHIYTGNTEPLCN------
epiR8_ACS_WA               MAHWREAMGK----------------------------------VSGSRSEFF-------
ENH99968.1_ACS_BM48331     MAHWREAM--------------------------------------GESSDT--------
EMD85580.1_ACS_BMC5        MAHWREAM--------------------------------------GESSDT--------
EUC30035.1_ACS_BZ          MAHWREAMVR-GIKLFRP-----------------------SHIYTGDTEPVYT------
XP_014553700.1_ACS_BV      MAHWREAMVR-GIKLFRP-----------------------SHIYTGDTEPVYT------
QTE76003.1_ACS_SA          MAHWRESMTR-DIKLFRP-----------------------SHIYTGPTTPVAD------
ASK38711.1_ACS_PV          LAHYRESMLKNKIRLFRP-----------------------THVYTGETEPVLE------
OKL57051.1_ACS_TA          MAHYRESMRK----LFRP-----------------------THVYIGETDPVQD----KV
vibL3_ACS                  MAHWRESMRM-YFQLSVPKKADLAHNQSTKYQVVPPYTRLHRRYRAGSRYENGRNEGTIK
EED18839.1_ACS_TS2         MAHWREAMLR-KVKLFRP-----------------------THIYTGETEPVEH------
ACS_TF                     MAHWREAMLR-KVKLFRP-----------------------THVYTGETEPVEH------
                           :.*:*: *                                      *

BBU42028.1_ACS_ZC          -----ARL
QTE75995.1_ACS_DC          -----ARL
BBG28507.1_ACS_phiJ        GILAQARL
EED15409.1_ACS_TS1         EPKIQSRL
OJJ86352.1_ACS_AG          ---LSAKL
ANF07286.1_ACS_BF          ---FSPKL
epiR8_ACS_WA               --------
ENH99968.1_ACS_BM48331     --------
EMD85580.1_ACS_BMC5        --------
EUC30035.1_ACS_BZ          ---ASAKL
XP_014553700.1_ACS_BV      ---ASAKL
QTE76003.1_ACS_SA          ---LLSKL
ASK38711.1_ACS_PV          MTAPSAKL
OKL57051.1_ACS_TA          IIPREKKL
vibL3_ACS                  TLNYKAGN
EED18839.1_ACS_TS2         -IKIPSKL
ACS_TF                     -TRVSSKL
```

Figure S44: Multiple sequence alignment of maleidride alkylcitrate synthases (ACSs) using MUSCLE[1]. An asterisk (*) marks completely conserved amino acid residues, whereas a colon (:) identifies highly conserved residues and a period (.) indicates moderately conserved residues. Catalytic residues are conserved (depicted in **red**). Acyl CoA binding residues (depicted in **purple**) and oxaloacetate binding residues (depicted in **blue**) are also largely conserved, with the exception of the residue highlighted in yellow.

```
                                              Motif I
Q9I4V0.1_NQRED        MGVFRTRFTETFGVEHPIMQGGMQWVGRAEMAAAVANAGGLATLSALTQPSPEALAAEIA
Ta_OKL57052.1         -MPFNTRLTRALGIQIPIVQGGMQWVGYAELASAVSNAGGLGIVTALTQPSPEDLRKEIR
Czm_KAF2215726.1      -MPFDTQLTRALGIRLPIVQGGMQWVGYAELAAAVSNAGGLGMLTALTQPTPEDLRNEIR
                         * *.:* ::*:   **:******** **:*:**:*****. ::*****:** *   **
                               Motif II
Q9I4V0.1_NQRED        RCRELTDRPFGVNLTLLPTQKPVPYAEYRAAIIEAGIRVVETAGNDPGEHIAEFRRHGVK
Ta_OKL57052.1         RCKNMTSRPFGVNITILPAMVPPDYRAYAQVVIEEGVRVVETAGNNPEQIITQLKDAGVI
Czm_KAF2215726.1      RCRSMTPNPFGVNITMLPALQPPPYVEYAHVAIQEGVKIIETAGNSPAALISLLKAGGCI
                      **..:* .*****:*:**:   *  *  *  . *: *:.::*****.*   *: :.  *
                       Motif III                Motif IV
Q9I4V0.1_NQRED        VIHKCTAVRHALKAERLGVDAVSIDGFECAGHPGEDDIPGLVLLPAAANRLRVPIIASGG
Ta_OKL57052.1         ILHKCTSIRHAQRAARLGADFLSIDGFECAGHVGETDITNLILLSRARQSLTTPFIASGG
Czm_KAF2215726.1      VIHKCTTIRHAQSAAAMGVDFVSMDGFECAGHIGETDIANTVLLSRARQSLKIPFIASGG
                      ::****::***   *   :*.* :*:******** ** **.. :**. * : *   *:*****
                         Motif V                                         Motif VI
Q9I4V0.1_NQRED        FADGRGLVAALALGADAINMGTRFLATRECPIHPAVKAAIRAADERSTDLIMRSLRNTAR
Ta_OKL57052.1         FADGHGLAAALVLGAEGINMGTRFLCTVESPIHANIKEEIVKAQETDTVLVLRRWRNTMR
Czm_KAF2215726.1      FADGRGLAAALILGAQGINMGTRFLCTAEAPIHIEIKKRIVRAKETDTRLVLRRWKNTTR
                      ****.**.*** ***:.********.* *.***   :*   *   *.* .* *::*   .** *

Q9I4V0.1_NQRED        VARNAISQEVLAIEA--RGGAGYADIAALVSGQRGRQVYQQGDTDLGIWSAGMVQGLIDD
Ta_OKL57052.1         LFKNKVTREALEIEKSTSSSDDFGEIAPYVSGKRGREVFRLGDPDYGVWTAGQVLGLIRD
Czm_KAF2215726.1      LYDNKVARKASEVERS-SASGNFEEIAPLVSGKRGREVFRLGDPDFGVWTAGQVIGLIDD
                      :  * ::.:.  :*      .. .: :**. ***:***:*:. **.* *:*:** * *** *

Q9I4V0.1_NQRED        EPACAELLRDIVEQARQLVRQR--LEGMLAGV
Ta_OKL57052.1         IPTCSGLCTRIEKEAEITLAQTTALIVRQAHL
Czm_KAF2215726.1      IPTVSELMARIEREVEAALRVSQALVGLEAKL
                       *: : *    *  :.    :      *    * :
```

Figure S45: Multiple sequence alignment using MUSCLE[1] of the putative quinone reductases KAF2215726.1 and OKL57052.1, aligned with the known quinone reductase PA1024 (Q9I4V0.1) from *Pseudomonas aeruginosa*. An asterisk (*) marks completely conserved amino acid residues, whereas a colon (:) identifies highly conserved residues and a period (.) indicates moderately conserved residues. Motifs which define this class of enzyme are depicted in red.[15] Where the maleidride sequences differ from the consensus sequence residues are highlighted in yellow. However, in all cases the substitution is for another hydrophobic residue.

```
A0A3G1DJG9.1-Mfr1                    -----MATAILPSTSGVIGL-WDGTTDGKEGFMDYAN-----------------------      31
A0A3G1DJF4.1-Mfr2                    ---MATATTTLHSTTGTVYV-ADGTTDGKVGYYNHTD-----------------------      33
OKL57227.1-T.atroroseus              ------MASSSM---------KPQDVLAHFTYLQWKDDFLKERPIEVQHD---QDPRKTN      42
QTE75998.1-ScyL2                     ---MATETVTIH---------QPKDALAKFTYLEWHDHYRTERPFQALDILHNAVDKREG      48
EED18844.1-TsRbtG                    ---MATAKITTQ---------HTYDIPAQLTYLEWHDHYETEKPFMVIRYPDDPPEMTGG      48
EED18849.1a_mod-TsRbtB               MATTTTLATTSK---------NGGDVPAKLTYIEWHDHYETEEPHFVLNTPDDPPDAYAG      51
B6HLP7.1-ChyM                        ---MGSITPSLHRGKFRYLTRGAKPAQSKEAYLLPPL-------------------SE--      36
S0E2Y4.1-Des                         ---MP------HKDNLLESPVG-KSVTATIAYHSGPA-LPT-----------SPIAGV--      36
B8N0E7.1-AsaB                        ---------------MRF---R-QQVETCLNYWPAEGPVQR-----------ELIL---G      27
AZL87943.1-AsaB                      ---------------MRF---R-QQVETCLNYWPAEGPVQK-----------ELIL---G      27
QTE75983.1-ZopL9                     ---MA------TATVTTA---P-TVVRTTADYYDAPPVLKI-----------HTYTRESY      36
BBU42017.1-ZopK                      ---MA------TATVTTA---P-TVVRTTADYYDAPPVLKI-----------HTYTRESY      36
RDW56970.1-C.crateriformis           ---MP------HNTSLDT---H-TPVEAFIDYYQPNA--DG-----------SPPDGMDL      34
BBG28508.1-PhiK                      ---MP------HSVEAE----Q-GVVEAFLEYYAPNS--DG-----------SLPDANDL      33
EED15414.1-T.stipitatus              ---MP------HSNY--------ETPEVILQYFAPNQ--DG-----------TPPDANDL      30
                                                                    :

A0A3G1DJG9.1-Mfr1                    -GDTNVKQPKEYEIQVHDIR-----KLDPQPTLLKNGYELVDIPTVVTDEQFIESGKSDE      85
A0A3G1DJF4.1-Mfr2                    -DSTNVI-RKPIPIEVEDAR-----TLSKSPTTKAEGYQLVNFHTKIPEEHFLNSK-LPE      85
OKL57227.1-T.atroroseus              ----MTWYKSGNAEIVRDAR-----GRESEYNVDTHGFTFVKHDTGLTT-----DDF--Y      86
QTE75998.1-ScyL2                     ----NVSFKEGGKEVVHDVR-----GHEQDFTLDKHGFLFANAPTSLSP-----SDF--Q      92
EED18844.1-TsRbtG                    ----NVTFKEGEEETIHDIR-----GHEDDFTLDGNGFLFTHAPTSLAP-----SDF--L      92
EED18849.1a_mod-TsRbtB               ----NVTFKEGEEEIIHDIR-----GHEDKFTLDKQGFVFTKAPTSLSP-----SEF--L      95
B6HLP7.1-ChyM                        -------FGDVLALPLVDMKPSLDLGDDSPYKLSVHGFTARRHHSALHAAPYERRSW--N      87
S0E2Y4.1-Des                         ---TTLQDCTQQAVAVTDIR-----PSVSSFTLDGNGFQVVKHTSAVGSPPYDHSSW--T      86
B8N0E7.1-AsaB                        TVGAYRRPIDSRPVLIQDVR-----GQEGTFTLDIHGFQFIKHISQH-------VAS--F      73
AZL87943.1-AsaB                      TVGAYRRPIDSRPVLIQDVR-----GQEGTFTLDIHGFQFIKHISQH-------VAS--F      73
QTE75983.1-ZopL9                     EEQFGNKSVIHHPINLKDIR-----AAN--INLQNNGFQLIKLQSKLTNP----DDY--L      83
BBU42017.1-ZopK                      EEQFGNKSVIHHPINLKDIR-----AAN--INLQNNGFQLIKLQSKLTNP----DDY--L      83
RDW56970.1-C.crateriformis           EIQYGQKAIDHHPVKMRDLR-----AGN--FTLEKNGFQLVPHHSKV-------TDF--N      78
BBG28508.1-PhiK                      EVQYGRKNLDLKSVKVGDMR-----SRD--FKLEEDGFTLMKHESAM-------TDF--T      77
EED15414.1-T.stipitatus              EIQYGTKNLHLAAVTLNDLR-----PDKDKITLTDHGLQLINHDTAMS-Y----EDF--N      78
                                                    : * :           .   .*       :

A0A3G1DJG9.1-Mfr1                    GNAYIKDVYFAECKRIIEEVSGGVDLIIPVSFRMREQKGEKESTT---------------      130
A0A3G1DJF4.1-Mfr2                    NKELIEEVYFDECRRLVQEVTGAAEA-YPYVYRVRNQEQNAKES----------------      128
OKL57227.1-T.atroroseus              DRNQVIERYIPQAIDLYKKVFGDVDEVYIFHWQMVSKLRST-------------------      127
QTE75998.1-ScyL2                     DDEKIKEKYLPECETYLKQYFDNVDQVHIIHYRVRCTN-SS-------------------      132
EED18844.1-TsRbtG                    DDEKIKTKYLPECEAYLKGLLD-ADQVIIFHYRVRNTITSD-------------------      132
EED18849.1a_mod-TsRbtB               DEEKIKEKYLPECEKYYREYFKGIDEVVFIHYRARNSITAD-------------------      136
B6HLP7.1-ChyM                        DEHLLREIYFPEVQELVQKVTG-CKKVVVSAAVLRNELYTEGDPASATQTEQNGQDAVKK      146
S0E2Y4.1-Des                         DPVVRKEVYDPEIIELAKSLTG-AKKVMILLASSRNVPFKEPELAPPYPMPGKSSSGSKE      145
B8N0E7.1-AsaB                        DEAS--------------------------------------------------------      77
AZL87943.1-AsaB                      DEAYVKTHMYPEAESILKNVTG-ATRAHVFSHITRTAPYESVEAMA--------------      118
QTE75983.1-ZopL9                     DEETVKRVYIPELAEAVKKLTG-ATEVRVLNPKVRDSSTEKDGFE---------------      127
BBU42017.1-ZopK                      DEETVKRVYIPELAEAVKKLTG-ATEVRVLNPKVRDSSTEKDGFE---------------      127
RDW56970.1-C.crateriformis           DKETVKRDYYPEVGETIRKYTG-ASKVYVLTHNTRSSAVPKMDLA---------------      122
BBG28508.1-PhiK                      DREKVKKEYYPEVAEAIKKHTG-ASKVICFNHNVRSTALPGLDLQ---------------      121
EED15414.1-T.stipitatus              DPEKIKSQYYLEVAEAIKKATG-AQKVICFNHNVRSEAAPRLDIS---------------      122
                                     .
                                                                  HXDXn
A0A3G1DJG9.1-Mfr1                    ------------KKL-GNIESRYAPRPVAHLDRDTPTAITVLEETV--------------      163
A0A3G1DJF4.1-Mfr2                    -------------NK-SNFH--TDFVPIVHVDRDDVTAPQRLRASL--------------      158
OKL57227.1-T.atroroseus              ----------------AQQRKLREQNKDVHVDQTQSEVERHVRNLL--------------      157
QTE75998.1-ScyL2                     ----------------DPN-SPTGPAKVVHVDQSGPHVTERIHKAF--------------      161
EED18844.1-TsRbtG                    ----------------DPY-SDTGPARSAHVDLSSQTIRERIRNRY--------------      161
EED18849.1a_mod-TsRbtB               ----------------DHN-SPTGPARVAHVDLSGPEINARIRKAF--------------      165
B6HLP7.1-ChyM                        ---DMSEL--FPPIVGNSPTDGICPAPKVHLDLTPKGARYHIRKYHREVTSAAEQVIEAE      201
S0E2Y4.1-Des                         REAIPANELPTTRAKGFQKGEEEGPVRKPHKDWGPSGAWNTLRNWSQELIDEAGDIIKAG      205
B8N0E7.1-AsaB                        ----------------------------------------VLKDNM--------------      83
AZL87943.1-AsaB                      ----------DSADPDAKATSVMVPARHVHVDQSESGAFEVLKDNM--------------      154
QTE75983.1-ZopL9                     ----------------NNWGKNNGAVRRIHIDLAPGGVEEALYPIF--------------      157
BBU42017.1-ZopK                      ----------------NNWGKNNGAVRRIHIDLAPGGVEEALYPIF--------------      157
RDW56970.1-C.crateriformis           ----------------TQKIEHVGPMRRCHVDVAPGGVEAAVIKRI--------------      152
BBG28508.1-PhiK                      ----------------KQKVDHVGPMRRVHIDVAPGGVSEAVTKRA--------------      151
EED15414.1-T.stipitatus              ----------------KHKVDHIGPMRRVHVDVAPRGTYEAVEKRA--------------      152
                                                                              :

A0A3G1DJG9.1-Mfr1                    ----------------GKEKAQELLSKHKRWAQVNVWRPIGNPATMWPLCFLNHDRIPTW      207
A0A3G1DJF4.1-Mfr2                    ----------------GAEKADMLLSKYKSYGSINVWRPVKNMVQKWPLMLVDHKSIEDW      202
OKL57227.1-T.atroroseus              -----------------PDRA--EYLLKGRVRNVNMWRPVRGPVQNWPLAVCDSRSVSAE      198
QTE75998.1-ScyL2                     -----------------PDCA--DFLLRGHVRLINLWRPINGPIQNWPLAVCDANSLPEE      202
EED18844.1-TsRbtG                    -----------------PDRA--DFLLSGRVRLINLWRPINGPIQNWPLAVCDGNTLPEK      202
EED18849.1a_mod-TsRbtB               -----------------PDRA--DFILRGRVRLVNLWRPINGPLQNWPLCVADCNSIQEK      206
B6HLP7.1-ChyM                        NRLLESGVQWDDLKDYYQGKE--SDDGVPRFALFSIWRPLK-PVHRDPLALASCASFPES      258
S0E2Y4.1-Des                         DEAA----------KLPGGRA--KNYQGRRWALYTTWRPLK-TVKRDPMAYVDYWTADEE      252
B8N0E7.1-AsaB                        ----------------TALEA--EHLLKTRWAIVNIWRPLK-PVPRDPLAVSDARSFHDK      124
AZL87943.1-AsaB                      ----------------TAVEA--ERLLKTRWAIVNIWRPLK-PVPRDPLAVSDAGSFHDE      195
QTE75983.1-ZopL9                     -----------------GEEY--MKSIAGRWRLINAWKPTR-PVERDPLAVC--DRVPDE      195
BBU42017.1-ZopK                      -----------------GEEY--MKSIAGRWRLINAWKPTR-PVERDPLAVC--DRVPDE      195
RDW56970.1-C.crateriformis           -----------------GEDF--MKPLAGRWKIVNAWKPVK-TVERDPLAVA--DLIPDE      190
BBG28508.1-PhiK                      -----------------GEEF--MSKFQGRWKIMNAWKPIR-TVERDPLGVAVAASVPDE      191
EED15414.1-T.stipitatus              -----------------GEAL--MSSIRGRWKIINSWKPLK-TVQRDPLAIATGPSCPDE      192
                                                                       . *:*        *:
```

```
A0A3G1DJG9.1-Mfr1                NYDTHV-----GHV--------WSLNDPRVSD--------R-----GQKTYDCVVKHDDR      241
A0A3G1DJF4.1-Mfr2                DYSTHM-----FTL--------HSSNDERVAT--------R-----GAKEHETILTHDKR      236
OKL57227.1-T.atroroseus          KYVEFDRILPKGNMTA------------------------------RMVL------EHPT      222
QTE75998.1-ScyL2                 NLIETDRIRKAQKGNT------------------------------RFVI------QAPS      226
EED18844.1-TsRbtG                NLVLTERIRTRDKAIA------------------------------RFVV------HSPA      226
EED18849.1a_mod-TsRbtB           HLVATKRIRKTHQAVT------------------------------RLVV------HEPT      230
B6HLP7.1-ChyM                    DYVPSEQLEPMDHQISAHLSRIIDPNAPKLSVENERQVQGGTCQTYGYLAYGPRDKEQKA      318
S0E2Y4.1-Des                     DGVSFWRNPPGVHGT------------------FESD-----------VLL---TKANPK      280
B8N0E7.1-AsaB                    DLLEIYGRVPGRQAK------------------KDYDA-ATKGSGFGMLY----GKYSPG      161
AZL87943.1-AsaB                  DLLEIYGRVPGKQVK------------------KDYDA-ATKGSGFGMLY----GKYSPG      232
QTE75983.1-ZopL9                 DLVPLQRVVPGKALM------------------EQR---------YHLKT------GKKD      222
BBU42017.1-ZopK                  DLVPLQRVVPGKALM------------------EQR---------YHLKT------GKKD      222
RDW56970.1-C.crateriformis       DLLAIQRFRADGSIS------------------EER---------YMVKAA---SPGKEE      220
BBG28508.1-PhiK                  DLINLKRYRPDGSLS------------------ESR---------YAVKA-------GEG      217
EED15414.1-T.stipitatus          DLVELTRYRPDGSLS------------------ESR---------YSVLY-------DQR      218
                                 .

                                                                           H
A0A3G1DJG9.1-Mfr1                YDYHYVSDLRPEECLVFCSFDSI------------------PKYAMPHSAFWDNNVPADA      283
A0A3G1DJF4.1-Mfr2                YRYIYASDMTPEEAWLFFAFHSD------------------PALGIPHGAFWDDSTKEEA      278
OKL57227.1-T.atroroseus          LKWHYLSQQQPDEAIIFKCTDS---HEG-------------VAKCVPHASIELPNTNSGT      266
QTE75998.1-ScyL2                 MKWYYQSKMEDNTLLVFKSYES---QDG-------------VAKYASHCSFPLPTAGPMT      270
EED18844.1-TsRbtG                MKWYYQSGMEDGTLLVLKNYDSHAEEGG-------------VARYSAHCSFPLPTAGPDT      273
EED18849.1a_mod-TsRbtB           LKWYYQSGMEDDTLLVLKNYDS---EDG-------------VAKYVPHCSFSLPTATAST      274
B6HLP7.1-ChyM                    HGWHFVSEQQPSDVLIIQLFDNEMEAHARAPQEGTNKMSNLGVGGAIHSAFELEGQDIDA      378
S0E2Y4.1-Des                     HKWYWISDQTPDEVLLMKIMDTESEKDGSE-----------IAGGVHHCSFHLPGTEK-E      328
B8N0E7.1-AsaB                    QQWFYMSDMKPDEALLIKCYD---SKDDGR-----------TARRTPHTAFVDPRTRDVK      207
AZL87943.1-AsaB                  QKWFYMSDMKPDEALLIKCYD---SRDDGR-----------TARRTPHTAFVDPRTRDVK      278
QTE75983.1-ZopL9                 HDWYYASNQQPDEVLLFTQYS---DFPNRN-----------TADRVPHVSVKLPGQED-K      267
BBU42017.1-ZopK                  HDWYYASNQQPDEVLLFTQYS---DFPNRN-----------TADRVPHVSVKLPGQED-K      267
RDW56970.1-C.crateriformis       HSWYFAPEQRPDELLLFNQYS---DKADRG-----------IADRVAHAAFVLPGTED-K      265
BBG28508.1-PhiK                  HQWYYVPQQRTDEMILFTQYS---DNPNRG-----------IADRVAHCAFILPGTED-K      262
EED15414.1-T.stipitatus          HKWYWVPMQKPNEMLLFNQYS---DDPNRT-----------LADRVAHCGFTLPGAED-K      263
                                   : :           ::                             * ..

                                   RXS
A0A3G1DJG9.1-Mfr1                PNRRSIEVRSLVFF-------------------------     297
A0A3G1DJF4.1-Mfr2                LTRCSIEVRIWVFFD------------------------     293
OKL57227.1-T.atroroseus          PARESIEIRAYLFSYPKNEA-------------------     286
QTE75998.1-ScyL2                 PPRESIELRAFVFTYPRDEST------------------     291
EED18844.1-TsRbtG                PPRESVEVRAVVLNYPRDDPSLVEPKPAISNVDSASITV     312
EED18849.1a_mod-TsRbtB           PPRESIEVRAFLFNYPRNEKSV-----------------     296
B6HLP7.1-ChyM                    EARESIEVRCAAFW-------------------------     392
S0E2Y4.1-Des                     EVRESIETKFIAFW-------------------------     342
B8N0E7.1-AsaB                    VARESLELRCLVFFEDQPLA-------------------     227
AZL87943.1-AsaB                  EARESLELRCLVFFEDQPLV-------------------     298
QTE75983.1-ZopL9                 PRRTSVDARCLVVW-------------------------     281
BBU42017.1-ZopK                  PRRTSVDARCLVVW-------------------------     281
RDW56970.1-C.crateriformis       PTRESIEVRALVVY-------------------------     279
BBG28508.1-PhiK                  PVRESVEVRALVVF-------------------------     276
EED15414.1-T.stipitatus          EIRESVEVRALVIY-------------------------     277
                                   * *:: :    .
```

Figure S46: A clustal omega[9] alignment of various asaB-like αKGDDs (domain IPR044053), including those identified in all known and putative maleidride BGCs. An asterisk (*) marks completely conserved amino acid residues, whereas a colon (:) identifies highly conserved residues and a period (.) indicates moderately conserved residues. The conserved $Fe^{+}$ binding H-X-D/E-$X_n$-H motif is shown in red.[16] The R-X-S motif, thought to bind αKG, is shown in blue.[17] Two publicly available protein sequences for AsaB from *A. flavus* are shown in this alignment (differing in annotation, but representing translations of the same gene from different strains). As can be seen in the alignment, the protein sequence with accession number B8N0E7.1 is missing the highly conserved iron-binding motif. It is therefore considered likely that this annotation is incorrect. The protein sequence with accession number AZL87943.1 was therefore used in the phylogenetic analysis and is shown in the percentage identity matrix.

```
EED18830.1a_mod-TsRbtU           ------------------------------------------------------------ 0
EED18846.1-TsRbtE                ------------------------------------------------------------ 0
TfL12-T.funiculosus              ------------------------------------------------------------ 0
Q2TXF3.1-OryG_(oryzines)         ------------------------------------------MHSTKVTY---------- 8
KIN05358.1-O.maius               MAPSAIESPVVNITEAPPQPKWFVPELLPESVKERMEKAGIDMTEYPTPPVVPFYYQEAV 60
PVH77204.1-Cadophora             MAPSAIEPT---VTDVPPQPKWHVPA-CPDTLKERLSKAGIDESTYPGPPTIPYYYQDAM 56
sA0A0A2IJP3.1-CnsP_(communesin)  MST-------------------TTVITPGTITREKNENGA----PLYPDYMPFYDPLEK 36

EED18830.1a_mod-TsRbtU           ----------------MSGDA-----ERRDLTIVANKVGAGADVLGFDFDTM-----PSH 34
EED18846.1-TsRbtE                ----------------MV-QN-----GAGTIQVIPIQASCGADIIGFDFEHL-----YPD 33
TfL12-T.funiculosus              ----------------MV-QN-----GESAVSVVPIEASCGADIVGFDFEHL-----PPG 33
Q2TXF3.1-OryG_(oryzines)         ------------------PEPMQLSGILDQYESFQVTPCIGTEFPKANLAEWLHSPNADA 50
KIN05358.1-O.maius               TLKGDPWEYNDVAAR---ADKTKSSLLSAATKVTNLTAHIGTEIEGLQLKDL-----TDQ 112
PVH77204.1-Cadophora             AVKKDPWEYNDAGIRAMQTDKTKSSLLNAATKVTDLTTNIGTEIEGLQLKDL-----TSQ 111
sA0A0A2IJP3.1-CnsP_(communesin)  VEDIGAFEHFDPGHR---ADPKLPNLLKNATKVWELSPHVGTEIHGVQLSQL-----DSA 88
                                                    :                    *::.    ::

EED18830.1a_mod-TsRbtU           QVGALRSAWLKYGILRFRGYDLTDE---HQLKLTKLIGSFLKREEDGAP------TTYKD 85
EED18846.1-TsRbtE                QVDAVRAAWRDYGVLRFRGYDITTQ---QHAKFSNLFGHYVPVKGT--S------IAHHD 82
TfL12-T.funiculosus              QVETVKQAWRDYGVLRFRGYDITTQ---QHVNFSNLFGRHVPVKSA--S------IGHEE 82
Q2TXF3.1-OryG_(oryzines)         LLRDLAITIAQRGVVFFRAQTDLDGE--LQKELTHRLGVQSGKPAGHRLSKHPLHLIRKD 108
KIN05358.1-O.maius               QRDELALLIAERSVVFFRDQDLSPQ---KQEELGKYWGRI----EYHPHVPHV---PGVP 162
PVH77204.1-Cadophora             QRDELALLIAERCVVFFRDQDITPQ---QQEELGKYYGRV----EIHPHVPHV---PGAE 161
sA0A0A2IJP3.1-CnsP_(communesin)  GLDELALLAAQRGALVFRDQDFVNIGFDAQKKLVSHFGPL----HIHGWAPHP---AAGS 141
                                         :      .    : **               : ::     *
```

```
                                                                    HXDXn
EED18830.1a_mod-TsRbtU               DEKVTVMT-NL--ING--VPSGAGSNVELEWHTDSWFWEYPPVGEILRAMEL-PQTGGDT 139
EED18846.1-TsRbtE                    QKEITVISNAK--VDG--KPVGTLGNVDLEWHTDSWYFDKPPCGQILRALEL-PRTGGDT 137
TfL12-T.funiculosus                  QEEITVISNVK--VDG--KPIGTLGSVDLEWHSDSWYFDKPPCGQILHALQV-PRIGGNT 137
Q2TXF3.1-OryG_(oryzines)             DPEMGVLDPGRQQKLHGVENTQKRQRAVLEYHSDGSYEVCPPDFTMLRMTEI-PPTGGDT 167
KIN05358.1-O.maius                   GAS-VVWDGLK--PGVPRKSTYRNPGGTNRWHSDTSHEPQTPSYTHLHLDAL-PSTGGDT 218
PVH77204.1-Cadophora                 GAT-VIWDALK--SEGRKSGTFRNPGGTYRWHSDIAHERQPPAYAHLHNDTI-PSTGGDT 217
sA0A0A2IJP3.1-CnsP_(communesin)      EEHMIIYDH-K--DDLRVRQSW-AGRSPVQWHTDQSPEQQPPGTTFIAMLESPTTAGGDT 197
                                          :                       .:*:*       *    :         **:*

EED18830.1a_mod-TsRbtU               YWADMYAVYDALPEDLRSTIEGRLIQFDTVYNGHG------NLRKGKEAPKTDDFRLWEH 193
EED18846.1-TsRbtE                    YWVNMYAVYDALPEFTRKIIEGRLIQFNIVYDAVG------RVRPGQEKPETDDFRLWKH 191
TfL12-T.funiculosus                  YWVNMYAVYDALPESTRKIIEGRLIQFDIVYDGYG------RLRPGQEKPEEEDFRLWKH 191
Q2TXF3.1-OryG_(oryzines)             LWASGYELYDRLSTPYQKFFESLTAQHEVPSLRKLAETEPGIYDGPRGAP--ANTDMQFK 225
KIN05358.1-O.maius                   VWASGYAAYDKLSPAFREFIDGKMAVFSSTHTYID-----------RNDP--YAGPKFIQ 265
PVH77204.1-Cadophora                 VWASGYAAYDKLSPAFRQFIDGKKAVFRSTHSYVD-----------RDDP--HGARRYNE 264
sA0A0A2IJP3.1-CnsP_(communesin)      LVSSSVRAYSSLSPRFRKRLEGLTAIHTNNDGVSQ---EL----KHGQQA--VMRRGVLQ 248
                                        .    *. *    :. ::.    .                                .

EED18830.1a_mod-TsRbtU               IRHPIIRTHPESGRKAVFVGQSKHEKNWIVGLPLEESKEILAKILSYVEK-PEFQLHQKW 252
EED18846.1-TsRbtE                    VRHPIVRTNPESGRKAVYIGYFDSTKNWIVGLPLEQSKAILEEIYSLIDS-GKFVFQQKW 250
TfL12-T.funiculosus                  VRHPIVRTNPYSGKKAVYVGYLNAERNWIVGLSLETSTAILNEIFSYINS-GKYVFKQVW 250
Q2TXF3.1-OryG_(oryzines)             QSHPMVRTHPVTGWKTLFAGGLHCR--RVNDVTDFESEQLLSKIISLVGDNHDLQVRFRW 283
KIN05358.1-O.maius                   NIHPVVRVHPVTGWKSLWVNRGYTR--RIVGLEPGESDAILNYLYNVYESNLDIQVRFKW 323
PVH77204.1-Cadophora                 NIHPIVRVHPVTGWKSLWLNRNYTQ--RIVGLEKAESDAILNYLYDVFEHNLDIQVRFKW 322
sA0A0A2IJP3.1-CnsP_(communesin)      AEHPVVLVHPVTKQKALYVNPVYTK--KIVGFDQEESDCILKFLFDHIAKRQDFSCRIRY 306
                                       **:: .:* :  *::: .        : ..    *  :*  : .      .   :  :

                                                    H           R
EED18830.1a_mod-TsRbtU               Q-PGDTVIWDNRCTMHRRE-TWPDDQTRIMHRTTCNTKGQPRPFYVY------------- 297
EED18846.1-TsRbtE                    Q-PNDIIMWDNRCTMHRRD-GWNETDMRIMHRTGT---GTETPIYVY------------- 292
TfL12-T.funiculosus                  Q-PHDIVMWDNRCTMHRRD-GWEGNDARVMHRTGT---GMETPIYVC------------- 292
Q2TXF3.1-OryG_(oryzines)             NNPGDVAIWDNRCVLHCPTQDHYGLGGRMGYRTMG---IAEKPYLDPNSPSRQEALAAAA 340
KIN05358.1-O.maius                   T-PRSSALWDNRITTHNAMWDYEGKEPRHGTRVMT---LGERPYFDKDAPSRR-----QA 374
PVH77204.1-Cadophora                 T-PNASALWDNRVTIHNAIWDYEGREPRHGTRVMT---LGEKPYFDENAVSRR-----QA 373
sA0A0A2IJP3.1-CnsP_(communesin)      E-AGTVLVWDQRVTNHSQTLDYPIGDRRHGFRLTP---LANKPIPAKIEEDDEEFSTDDA 362
                                            :**:* . *           *   *          *

EED18830.1a_mod-TsRbtU               --------         297
EED18846.1-TsRbtE                    --------         292
TfL12-T.funiculosus                  --------         292
Q2TXF3.1-OryG_(oryzines)             K-------         341
KIN05358.1-O.maius                   LGLDGPE-         381
PVH77204.1-Cadophora                 LGLEDSE-         380
sA0A0A2IJP3.1-CnsP_(communesin)      RHLVGNAS         370
```

Figure S47: A clustal omega[9] alignment of various TauD-like αKGDDs (domain IPR042098). An asterisk (*) marks completely conserved amino acid residues, whereas a colon (:) identifies highly conserved residues and a period (.) indicates moderately conserved residues. Residues of the conserved $Fe^{+}$ binding H-X-D/E-$X_n$-H motif are shown in red.[16] The invariant arginine residue, thought to bind αKG, is shown in blue.[18]

```
A0A159BP93.1-CitB      -----------------MPISTKSSFYL-----------PAVDISPYLQDPNSDAARKVI    32
ASK38712.1-PvL5        ----------------MASKETFSK-YPAFPDNIPTAAVPKISLRQILSRD-----PTVS    38
Q4WKX0.1-FgnB          MTVNGKDIDSPNAQYVAAGIDMSDLFPAPFPTNVKTVHLETLSLAKLLQRD-----EDEL    55
                                               .  .                    :.:    *.

A0A159BP93.1-CitB      DDVRAACTSTGFFQLLGHGI--SPALQQSVFAA---AAKFFALPS-D----VKSRCRNVG    82
ASK38712.1-PvL5        KRLVDAGKEFGCFKVDLTDAIDGPVLCQGVERGFDLGKAFF-DQDIETKKAYKLSHENVG    97
Q4WKX0.1-FgnB          RRIYENCKDPGFFQLDLTDDEQGVQLLQDAVDCARLMKQLLPNMSVEEKRMYKQ-HSRVG    114
                         :    .. * *::    .    .   * *..          ::    . :      *    .**

A0A159BP93.1-CitB      FRGYDPMASQSYELGVLPDLKEGFIAGKDIPL------DDPRVASQRFFMGQNAWPPSEL    136
ASK38712.1-PvL5        YKQA---GVLVI--TKER---RDQVETCSV---SRDDLAASRPDL-----------P---    132
Q4WKX0.1-FgnB          VYSK---GYQVY--DVLPNGQPKYNETVNFPMTEMLGYGDSTVDL-----------P---    155
                               .                  ..                            *

A0A159BP93.1-CitB      LPEANFRRPIEEYYQAMLK----LCWVVLD--LVAATLPYGPHVFDEFKENDPACPLRLL    190
ASK38712.1-PvL5        ----SVFDQQRLLLQGLVAQLGQLSHLAVYHLSEGLGLDYGVVSARHDPRDQSATMIRFL    188
Q4WKX0.1-FgnB          ----DWLSPHRELFQRTMRSGNKIANIVLAALEVGLQVPRGALTDAHRIQDPSDDFLRLL    211
                           .      .    *   :      :. :.:      .   :    *      .   .:      :*:*

                                                 HXDXn
A0A159BP93.1-CitB      HYPPAPAPDVA------KGRQLGSSAHTDFGAITLLLQDDHSGLEVQ-------------    231
ASK38712.1-PvL5        HNPPQRGPREELPSTEDPGSRAYLMGHSDGGTVTILFNV-LGGLQLQRQQP---------    238
Q4WKX0.1-FgnB          RYPGLQP--------GQPRDDLCFPAHKDFTSLGILFTW-LGGLQLLASASAPGVTSGTM    262
                       : *                        .*.*   :: :*:     .**::

                                                                    H            RXS
A0A159BP93.1-CitB      ----DCETGEWIGVPPNKDAYVVNLGDMMSRITRGHYKSSIHRVIN----QNLTDRYSVV    283
ASK38712.1-PvL5        -----DGSIEWQYIPPEPGCALIMVGDAFKSFTDGEVPSCVHRVIQPPGEQDRFDRYALG    293
Q4WKX0.1-FgnB          TGPLDIAEDAWRWVQPVPGTAIVNVGNALEILTNKALTSGLHRVVRAPGEQLPFDRYSVL    322
                                *   :  *   .   :: :*: :. :*      *  :***:.     *    ***::

A0A159BP93.1-CitB      FFFDGNLDYRLRPLDRVGQN-----------------WDEEDTLTVEEHMLERTTTTYNL    326
ASK38712.1-PvL5        FFLKPANGASIGPVPRRGV--------TE--------NGVNKASDYGEWAKNKNAALYNE    337
Q4WKX0.1-FgnB          VGTRPANSFPMKPLQPSPQISPVLDPAAAEIATMTSGQWGTHNIGSFNNWVKARTE-----    377
                       .       .   :  *:                               .  ..       :     :.

A0A159BP93.1-CitB      KVK------          329
ASK38712.1-PvL5        MRQENVAI-          345
Q4WKX0.1-FgnB          -RQEVLIIP          385
                             :
```

Figure S48: A clustal omega[9] alignment of PvL5 from the cornexistin BGC with two known IPNS-like (domain IPR027443) αKGDDs. An asterisk (*) marks completely conserved amino acid residues, whereas a colon (:) identifies highly conserved residues and a period (.) indicates moderately conserved residues. Residues of the conserved $Fe^{+}$ binding H-X-D/E-$X_n$-H motif are shown in red.[16] The RXS motif, usually present in IPNS-like αKGDDs, and thought to bind αKG, is shown in blue.[17] In PvL5 the serine of the RXS motif is substituted with alanine (highlighted in yellow). This may not impact catalytic function, as the serine at this position is frequently absent in αKGDDs, such as TauD-like αKGDDs (see Figure S46).

```
OKL57048.1-T.atroroseus     ------------------------------------------------------------    0
A0A2I1BSW6.2-NvfE           ------------------------------------------------------------    0
Q5AR53.1-AsqJ               MGYPKAFTSSDSEPEPDLSRDLGNPVMGNPGVVSRSSSTVAQHSVRNNPTGPDGRLAGLW    60
Q5AR34-AusE                 ------------------------------------------------------------    0

OKL57048.1-T.atroroseus     ---------------------------------MPTT---------IGAYTKEEPKSSKS    18
A0A2I1BSW6.2-NvfE           ----------------------------------------------MGRDQVSHKRSQNS    14
Q5AR53.1-AsqJ               NARALLRFPAEVNGVRLDFKSVSSRRPTSRLSLLPYSLCSICPSSQATMTSKDHVKSQIP    120
Q5AR34-AusE                 --------------------------------------------------MGSATPSRLQ    10
                                                                                .   *

OKL57048.1-T.atroroseus     PIPPVLELDAST-----CTSADLVSALKVAGGVIVRNILTAEEIQQIESDVRPWLEQD--    71
A0A2I1BSW6.2-NvfE           NVSEIPDLDSSLHRSEDTLVAEIIEAMTLAGVCVVRNLFTKSLVDQVLKDFEPHVSST--    72
Q5AR53.1-AsqJ               RLSAIN------------DLHKIWPTVEEHGAAIIESFLSLDIVRRLNEEVDPFVKIEPI    168
Q5AR34-AusE                 KFPATA------------PADEIYAAFKEDGCVIIEGFVPPDQMARFSQEIQPAMEKIQV    58
                             .                .:  :.    *  ::..:.  . : :. .:. * :.

OKL57048.1-T.atroroseus     -----KPWNGDFFPQGTRRAFGLVGKSKAFALRLVDHELWLQVVDALLTSENPNNWVGDK    126
A0A2I1BSW6.2-NvfE           -----KLFDG--YPGNGCHLTGLLSKSEIYAHMVVGNSVFEKVRNHF-LSTTFRSWIGGK    124
Q5AR53.1-AsqJ               PAAKTKDHPNHVLSTTTTRLVNVLAPISKAYREDVLNSKVLHRICSDA-----FHVY-GD-    221
Q5AR34-AusE                 QVTNDGNSND-R----VKRFSKLVTTSPTFRHEILENDLMHELLQRV-----FSKP-GEG    107
                                     .              *     *   :     ::   .:   .: .           *

                                             Q   HXDXn
OKL57048.1-T.atroroseus     NEVSVCKPQLNNTIVFSIGPGARDQSLHRDDQIHHQNHRAVAKHEPGRDTGIGFFVAGKK    186
A0A2I1BSW6.2-NvfE           MMTFTSPPQLDSTICSYINPQSPGEHLHRDDAIHYGWNEAASEYTVGRDISMSMFLALTE    184
Q5AR53.1-AsqJ               -------YWVLMGAVMELAPSNPAQPLHRDMRFSHPIVEYLKPDAP--ATSINFLVALSP    272
Q5AR34-AusE                 M-----GYHFNDTMVIEVQPGAPAQRLHRDQEL-YPWWNSMGPDAP--ECLVNFFCAVTP    159
                                    .          :  *       :  ****  :  :     .          :.:: *  .

                                                                                  H
OKL57048.1-T.atroroseus     TTRQNGATRFIPGSHLWDYAE-----GPAH--EDQTVYAELNPGDGFMVLSGCFHGGSAN    239
A0A2I1BSW6.2-NvfE           STRENGTTRFFPGSHLWDYSQ-----DFPSADDTRIRYAELHPGDCYFMLSSVTHSSTDN    239
Q5AR53.1-AsqJ               FTAENGATHVILGSHKWQNLSNVSM--------DATVRALMNPGDALLITDSTIHCGGAE    324
Q5AR34-AusE                 FTVENGATRLVPGSNRWPELTLINATDCPQYGKIDSVPAIMQPGDCYMMSGKVIHGAGHN    219
                             * :**:*:.. **: *                            * ::***   :: .    *  .   :
```

```
                                      R
OKL57048.1-T.atroroseus      KTENEERLVYSCFYTRSWLRQEENQYLANDKTKILELPNCLQERVGWGLSTPFLG-----      294
A0A2I1BSW6.2-NvfE            RSTNRPRVLAATIVTRSHLRQEENQYLTYDPITVGRFPTWLQRLVGYAPSAPFLG-----      294
Q5AR53.1-AsqJ                TTGTETRRLLTITMGISQLTPLES-NLAVPRPVIESLTPLAQRLLGWASQRSAAPRD-IG      382
Q5AR34-AusE                  ATLSDQRRALAFSTIRRELRPVQAFPLWIPMQIATELSPRTQAMFGFRSSTQHCDVDTVH      279
                              : .  *   :       *   :   *        :    *  .*:  .

OKL57048.1-T.atroroseus      --------------------------------------------   294
A0A2I1BSW6.2-NvfE            WVDKRDP--RCVI--------DPKAADDHCGGEYYETNEETLN   327
Q5AR53.1-AsqJ                LLTIRGNSIEKTMNLKAEQPLHDDEAEPLCRETI---------   416
Q5AR34-AusE                  FWGNDGKDIGEHLGLISSA------------------------   298
```

Figure S49: A clustal omega[9] multiple sequence alignment of PhyH-like αKGDDs, showing residues of the conserved $Fe^{+}$ binding H-X-D/E-$X_n$-H motif in red.[16] An asterisk (*) marks completely conserved amino acid residues, whereas a colon (:) identifies highly conserved residues and a period (.) indicates moderately conserved residues. The invariant arginine residue thought to bind αKG is shown in blue.[18] PhyH-like αKGDDs are known to contain a conserved glutamine residue (Q: in green),[19] which is also thought to be involved in binding αKG. NvfE has glutamic acid substitution at this position and is known to be an unusual, cofactor free, isomerase.[20] OKL57048.1 from *T. atroroseus* – which is putatively involved in maleidride biosynthesis – does contain the conserved glutamine (Q).

```
PvL7_ASK38710.1       --MHFNL--LLVLTVL--------------LRQATALVLPPSNST---------SNSTGK
A0A4P8GEA3_EUPF       --MRYHLSALVLVFTA--------------FRETLTAPTPGNNTI--------------P
A0A2U8U2M1_ASR5       MRRSFLISAALGLSMSTPALAASIQSVLGYLRPTSHHHAPCADDVVLKQSAGSDSAAPDP
                          : :   : :                 :* :     *  :

PvL7_ASK38710.1       FSARTVFQFPQGYWLENLAVRGNGQVLATTYMPSAGLYLIDPTPNASYPAVLVHQFENST
A0A4P8GEA3_EUPF       LPNRLLHQWPNGTWVENISVRPNGNLLVTTSTPDGSVWQVKEPWKENPEVERVFNFDEWV
A0A2U8U2M1_ASR5       LPSRVVHNWPNGTWIENISVRPNGNLLVSQSTPRGRVWQVKEPWLDEPKVELAYDFDEWV
                      :. * :.::*:* *:**::** **::*.:   * . :: :. .   .  .  ..:*:: .

PvL7_ASK38710.1       SALGIVEAEGTEPDTFYLATLNFSAADGFVPRTSQVWRVDMS-SFHYSPQTGQVSGKAAV
A0A4P8GEA3_EUPF       DRL--IGIGETQDDKYVVVGSRFYSTD---AQSSHVARTFCAMELDFSGNTTEPSARL--
A0A2U8U2M1_ASR5       DRI--IGIGETTPDKYVVVGSRFYSLD---PQSSQVERTFCAMELDFT-KGEKPSARL--
                      . :  :    *  *.: :.  .* : *   ..:*:* *.  : .: :: :  : *..

PvL7_ASK38710.1       SHVTTLSSVGMANGMTLLAPDSSHILIADSL--------------RGAIWDLDTATGHYG
A0A4P8GEA3_EUPF       --IAWMPESYLLQGVAALPWDRDTVLISDQYVLRPRAVQIDWTPSPGQIWVLDTRTGEYG
A0A2U8U2M1_ASR5       --VARFPHANLLQSVSALPWDRSVVLISDQYLLHPRADWEDLTPGPGQIWRLDTKTGHHE
                        :: :.   : :.:: *. * . :**:*.                * ** *** ** :

PvL7_ASK38710.1       LS-SSFPAMRSDNPARRFLGIDGVKVHQGSLYFNNAGEFTLARMPIHSDG-----IAKGE
A0A4P8GEA3_EUPF       LVMTDYAELNTTYAKGPDVGIDGIKIRDHDLFWVNQDDSGIYRVKIDDAG--VPVAPVKP
A0A2U8U2M1_ASR5       IVMTNYAEMNTTYNHGLDVGINGIKIHGDHLYWINMDTGGAYRVRIDKYGYPTPLNAVPE
                      :  :.:. :.:       :**:*:*:.   *:: * .     *: * . *      .

PvL7_ASK38710.1       PVVLATDLYSDGFCL--------YDADTVLVTMNIDNGLAALDLESHRRWMVAGNMPDGV
A0A4P8GEA3_EUPF       QLVASYNTMWDDMAF------DPFNENVIWATGLNAVFAATLD-GQIVPVDGVGTSDNLT
A0A2U8U2M1_ASR5       TLGVAEDALWDDFAMHGTRIGEESDDTTMFATSIVNLMAISPENGTIVPLAGVGTSEPMG
                       :  : :   *.:.:         :  .: .*        : :         .*.

PvL7_ASK38710.1       FTTPTSVELGRGEDAGKLAYVTMGGTYVATGAEDLVGGSLVVVDLKSATTDPKGTGRGGL
A0A4P8GEA3_EUPF       LPGPTACAFGRTEKDKSILYVT---------------GNLLTVPESLLDVKLGG------
A0A2U8U2M1_ASR5       FPGPTSAQFGRTEKDSHILYVT---------------GKLFNVPPSIRDVVIQG------
                      :. **:  :** *.   : ***               *.*. *  .   .   *

PvL7_ASK38710.1       LVQETETIWLEL
A0A4P8GEA3_EUPF       W-----------
A0A2U8U2M1_ASR5       WVRAIDTTGFHF
```

Figure S50: Multiple sequence alignment using MUSCLE[1] of PvL7 compared to two putative Diels Alderases, EupF[21] and Asr5.[22] An asterisk (*) marks completely conserved amino acid residues, whereas a colon (:) identifies highly conserved residues and a period (.) indicates moderately conserved residues.

### Phylogenetic analyses

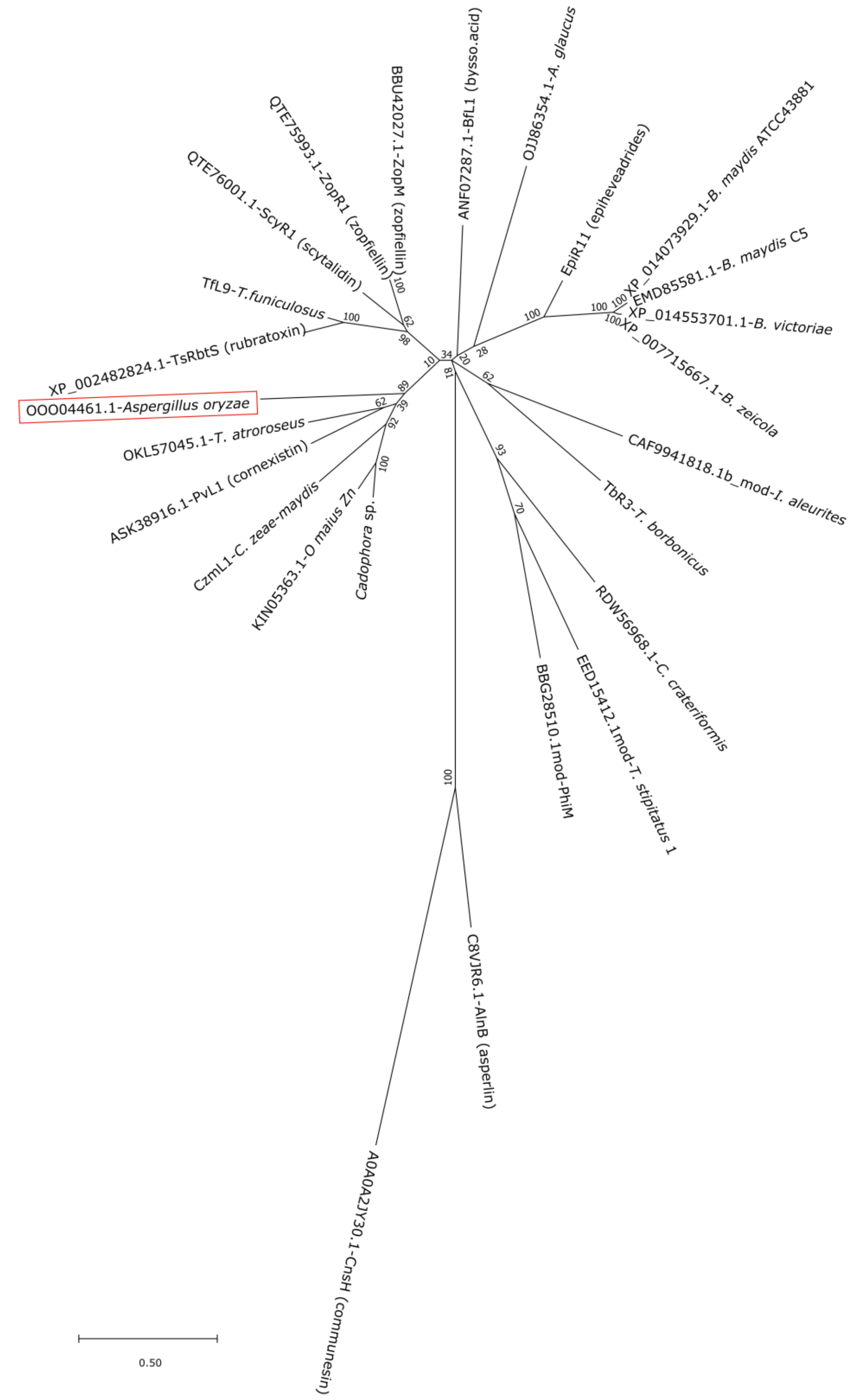

Figure S51: An unrooted phylogenetic analysis of the predicted maleidride polyketide hydrolases, with hydrolases from other polyketide BGCs (asperlin and communesin), and the putative homologue identified within the genome of *Aspergillus oryzae* BCC7051. This predicted protein clearly clades with the maleidride serine hydrolases. The evolutionary history was inferred by using the Maximum Likelihood method and Le_Gascuel_2008 model.[23] The tree with the highest log likelihood (-6067.26) is shown. The percentage of trees in which the associated taxa clustered together is shown next to the branches. A discrete Gamma distribution was used to model evolutionary rate differences among sites (5 categories (+G, parameter = 1.8561)). The rate variation model allowed for some sites to be evolutionarily invariable ([+I], 5.18% sites). The tree is drawn to scale, with branch lengths measured in the number of substitutions per site. This analysis involved 25 amino acid sequences. There were a total of 193 positions in the final dataset. Evolutionary analyses were conducted in MEGA X.[24]

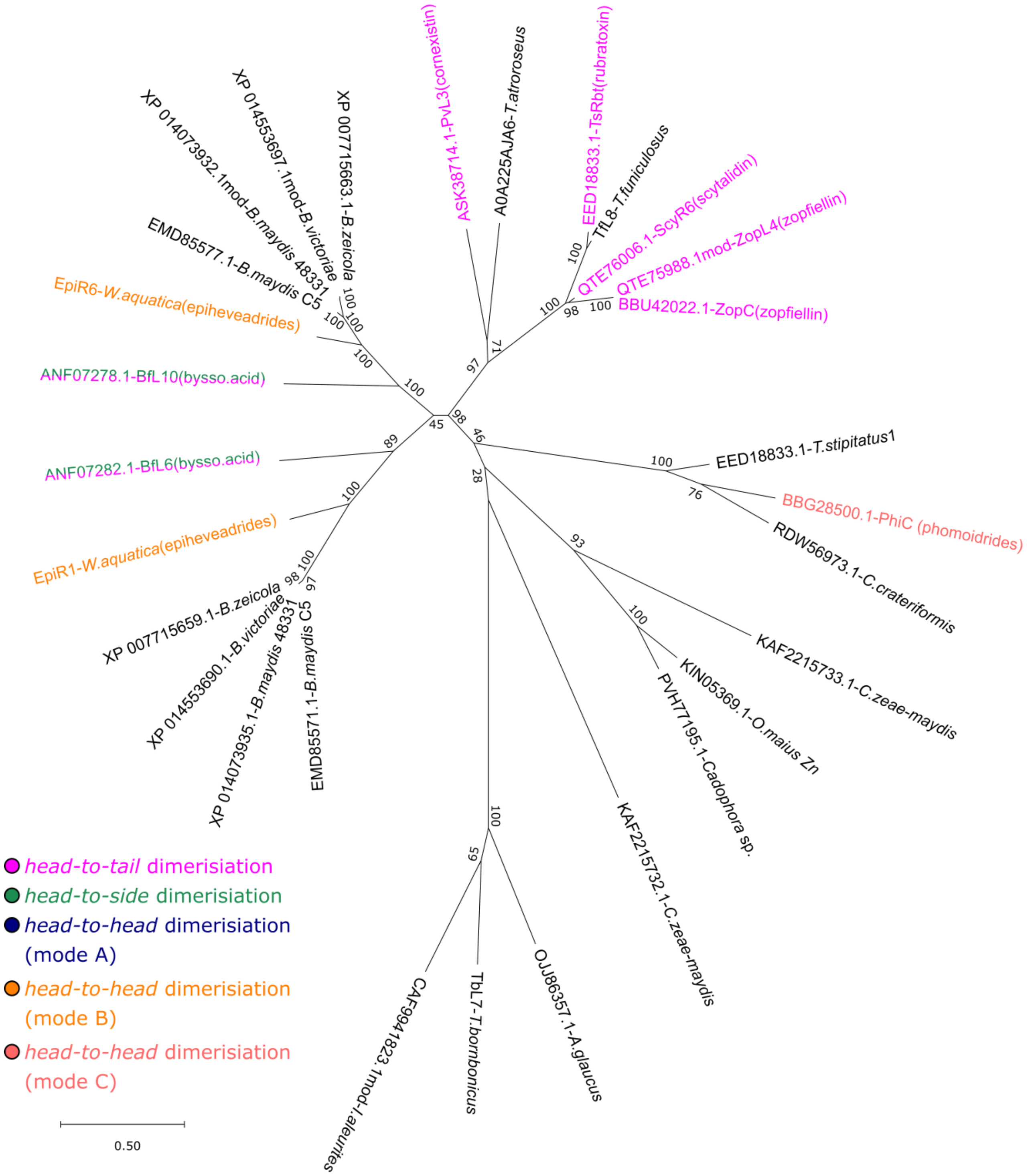

Figure S52: An unrooted phylogenetic analysis of maleidride dimerising cyclases. MDCs do not appear to clade according to mode of dimerisation. The evolutionary history was inferred by using the Maximum Likelihood method and Le_Gascuel_2008 model. [23] The tree with the highest log likelihood (-6093.34) is shown. A discrete Gamma distribution was used to model evolutionary rate differences among sites (5 categories (+G, parameter = 1.2572)). The rate variation model allowed for some sites to be evolutionarily invariable ([+I], 5.34% sites). The tree is drawn to scale, with branch lengths measured in the number of substitutions per site. This analysis involved 29 amino acid sequences. There were a total of 178 positions in the final dataset. Evolutionary analyses were conducted in MEGA X.[24]

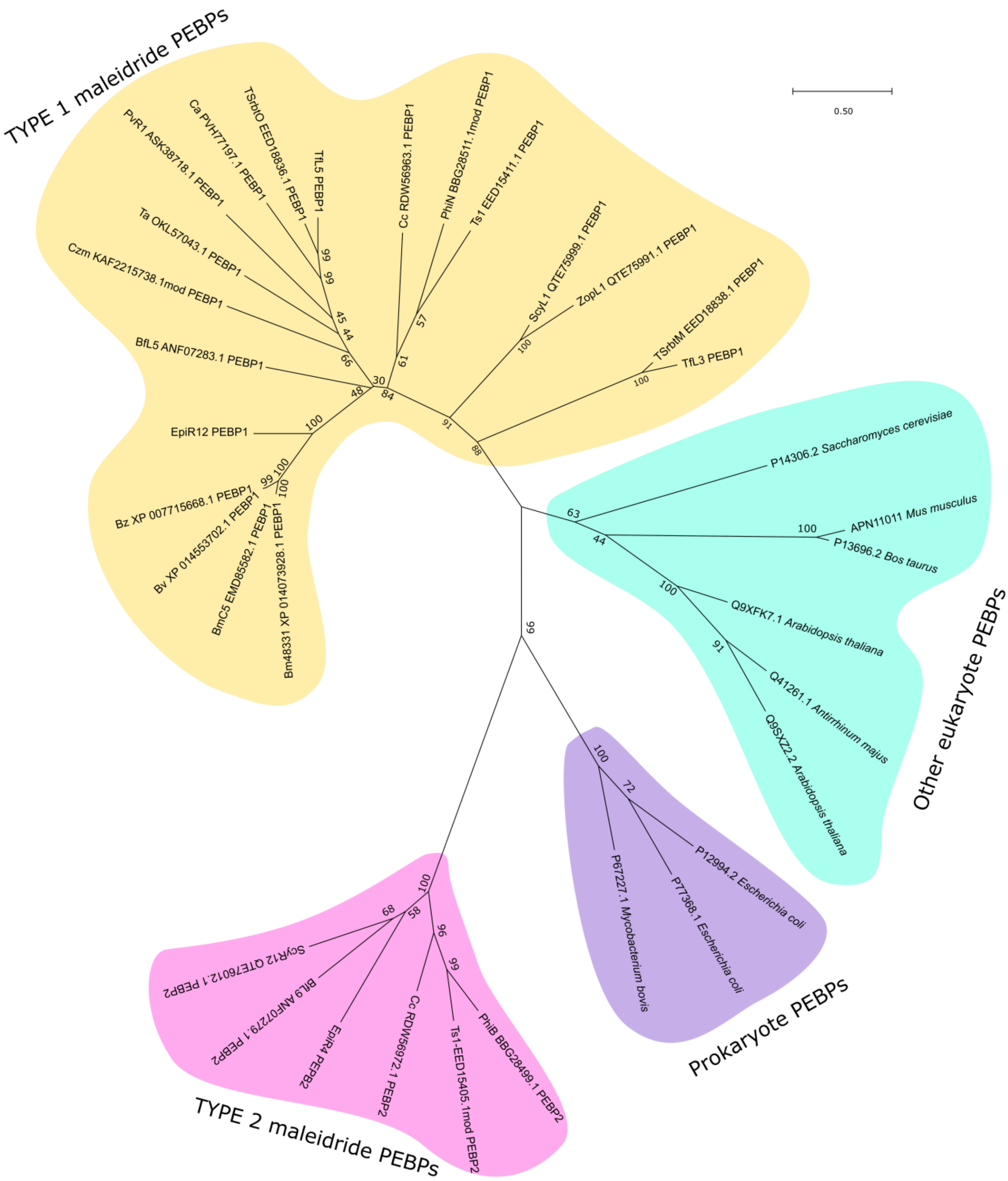

Figure S53: An unrooted phylogenetic analysis of PEBP sequences from maleidride BGCs and from characterised eukaryote and prokaryote PEBPs. The initial multiple sequence alignment was conducted using T-coffee.[25] Evolutionary analyses were conducted in MEGA X.[24] The evolutionary history was inferred by using the Maximum Likelihood method and Whelan and Goldman model.[26] The tree with the highest log likelihood (-6440.32) is shown. The percentage of trees in which the associated taxa clustered together is shown next to the branches. A discrete Gamma distribution was used to model evolutionary rate differences among sites (5 categories (+*G*, parameter = 2.0270)). The rate variation model allowed for some sites to be evolutionarily invariable ([+*I*], 4.13% sites). The tree is drawn to scale, with branch lengths measured in the number of substitutions per site. This analysis involved 34 amino acid sequences. There were a total of 117 positions in the final dataset.

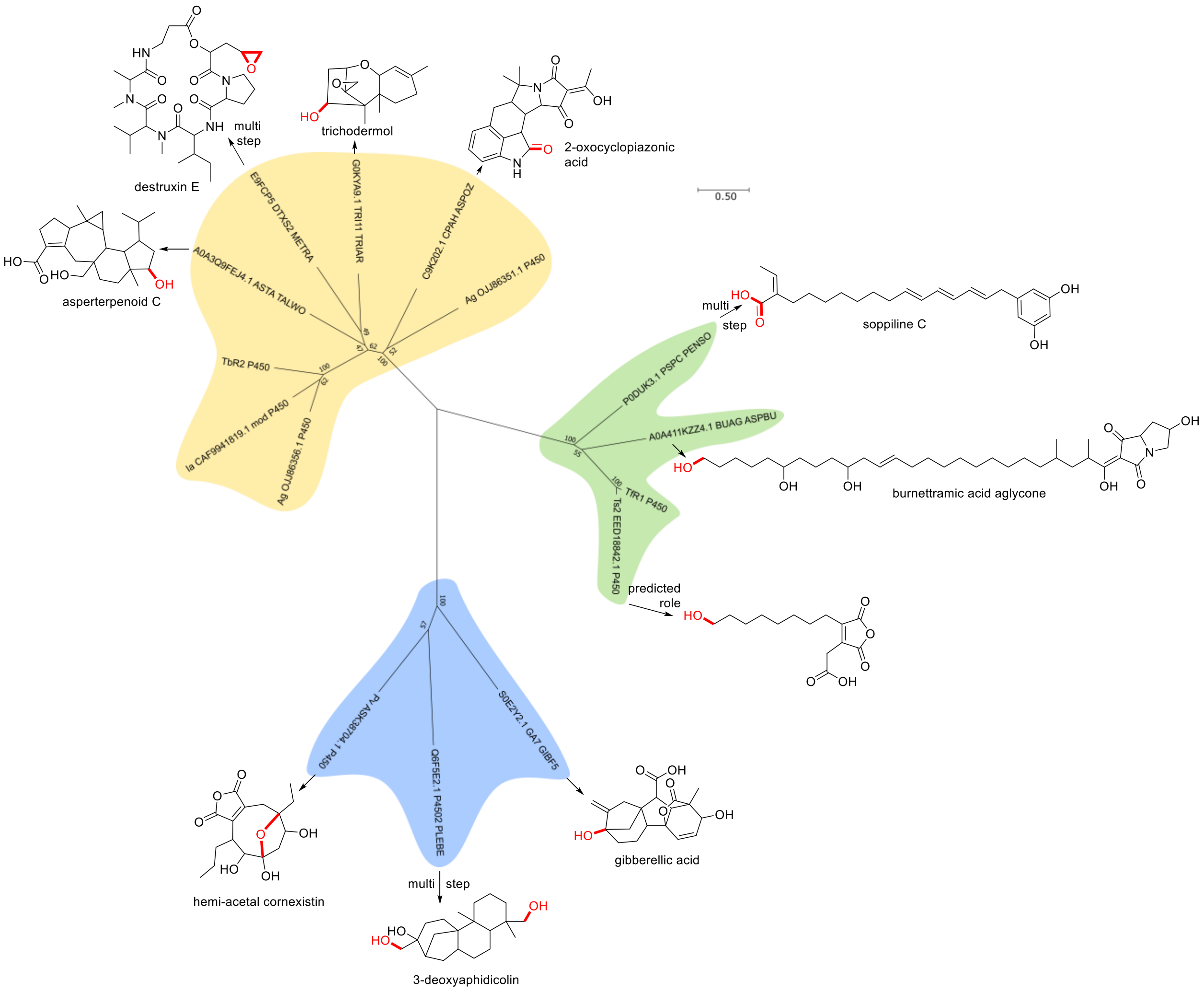

Figure S54: An unrooted phylogenetic analysis of the predicted maleidride CYP enzymes, with characterised CYPs. The evolutionary history was inferred by using the Maximum Likelihood method and Le_Gascuel_2008 model.[23] The tree with the highest log likelihood (-11238.81) is shown. A discrete Gamma distribution was used to model evolutionary rate differences among sites (5 categories (+G, parameter = 2.7828)). The rate variation model allowed for some sites to be evolutionarily invariable ([+I], 1.22% sites). The tree is drawn to scale, with branch lengths measured in the number of substitutions per site. This analysis involved 15 amino acid sequences. There were a total of 368 positions in the final dataset. Evolutionary analyses were conducted in MEGA X .[24]

### Percentage identity matrices

#### PKS

Table S4: Percentage identity matrix of maleidride PKSs generated using MUSCLE[1].

| | 1: | 2: | 3: | 4: | 5: | 6: | 7: | 8: | 9: | 10: | 11: | 12: | 13: | 14: | 15: | 16: | 17: | 18: | 19: | 20: | 21: | 22: | 23: | 24: |
|---|---|---|---|---|---|---|---|---|---|---|---|---|---|---|---|---|---|---|---|---|---|---|---|---|
| 1: EpiPKS | 100.00 | 63.12 | 63.12 | 62.87 | 62.62 | 48.30 | 51.75 | 50.76 | 51.28 | 50.34 | 44.43 | 53.24 | 47.52 | 47.61 | 47.39 | 47.15 | 47.94 | 48.88 | 47.76 | 53.87 | 52.94 | 52.94 | 47.38 | 47.15 |
| 2: BmATTC | 63.12 | 100.00 | 100.00 | 88.30 | 88.17 | 44.59 | 48.23 | 47.11 | 47.07 | 47.65 | 43.17 | 50.54 | 45.42 | 45.10 | 44.98 | 45.06 | 45.03 | 46.48 | 45.48 | 49.79 | 49.10 | 49.10 | 44.41 | 44.48 |
| 3: BmC5 | 63.12 | 100.00 | 100.00 | 88.30 | 88.17 | 44.59 | 48.23 | 47.11 | 47.07 | 47.65 | 43.17 | 50.54 | 45.42 | 45.10 | 44.98 | 45.06 | 45.03 | 46.48 | 45.48 | 49.79 | 49.10 | 49.10 | 44.41 | 44.48 |
| 4: BZ | 62.87 | 88.30 | 88.30 | 100.00 | 98.88 | 44.97 | 47.99 | 46.75 | 46.86 | 47.05 | 42.62 | 49.58 | 44.78 | 44.51 | 44.65 | 44.56 | 44.90 | 45.84 | 45.10 | 49.21 | 48.27 | 48.27 | 44.24 | 44.23 |
| 5: Bv | 62.62 | 88.17 | 88.17 | 98.88 | 100.00 | 45.10 | 47.95 | 46.75 | 46.90 | 47.01 | 42.79 | 49.46 | 44.87 | 44.55 | 44.65 | 44.51 | 44.95 | 45.84 | 45.18 | 49.29 | 48.39 | 48.39 | 44.24 | 44.31 |
| 6: Czm | 48.30 | 44.59 | 44.59 | 44.97 | 45.10 | 100.00 | 54.59 | 54.94 | 51.27 | 50.75 | 42.25 | 44.70 | 44.74 | 44.15 | 44.45 | 44.15 | 44.54 | 44.78 | 44.59 | 49.37 | 49.22 | 49.22 | 43.31 | 44.47 |
| 7: CadPKS1 | 51.75 | 48.23 | 48.23 | 47.99 | 47.95 | 54.59 | 100.00 | 72.18 | 56.90 | 56.28 | 45.54 | 48.23 | 48.50 | 46.98 | 47.11 | 47.83 | 47.73 | 47.92 | 47.24 | 54.97 | 54.44 | 54.44 | 46.94 | 47.58 |
| 8: OmPKS2 | 50.76 | 47.11 | 47.11 | 46.75 | 46.75 | 54.94 | 72.18 | 100.00 | 57.12 | 55.49 | 45.17 | 47.63 | 48.26 | 47.09 | 46.90 | 46.88 | 47.14 | 47.85 | 47.13 | 55.31 | 53.53 | 53.53 | 46.99 | 47.53 |
| 9: Ta | 51.28 | 47.07 | 47.07 | 46.86 | 46.90 | 51.27 | 56.90 | 57.12 | 100.00 | 55.55 | 45.35 | 49.10 | 48.95 | 49.51 | 49.88 | 49.33 | 49.25 | 48.73 | 48.77 | 55.62 | 55.25 | 55.25 | 47.45 | 46.77 |
| 10: Pvpks1 | 50.34 | 47.65 | 47.65 | 47.05 | 47.01 | 50.75 | 56.28 | 55.49 | 55.55 | 100.00 | 46.49 | 50.23 | 51.21 | 50.04 | 49.85 | 49.94 | 50.32 | 49.73 | 48.83 | 56.25 | 54.97 | 54.97 | 47.70 | 48.55 |
| 11: Ag | 44.43 | 43.17 | 43.17 | 42.62 | 42.79 | 42.25 | 45.54 | 45.17 | 45.35 | 46.49 | 100.00 | 44.02 | 47.46 | 46.24 | 45.92 | 46.21 | 46.71 | 47.08 | 46.60 | 51.34 | 51.86 | 51.86 | 54.50 | 53.96 |
| 12: Bfpks1 | 53.24 | 50.54 | 50.54 | 49.58 | 49.46 | 44.70 | 48.23 | 47.63 | 49.10 | 50.23 | 44.02 | 100.00 | 47.14 | 46.51 | 46.74 | 46.36 | 46.25 | 47.07 | 45.71 | 52.84 | 51.75 | 51.75 | 44.94 | 45.35 |
| 13: Cc | 47.52 | 45.42 | 45.42 | 44.78 | 44.87 | 44.74 | 48.50 | 48.26 | 48.95 | 51.21 | 47.46 | 47.14 | 100.00 | 62.78 | 63.67 | 63.35 | 63.23 | 57.36 | 56.43 | 55.19 | 54.97 | 54.97 | 48.68 | 48.91 |
| 14: TSrbtJ | 47.61 | 45.10 | 45.10 | 44.51 | 44.55 | 44.15 | 46.98 | 47.09 | 49.51 | 50.04 | 46.24 | 46.51 | 62.78 | 100.00 | 85.07 | 70.49 | 72.89 | 56.27 | 56.23 | 54.74 | 54.36 | 54.36 | 48.33 | 48.72 |
| 15: Tfpks | 47.39 | 44.98 | 44.98 | 44.65 | 44.65 | 44.45 | 47.11 | 46.90 | 49.88 | 49.85 | 45.92 | 46.74 | 63.67 | 85.07 | 100.00 | 71.10 | 73.74 | 56.41 | 56.44 | 55.35 | 54.80 | 54.80 | 48.55 | 48.65 |
| 16: Cad2 | 47.15 | 45.06 | 45.06 | 44.56 | 44.51 | 44.15 | 47.83 | 46.88 | 49.33 | 49.94 | 46.21 | 46.36 | 63.35 | 70.49 | 71.10 | 100.00 | 74.97 | 56.81 | 56.46 | 55.60 | 54.36 | 54.36 | 48.89 | 49.01 |
| 17: OmPKS1 | 47.94 | 45.03 | 45.03 | 44.90 | 44.95 | 44.54 | 47.73 | 47.14 | 49.25 | 50.32 | 46.71 | 46.25 | 63.23 | 72.89 | 73.74 | 74.97 | 100.00 | 56.66 | 57.19 | 55.53 | 55.14 | 55.14 | 49.08 | 48.62 |
| 18: phiA | 48.88 | 46.48 | 46.48 | 45.84 | 45.84 | 44.78 | 47.92 | 47.85 | 48.73 | 49.73 | 47.08 | 47.07 | 57.36 | 56.27 | 56.41 | 56.81 | 56.66 | 100.00 | 64.05 | 54.91 | 54.32 | 54.32 | 49.30 | 50.14 |
| 19: Ts1 | 47.76 | 45.48 | 45.48 | 45.10 | 45.18 | 44.59 | 47.24 | 47.13 | 48.77 | 48.83 | 46.60 | 45.71 | 56.43 | 56.23 | 56.44 | 56.46 | 57.19 | 64.05 | 100.00 | 54.54 | 53.89 | 53.89 | 48.74 | 48.70 |
| 20: ScyPKS | 53.87 | 49.79 | 49.79 | 49.21 | 49.29 | 49.37 | 54.97 | 55.31 | 55.62 | 56.25 | 51.34 | 52.84 | 55.19 | 54.74 | 55.35 | 55.60 | 55.53 | 54.91 | 54.54 | 100.00 | 81.18 | 81.18 | 55.86 | 56.43 |
| 21: ZopPKS | 52.94 | 49.10 | 49.10 | 48.27 | 48.39 | 49.22 | 54.44 | 53.53 | 55.25 | 54.97 | 51.86 | 51.75 | 54.97 | 54.36 | 54.80 | 54.36 | 55.14 | 54.32 | 53.89 | 81.18 | 100.00 | 100.00 | 55.78 | 55.32 |
| 22: ZopA | 52.94 | 49.10 | 49.10 | 48.27 | 48.39 | 49.22 | 54.44 | 53.53 | 55.25 | 54.97 | 51.86 | 51.75 | 54.97 | 54.36 | 54.80 | 54.36 | 55.14 | 54.32 | 53.89 | 81.18 | 100.00 | 100.00 | 55.78 | 55.32 |
| 23: TbPKS1 | 47.38 | 44.41 | 44.41 | 44.24 | 44.24 | 43.31 | 46.94 | 46.99 | 47.45 | 47.70 | 54.50 | 44.94 | 48.68 | 48.33 | 48.55 | 48.89 | 49.08 | 49.30 | 48.74 | 55.86 | 55.78 | 55.78 | 100.00 | 60.88 |
| 24: Ia | 47.15 | 44.48 | 44.48 | 44.23 | 44.31 | 44.47 | 47.58 | 47.53 | 46.77 | 48.55 | 53.96 | 45.35 | 48.91 | 48.72 | 48.65 | 49.01 | 48.62 | 50.14 | 48.70 | 56.43 | 55.32 | 55.32 | 60.88 | 100.00 |

#### Hydrolase

Table S5: Percentage identity matrix of hydrolases generated using MUSCLE,[1] including the maleidride hydrolases, identified homologues from the asperlin and communesin pathways, and a homologue present within the *A. oryzae* genome.

| | 1: | 2: | 3: | 4: | 5: | 6: | 7: | 8: | 9: | 10: | 11: | 12: | 13: | 14: | 15: | 16: | 17: | 18: | 19: | 20: | 21: | 22: | 23: | 24: | 25: |
|---|---|---|---|---|---|---|---|---|---|---|---|---|---|---|---|---|---|---|---|---|---|---|---|---|---|
| 1: A0A0A2JY30.1_CnsH | 100.00 | 30.13 | 21.26 | 24.28 | 19.78 | 24.71 | 21.39 | 21.39 | 21.39 | 22.54 | 22.54 | 26.01 | 26.59 | 25.43 | 27.75 | 26.01 | 26.01 | 23.70 | 26.01 | 28.90 | 28.90 | 23.70 | 26.01 | 20.11 | 24.28 |
| 2: C8VJR6.1_AlnB | 30.13 | 100.00 | 28.77 | 34.95 | 26.48 | 31.75 | 33.33 | 31.58 | 31.58 | 31.58 | 31.58 | 33.01 | 33.97 | 34.93 | 38.28 | 38.76 | 38.76 | 30.65 | 35.55 | 32.23 | 33.81 | 34.93 | 35.71 | 25.59 | 27.14 |
| 3: RDW56968.1-Cc | 21.26 | 28.77 | 100.00 | 41.36 | 40.36 | 42.86 | 43.24 | 38.91 | 38.91 | 38.46 | 38.46 | 45.25 | 44.80 | 46.15 | 44.34 | 48.87 | 48.87 | 42.64 | 42.79 | 41.26 | 43.44 | 42.08 | 43.89 | 46.19 | 42.08 |
| 4: TbR3 | 24.28 | 34.95 | 41.36 | 100.00 | 42.20 | 50.68 | 46.33 | 49.08 | 49.08 | 47.71 | 47.71 | 51.83 | 46.33 | 47.25 | 50.92 | 51.38 | 51.38 | 44.39 | 46.54 | 49.32 | 49.31 | 48.17 | 48.39 | 41.82 | 39.63 |
| 5: OJJ86354.1_Ag | 19.78 | 26.48 | 40.36 | 42.20 | 100.00 | 43.95 | 50.23 | 47.51 | 47.51 | 47.51 | 47.51 | 47.06 | 48.87 | 50.23 | 52.49 | 50.23 | 50.23 | 40.61 | 47.06 | 44.80 | 46.15 | 44.59 | 46.82 | 39.73 | 39.64 |
| 6: CAF9941818.1b-Ia | 24.71 | 31.75 | 42.86 | 50.68 | 43.95 | 100.00 | 51.13 | 49.32 | 49.32 | 50.68 | 51.13 | 52.04 | 48.42 | 50.23 | 48.42 | 51.58 | 51.58 | 43.88 | 50.68 | 47.51 | 50.68 | 50.68 | 47.73 | 43.24 | 41.36 |
| 7: EpiR11_Wa_ | 21.39 | 33.33 | 43.24 | 46.33 | 50.23 | 51.13 | 100.00 | 70.14 | 70.14 | 70.14 | 71.04 | 57.01 | 54.75 | 53.39 | 57.01 | 56.11 | 56.11 | 47.96 | 48.87 | 45.25 | 49.09 | 52.73 | 51.13 | 43.24 | 41.82 |
| 8: XP_014073929.1_Bm | 21.39 | 31.58 | 38.91 | 49.08 | 47.51 | 49.32 | 70.14 | 100.00 | 100.00 | 90.95 | 91.86 | 56.11 | 52.94 | 55.66 | 57.01 | 54.30 | 54.30 | 47.18 | 48.64 | 46.82 | 50.00 | 50.91 | 48.18 | 40.72 | 42.01 |
| 9: EMD85581.1_BmC5 | 21.39 | 31.58 | 38.91 | 49.08 | 47.51 | 49.32 | 70.14 | 100.00 | 100.00 | 90.95 | 91.86 | 56.11 | 52.94 | 55.66 | 57.01 | 54.30 | 54.30 | 47.18 | 48.64 | 46.82 | 50.00 | 50.91 | 48.18 | 40.72 | 42.01 |
| 10: XP_014553701.1_Bv | 22.54 | 31.58 | 38.46 | 47.71 | 47.51 | 50.68 | 70.14 | 90.95 | 90.95 | 100.00 | 99.10 | 54.30 | 51.13 | 52.04 | 53.39 | 52.04 | 52.04 | 46.67 | 49.09 | 45.45 | 49.09 | 50.45 | 47.27 | 39.37 | 40.18 |
| 11: XP_007715667.1_Bz | 22.54 | 31.58 | 38.46 | 47.71 | 47.51 | 51.13 | 71.04 | 91.86 | 91.86 | 99.10 | 100.00 | 54.30 | 52.04 | 52.94 | 54.30 | 52.49 | 52.49 | 47.18 | 49.09 | 45.45 | 49.09 | 50.91 | 47.73 | 39.37 | 40.64 |
| 12: ANF07287.1_BfL1_ | 26.01 | 33.01 | 45.25 | 51.83 | 47.06 | 52.04 | 57.01 | 56.11 | 56.11 | 54.30 | 54.30 | 100.00 | 53.85 | 55.20 | 57.92 | 58.82 | 58.82 | 48.72 | 50.91 | 50.45 | 54.09 | 58.64 | 55.91 | 44.34 | 42.92 |
| 13: XP002482824_TsRbtS | 26.59 | 33.97 | 44.80 | 46.33 | 48.87 | 48.42 | 54.75 | 52.94 | 52.94 | 51.13 | 52.04 | 53.85 | 100.00 | 82.35 | 65.61 | 64.25 | 64.25 | 45.13 | 50.45 | 46.82 | 49.55 | 53.18 | 53.18 | 45.70 | 41.10 |
| 14: tfL9_Tf_ | 25.43 | 34.93 | 46.15 | 47.25 | 50.23 | 50.23 | 53.39 | 55.66 | 55.66 | 52.04 | 52.94 | 55.20 | 82.35 | 100.00 | 68.78 | 69.23 | 69.23 | 48.21 | 49.55 | 49.09 | 51.82 | 53.64 | 50.00 | 45.70 | 42.92 |
| 15: QTE76001.1_ScyR1 | 27.75 | 38.28 | 44.34 | 50.92 | 52.49 | 48.42 | 57.01 | 57.01 | 57.01 | 53.39 | 54.30 | 57.92 | 65.61 | 68.78 | 100.00 | 72.85 | 72.85 | 50.26 | 55.45 | 51.36 | 55.45 | 57.27 | 57.27 | 45.25 | 44.75 |
| 16: QTE75993.1_ZopR1 | 26.01 | 38.76 | 48.87 | 51.38 | 50.23 | 51.58 | 56.11 | 54.30 | 54.30 | 52.04 | 52.49 | 58.82 | 64.25 | 69.23 | 72.85 | 100.00 | 100.00 | 52.82 | 55.91 | 56.36 | 60.45 | 63.18 | 57.27 | 46.61 | 42.01 |
| 17: BBU42027.1_zopM | 26.01 | 38.76 | 48.87 | 51.38 | 50.23 | 51.58 | 56.11 | 54.30 | 54.30 | 52.04 | 52.49 | 58.82 | 64.25 | 69.23 | 72.85 | 100.00 | 100.00 | 52.82 | 55.91 | 56.36 | 60.45 | 63.18 | 57.27 | 46.61 | 42.01 |
| 18: OO004461.1-**A.oryzae** | 23.70 | 30.65 | 42.64 | 44.39 | 40.61 | 43.88 | 47.96 | 47.18 | 47.18 | 46.67 | 47.18 | 48.72 | 45.13 | 48.21 | 50.26 | 52.82 | 52.82 | 100.00 | 53.57 | 56.63 | 55.38 | 51.01 | 57.65 | 40.91 | 36.73 |
| 19: Czm | 26.01 | 35.55 | 42.79 | 46.54 | 47.06 | 50.68 | 48.87 | 48.64 | 48.64 | 49.09 | 49.09 | 50.91 | 50.45 | 49.55 | 55.45 | 55.91 | 55.91 | 53.57 | 100.00 | 64.86 | 65.16 | 57.27 | 60.63 | 42.99 | 41.10 |
| 20: Cad | 28.90 | 32.23 | 41.26 | 49.32 | 44.80 | 47.51 | 45.25 | 46.82 | 46.82 | 45.45 | 45.45 | 50.45 | 46.82 | 49.09 | 51.36 | 56.36 | 56.36 | 56.63 | 64.86 | 100.00 | 79.19 | 59.09 | 60.18 | 40.72 | 37.44 |
| 21: KIN05363.1-Om | 28.90 | 33.81 | 43.44 | 49.31 | 46.15 | 50.68 | 49.09 | 50.00 | 50.00 | 49.09 | 49.09 | 54.09 | 49.55 | 51.82 | 55.45 | 60.45 | 60.45 | 55.38 | 65.16 | 79.19 | 100.00 | 60.91 | 62.73 | 43.64 | 40.37 |
| 22: ASK38716.1_PvL1 | 23.70 | 34.93 | 42.08 | 48.17 | 44.59 | 50.68 | 52.73 | 50.91 | 50.91 | 50.45 | 50.91 | 58.64 | 53.18 | 53.64 | 57.27 | 63.18 | 63.18 | 51.01 | 57.27 | 59.09 | 60.91 | 100.00 | 65.00 | 41.89 | 39.09 |
| 23: XP_020117166_Ta | 26.01 | 35.71 | 43.89 | 48.39 | 46.82 | 47.73 | 51.13 | 48.18 | 48.18 | 47.27 | 47.73 | 55.91 | 53.18 | 50.00 | 57.27 | 57.27 | 57.27 | 57.65 | 60.63 | 60.18 | 62.73 | 65.00 | 100.00 | 42.08 | 40.18 |
| 24: XM_002485320_ts1R10 | 20.11 | 25.59 | 46.19 | 41.82 | 39.73 | 43.24 | 43.24 | 40.72 | 40.72 | 39.37 | 39.37 | 44.34 | 45.70 | 45.70 | 45.25 | 46.61 | 46.61 | 40.91 | 42.99 | 40.72 | 43.64 | 41.89 | 42.08 | 100.00 | 48.21 |
| 25: BBG28510.1mod_PhiM | 24.28 | 27.14 | 42.08 | 39.63 | 39.64 | 41.36 | 41.82 | 42.01 | 42.01 | 40.18 | 40.64 | 42.92 | 41.10 | 42.92 | 44.75 | 42.01 | 42.01 | 36.73 | 41.10 | 37.44 | 40.37 | 39.09 | 40.18 | 48.21 | 100.00 |

#### ACS

Table S6: Percentage identity matrix of maleidride alkylcitrate synthases generated using MUSCLE.[1]

| | 1: | 2: | 3: | 4: | 5: | 6: | 7: | 8: | 9: | 10: | 11: | 12: | 13: | 14: | 15: | 16: | 17: | 18: | 19: | 20: | 21: | 22: |
|---|---|---|---|---|---|---|---|---|---|---|---|---|---|---|---|---|---|---|---|---|---|---|
| 1: BBU42028.1_ZC | 100.00 | 100.00 | 61.40 | 52.82 | 56.36 | 45.50 | 49.08 | 47.11 | 54.27 | 48.73 | 50.35 | 51.04 | 53.12 | 51.73 | 50.70 | 48.27 | 54.50 | 50.00 | 49.52 | 49.52 | 49.65 | 49.88 |
| 2: QTE75995.1_DC | 100.00 | 100.00 | 61.40 | 52.82 | 56.36 | 45.50 | 49.08 | 47.11 | 54.27 | 48.73 | 50.35 | 51.04 | 53.12 | 51.73 | 50.70 | 48.27 | 54.50 | 50.00 | 49.52 | 49.52 | 49.65 | 49.88 |
| 3: Cc\|RDW56963.1 | 61.40 | 61.40 | 100.00 | 52.94 | 56.41 | 46.23 | 44.94 | 44.16 | 51.95 | 46.23 | 46.49 | 47.79 | 46.75 | 46.75 | 48.31 | 47.27 | 48.31 | 48.56 | 44.42 | 44.42 | 43.90 | 44.42 |
| 4: EED15409.1_TS1 | 52.82 | 52.82 | 52.94 | 100.00 | 58.22 | 44.47 | 45.55 | 43.53 | 50.35 | 41.92 | 44.96 | 46.84 | 46.60 | 45.43 | 47.07 | 46.26 | 50.12 | 46.68 | 44.06 | 44.06 | 43.29 | 44.00 |
| 5: BBG28507.1_phiJ | 56.36 | 56.36 | 56.41 | 58.22 | 100.00 | 45.33 | 46.23 | 44.87 | 52.39 | 44.90 | 46.26 | 47.17 | 47.62 | 47.85 | 49.43 | 49.32 | 51.03 | 48.93 | 47.85 | 47.85 | 46.47 | 46.92 |
| 6: OJJ86352.1_AG | 45.50 | 45.50 | 46.23 | 44.47 | 45.33 | 100.00 | 60.47 | 62.81 | 56.46 | 50.11 | 49.66 | 51.25 | 53.29 | 52.83 | 54.34 | 52.83 | 52.61 | 53.19 | 50.95 | 50.95 | 51.70 | 51.93 |
| 7: Ia\|CAF9941815.1\| | 49.08 | 49.08 | 44.94 | 45.55 | 46.23 | 60.47 | 100.00 | 71.06 | 59.43 | 48.58 | 50.39 | 51.42 | 54.78 | 52.20 | 55.30 | 54.78 | 52.45 | 53.51 | 50.90 | 50.90 | 50.39 | 50.90 |
| 8: Tb\|TbR4 | 47.11 | 47.11 | 44.16 | 43.53 | 44.87 | 62.81 | 71.06 | 100.00 | 59.86 | 49.66 | 50.57 | 50.11 | 53.74 | 52.61 | 56.16 | 54.65 | 54.42 | 53.43 | 51.19 | 51.19 | 51.02 | 51.70 |
| 9: QTE76003.1_SA | 54.27 | 54.27 | 51.95 | 50.35 | 52.39 | 56.46 | 59.43 | 59.86 | 100.00 | 59.41 | 59.86 | 60.77 | 65.53 | 65.99 | 64.16 | 59.41 | 62.36 | 61.23 | 57.62 | 57.62 | 59.64 | 60.09 |
| 10:Czm\|KAF2215727.1\| | 48.73 | 48.73 | 46.23 | 41.92 | 44.90 | 50.11 | 48.58 | 49.66 | 59.41 | 100.00 | 62.75 | 61.63 | 66.82 | 64.79 | 62.50 | 61.85 | 56.46 | 57.45 | 57.38 | 57.38 | 58.28 | 57.82 |
| 11:EED18839.1_TS2 | 50.35 | 50.35 | 46.49 | 44.96 | 46.26 | 49.66 | 50.39 | 50.57 | 59.86 | 62.75 | 100.00 | 89.39 | 77.43 | 74.04 | 67.95 | 65.69 | 55.33 | 58.63 | 58.10 | 58.10 | 58.73 | 58.96 |
| 12:TF\|TfL2\| | 51.04 | 51.04 | 47.79 | 46.84 | 47.17 | 51.25 | 51.42 | 50.11 | 60.77 | 61.63 | 89.39 | 100.00 | 76.75 | 75.17 | 70.23 | 67.04 | 55.56 | 60.28 | 57.62 | 57.62 | 58.05 | 58.05 |
| 13:Om\|KIN05361.1\| | 53.12 | 53.12 | 46.75 | 46.60 | 47.62 | 53.29 | 54.78 | 53.74 | 65.53 | 66.82 | 77.43 | 76.75 | 100.00 | 81.94 | 70.00 | 69.30 | 61.45 | 61.94 | 60.71 | 60.71 | 61.22 | 60.32 |
| 14:Cad\|PVH77201.1\| | 51.73 | 51.73 | 46.75 | 45.43 | 47.85 | 52.83 | 52.20 | 52.61 | 65.99 | 64.79 | 74.04 | 75.17 | 81.94 | 100.00 | 74.55 | 71.78 | 59.64 | 63.12 | 61.67 | 61.67 | 61.45 | 60.54 |
| 15:OKL57051.1_TA | 50.70 | 50.70 | 48.31 | 47.07 | 49.43 | 54.34 | 55.30 | 56.16 | 64.16 | 62.50 | 67.95 | 70.23 | 70.00 | 74.55 | 100.00 | 75.74 | 61.19 | 64.52 | 60.19 | 60.19 | 60.05 | 60.73 |
| 16:ASK38711.1_PV | 48.27 | 48.27 | 47.27 | 46.26 | 49.32 | 52.83 | 54.78 | 54.65 | 59.41 | 61.85 | 65.69 | 67.04 | 69.30 | 71.78 | 75.74 | 100.00 | 58.28 | 61.70 | 59.76 | 59.76 | 59.86 | 60.54 |
| 17:ANF07286.1_BF | 54.50 | 54.50 | 48.31 | 50.12 | 51.03 | 52.61 | 52.45 | 54.42 | 62.36 | 56.46 | 55.33 | 55.56 | 61.45 | 59.64 | 61.19 | 58.28 | 100.00 | 66.67 | 63.57 | 63.57 | 64.40 | 63.95 |
| 18:epiR8_WA | 50.00 | 50.00 | 48.56 | 46.68 | 48.93 | 53.19 | 53.51 | 53.43 | 61.23 | 57.45 | 58.63 | 60.28 | 61.94 | 63.12 | 64.52 | 61.70 | 66.67 | 100.00 | 76.08 | 76.08 | 74.70 | 73.76 |
| 19:XP_014073930.1_BM | 49.52 | 49.52 | 44.42 | 44.06 | 47.85 | 50.95 | 50.90 | 51.19 | 57.62 | 57.38 | 58.10 | 57.62 | 60.71 | 61.67 | 60.19 | 59.76 | 63.57 | 76.08 | 100.00 | 100.00 | 92.14 | 91.43 |
| 20:EMD85580.1_BMC5 | 49.52 | 49.52 | 44.42 | 44.06 | 47.85 | 50.95 | 50.90 | 51.19 | 57.62 | 57.38 | 58.10 | 57.62 | 60.71 | 61.67 | 60.19 | 59.76 | 63.57 | 76.08 | 100.00 | 100.00 | 92.14 | 91.43 |
| 21:XP_007715666.1_BZ | 49.65 | 49.65 | 43.90 | 43.29 | 46.47 | 51.70 | 50.39 | 51.02 | 59.64 | 58.28 | 58.73 | 58.05 | 61.22 | 61.45 | 60.05 | 59.86 | 64.40 | 74.70 | 92.14 | 92.14 | 100.00 | 98.41 |
| 22:XP_014553700.1_BV | 49.88 | 49.88 | 44.42 | 44.00 | 46.92 | 51.93 | 50.90 | 51.70 | 60.09 | 57.82 | 58.96 | 58.05 | 60.32 | 60.54 | 60.73 | 60.54 | 63.95 | 73.76 | 91.43 | 91.43 | 98.41 | 100.00 |

### ACDH

Table S7: Percentage identity matrix of maleidride alkylcitrate dehydratases generated using MUSCLE.[1]

| | 1: | 2: | 3: | 4: | 5: | 6: | 7: | 8: | 9: | 10: | 11: | 12: | 13: | 14: | 15: | 16: | 17: | 18: | 19: | 20: | 21: | 22: |
|---|---|---|---|---|---|---|---|---|---|---|---|---|---|---|---|---|---|---|---|---|---|---|
| 1: Ts1_EED15409.1 | 100.00 | 56.70 | 59.05 | 51.95 | 52.57 | 50.10 | 50.31 | 50.10 | 50.10 | 51.95 | 52.25 | 52.25 | 53.20 | 52.89 | 50.92 | 51.33 | 51.95 | 54.62 | 53.39 | 51.23 | 50.92 | 52.46 |
| 2: Cc_RDW56964.1 | 56.70 | 100.00 | 61.36 | 57.88 | 59.92 | 55.79 | 55.99 | 57.02 | 57.02 | 58.88 | 60.33 | 60.33 | 60.50 | 59.75 | 54.75 | 58.47 | 59.50 | 58.47 | 59.09 | 54.75 | 57.53 | 57.44 |
| 3: BBG28506.1_PhiI | 59.05 | 61.36 | 100.00 | 55.79 | 57.82 | 54.53 | 54.73 | 56.17 | 56.17 | 57.82 | 58.23 | 58.23 | 57.97 | 57.64 | 53.18 | 55.14 | 57.61 | 57.41 | 58.64 | 52.78 | 55.14 | 53.91 |
| 4: Ta_OKL57050.1 | 51.95 | 57.88 | 55.79 | 100.00 | 63.17 | 58.44 | 58.44 | 59.26 | 59.26 | 61.52 | 60.57 | 60.57 | 61.98 | 61.86 | 56.79 | 58.93 | 59.96 | 59.34 | 59.75 | 58.32 | 60.66 | 61.15 |
| 5: Wa_epiR8 | 52.57 | 59.92 | 57.82 | 63.17 | 100.00 | 72.54 | 72.75 | 73.16 | 73.16 | 67.62 | 64.75 | 64.75 | 65.98 | 65.77 | 59.14 | 64.55 | 63.93 | 63.52 | 63.73 | 57.08 | 58.40 | 59.63 |
| 6: Bv_XP_014553699.1 | 50.10 | 55.79 | 54.53 | 58.44 | 72.54 | 100.00 | 99.39 | 89.96 | 89.96 | 61.48 | 60.04 | 60.04 | 61.24 | 61.86 | 54.83 | 57.99 | 57.17 | 58.61 | 57.38 | 50.92 | 54.51 | 56.97 |
| 7: Bz_EUC30034.1 | 50.31 | 55.99 | 54.73 | 58.44 | 72.75 | 99.39 | 100.00 | 89.96 | 89.96 | 61.68 | 60.25 | 60.25 | 61.44 | 62.06 | 55.03 | 57.99 | 57.38 | 58.81 | 57.58 | 51.13 | 54.71 | 56.97 |
| 8: Bm_C5_EMD85579.1 | 50.10 | 57.02 | 56.17 | 59.26 | 73.16 | 89.96 | 89.96 | 100.00 | 100.00 | 61.07 | 60.86 | 60.86 | 62.06 | 62.68 | 56.06 | 58.81 | 59.02 | 59.84 | 58.40 | 51.75 | 56.35 | 57.99 |
| 9: Bm_48331_ENH99969.1 | 50.10 | 57.02 | 56.17 | 59.26 | 73.16 | 89.96 | 89.96 | 100.00 | 100.00 | 61.07 | 60.86 | 60.86 | 62.06 | 62.68 | 56.06 | 58.81 | 59.02 | 59.84 | 58.40 | 51.75 | 56.35 | 57.99 |
| 10: Bf_ANF07285.1 | 51.95 | 58.88 | 57.82 | 61.52 | 67.62 | 61.48 | 61.68 | 61.07 | 61.07 | 100.00 | 64.14 | 64.14 | 67.63 | 63.92 | 58.93 | 61.48 | 63.11 | 64.14 | 63.73 | 56.06 | 56.15 | 59.02 |
| 11: Dc_QTE75987.1 | 52.25 | 60.33 | 58.23 | 60.57 | 64.75 | 60.04 | 60.25 | 60.86 | 60.86 | 64.14 | 100.00 | 100.00 | 81.39 | 64.95 | 57.99 | 62.17 | 64.01 | 65.44 | 64.83 | 55.44 | 57.06 | 58.90 |
| 12: Zc_BBU42021.1 | 52.25 | 60.33 | 58.23 | 60.57 | 64.75 | 60.04 | 60.25 | 60.86 | 60.86 | 64.14 | 100.00 | 100.00 | 81.39 | 64.95 | 57.99 | 62.17 | 64.01 | 65.44 | 64.83 | 55.44 | 57.06 | 58.90 |
| 13: Sa_QTE76007.1 | 53.20 | 60.50 | 57.97 | 61.98 | 65.98 | 61.24 | 61.44 | 62.06 | 62.06 | 67.63 | 81.39 | 81.39 | 100.00 | 68.26 | 60.21 | 64.61 | 66.26 | 64.61 | 65.43 | 54.75 | 58.23 | 60.08 |
| 14: Pv_ASK38715.1 | 52.89 | 59.75 | 57.64 | 61.86 | 65.77 | 61.86 | 62.06 | 62.68 | 62.68 | 63.92 | 64.95 | 64.95 | 68.26 | 100.00 | 65.70 | 67.42 | 69.90 | 68.45 | 68.45 | 57.32 | 58.97 | 61.24 |
| 15: Czm_KAF2215728.1 | 50.92 | 54.75 | 53.18 | 56.79 | 59.14 | 54.83 | 55.03 | 56.06 | 56.06 | 58.93 | 57.99 | 57.99 | 60.21 | 65.70 | 100.00 | 66.94 | 70.41 | 69.39 | 70.20 | 53.29 | 55.53 | 54.10 |
| 16: Ca_PVH77202.1 | 51.33 | 58.47 | 55.14 | 58.93 | 64.55 | 57.99 | 57.99 | 58.81 | 58.81 | 61.48 | 62.17 | 62.17 | 64.61 | 67.42 | 66.94 | 100.00 | 74.95 | 72.10 | 73.32 | 54.21 | 55.21 | 55.42 |
| 17: Om_KIN05360.1 | 51.95 | 59.50 | 57.61 | 59.96 | 63.93 | 57.17 | 57.38 | 59.02 | 59.02 | 63.11 | 64.01 | 64.01 | 66.26 | 69.90 | 70.41 | 74.95 | 100.00 | 79.84 | 81.67 | 54.83 | 56.85 | 56.24 |
| 18: Ts2_EED18840.1 | 54.62 | 58.47 | 57.41 | 59.34 | 63.52 | 58.61 | 58.81 | 59.84 | 59.84 | 64.14 | 65.44 | 65.44 | 64.61 | 68.45 | 69.39 | 72.10 | 79.84 | 100.00 | 87.58 | 55.24 | 58.90 | 57.67 |
| 19: TfL1 | 53.39 | 59.09 | 58.64 | 59.75 | 63.73 | 57.38 | 57.58 | 58.40 | 58.40 | 63.73 | 64.83 | 64.83 | 65.43 | 68.45 | 70.20 | 73.32 | 81.67 | 87.58 | 100.00 | 56.26 | 57.87 | 58.08 |
| 20: Ag_OJJ86350.1 | 51.23 | 54.75 | 52.78 | 58.32 | 57.08 | 50.92 | 51.13 | 51.75 | 51.75 | 56.06 | 55.44 | 55.44 | 54.75 | 57.32 | 53.29 | 54.21 | 54.83 | 55.24 | 56.26 | 100.00 | 61.15 | 62.30 |
| 21: TbR1 | 50.92 | 57.53 | 55.14 | 60.66 | 58.40 | 54.51 | 54.71 | 56.35 | 56.35 | 56.15 | 57.06 | 57.06 | 58.23 | 58.97 | 55.53 | 55.21 | 56.85 | 58.90 | 57.87 | 61.15 | 100.00 | 66.94 |
| 22: Ia_CAF9941813.1 | 52.46 | 57.44 | 53.91 | 61.15 | 59.63 | 56.97 | 56.97 | 57.99 | 57.99 | 59.02 | 58.90 | 58.90 | 60.08 | 61.24 | 54.10 | 55.42 | 56.24 | 57.67 | 58.08 | 62.30 | 66.94 | 100.00 |

### MDC

Table S8: Percentage identity matrix of maleidride dimerising cyclases generated using MUSCLE.[1]

| | 1: | 2: | 3: | 4 | 5: | 6: | 7: | 8: | 9: | 10: | 11: | 12: | 13: | 14: | 15: | 16: | 17: | 18: | 19: | 20: | 21: | 22: | 23: | 24: | 25: | 26: | 27: | 28: | 29: |
|---|---|---|---|---|---|---|---|---|---|---|---|---|---|---|---|---|---|---|---|---|---|---|---|---|---|---|---|---|---|
| 1: OJJ86357.1-Ag | 100.00 | 53.48 | 53.98 | 29.73 | 28.73 | 31.11 | 32.43 | 34.43 | 34.78 | 33.70 | 31.72 | 32.61 | 33.88 | 33.88 | 32.79 | 33.33 | 31.22 | 32.80 | 34.97 | 34.97 | 35.52 | 35.52 | 31.18 | 30.98 | 32.97 | 31.89 | 31.89 | 32.80 | 29.79 |
| 2: TbL7 | 53.48 | 100.00 | 56.76 | 24.87 | 24.34 | 25.93 | 28.93 | 29.69 | 31.61 | 25.00 | 29.85 | 31.05 | 30.69 | 30.69 | 29.10 | 29.63 | 29.50 | 34.87 | 34.41 | 34.41 | 34.41 | 34.41 | 28.35 | 26.70 | 30.05 | 28.50 | 28.50 | 31.47 | 29.15 |
| 3: CAF9941823.1mod-Ia | 53.98 | 56.76 | 100.00 | 26.40 | 25.99 | 29.38 | 28.49 | 31.64 | 33.90 | 23.89 | 30.05 | 32.76 | 31.03 | 31.03 | 30.46 | 30.46 | 30.77 | 33.15 | 34.29 | 34.29 | 34.29 | 34.29 | 30.86 | 31.21 | 32.18 | 31.61 | 31.61 | 29.78 | 28.89 |
| 4: EED18833.1-Ts1 | 29.73 | 24.87 | 26.40 | 100.00 | 61.75 | 60.19 | 37.21 | 33.02 | 34.91 | 32.08 | 39.81 | 37.56 | 37.25 | 37.25 | 36.27 | 37.25 | 37.73 | 34.42 | 33.66 | 33.66 | 33.66 | 33.66 | 35.98 | 37.26 | 38.50 | 36.15 | 36.15 | 34.70 | 33.95 |
| 5: RDW56973.1-Cc | 28.73 | 24.34 | 25.99 | 61.75 | 100.00 | 59.64 | 27.49 | 30.37 | 33.80 | 31.13 | 36.27 | 36.32 | 36.32 | 36.32 | 34.33 | 35.32 | 35.00 | 33.18 | 30.30 | 30.30 | 30.30 | 30.30 | 34.58 | 36.02 | 38.03 | 35.68 | 35.68 | 34.72 | 33.64 |
| 6: BBG28500.1-PhiC | 31.11 | 25.93 | 29.38 | 60.19 | 59.64 | 100.00 | 31.10 | 33.80 | 36.45 | 31.28 | 38.73 | 34.50 | 36.00 | 36.00 | 35.00 | 36.00 | 34.40 | 33.02 | 31.98 | 31.98 | 31.98 | 31.98 | 34.42 | 35.85 | 34.58 | 33.18 | 33.18 | 35.05 | 30.19 |
| 7: KAF2215733.1-Czm | 32.43 | 28.93 | 28.49 | 37.21 | 27.49 | 31.10 | 100.00 | 47.98 | 42.86 | 30.14 | 35.32 | 36.67 | 36.84 | 36.84 | 35.89 | 35.89 | 33.63 | 29.60 | 32.38 | 32.38 | 32.38 | 32.38 | 33.18 | 33.94 | 33.93 | 33.04 | 33.04 | 30.43 | 32.30 |
| 8: KIN05369.1-Om | 34.43 | 29.69 | 31.64 | 33.02 | 30.37 | 33.80 | 47.98 | 100.00 | 70.61 | 34.72 | 37.91 | 40.58 | 40.10 | 40.10 | 39.61 | 40.10 | 34.53 | 32.57 | 36.95 | 36.95 | 36.95 | 36.95 | 37.33 | 38.14 | 39.91 | 38.07 | 38.07 | 31.70 | 35.59 |
| 9: PVH77195.1-Ca | 34.78 | 31.61 | 33.90 | 34.91 | 33.80 | 36.45 | 42.86 | 70.61 | 100.00 | 35.62 | 39.91 | 41.83 | 41.35 | 41.35 | 39.90 | 40.38 | 34.07 | 33.94 | 37.13 | 37.13 | 37.13 | 37.13 | 39.55 | 40.55 | 41.63 | 39.82 | 39.82 | 33.92 | 36.89 |
| 10: KAF2215732.1-Czm | 33.70 | 25.00 | 23.89 | 32.08 | 31.13 | 31.28 | 30.14 | 34.72 | 35.62 | 100.00 | 35.75 | 37.81 | 37.50 | 37.50 | 35.50 | 35.50 | 32.17 | 30.32 | 34.63 | 34.63 | 34.63 | 34.63 | 31.65 | 33.64 | 33.49 | 32.11 | 32.11 | 30.54 | 30.60 |
| 11: ANF07278.1-BfL10 | 31.72 | 29.85 | 30.05 | 39.81 | 36.27 | 38.73 | 35.32 | 37.91 | 39.91 | 35.75 | 100.00 | 61.46 | 62.75 | 62.75 | 61.76 | 61.76 | 49.10 | 46.08 | 48.28 | 48.28 | 47.78 | 47.78 | 43.87 | 43.75 | 46.23 | 47.17 | 47.17 | 43.05 | 44.50 |
| 12: EpiR6-Wa | 32.61 | 31.05 | 32.76 | 37.56 | 36.32 | 34.50 | 36.67 | 40.58 | 41.83 | 37.81 | 61.46 | 100.00 | 74.30 | 74.30 | 71.96 | 71.03 | 47.62 | 45.15 | 46.77 | 46.77 | 46.77 | 46.77 | 43.48 | 44.17 | 43.69 | 43.20 | 43.20 | 45.75 | 48.08 |
| 13: EMD85577.1-BmC5 | 33.88 | 30.69 | 31.03 | 37.25 | 36.32 | 36.00 | 36.84 | 40.10 | 41.35 | 37.50 | 62.75 | 74.30 | 100.00 | 100.00 | 90.65 | 89.72 | 49.28 | 46.83 | 51.00 | 51.00 | 51.00 | 51.00 | 46.12 | 46.83 | 46.34 | 45.37 | 45.37 | 46.92 | 49.28 |
| 14: XP_014073932.1mod-Bm48331 | 33.88 | 30.69 | 31.03 | 37.25 | 36.32 | 36.00 | 36.84 | 40.10 | 41.35 | 37.50 | 62.75 | 74.30 | 100.00 | 100.00 | 90.65 | 89.72 | 49.28 | 46.83 | 51.00 | 51.00 | 51.00 | 51.00 | 46.12 | 46.83 | 46.34 | 45.37 | 45.37 | 46.92 | 49.28 |
| 15: XP_014553697.1mod-Bv | 32.79 | 29.10 | 30.46 | 36.27 | 34.33 | 35.00 | 35.89 | 39.61 | 39.90 | 35.50 | 61.76 | 71.96 | 90.65 | 90.65 | 100.00 | 99.07 | 47.37 | 44.88 | 49.00 | 49.00 | 49.00 | 49.00 | 44.17 | 45.37 | 45.37 | 44.88 | 44.88 | 44.55 | 46.38 |
| 16: XP_007715663.1-Bz | 33.33 | 29.63 | 30.46 | 37.25 | 35.32 | 36.00 | 35.89 | 40.10 | 40.38 | 35.50 | 61.76 | 71.03 | 89.72 | 89.72 | 99.07 | 100.00 | 47.85 | 44.88 | 49.50 | 49.50 | 49.50 | 49.50 | 43.20 | 44.39 | 45.37 | 44.88 | 44.88 | 45.02 | 45.89 |
| 17: ANF07282.1-BfL6 | 31.22 | 29.50 | 30.77 | 37.73 | 35.00 | 34.40 | 33.63 | 34.53 | 34.07 | 32.17 | 49.10 | 47.62 | 49.28 | 49.28 | 47.37 | 47.85 | 100.00 | 47.84 | 55.14 | 55.14 | 54.67 | 54.67 | 42.22 | 42.99 | 45.33 | 42.67 | 42.67 | 45.38 | 44.21 |
| 18: EpiR1-Wa | 32.80 | 34.87 | 33.15 | 34.42 | 33.18 | 33.02 | 29.60 | 32.57 | 33.94 | 30.32 | 46.08 | 45.15 | 46.83 | 46.83 | 44.88 | 44.88 | 47.84 | 100.00 | 60.85 | 60.85 | 60.85 | 60.85 | 40.00 | 39.45 | 40.91 | 40.00 | 40.00 | 44.10 | 41.52 |
| 19: EMD85571.1-BmC5 | 34.97 | 34.41 | 34.29 | 33.66 | 30.30 | 31.98 | 32.38 | 36.95 | 37.13 | 34.63 | 48.28 | 46.77 | 51.00 | 51.00 | 49.00 | 49.50 | 55.14 | 60.85 | 100.00 | 100.00 | 98.13 | 98.13 | 42.44 | 41.95 | 43.63 | 41.67 | 41.67 | 45.54 | 43.00 |
| 20: XP_014073935.1-Bm48331 | 34.97 | 34.41 | 34.29 | 33.66 | 30.30 | 31.98 | 32.38 | 36.95 | 37.13 | 34.63 | 48.28 | 46.77 | 51.00 | 51.00 | 49.00 | 49.50 | 55.14 | 60.85 | 100.00 | 100.00 | 98.13 | 98.13 | 42.44 | 41.95 | 43.63 | 41.67 | 41.67 | 45.54 | 43.00 |
| 21: XP_014553690.1-Bv | 35.52 | 34.41 | 34.29 | 33.66 | 30.30 | 31.98 | 32.38 | 36.95 | 37.13 | 34.63 | 47.78 | 46.77 | 51.00 | 51.00 | 49.00 | 49.50 | 54.67 | 60.85 | 98.13 | 98.13 | 100.00 | 100.00 | 42.93 | 42.44 | 44.12 | 42.16 | 42.16 | 44.60 | 42.51 |
| 22: XP_007715659.1-Bz | 35.52 | 34.41 | 34.29 | 33.66 | 30.30 | 31.98 | 32.38 | 36.95 | 37.13 | 34.63 | 47.78 | 46.77 | 51.00 | 51.00 | 49.00 | 49.50 | 54.67 | 60.85 | 98.13 | 98.13 | 100.00 | 100.00 | 42.93 | 42.44 | 44.12 | 42.16 | 42.16 | 44.60 | 42.51 |
| 23: EED18833.1-TsRbtR | 31.18 | 28.35 | 30.86 | 35.98 | 34.58 | 34.42 | 33.18 | 37.33 | 39.55 | 31.65 | 43.87 | 43.48 | 46.12 | 46.12 | 44.17 | 43.20 | 42.22 | 40.00 | 42.44 | 42.44 | 42.93 | 42.93 | 100.00 | 90.00 | 76.74 | 69.85 | 69.85 | 52.32 | 48.93 |
| 24: TfL8 | 30.98 | 26.70 | 31.21 | 37.26 | 36.02 | 35.85 | 33.94 | 38.14 | 40.55 | 33.64 | 43.75 | 44.17 | 46.83 | 46.83 | 45.37 | 44.39 | 42.99 | 39.45 | 41.95 | 41.95 | 42.44 | 42.44 | 90.00 | 100.00 | 76.38 | 70.54 | 70.54 | 51.50 | 48.91 |
| 25: QTE76006.1-ScyR6 | 32.97 | 30.05 | 32.18 | 38.50 | 38.03 | 34.58 | 33.93 | 39.91 | 41.63 | 33.49 | 46.23 | 43.69 | 46.34 | 46.34 | 45.37 | 45.37 | 45.33 | 40.91 | 43.63 | 43.63 | 44.12 | 44.12 | 76.74 | 76.38 | 100.00 | 80.69 | 80.69 | 52.94 | 51.71 |
| 26: QTE75988.1mod-ZopL4 | 31.89 | 28.50 | 31.61 | 36.15 | 35.68 | 33.18 | 33.04 | 38.07 | 39.82 | 32.11 | 47.17 | 43.20 | 45.37 | 45.37 | 44.88 | 44.88 | 42.67 | 40.00 | 41.67 | 41.67 | 42.16 | 42.16 | 69.85 | 70.54 | 80.69 | 100.00 | 100.00 | 52.94 | 49.57 |
| 27: BBU42022.1-ZopC | 31.89 | 28.50 | 31.61 | 36.15 | 35.68 | 33.18 | 33.04 | 38.07 | 39.82 | 32.11 | 47.17 | 43.20 | 45.37 | 45.37 | 44.88 | 44.88 | 42.67 | 40.00 | 41.67 | 41.67 | 42.16 | 42.16 | 69.85 | 70.54 | 80.69 | 100.00 | 100.00 | 52.94 | 49.57 |
| 28: ASK38714.1-PvL3 | 32.80 | 31.47 | 29.78 | 34.70 | 34.72 | 35.05 | 30.43 | 31.70 | 33.92 | 30.54 | 43.05 | 45.75 | 46.92 | 46.92 | 44.55 | 45.02 | 45.38 | 44.10 | 45.54 | 45.54 | 44.60 | 44.60 | 52.32 | 51.50 | 52.94 | 52.94 | 52.94 | 100.00 | 61.26 |
| 29: A0A225AJA6-Ta | 29.79 | 29.15 | 28.89 | 33.95 | 33.64 | 30.19 | 32.30 | 35.59 | 36.89 | 30.60 | 44.50 | 48.08 | 49.28 | 49.28 | 46.38 | 45.89 | 44.21 | 41.52 | 43.00 | 43.00 | 42.51 | 42.51 | 48.93 | 48.91 | 51.71 | 49.57 | 49.57 | 61.26 | 100.00 |

### PEBP

Table S9: Percentage identity matrix of maleidride PEBPs types 1 and 2 generated using T-coffee.[25]

| | 1: | 2: | 3: | 4 | 5: | 6: | 7: | 8: | 9: | 10: | 11: | 12: | 13: | 14: | 15: | 16: | 17: | 18: | 19: | 20: | 21: | 22: | 23: | 24: | 25: |
|---|---|---|---|---|---|---|---|---|---|---|---|---|---|---|---|---|---|---|---|---|---|---|---|---|---|
| 1: BfL5_ANF07283.1_PEBP1 | 100.00 | 19.75 | 45.63 | 45.63 | 46.19 | 46.67 | 39.13 | 37.56 | 19.50 | 38.33 | 47.20 | 15.92 | 15.62 | 40.21 | 38.28 | 38.05 | 41.83 | 39.42 | 27.98 | 42.31 | 20.25 | 29.76 | 35.90 | 23.57 | 36.46 |
| 2: BfL9_ANF07279.1_PEBP2 | 19.75 | 100.00 | 22.93 | 22.93 | 24.22 | 24.22 | 24.38 | 19.88 | 38.35 | 17.31 | 20.99 | 45.10 | 35.78 | 20.50 | 16.77 | 20.61 | 19.14 | 20.50 | 23.08 | 17.90 | 37.07 | 21.68 | 20.61 | 45.59 | 23.87 |
| 3: Bm48331_XP_014073928.1_PEBP1 | 45.63 | 22.93 | 100.00 | 100.00 | 84.54 | 84.54 | 42.08 | 34.50 | 22.08 | 37.14 | 62.32 | 17.11 | 20.65 | 38.10 | 37.25 | 38.50 | 40.59 | 39.90 | 26.19 | 41.58 | 21.57 | 27.38 | 36.84 | 23.03 | 38.20 |
| 4: BmC5_EMD85582.1_PEBP1 | 45.63 | 22.93 | 100.00 | 100.00 | 84.54 | 84.54 | 42.08 | 34.50 | 22.08 | 37.14 | 62.32 | 17.11 | 20.65 | 38.10 | 37.25 | 38.50 | 40.59 | 39.90 | 26.19 | 41.58 | 21.57 | 27.38 | 36.84 | 23.03 | 38.20 |
| 5: Bv_XP_014553702.1_PEBP1 | 46.19 | 24.22 | 84.54 | 84.54 | 100.00 | 97.17 | 41.95 | 33.66 | 21.38 | 37.43 | 61.32 | 19.23 | 21.38 | 38.14 | 37.80 | 37.56 | 42.03 | 41.83 | 26.79 | 41.55 | 22.93 | 28.57 | 35.90 | 22.44 | 38.33 |
| 6: Bz_XP_007715668.1_PEBP1 | 46.67 | 24.22 | 84.54 | 84.54 | 97.17 | 100.00 | 41.95 | 33.17 | 21.38 | 36.87 | 61.32 | 19.87 | 22.64 | 38.14 | 37.32 | 37.07 | 42.03 | 41.35 | 27.38 | 42.03 | 23.57 | 28.57 | 34.87 | 22.44 | 38.33 |
| 7: Ca_PVH77197.1_PEBP1 | 39.13 | 24.38 | 42.08 | 42.08 | 41.95 | 41.95 | 100.00 | 35.15 | 17.31 | 41.67 | 41.83 | 18.71 | 14.65 | 35.79 | 43.35 | 30.50 | 56.07 | 42.57 | 29.59 | 55.14 | 17.31 | 28.24 | 31.05 | 21.29 | 37.02 |
| 8: Cc_RDW56963.1_PEBP1 | 37.56 | 19.88 | 34.50 | 34.50 | 33.66 | 33.17 | 35.15 | 100.00 | 18.12 | 39.13 | 37.32 | 17.20 | 18.87 | 43.00 | 34.30 | 33.82 | 36.06 | 37.14 | 29.41 | 34.13 | 22.15 | 29.82 | 34.52 | 22.29 | 42.25 |
| 9: Cc_RDW56972.1_PEBP2 | 19.50 | 38.35 | 22.08 | 22.08 | 21.38 | 21.38 | 17.31 | 18.12 | 100.00 | 15.79 | 19.63 | 39.51 | 50.00 | 19.88 | 15.00 | 18.40 | 16.05 | 16.15 | 19.15 | 16.67 | 51.44 | 17.73 | 19.02 | 44.39 | 19.61 |
| 10: Czm_KAF2215738.1mod_PEBP1 | 38.33 | 17.31 | 37.14 | 37.14 | 37.43 | 36.87 | 41.67 | 39.13 | 15.79 | 100.00 | 40.88 | 13.82 | 13.07 | 37.70 | 40.22 | 35.14 | 43.72 | 41.44 | 25.44 | 45.36 | 18.42 | 27.06 | 30.81 | 14.47 | 40.88 |
| 11: EpiR12_PEBP1 | 47.20 | 20.99 | 62.32 | 62.32 | 61.32 | 61.32 | 41.83 | 37.32 | 19.63 | 40.88 | 100.00 | 19.11 | 18.63 | 36.87 | 40.76 | 38.83 | 41.01 | 42.92 | 27.98 | 41.47 | 20.75 | 27.98 | 35.71 | 22.29 | 38.59 |
| 12: EpiR4_PEPB2 | 15.92 | 45.10 | 17.11 | 17.11 | 19.23 | 19.87 | 18.71 | 17.20 | 39.51 | 13.82 | 19.11 | 100.00 | 35.82 | 16.67 | 14.74 | 16.35 | 14.01 | 16.67 | 14.89 | 13.38 | 35.61 | 16.31 | 17.61 | 43.41 | 15.89 |
| 13: PhiB_BBG28499.1_PEBP2 | 15.62 | 35.78 | 20.65 | 20.65 | 21.38 | 22.64 | 14.65 | 18.87 | 50.00 | 13.07 | 18.63 | 35.82 | 100.00 | 18.75 | 13.84 | 17.90 | 11.25 | 15.00 | 17.02 | 10.62 | 57.64 | 15.60 | 17.90 | 39.80 | 18.30 |
| 14: PhiN_BBG28511.1mod_PEBP1 | 40.21 | 20.50 | 38.10 | 38.10 | 38.14 | 38.14 | 35.79 | 43.00 | 19.88 | 37.70 | 36.87 | 16.67 | 18.75 | 100.00 | 31.98 | 39.90 | 36.22 | 31.84 | 27.81 | 35.71 | 22.15 | 26.47 | 35.03 | 19.23 | 45.74 |
| 15: PvR1_ASK38719.1_PEBP1 | 38.28 | 16.77 | 37.25 | 37.25 | 37.80 | 37.32 | 43.35 | 34.30 | 15.00 | 40.22 | 40.76 | 14.74 | 13.84 | 31.98 | 100.00 | 30.58 | 43.48 | 44.08 | 24.40 | 44.93 | 15.92 | 25.60 | 32.65 | 16.03 | 36.46 |
| 16: ScyL1_QTE75999.1_PEBP1 | 38.05 | 20.61 | 38.50 | 38.50 | 37.56 | 37.07 | 30.50 | 33.82 | 18.40 | 35.14 | 38.83 | 16.35 | 17.90 | 39.90 | 30.58 | 100.00 | 32.02 | 32.37 | 27.65 | 30.54 | 19.38 | 29.24 | 65.69 | 21.38 | 37.70 |
| 17: TSrbtO_EED18836.1_PEBP1 | 41.83 | 19.14 | 40.59 | 40.59 | 42.03 | 42.03 | 56.07 | 36.06 | 16.05 | 43.72 | 41.01 | 14.01 | 11.25 | 36.22 | 43.48 | 32.02 | 100.00 | 42.79 | 31.95 | 71.05 | 14.47 | 33.53 | 31.09 | 15.29 | 38.17 |
| 18: Ta_OKL57043.1_PEBP1 | 39.42 | 20.50 | 39.90 | 39.90 | 41.83 | 41.35 | 42.57 | 37.14 | 16.15 | 41.44 | 42.92 | 16.67 | 15.00 | 31.84 | 44.08 | 32.37 | 42.79 | 100.00 | 25.60 | 42.31 | 15.82 | 26.79 | 31.98 | 19.23 | 37.84 |
| 19: TfL3_PEBP1 | 27.98 | 23.08 | 26.19 | 26.19 | 26.79 | 27.38 | 29.59 | 29.41 | 19.15 | 25.44 | 27.98 | 14.89 | 17.02 | 27.81 | 24.40 | 27.65 | 31.95 | 25.60 | 100.00 | 30.18 | 17.73 | 79.43 | 26.47 | 19.86 | 30.00 |
| 20: TfL5_PEBP1 | 42.31 | 17.90 | 41.58 | 41.58 | 41.55 | 42.03 | 55.14 | 34.13 | 16.67 | 45.36 | 41.47 | 13.38 | 10.62 | 35.71 | 44.93 | 30.54 | 71.05 | 42.31 | 30.18 | 100.00 | 14.47 | 30.00 | 30.05 | 17.20 | 39.78 |
| 21: Ts1_EED15405.1mod_PEBP2 | 20.25 | 37.07 | 21.57 | 21.57 | 22.93 | 23.57 | 17.31 | 22.15 | 51.44 | 18.42 | 20.75 | 35.61 | 57.64 | 22.15 | 15.92 | 19.38 | 14.47 | 15.82 | 17.73 | 14.47 | 100.00 | 17.73 | 23.12 | 43.90 | 22.37 |
| 22: TsRbtM_EED18838.1_PEBP1 | 29.76 | 21.68 | 27.38 | 27.38 | 28.57 | 28.57 | 28.24 | 29.82 | 17.73 | 27.06 | 27.98 | 16.31 | 15.60 | 26.47 | 25.60 | 29.24 | 33.53 | 26.79 | 79.43 | 30.00 | 17.73 | 100.00 | 28.07 | 19.86 | 29.24 |
| 23: ZopL1_QTE75991.1_PEBP1 | 35.90 | 20.61 | 36.84 | 36.84 | 35.90 | 34.87 | 31.05 | 34.52 | 19.02 | 30.81 | 35.71 | 17.61 | 17.90 | 35.03 | 32.65 | 65.69 | 31.09 | 31.98 | 26.47 | 30.05 | 23.12 | 28.07 | 100.00 | 23.27 | 34.43 |
| 24: scyR12_QTE76012.1_PEBP2 | 23.57 | 45.59 | 23.03 | 23.03 | 22.44 | 22.44 | 21.29 | 22.29 | 44.39 | 14.47 | 22.29 | 43.41 | 39.80 | 19.23 | 16.03 | 21.38 | 15.29 | 19.23 | 19.86 | 17.20 | 43.90 | 19.86 | 23.27 | 100.00 | 20.53 |
| 25: Ts1_EED15411.1_PEBP1 | 36.46 | 23.87 | 38.20 | 38.20 | 38.33 | 38.33 | 37.02 | 42.25 | 19.61 | 40.88 | 38.59 | 15.89 | 18.30 | 45.74 | 36.46 | 37.70 | 38.17 | 37.84 | 30.00 | 39.78 | 22.37 | 29.24 | 34.43 | 20.53 | 100.00 |

### AMP CoA ligase

Table S10: Percentage identity matrix of maleidride AMP CoA ligases generated using MUSCLE,[1] including AMP CoA ligases from the similar oryzine biosynthetic pathway and the squalestatin pathway.

| | 1: | 2: | 3: | 4: | 5: | 6: | 7: | 8: | 9: | 10: | 11: | 12: | 13: |
|---|---|---|---|---|---|---|---|---|---|---|---|---|---|
| 1: P9WEZ0-OryP_oryzines | 100.00 | 20.87 | 19.50 | 20.58 | 20.58 | 19.59 | 19.80 | 23.16 | 22.22 | 22.15 | 22.76 | 21.95 | 22.15 |
| 2: A0A3G1DJF8.1-Mfm9_squalestatin | 20.87 | 100.00 | 23.97 | 21.88 | 21.88 | 22.07 | 22.07 | 25.29 | 25.09 | 27.51 | 26.19 | 25.57 | 25.62 |
| 3: EpiR12_Wa | 19.50 | 23.97 | 100.00 | 69.88 | 69.88 | 68.60 | 68.77 | 48.29 | 50.89 | 50.89 | 50.18 | 51.69 | 51.42 |
| 4: XP_014073927.1-Bm48331 | 20.58 | 21.88 | 69.88 | 100.00 | 100.00 | 88.48 | 88.66 | 45.16 | 47.26 | 47.43 | 48.50 | 48.06 | 47.61 |
| 5: EMD85583.1-BmC5 | 20.58 | 21.88 | 69.88 | 100.00 | 100.00 | 88.48 | 88.66 | 45.16 | 47.26 | 47.43 | 48.50 | 48.06 | 47.61 |
| 6: XP_007715669.1-Bz | 19.59 | 22.07 | 68.60 | 88.48 | 88.48 | 100.00 | 98.97 | 44.62 | 48.08 | 47.74 | 48.08 | 48.00 | 47.74 |
| 7: XP_014553703.1-Bv | 19.80 | 22.07 | 68.77 | 88.66 | 88.66 | 98.97 | 100.00 | 44.62 | 48.26 | 47.74 | 48.08 | 48.17 | 47.91 |
| 8: KAF2215735.1-Czm | 23.16 | 25.29 | 48.29 | 45.16 | 45.16 | 44.62 | 44.62 | 100.00 | 55.36 | 59.00 | 59.17 | 59.97 | 58.30 |
| 9: OKL57044.1-Ta | 22.22 | 25.09 | 50.89 | 47.26 | 47.26 | 48.08 | 48.26 | 55.36 | 100.00 | 59.83 | 60.00 | 61.64 | 61.71 |
| 10: EED18837.1-TsRbtN | 22.15 | 27.51 | 50.89 | 47.43 | 47.43 | 47.74 | 47.74 | 59.00 | 59.83 | 100.00 | 87.05 | 73.38 | 71.04 |
| 11: TfL4 | 22.76 | 26.19 | 50.18 | 48.50 | 48.50 | 48.08 | 48.08 | 59.17 | 60.00 | 87.05 | 100.00 | 73.04 | 72.06 |
| 12: KIN05362.1-Om | 21.95 | 25.57 | 51.69 | 48.06 | 48.06 | 48.00 | 48.17 | 59.97 | 61.64 | 73.38 | 73.04 | 100.00 | 74.62 |
| 13: PVH77196.1-Ca | 22.15 | 25.62 | 51.42 | 47.61 | 47.61 | 47.74 | 47.91 | 58.30 | 61.71 | 71.04 | 72.06 | 74.62 | 100.00 |

#### Isochorismatase(ICM)-like enzymes

Table S11: Percentage identity matrix of maleidride isochorismatase-like enzymes generated using MUSCLE.[1]

| | 1: | 2: | 3: | 4: | 5: | 6: | 7: | 8: | 9: | 10: | 11: | 12: | 13: | 14: |
|---|---|---|---|---|---|---|---|---|---|---|---|---|---|---|
| 1: KIN05357.1-Om | 100.00 | 68.87 | 44.29 | 48.06 | 26.70 | 26.70 | 25.13 | 25.13 | 25.14 | 27.62 | 27.62 | 27.78 | 25.56 | 28.33 |
| 2: PVH77207.1-Ca | 68.87 | 100.00 | 44.93 | 44.83 | 24.08 | 24.08 | 22.99 | 22.99 | 25.14 | 23.76 | 23.76 | 25.00 | 23.33 | 25.00 |
| 3: KAF2215730.1-Czm | 44.29 | 44.93 | 100.00 | 60.19 | 20.63 | 20.63 | 22.70 | 22.70 | 22.47 | 24.44 | 24.44 | 25.70 | 22.35 | 24.58 |
| 4: KAF2215740.1-Czm | 48.06 | 44.83 | 60.19 | 100.00 | 21.69 | 21.69 | 21.08 | 21.08 | 20.22 | 25.00 | 25.00 | 24.58 | 24.02 | 26.26 |
| 5: EMD85567.1-BmC5 | 26.70 | 24.08 | 20.63 | 21.69 | 100.00 | 100.00 | 91.00 | 91.00 | 40.76 | 37.10 | 37.10 | 40.00 | 40.00 | 37.30 |
| 6: XP_014073915.1-Bm48331 | 26.70 | 24.08 | 20.63 | 21.69 | 100.00 | 100.00 | 91.00 | 91.00 | 40.76 | 37.10 | 37.10 | 40.00 | 40.00 | 37.30 |
| 7: XP_014553686.1-Bv | 25.13 | 22.99 | 22.70 | 21.08 | 91.00 | 91.00 | 100.00 | 100.00 | 40.00 | 35.16 | 35.16 | 38.67 | 38.12 | 34.25 |
| 8: XP_007715656.1-Bz | 25.13 | 22.99 | 22.70 | 21.08 | 91.00 | 91.00 | 100.00 | 100.00 | 40.00 | 35.16 | 35.16 | 38.67 | 38.12 | 34.25 |
| 9: OKL57229.1_Ta | 25.14 | 25.14 | 22.47 | 20.22 | 40.76 | 40.76 | 40.00 | 40.00 | 100.00 | 43.48 | 43.48 | 43.17 | 43.17 | 42.08 |
| 10: QTE75990.1-ZopL2 | 27.62 | 23.76 | 24.44 | 25.00 | 37.10 | 37.10 | 35.16 | 35.16 | 43.48 | 100.00 | 100.00 | 70.27 | 68.65 | 70.27 |
| 11: BBU42024.1-ZopQ | 27.62 | 23.76 | 24.44 | 25.00 | 37.10 | 37.10 | 35.16 | 35.16 | 43.48 | 100.00 | 100.00 | 70.27 | 68.65 | 70.27 |
| 12: QTE76004.1-ScyR4 | 27.78 | 25.00 | 25.70 | 24.58 | 40.00 | 40.00 | 38.67 | 38.67 | 43.17 | 70.27 | 70.27 | 100.00 | 71.35 | 68.11 |
| 13: EED18834.1mod-TsRbtQ | 25.56 | 23.33 | 22.35 | 24.02 | 40.00 | 40.00 | 38.12 | 38.12 | 43.17 | 68.65 | 68.65 | 71.35 | 100.00 | 81.08 |
| 14: TfL7 | 28.33 | 25.00 | 24.58 | 26.26 | 37.30 | 37.30 | 34.25 | 34.25 | 42.08 | 70.27 | 70.27 | 68.11 | 81.08 | 100.00 |

#### Enoyl CoA isomerases

Table S12: Percentage identity matrix generated using MUSCLE[1] of maleidride enoyl CoA isomerases, as well as an identified homologue from the *A. oryzae* genome (BAE65732.1).

| | 1: | 2: | 3: | 4: | 5: | 6: | 7: | 8: | 9: | 10 | 11: |
|---|---|---|---|---|---|---|---|---|---|---|---|
| 1: TsR1 | 100.00 | 15.91 | 21.76 | 20.27 | 19.63 | 19.63 | 19.35 | 16.44 | 17.24 | 18.94 | 15.91 |
| 2: EpiR10 | 15.91 | 100.00 | 23.20 | 21.7 | 18.78 | 18.78 | 23.90 | 21.94 | 22.51 | 20.78 | 22.94 |
| 3: Cc-RDW56966.1 | 21.76 | 23.20 | 100.00 | 21.36 | 26.51 | 26.51 | 27.36 | 26.98 | 23.56 | 22.37 | 22.51 |
| 4: BfL10-ANF07277.1 | 20.27 | 21.70 | 21.36 | 100.00 | 23.27 | 23.27 | 28.37 | 24.62 | 25.00 | 22.41 | 20.69 |
| 5: ZopR2-QTE75994.1 | 19.63 | 18.78 | 26.51 | 23.27 | 100.00 | 100.00 | 25.59 | 24.74 | 29.78 | 26.32 | 23.14 |
| 6: ZopS | 19.63 | 18.78 | 26.51 | 23.27 | 100.00 | 100.00 | 25.59 | 24.74 | 29.78 | 26.32 | 23.14 |
| 7: ScyR2-QTE76002.1 | 19.35 | 23.90 | 27.36 | 28.37 | 25.59 | 25.59 | 100.00 | 30.37 | 25.99 | 30.84 | 29.52 |
| 8: BAE65732.1 A. oryzae | 16.44 | 21.94 | 26.98 | 24.62 | 24.74 | 24.74 | 30.37 | 100.00 | 45.83 | 47.08 | 44.17 |
| 9: Ag-OJJ86355.1 | 17.24 | 22.51 | 23.56 | 25.00 | 29.78 | 29.78 | 25.99 | 45.83 | 100.00 | 55.20 | 51.61 |
| 10: Ia-CAF9941821.1 | 18.94 | 20.78 | 22.37 | 22.41 | 26.32 | 26.32 | 30.84 | 47.08 | 55.20 | 100.00 | 62.06 |
| 11: TbL5 | 15.91 | 22.94 | 22.51 | 20.69 | 23.14 | 23.14 | 29.52 | 44.17 | 51.61 | 62.06 | 100.00 |

#### Maleidride conserved proteins

Table S13: Percentage identity matrix generated using MUSCLE[1] of maleidride conserved proteins, as well as identified homologues.

| | 1: | 2: | 3: | 4: | 5: | 6: | 7: | 8: | 9: | 10: | 11: | 12: | 13: | 14: |
|---|---|---|---|---|---|---|---|---|---|---|---|---|---|---|
| 1: AlnI-C8VJQ9.1 | 100.00 | 12.72 | 18.95 | 19.80 | 22.05 | 22.05 | 23.04 | 15.31 | 15.31 | 15.82 | 15.82 | 13.71 | 17.39 | 16.10 |
| 2: BAE60519.1 | 12.72 | 100.00 | 36.00 | 19.53 | 21.56 | 21.56 | 19.64 | 19.88 | 19.88 | 21.08 | 21.08 | 20.73 | 18.93 | 23.67 |
| 3: BfL8-ANF07280.1 | 18.95 | 36.00 | 100.00 | 26.42 | 24.61 | 24.61 | 26.56 | 30.39 | 30.39 | 29.28 | 29.28 | 25.14 | 29.02 | 29.69 |
| 4: ScyR11-QTE76011.1 | 19.80 | 19.53 | 26.42 | 100.00 | 55.91 | 55.91 | 28.18 | 29.10 | 29.10 | 27.51 | 27.51 | 24.34 | 26.58 | 25.74 |
| 5: ZopL8-QTE75984.1 | 22.05 | 21.56 | 24.61 | 55.91 | 100.00 | 100.00 | 27.52 | 26.74 | 26.74 | 25.67 | 25.67 | 22.99 | 25.91 | 25.50 |
| 6: ZopP-BBU42018.1 | 22.05 | 21.56 | 24.61 | 55.91 | 100.00 | 100.00 | 27.52 | 26.74 | 26.74 | 25.67 | 25.67 | 22.99 | 25.91 | 25.50 |
| 7: EpiR3 | 23.04 | 19.64 | 26.56 | 28.18 | 27.52 | 27.52 | 100.00 | 47.96 | 47.96 | 47.45 | 47.45 | 27.55 | 28.07 | 29.61 |
| 8: BmC5-EMD85573.1 | 15.31 | 19.88 | 30.39 | 29.10 | 26.74 | 26.74 | 47.96 | 100.00 | 100.00 | 86.29 | 86.29 | 25.13 | 28.72 | 26.67 |
| 9: Bm-XP_014073934.1 | 15.31 | 19.88 | 30.39 | 29.10 | 26.74 | 26.74 | 47.96 | 100.00 | 100.00 | 86.29 | 86.29 | 25.13 | 28.72 | 26.67 |
| 10: Bv-XP_014553692.1 | 15.82 | 21.08 | 29.28 | 27.51 | 25.67 | 25.67 | 47.45 | 86.29 | 86.29 | 100.00 | 100.00 | 24.61 | 26.67 | 26.67 |
| 11: Bz-XP_007715652.1 | 15.82 | 21.08 | 29.28 | 27.51 | 25.67 | 25.67 | 47.45 | 86.29 | 86.29 | 100.00 | 100.00 | 24.61 | 26.67 | 26.67 |
| 12: Czm-KAF2215736.1 | 13.71 | 20.73 | 25.14 | 24.34 | 22.99 | 22.99 | 27.55 | 25.13 | 25.13 | 24.61 | 24.61 | 100.00 | 27.41 | 31.61 |
| 13: PvL16-ASK38701.1 | 17.39 | 18.93 | 29.02 | 26.58 | 25.91 | 25.91 | 28.07 | 28.72 | 28.72 | 26.67 | 26.67 | 27.41 | 100.00 | 32.39 |
| 14: TaR7 | 16.10 | 23.67 | 29.69 | 25.74 | 25.50 | 25.50 | 29.61 | 26.67 | 26.67 | 26.67 | 26.67 | 31.61 | 32.39 | 100.00 |

#### Quinone reductase

Table S14: Percentage identity matrix generated using MUSCLE[1] of maleidride quinone reductases compared to a homologue from *Pseudomonas aeruginosa*.

| | 1: | 2: | 3: |
|---|---|---|---|
| 1: Q9I4V0.1_NQRED_PSEAE | 100.00 | 52.91 | 51.99 |
| 2: Ta_OKL57052.1 | 52.91 | 100.00 | 71.21 |
| 3: Czm_KAF2215726.1 | 51.99 | 71.21 | 100.00 |

### α-ketoglutarate dependant dioxygenases (αKGDDs)

#### AsaB-like

Table S15: Percentage identity matrix for AsaB-like αKGDDs generated using MUSCLE.[1]

| | 1: | 2: | 3: | 4: | 5: | 6: | 7: | 8: | 9: | 10: | 11: | 12: | 13: | 14: |
|---|---|---|---|---|---|---|---|---|---|---|---|---|---|---|
| 1: A0A3G1DJG9.1-Mfr1 | 100.00 | 41.38 | 19.05 | 24.30 | 24.60 | 26.98 | 21.05 | 17.74 | 21.37 | 21.51 | 21.51 | 22.00 | 20.16 | 20.48 |
| 2: A0A3G1DJF4.1-Mfr2 | 41.38 | 100.00 | 21.37 | 25.30 | 23.60 | 24.40 | 21.43 | 21.92 | 21.88 | 23.08 | 23.08 | 23.58 | 20.49 | 23.27 |
| 3: OKL57227.1-T.atroroseus | 19.05 | 21.37 | 100.00 | 38.38 | 33.80 | 37.89 | 24.90 | 27.34 | 29.34 | 26.88 | 26.88 | 26.80 | 27.60 | 23.81 |
| 4: QTE75998.1-ScyL2 | 24.30 | 25.30 | 38.38 | 100.00 | 63.79 | 61.17 | 21.71 | 24.81 | 28.96 | 23.92 | 23.92 | 26.19 | 28.17 | 28.74 |
| 5: EED18844.1-TsRbtG | 24.60 | 23.60 | 33.80 | 63.79 | 100.00 | 65.75 | 21.76 | 27.10 | 27.59 | 26.46 | 26.46 | 26.38 | 27.56 | 29.69 |
| 6: EED18849.1a_mod-TsRbtB | 26.98 | 24.40 | 37.89 | 61.17 | 65.75 | 100.00 | 21.62 | 26.25 | 27.31 | 24.61 | 24.61 | 28.46 | 26.48 | 27.06 |
| 7: B6HLP7.1-ChyM | 21.05 | 21.43 | 24.90 | 21.71 | 21.76 | 21.62 | 100.00 | 29.75 | 25.00 | 25.38 | 25.38 | 27.27 | 26.82 | 28.24 |
| 8: S0E2Y4.1-Des | 17.74 | 21.92 | 27.34 | 24.81 | 27.10 | 26.25 | 29.75 | 100.00 | 28.62 | 29.30 | 29.30 | 33.82 | 29.74 | 29.26 |
| 9: AZL87943.1-AsaB | 21.37 | 21.88 | 29.34 | 28.96 | 27.59 | 27.31 | 25.00 | 28.62 | 100.00 | 29.37 | 29.37 | 30.86 | 30.71 | 29.32 |
| 10: QTE75983.1-ZopL9 | 21.51 | 23.08 | 26.88 | 23.92 | 26.46 | 24.61 | 25.38 | 29.30 | 29.37 | 100.00 | 100.00 | 46.38 | 43.80 | 39.56 |
| 11: BBU42017.1-ZopK | 21.51 | 23.08 | 26.88 | 23.92 | 26.46 | 24.61 | 25.38 | 29.30 | 29.37 | 100.00 | 100.00 | 46.38 | 43.80 | 39.56 |
| 12: RDW56970.1-C.crateriformis | 22.00 | 23.58 | 26.80 | 26.19 | 26.38 | 28.46 | 27.27 | 33.82 | 30.86 | 46.38 | 46.38 | 100.00 | 62.04 | 54.61 |
| 13: BBG28508.1-PhiK | 20.16 | 20.49 | 27.60 | 28.17 | 27.56 | 26.48 | 26.82 | 29.74 | 30.71 | 43.80 | 43.80 | 62.04 | 100.00 | 63.74 |
| 14: EED15414.1-T.stipitatus | 20.48 | 23.27 | 23.81 | 28.74 | 29.69 | 27.06 | 28.24 | 29.26 | 29.32 | 39.56 | 39.56 | 54.61 | 63.74 | 100.00 |

#### TauD-like

Table S16: Percentage identity matrix for TauD-like αKGDDs generated using MUSCLE.[1]

| | 1: | 2: | 3: | 4: | 5: | 6: | 7: |
|---|---|---|---|---|---|---|---|
| 1: EED18830.1mod-TsRbtU | 100.00 | 52.92 | 48.80 | 25.00 | 25.27 | 25.81 | 22.30 |
| 2: EED18846.1-TsRbtE | 52.92 | 100.00 | 77.05 | 21.68 | 25.82 | 27.44 | 22.83 |
| 3: TfL12-T.funiculosus | 48.80 | 77.05 | 100.00 | 20.63 | 26.55 | 27.80 | 21.74 |
| 4: Q2TXF3.1-OryG | 25.00 | 21.68 | 20.63 | 100.00 | 32.79 | 31.82 | 27.16 |
| 5: KIN05358.1-O.maius | 25.27 | 25.82 | 26.55 | 32.79 | 100.00 | 73.47 | 32.39 |
| 6: PVH77204.1-Cadophora | 25.81 | 27.44 | 27.80 | 31.82 | 73.47 | 100.00 | 32.77 |
| 7: sA0A0A2IJP3.1-CnsP | 22.30 | 22.83 | 21.74 | 27.16 | 32.39 | 32.77 | 100.00 |

#### PhyH-like

Table S17: Percentage identity matrix for PhyH-like αKGDDs generated using MUSCLE.[1]

| | 1: | 2: | 3: | 4: |
|---|---|---|---|---|
| 1: OKL57048.1-T.atroroseus | 100.00 | 35.54 | 24.91 | 24.23 |
| 2: A0A2I1BSW6.2-NvfE | 35.54 | 100.00 | 20.14 | 20.96 |
| 3: Q5AR53.1-AsqJ | 24.91 | 20.14 | 100.00 | 27.02 |
| 4: Q5AR34-AusE | 24.23 | 20.96 | 27.02 | 100.00 |

#### IPNS-like

Table S18: Percentage identity matrix for IPNS-like αKGDDs generated using MUSCLE.[1]

| | 1: | 2: | 3: |
|---|---|---|---|
| 1: A0A159BP93.1-CitB | 100.00 | 23.02 | 24.22 |
| 2: ASK38712.1-PvL5 | 23.02 | 100.00 | 29.09 |
| 3: Q4WKX0.1-FgnB | 24.22 | 29.09 | 100.00 |

### Isochorismatase I-Tasser analysis

Table S19: I-Tasser analysis for a selection of isochorismatase-like enzymes encoded by maleidride BGCs.

| Species | Isochorismatase | Predicted Ligand | C-score | Ligand Binding Site Residues |
|---|---|---|---|---|
| ***O. maius* Zn** | KIN05357.1 | Maleic acid | 0.45 | 30,35,70,82,91,149,150,153,154,179 |
| ***S. album*** | QTE76004.1 | Maleic acid | 0.37 | 10,15,47,62,124,125,128,129,154 |
| ***B. maydis*** | XP_014073915.1 | Maleic acid | 0.37 | 15,20,62,83,154,155,158,159,184 |

C-score is the confidence score of the prediction. C-score ranges [0-1], where a higher score indicates a more reliable prediction.

### Genbank files

#### CaL1

```
LOCUS       PCYN01000097            2278 bp    DNA     linear   PLN 30-APR-2018
DEFINITION  Cadophora sp. DSE1049 DL98scaffold_72_Cont97, whole genome shotgun
            sequence.
ACCESSION   PCYN01000097 REGION: 18734..21011
VERSION     PCYN01000097.1
DBLINK      BioProject: PRJNA243951
            BioSample: SAMN02745222
KEYWORDS    WGS.
SOURCE      Cadophora sp. DSE1049
  ORGANISM  Cadophora sp. DSE1049
            Eukaryota; Fungi; Dikarya; Ascomycota; Pezizomycotina;
            Leotiomycetes; Helotiales; Helotiales incertae sedis; Cadophora.
REFERENCE   1  (bases 1 to 2278)
  AUTHORS   Knapp,D.G., Nemeth,J.B., Barry,K., Hainaut,M., Henrissat,B.,
            Johnson,J., Kuo,A., Lim,J.H.P., Lipzen,A., Nolan,M., Ohm,R.A.,
            Tamas,L., Grigoriev,I.V., Spatafora,J.W., Nagy,L.G. and Kovacs,G.M.
  TITLE     Comparative genomics provides insights into the lifestyle and
            reveals functional heterogeneity of dark septate endophytic fungi
  JOURNAL   Sci Rep 8 (1), 6321 (2018)
   PUBMED   29679020
  REMARK    Publication Status: Online-Only
REFERENCE   2  (bases 1 to 2278)
  AUTHORS   Ohm,R., Kuo,A., Knapp,D.G., Nemeth,J.B., Barry,K., Hainaut,M.,
            Henrissat,B., Johnson,J., Lim,J., Lipzen,A., Nolan,M., Tamas,L.,
            Grigoriev,I.V., Spatafora,J.W., Nagy,L.G., Kovacs,G.M.,
            Nordberg,H.P., Cantor,M.N. and Hua,S.X.
  CONSRTM   DOE Joint Genome Institute
  TITLE     Direct Submission
  JOURNAL   Submitted (29-SEP-2017) DOE Joint Genome Institute, 2800 Mitchell
            Drive, Walnut Creek, CA 94598-1698, USA
COMMENT     URL -- http://genome.jgi.doe.gov/Cadsp1
            JGI Project ID: 1025590
            The DNA was provided by Gabor M. Kovacs, Daniel G. Knapp
           
            The strain is available from Gabor M. Kovacs, Daniel G. Knapp
            Contacts: Gabor  Kovacs
            Assembly and annotation done by JGI.
            The JGI and collaborators endorse the principles for the
            distribution and use of large scale sequencing data adopted by the
            larger genome sequencing community and urge users of this data to
            follow them. It is our intention to publish the work of this
            project in a timely fashion and we welcome collaborative
            interaction on the project and analysis.
            (http://www.genome.gov/page.cfm?pageID=10506376).

            ##Metadata-START##
            Organism Display Name :: Cadophora sp. DSE1049 v1.0
            GOLD Stamp ID         :: Gp0046570
            ##Metadata-END##
```

```
            ##Genome-Assembly-Data-START##
            Assembly Date          :: 14-FEB-2014
            Assembly Method        :: AllPathsLG v. R47710
            Assembly Name          :: Cadsp1
            Long Assembly Name     :: Cadophora sp. DSE1049 v1.0
            Genome Coverage        :: 79.2x
            Sequencing Technology :: Illumina
            ##Genome-Assembly-Data-END##
FEATURES             Location/Qualifiers
     CDS             join(812..884,1002..1603)
                     /gene="caL1"
                     /note="Hydrolase"
BASE COUNT      637 a    477 c    491 g    673 t
ORIGIN
        1 gagcacgcgg gctaggacga aattcactta ttgtatgagt aagagataag taaccggtga
       61 ggacttattt gctcttaaaa tccttgaata caaaatagct cgggtatcaa aggcacgaat
      121 aaatactaga aggtattaag gcaatccatg tgctagtagg atccttctat ctcatcacat
      181 atcaccacta gcactaaata ttttcggatt caagaataat taccttttta tctctctata
      241 tcctatataa cttccctagc ttagatttaa tcgagcaagt attgtgaccc ttagaactcc
      301 cacacttgca ctattgtcgc tcattcaaga gcatttttgc taacaccagc attattaata
      361 aaatatcttt aattttgtaa atgcgccggc aactttacgc aaaagtctgc ttggttaatt
      421 gagctgtcta gttgagtata tgtcggcaaa gcacgaaatt acatgcgtcg aacagctgga
      481 aatttagcca ttggttaata gtaaccgaaa accctttttt tggtaccatt cgagacacta
      541 aggtttctga tacccttgaa cagtcggaaa catttggaac atatcatatt caaagcccta
      601 attagtctat gtcttgggaa tatttgttga ttctgctcga gtgtaaaaat actctcaaca
      661 ccacagtatc ggcacatttc cgatccagtg taacttgtct tgattgtcga ctactccaat
      721 acttcgccta gatttcatat cttgatctca ctcctttgaa aagaattact tgacattaaa
      781 gtttagacaa aaatctctct ggactaggaa gatgcctcta aaattcctct gtcttcatgg
      841 ctggggtacc aactccaagg taagtactct cctcgatggg gtcggtccat tctggggatg
      901 atcgctgata cttaggtctt cccatagatt ttagaatcgc agcttggtat gagattcttc
      961 aagacttagc aactttggcg ttaatttaat aatgtacata ggtgctttga tgcgagagct
     1021 caaacgagac aacaccgccg aattttattt ctttgaaggc gacctggact ccgagcctgg
     1081 cccaggcatc gaaggttact acacaggtcc tttctacagc tactacaagt tcccacggtc
     1141 attcgaagat agcgatgagt cgatgttaga ggcatacgaa ctgttggacg agatcgtgga
     1201 ggaagaaggg cctttcgacg gcgtgctagg gttctcccac ggagggaccc tcgcaagcgg
     1261 gtggctaata catcatgcag cgaagtatcc tttggagccg ctgccagtta gatgcgttgt
     1321 ctttctgaat tcgttgccac cgttccgtat gaaggctggc gaggatcctg tcgtagatga
     1381 aggttttaag gatggatgca ttcagatacc gagtgtcagt gttgctggta ccaaggactt
     1441 tgtttatgat tactctatca agcttcataa gctctgcgac ccacgaaagt ctcagctggt
     1501 aatccatgac aaaggtcatg atattcctag cgacgccaag aatgtggggg ccatggcaag
     1561 ggcaatcagg aaaatgtcag cggacgcaat acaagcatgg tagaatgaga tttgtcttcc
     1621 agagaaagtg gctagatatg agtaggttat ttgaacagac aataaatcca ctgaattcca
     1681 aatttacctt cgcttttctt cctgagccat ttcccacacc gcgatagata cgtactcatg
     1741 gtgctacaga gcatcatttt tttggggctt aatcagccag cactgagagc tcttttaagt
     1801 atctctactt aaaagaacaa acaagaacgt ggtgatcttg cgtcggctat ttctgtcagt
     1861 gctatagaaa ttctagtaaa ccgacacact tatctaagcg atgtgaaatg ctccgacaaa
     1921 gcccgtcaaa actgcgatgg ccgagaaaca ttcaagctta attgaagcgg tgaacagttc
     1981 tataaggcta gttgggtcat ttaattaggt agataattta gcatagaggt caggccaggc
     2041 catattttga ataatttttg tactttggat atgcacatga atttcgcctc aggcgatttt
     2101 ttatttttaa atctggattt gcccgagctt agggggcctt tctttgactc ggacgggttt
     2161 agccagattc gacccaaatt taacacgctg attgacggac tccaatctgg tctcgaagcg
     2221 ttcttgatgc ttagaatcct gctttgtctc gaataaatcc gtagttgtct tgaaatgg
//
```

#### CzmL1

```
LOCUS       JAAEIV010000103         1984 bp    DNA     linear   PLN 31-JAN-2020
DEFINITION  Cercospora zeae-maydis SCOH1-5 CERZMscaffold_4_Cont103, whole
            genome shotgun sequence.
ACCESSION   JAAEIV010000103 REGION: 80080..82063
VERSION     JAAEIV010000103.1
DBLINK      BioProject: PRJNA68833
            BioSample: SAMN00773058
KEYWORDS    WGS.
SOURCE      Cercospora zeae-maydis SCOH1-5
  ORGANISM  Cercospora zeae-maydis SCOH1-5
            Eukaryota; Fungi; Dikarya; Ascomycota; Pezizomycotina;
            Dothideomycetes; Dothideomycetidae; Capnodiales;
            Mycosphaerellaceae; Cercospora.
REFERENCE   1  (bases 1 to 1984)
  AUTHORS   Haridas,S., Albert,R., Binder,M., Bloem,J., LaButti,K., Salamov,A.,
            Andreopoulos,B., Baker,S., Barry,K., Bills,G., Bluhm,B., Cannon,C.,
            Castanera,R., Culley,D., Daum,C., Ezra,D., Gonzalez,J.,
            Henrissat,B., Kuo,A., Liang,C., Lipzen,A., Lutzoni,F., Magnuson,J.,
```

```
            Mondo,S., Nolan,M., Ohm,R., Pangilinan,J., Park,H.-J., Ramirez,L.,
            Alfaro,M., Sun,H., Tritt,A., Yoshinaga,Y., Zwiers,L.-H.,
            Turgeon,B., Goodwin,S., Spatafora,J., Crous,P. and Grigoriev,I.
  TITLE     101 Dothideomycetes genomes: A test case for predicting lifestyles
            and emergence of pathogens
  JOURNAL   Stud. Mycol. 96, 141-153 (2020)
REFERENCE   2  (bases 1 to 1984)
  AUTHORS   Haridas,S., Albert,R., Binder,M., Bloem,J., Labutti,K., Salamov,A.,
            Andreopoulos,B., Baker,S.E., Barry,K., Bills,G., Bluhm,B.H.,
            Cannon,C., Castanera,R., Culley,D.E., Daum,C., Ezra,D.,
            Gonzalez,J.B., Henrissat,B., Kuo,A., Liang,C., Lipzen,A.,
            Lutzoni,F., Magnuson,J., Mondo,S., Nolan,M., Ohm,R., Pangilinan,J.,
            Park,H.-J., Ramirez,L., Alfaro,M., Sun,H., Tritt,A., Yoshinaga,Y.,
            Zwiers,L.-H., Turgeon,B.G., Goodwin,S.B., Spatafora,J.W.,
            Crous,P.W. and Grigoriev,I.V.
  CONSRTM   DOE Joint Genome Institute
  TITLE     Direct Submission
  JOURNAL   Submitted (28-JAN-2020) DOE Joint Genome Institute, 2800 Mitchell
            Drive, Walnut Creek, CA 94598-1698, USA
COMMENT     URL -- http://genome.jgi.doe.gov/Cerzm1
            JGI Project ID: 401984
            The DNA was provided by Stephen Goodwin
            The strain is available from Stephen Goodwin
            Contacts: Stephen Goodwin
            Assembly and annotation done by JGI.
            The JGI and collaborators endorse the principles for the
            distribution and use of large scale sequencing data adopted by the
            larger genome sequencing community and urge users of this data to
            follow them. It is our intention to publish the work of this
            project in a timely fashion and we welcome collaborative
            interaction on the project and analysis.
            (http://www.genome.gov/page.cfm?pageID=10506376)

            This draft release was produced using Roche (454), Fosmid (Sanger),
            and shredded consensus from a velvet assembly of Illumina data.

            ##Metadata-START##
            Organism Display Name :: Cercospora zeae-maydis SCOH1-5
            GOLD Stamp ID         :: Gp0009719
            ##Metadata-END##

            ##Genome-Assembly-Data-START##
            Assembly Date         :: 2011
            Assembly Method       :: Velvet v. unknown
            Assembly Name         :: Cerzm1
            Genome Coverage       :: 39.29x
            Sequencing Technology :: Roche (454); Fosmid (Sanger); Illumina
            ##Genome-Assembly-Data-END##
FEATURES             Location/Qualifiers
     CDS             complement(join(883..1134,1183..1532,1590..1608,1664..1711))
                     /note="Hydrolase"
                     /gene="czmL1"
BASE COUNT      524 a    485 c    491 g    484 t
ORIGIN
        1 acgctgatac cttgctttgg atgttaagac gtggggacta ttgctaaggg gtatcgaaag
       61 gctccgaggc tattattttg gcgcgtcgtc ctgtggacgg accgagcgac cgtgatcagg
      121 cccgcggaag tgaacatcca tttcgtattc aactataagc aaagttttgg cgtcaacagg
      181 tcgcattgtg cattcctaca cccatattag tgaccgcggc cgggacgatt gctcaaaaag
      241 tccccgccgg caagatcaga tatgccgatc ggtcttcact attgcctgga attatgtcgc
      301 ttgagctcaa tctatcaaat gcaactacaa cgagcctttg aaacaaagac gcttgagaat
      361 ccgcgggtgc ggccggggct gtagcggtga gaaggaggaa tacggaagca taaagcatgc
      421 atcgaggccg agatactcaa cagccacaat ggttttggac tggccactgc acaagttttc
      481 cactgcagag cgtcgcactt cgcctcggta tacaaagtga atttccacgt aatgatattg
      541 tgattctgaa gaatctaaga attgatttgg agggcttgaa cattgtatgt tttcaaaatg
      601 tctatctgga ggcactgtat atgtacacag catttggctc tgttaggctc catctgaagc
      661 gggtcaagcc atcataagac agaaccttct tctggagtac acgttacatg atagggcaat
      721 cacatcaagt taaagtgtca tagactatat cccaatggtt gaaaaccact gtctcggaga
      781 tcagccggaa ttgatatgtt acgtactgct ctatgtcacg gcgggcagac tttcatatac
      841 tttatttgaa ctaaaccaag taaatcaaca aaagggcttc gttcagtacc cgttttgcac
      901 ctgtatagct aatttgcgga tcgcttgcgc tatggcagct gcgttcttgg catccctagg
      961 aatgtcgtgg ccccgcttgt gggtgacgag ctgcgcagcg cgagtatcgc acagatggta
     1021 cagccgaata gaatactcat agacaaaatc attttcgccc accacatgca ctgaaggtat
     1081 cgcgatacat ccatcctcca aaccttcgtc cactacagga tcctggccgg gatgctgtat
```

```
     1141 aaaatcagac atcagctgga gcgaagaaaa ggcgatacct accttgcgaa aaggtggcaa
     1201 cgcattgaag aatagcgcac accggaacgg caggggctct gacggtcgca tcgccatgtg
     1261 gtgcataatg aatccgctcg ccaaagttcc accatgggaa aagccgagga taccatcgaa
     1321 tggtccatcg tcctcgatca catcgtacag aagctcatag gcctcaagca tggactgatc
     1381 gctgtcctca aacgtccgtg gaaacttgta atagctgtag aatggcccct catagaatcc
     1441 ttcaatgccg gggcctggct ctgattcgat atctccctca tggaaatgga attcgactgt
     1501 gtgatccacc agcaggttct tccgcacggt gtctgtgtat cgcgtcaatg atccgcgaag
     1561 aagcaattca gggccatttg acgacgtacc aagctgtgac cgcaatatct gaatcgccag
     1621 caagcgtcag cgaacgtata tccatcactt acgtgtcgct tactgagctg tttgttcccc
     1681 atccgtgtag acaaaggaat ttcaaaggca tgtttccgac tcgatcaaat caccgacgac
     1741 gatgatcaaa ttgctcaggg ggagttactt tcactcgact aggatgggaa gaatatgttc
     1801 caagaatttg tcggaaggaa tcgctcttat atgcctgcgt ccggcataaa gtctagcggc
     1861 ggctttagtc tggcacaacg agctgccaac tccaaatatc cgcgacaagc gctcctgcga
     1921 agtgatgtaa gaggcaccct gtatctggtt ctgatgcatc gggggttttg catgccaatc
     1981 aaca
//
```

#### TaR7

```
LOCUS       LFMY01000012            2053 bp    DNA     linear   PLN 03-AUG-2017
DEFINITION  Talaromyces atroroseus strain IBT 11181 scaffold_11, whole genome
            shotgun sequence.
ACCESSION   LFMY01000012 REGION: 260998..263050
VERSION     LFMY01000012.1
DBLINK      BioProject: PRJNA275056
            BioSample: SAMN03339010
KEYWORDS    WGS.
SOURCE      Talaromyces atroroseus
  ORGANISM  Talaromyces atroroseus
            Eukaryota; Fungi; Dikarya; Ascomycota; Pezizomycotina;
            Eurotiomycetes; Eurotiomycetidae; Eurotiales; Trichocomaceae;
            Talaromyces.
REFERENCE   1  (bases 1 to 2053)
  AUTHORS   Rasmussen,K.B., Rasmussen,S., Petersen,B., Sicheritz-Ponten,T.,
            Mortensen,U.H. and Thrane,U.
  TITLE     Talaromyces atroroseus IBT 11181 draft genome
  JOURNAL   Unpublished
REFERENCE   2  (bases 1 to 2053)
  AUTHORS   Rasmussen,K.B., Rasmussen,S., Petersen,B., Sicheritz-Ponten,T.,
            Mortensen,U.H. and Thrane,U.
  TITLE     Direct Submission
  JOURNAL   Submitted (24-JUN-2015) DTU Systems Biology, Technical University
            of Denmark, Soeltofts Plads 223, Kgs. Lyngby 2800, Denmark
COMMENT     ##Genome-Assembly-Data-START##
            Assembly Method       :: ALLPATHS-LG v. MAY-2013
            Genome Coverage       :: 193.0x
            Sequencing Technology :: Illumina HiSeq
            ##Genome-Assembly-Data-END##
FEATURES             Location/Qualifiers
     CDS             complement(124..765)
                     /note="Conserved maleidride protein"
                     /gene="taR7"
BASE COUNT      550 a    509 c    430 g    564 t
ORIGIN
        1 acgaaagatc tattcaagta tcaataatac aatgtcgctc tttcaattta tctagattgc
       61 tgattatcaa ttgctctaca taatatggaa atcccatggt ttattctccg ataagcgcgg
      121 ccttcattta agaagtccca tcaaactgtc cagtcgttcc cccaagttgg aaaaggacat
      181 ctcgcaggct gtcagcactg ttctctgtct aggccaagcc acttcccata gccccataag
      241 tatctcgtgc aattcatcca atatattacc taaactcgca agccgtcgtc gaagaagcac
      301 tttaatcaat aactttgcgt cctctccttt cagttccagt ccaccccagc tgatctttgc
      361 tggctctgat attgaaacag atgacactcg tccctcgatt ttctcggtca gtggttgatt
      421 agcttcagca aattgtcgct gagtgccacg atggtgtcct tcatctgaca ctatatcagc
      481 acttaaatcc tctctggtga acgagtaggc tgcgcatgca gcctcatata aggcgagcat
      541 gtactctagt gtcggtctga gggtattaaa cacgccaaaa tgagctcgac agtgatcaca
      601 tgagatcaga atatcacact gtcgagtgaa ccgcgtgcca acctcgataa acgtctcaaa
      661 ctctatggcg gacttgtcct ggatttccat cataatttga ttgatgccac tgcagcagtt
      721 gcatggcgtc acagctcgtg tcatgtcgcc gcttgaggag ctcatattat gcgtttgctg
      781 agttaatgtt gattcggaac tcaaagccgt tatgattccc ctttctcccc tataccctag
      841 ctgtaagagg gattcacttg tagcgttagt aaaattcaat tttgcaacag tatggatatg
      901 atttgcttac actttgtgaa gggaagtcct ggtctgatgc tactcaaact ccaacgtcat
      961 cagcagccca gttgaccagc ctagtttatc ctgcctgtca tttcctattc tggcgcaagc
     1021 tgagttagaa tttggccaca cgcgcatcac aatacgggcc gaaggattcc agctgaatac
     1081 attgcagccc acctggggta ccaaggtact ctggcaactt ttgaaacaaa gagggggctc
     1141 cagtagagga agcttggaag cctttgagcg gtggggatat gactcttctc gtatatattt
```

```
     1201 ggtaacgact tttctcgcaa ggaaaaaaat atgatcacaa ttgtagaaac cctgacatta
     1261 ccaactccaa ccagaaggtc ttccaatggc agatactgtc ccattcttat atcttacctt
     1321 gtctagcatg ctacattacg cgtacagtct atggaagata catcttatgg ctgagcgagc
     1381 caccagggat cttgcttctc tggccttcaa aaatggttag atatcgctgc atccaatcat
     1441 aatgggacga agacagtgta ttgaggtata ttttattttc cgccataggg cgggtcctgc
     1501 ctctattgca tgtgcttgca atgcatatgc ggccatcaga tgcaacggac gcatatcctg
     1561 ccttcagctc accaatacat gcatatattt ctgcacagcc cgctggattc cagacacctg
     1621 cgtgtcttcg gcccaatcgt ggaagccatc ggagctgatc cggctggaat attcagttct
     1681 catcctcggg ggctacttgc ttgaatgacc gaacccgaac acaacctaaa tgttcattct
     1741 atggtcgacg gtcaacatgt tattggccac atgcatgcat ggatctacac aataaaatcg
     1801 tacttataga ccttgaatat gagacatgca tatctttttg gttgctaatc cgaaacacgg
     1861 cccggaaccg ggatgatacc taaagtatag ggccaaacat ctaaatcata tacccacgaa
     1921 acagccctac ggctgttgcc attcgaagta tgtactctga atagtaaata ttccccgaat
     1981 acttcggtta aagaagggag aaaaaacacc ccgtagatag tcgtcagacc aaaattcgcc
     2041 gtagcgcaga gtg
//
```

#### *Talaromyces borbonicus* putative maleidride BGC

```
LOCUS       NBSA01000026           52167 bp    DNA     linear   PLN 05-FEB-2018
DEFINITION  Talaromyces borbonicus strain SV-2017a Contig0000026, whole genome
            shotgun sequence.
ACCESSION   NBSA01000026 REGION: 13874..66040
VERSION     NBSA01000026.1
DBLINK      BioProject: PRJNA379116
            BioSample: SAMN06579453
KEYWORDS    WGS.
SOURCE      Talaromyces borbonicus
  ORGANISM  Talaromyces borbonicus
            Eukaryota; Fungi; Dikarya; Ascomycota; Pezizomycotina;
            Eurotiomycetes; Eurotiomycetidae; Eurotiales; Trichocomaceae;
            Talaromyces.
REFERENCE   1  (bases 1 to 52167)
  AUTHORS   Varriale,S., Houbraken,J., Granchi,Z., Cerullo,G., Pepe,O.,
            Ventorino,V., Woeng,T.C.A., Meijer,M., De Vries,R. and Faraco,V.
  TITLE     Genome sequence of the novel fungal species Talaromyces borbonicus
  JOURNAL   Unpublished
REFERENCE   2  (bases 1 to 52167)
  AUTHORS   Varriale,S., Houbraken,J., Granchi,Z., Cerullo,G., Pepe,O.,
            Ventorino,V., Woeng,T.C.A., Meijer,M., De Vries,R. and Faraco,V.
  TITLE     Direct Submission
  JOURNAL   Submitted (03-APR-2017) Chemical Sciences, University of Naples
            'Federico II', via Cinthia 21, Naples 80126, Italy
COMMENT     ##Genome-Assembly-Data-START##
            Assembly Method        :: ABySS v. 1.3.7
            Genome Representation  :: Full
            Expected Final Version :: Yes
            Genome Coverage        :: 518.0x
            Sequencing Technology  :: Illumina HiSeq
            ##Genome-Assembly-Data-END##
FEATURES             Location/Qualifiers
     CDS             join(912..1677,1929..1970,2059..2396)
                     /gene="tbR7"
                     /note="Transcription factor"
     CDS             complement(2849..4159)
                     /gene="tbR6"
                     /note="Long-chain-fatty-acid-CoA ligase"
     CDS             join(7115..7270,7333..8022)
                     /gene="tbR5"
                     /note="AusD-like methyltransferase"
     CDS             complement(join(8441..8520,8577..8659,8733..9482,9539..9736,9805..10016))
                     /gene="tbR4"
                     /note="Alkylcitrate synthase"
     CDS             complement(join(10738..11345,11470..11536))
                     /gene="tbR3"
                     /note="Hydrolase"
     CDS
complement(join(12831..12984,13039..13211,13263..13607,13650..14125,14179..14268,14326..14548,14612
..14701))
                     /gene="tbR2"
                     /note="Cytochrome P450"
     CDS             join(15187..15517,15568..16709)
                     /gene="tbR1"
                     /note="Alkylcitrate dehydratase"
```

```
     CDS
join(17271..17489,17550..17636,17689..17746,17801..18140,18190..24139,24185..24502,24548..25384)
                     /gene="tbpks1"
                     /note="Highly reducing polyketide synthase"
     CDS             complement(join(26153..26940,26995..27073,27107..27673))
                     /gene="tbL1"
                     /note="Peroxisomal-CoA synthetase"
     CDS             join(28197..29945,29997..30032)
                     /gene="tbL2"
                     /note="2-hydroxyacyl-CoA lyase"
     CDS             30806..32083
                     /gene="tbL3"
                     /note="Phosphonomutase"
     CDS             complement(join(32404..32586,32643..32696,32772..33302,33364..34296))
                     /gene="tbL4"
                     /note="MFS transporter"
     CDS             35892..36746
                     /gene="tbL5"
                     /note="Enoyl CoA isomerase-like"
     CDS             complement(join(37239..38224,38278..38548,38667..39011))
                     /gene="tbL6"
                     /note="MFS transporter"
     CDS             join(40361..40778,40846..41039)
                     /gene="tbL7"
                     /note="Maleidride dimerising cyclase"
     CDS             complement(41281..42726)
                     /gene="tbL8"
                     /note="Acyltransferase"
     CDS
join(43549..43563,43621..43699,43754..43833,43902..44162,44226..44432,44485..44501,44564..44632,446
91..44703)
                     /gene="tbL9"
                     /note="Dienelactone hydrolase"
     CDS             complement(45287..46288)
                     /gene="tbL10"
                     /note="6-bladed beta propeller"
     CDS
complement(join(48027..49227,49284..49712,49782..49854,49917..50653,50735..50859,51272..51296,51531
..51700))
                     /gene="tbL11"
                     /note="Transcripton factor"
BASE COUNT    14628 a  11326 c  11919 g  14294 t
ORIGIN
        1 aagtgccagc ttctggctat caaactcgca atatcatacc gtacgcacgc tgcagaattg
       61 ttcctaccat tacggagggg ccaaataccg tagcagtttt tttggattcg gttaccgagt
      121 atgagaatat cgattgtatg taaatattac tgtttctgtt tagcagatag atgatcatgg
      181 cttgtactaa gtgtacgcga gtatattgtg tttcgcctcg gccccaaatc tctacggagc
      241 caaatcctgc cggtaaagtt acctcctgcc actatatgtt gctgacgaag tctcagccgg
      301 cttgataaaa ccagttgatt tcgtgggggc tattttccaa ctagttccct aaaagaagtc
      361 aaattcagtg gagaacactg aataagacaa atcaaaattg aattaatgtt aactacttta
      421 ctactagcct agaaggattc ttagcaacgt gttagatcta accgtgtcca gttgctgccg
      481 aatctgtact tagattacca acatgcattg gtttgtcgtt gaaatcctcg tatctcaacc
      541 atattcctca ttttgcatta gagtgaggtc ggaataggca gccactattc cttttggatc
      601 acaataatgt attttgataa gtgcgtaatt atgaaaaatg tgactacatt tgaggcttat
      661 cgtcagcatt taaaccacca acaaatacga gacactaaat ttcagccgtc tgcacgccga
      721 taccactgat cacgtctgaa cggcttatgt gtgaattatc ctggttgtcc agaaggtgac
      781 cagcttctgg ttccgcagta ccaatactta gactgccaag ttatataata taagaaaggt
      841 gccggttctt aagatggctt gattctaagc tttttttaat gtcagatcac gtctcctcac
      901 tacaattaag catggaggac aaggctccac agccaacggc accccagcgt cgccaagccc
      961 cgaagatacg actttcttgt gacaactgtt cctatgccaa ggtgcgatgc gaccaagagc
     1021 gaccgtcttg ccggcgttgt gtctctagta acgtctcatg cgtctatagc atttcacggc
     1081 gcatgggcaa accaccaaaa gatcgtcgaa tgaacaggac tggaaatgga gaaaatagta
     1141 ccagtcaaaa caacagaaat gggaatgacg ggaccaaaag aacaccacca cctagcaact
     1201 ccacgacagt cgcgccgact ggtgatgcat ccgacagctc atcgccaggg aaagtgacaa
     1261 atactagtgc agagggcgtt gtaggaccgc tgctcacgaa tcaagcgacc ctcgacagtc
     1321 tctatgtcga ttttggcata cccgacgttc gaacgacgaa cgggcctact tttgcttccg
     1381 accacctcct ttcaatttcg tctgatgttt ctttggaaga ccagaattac tttggatcgg
     1441 tggcatggcc aggtaatgat gcatccagaa ggatatcgca gagccagccc ttcggtgtca
     1501 cggtgccagc cattgacggc atcgaagcat ggaactcgag tgttaattcc aatttcgccc
     1561 tccatactgc acctacagac tctggtacaa gtagtgtggc gacatccgtc gactatccgt
     1621 atcacctact acatggatct actggcatac ggcgatcagt cttcgattcc actcagagtc
     1681 atgcttcaac gtgcggccag tttgtttcaa acttgcttca tagtctttcc ttaccaagca
     1741 attttgtac cacatttccc aatcccgtgc aaccaagaac cacagataga gttctagatg
```

1801 ccagtagaca ggcattagct gcagtgcaag tactgctaca atgctcatgt tcacatgaaa
1861 gcagttctgc aatctcaatt gcctccctga tcctgaagat tcttgattca tattgtgcta
1921 tctctcagag cctaccagtt tcttctgtga cgaaccaaac attgagctct gtgagtgatg
1981 ttaatgggca atctgttcct gcatcatctc cctcaaccga tactctcagt actaggtcta
2041 tgcaaaatat ggttctagat accccaatca cgtttggttc ttacaaaatt gatgcgggcg
2101 acgaacagcg attcattctc cagctactgt ggatggaact tcgcaaggtc ggtcgattgg
2161 ttgatgcttt caatgccaga tacgtgggca gcaacatgcc aacaaggaat cggaacagcg
2221 agaacatagc tcgccgtgct cgtgacagtg aattcccttc tcaaacctgg agtaacgaag
2281 aagaggcaat ttttattgcg ctagaacaat tcatgagggt caagatacaa tcgactcgtc
2341 gtgagattaa tatggcactg tcgagaagtg atgatacagc gatggaattt acaatttgag
2401 caacgtgaag actgcgctag agtatgcctt ttcttttgga tgatgtctta tgattctaga
2461 aacttgtatt tgggctttag aaatttgaaa tggtttttat tatagattcg actgccgtcc
2521 acggggtgtg aatagtttgg aagcgtccaa gatatgattt tgtttctgga aacgacaatt
2581 aaacgacaac aatatgaagt atagaaaaaa tcccttaaga gaaatttgcg taattcgagc
2641 cggtgaatat ggccccgaac taggatgaaa ctagaggcgt gccttcgccg attctttgcc
2701 tttggatgct gtcttttcat agagttggtt taggtgatct cgaaaattct ttcttgcttg
2761 tactctgcga agcttgaatc tagaaaatag atcaaagggg taagtatagc tgtaatattt
2821 catcgtccaa gggagtgtgg taagcttacg tagcagttaa aagctcgttg gcaactgtaa
2881 aaggctcgat ctccaaatga atagcccgca cgatctcgta gctatttagt ccctttgctt
2941 ttcctatctc acatagaaga tcaagcaaag ctcccacaac tttttcatct ttcatagctt
3001 tttttatggc agccgtatta tccgcattta tatttgagcc taggattctt cctgcatatg
3061 aagcaaacgg cgcggggtcg attccgagga tagcaacggg aaaactttgc gaaggatctc
3121 cgtgtacgaa ggcctgagtc aatagactgc agtttgccaa ataaacgttt tcgatccttt
3181 ccggtgaaat atattcgcct tgagccaatt taaccacatt ctttttgcga tcgatgattc
3241 ggaaacgacc aaggttgtca acctcggcga tatccccggt gtggaaccag ccatcatggt
3301 ctatcaccgt cttggtatca tgaggctccc cataatactc acgaaataat gttgttccac
3361 gtagcaaaag ttcaccacgg ggaaatggct tgtcagtcac attgtaatcc atatcgggta
3421 ctgactgaag acaggcctca cccgagggca agactcctcc gcagttacct gtcgactcgt
3481 ccgagcccat ctgagccata gctatggcgt atgtttcagt aaggccatag ccttgaatga
3541 actcattatt gaagacatgt cgtagtagct gctgtaaatc agggtctaag ggtgcagctg
3601 cactcatcat tcgctttgct ttctgcaggc ccagttctct cacaatcttt gacgaaaatt
3661 ctgatttgta cgagttgcgt cttctgtctg tcgcatagtt gcttgagtgg ccgaccgtgg
3721 tctctgtgtt cttagtacca gcagcatttg tctgaggagt tgaagacctc tcagcctgtc
3781 tcacaagggc accaaagcga ttatatatgc gaggaacgga gttgaaccct gtaggctgta
3841 gaactttcat atcatccgcc agtgcagaga catctccgtg aaagtatcca atagaagcac
3901 cggacataag cgcacactgt tccaccacac gttcataaat gtgggcgaga ggaagatatg
3961 agaagatgac gtccgaattc gtgatgccga aaataagcct tgcaaccgag gcggctgcaa
4021 ctgcgttctc gtgtgtcagc acaacaccct ttggatcacc cgttgtccca gaagtgtaat
4081 tgatagttgc gatatcgcta ggttttggct cgcaaaatga tatatgcgat tgtgcgccaa
4141 tagtctccac ttccgccatg tgaaatatct caatactgga ctctggagca gacttggcta
4201 ccttggactg ccgacttagt ccttctgtat gttcaacctg taacgaatct atcgaaacga
4261 tgatcttcaa ttgtggacaa tgagatgaaa tttcgagcaa ggaaggaatg tgctcagccg
4321 atgtcacgac aatagggatt gctgtcttgt tgatgataaa ttttgtagtt ccaggcccta
4381 aagattcata cagcgacaca gaaaacaaag actgtgacgt gcagcccaag tctatgtttt
4441 cataagcaca ctcgcatttc cgcaatttca gcttcaggaa agcttaccaa caatctgcca
4501 ctcgggacga ttttgacacc agatgccaac cccaaaattc tgcgcagata ttccatgttt
4561 ttgacataat gcacgaatac cagccccaat attcctcctc ctttcttcca cctctttgta
4621 agtcatccat tcatatggac cccagcaagc ggtagctgta ttccatttac gttgacccag
4681 acatttagca tctggcttat tctgcagcga caatataaaa agatcgtgca gtgtttgtac
4741 tcgagaatca agcgttttca ggagcggagc attggtatac ctccaatggc gatagatggc
4801 gctacggcca cacctcgtgt cttcactcag aggcagtttt gctgagtatg gctgtccagg
4861 tggaggtggt tgttgtaaaa gctgaaaaag cttttctgga cttaaggatc gatgcagcaa
4921 gtccatatca tccattcaca ctcgcaaatg acacggtaga aggtcaaatt atgcacagca
4981 gattggagga gagagtcgag agcaatatgc aatatgcaat atgatttatt aaaagggttg
5041 tgatccaata ttcatgaagt tccccttttgc atgcaaatcg tgccttggta gttttgagct
5101 attcttaaaa acaggcttta taaattgata gtgaatctga aatctcttcg ataatgtaat
5161 actcgtcaac gtagcacccc ttctagaagt cttctagaaa ttgcaaaatc aacttaatta
5221 tatggtacct aatataaaag gtttctcaac tataaaactc tgctaatgat ttgatggccg
5281 acacacattg atcccgaaaa tacagtcaaa aattgagtac cgaaatgaga cttgccgcca
5341 aggcacttcg ctatccggat tacataactt gcacatgact agttaactta ccgagcataa
5401 cggtaaacaa gaatttaagt agctgatttg cttcatagat acattgaagg aaaaaagctc
5461 agtagagagg ctgtccaaca tctagtttaa caaaaaaaaa aaaaaaaaac atgcttgcag
5521 aactgtttag accctgacca gatgaagtaa taccgattta gaataaatta aacgatatag
5581 cttcaatagc agcttacaca catgaagagt agtttccatg cttgggaatg aaatatggtc
5641 atttaacatg aaccatgatt agtacgcgag aggctccgca gactggatgt aaatcacttc
5701 gtggctagga agtgataggt gtgcagcatt ttccttggga cactcagcca attgactacg
5761 aactcgtctc gtgtttttcg taaatgaagg aaatggctga tgctagcagg cagccagctt
5821 cgcaaaaccc tctgtgtatg gcgcaattta cgtgaaagga tataccagat gctgattgga
5881 acgatcggtt atcgaatcgg tcatcgaatc ggtcatcgaa aggattgtct gaatgcaatt
5941 gattctgaag cttggtcgac cggttccggg ctcacagcga caaacttgtg ttatctaaat
6001 tcagccctgc tccggaaatc cggggcccac atccatacga gggccggtga cggcccgttg
6061 tgcacgggta aaactttaat ttttatatag tgggtccaag taagacacgt accatattca
6121 atacgacatg cgtatgtgta tgtaatacgc ggctccagga aaattaccac attagcgtat

6181 tagcgctcat acttcctgaa cagttcaagt gtgaactttg aatgcttcag acttgactgt
6241 actcaactaa agttgaactt gcggtctcat ctgttggttt catcaatgtt cttcagctcg
6301 ccgagccgaa aatcaagcaa aagtgatgat cacttggtat gtcgagccca actaccgtgc
6361 gtttaacagt ccgcacatgc atcacaattt cgtttccgac gccgcgggat tcgtctaccg
6421 tttgaaggtc atggctttgt gtcatcagaa tcttacagta tttattttga ccccaaccat
6481 tgtttatcat ttatagcgag agctctttga gattttacat gttgtaccta ggtagccgag
6541 cgaattacta gaaaatatca ctaaataaga gactaatgcg actgcacttc aagaataacg
6601 agcaccgatg caatagaatt aacatactaa aataggtcca tcttggagat actttattca
6661 tcgggtattg agttatgaat ttggaatctt caatctgtgc tggcaatgca cacgtatatg
6721 gctgtgtact atttgtccaa tttttctatc taaaaaatgc tcgtaatcga acaatttatc
6781 atgatcatct cgagttggac aaattttccg atgcaataca ttcatttgac ctgagatgat
6841 gatttctaag acaccgaagc tgtaaaagat agtccacggg tatttgactg gattggattt
6901 tcaccatttt cgctgtccgg gaactatccg ggcttcagtg acattcgcag agggtggaac
6961 tttctcttta gaatatcggt cgctttcaat tctataaaca caatgctcta gtatggaagc
7021 tgatcctatc taatattgct tcaattcccc gtcaattcat atgtagagtc tgagccatta
7081 tttctgtcag tactacaata agcaataaga cgacatggca cgttcaagta ccgaacaaca
7141 gcagcggaag gtggatggct cagagtggtt tgctgaagaa attcctcgtg ttcccttagc
7201 aatgcgagaa ttgcttgagg aatactccgg tgtttctgag gatcgtgtgc tgccgcatat
7261 tttagaaatc gtgagactga tccaaaaaaa agcttcctga gttaaaccac ctaatatgat
7321 cttatttgat agcgccacaa agctttttca ttattccctt atccgtgcat tggacaattt
7381 cggttcatgg atctgaagct atcagagaac gctctctacc gcactgttct ttcacgacta
7441 aagagtggta atgaaaattt cctcgacgcc ggttgctgct tcggccagga gctccgtaag
7501 ctcgcttttg atggtgtacc aacacgagcc ttgcacggta ttgatctgga gcctggattc
7561 tgggatctag gatacgagct attcggagac cagaagcgaa tgaatgatgc gtctttcttt
7621 gcttgcaact tgttggacca aaccactatt ccccagctga cagggaaaat tgacattctt
7681 ttggcaaata gcttgttgca tctatttacc tgggacgatc agatcaaggc tgcttgcaaa
7741 ttggtcacat ttatgaaagc taagaaaaat tcattgatca ttggaaggca ggttggtgcc
7801 gttaatgcag gcgaatatcg aggactcagc gaagacagca caacgtttcg gcacaatgtc
7861 gagagctttc agcgtttgtg ggacatcgtt ggtgagaaga ctatgtcaaa gtggaatgtg
7921 gaagctactt tagacatgaa agacattgtc gatggtgtga atctgggtca aaaatggatg
7981 gagaacggaa ctcggcgttt acactttgta gtgacaaggc agtaaatagc ataatgcatg
8041 gatgtatgac ttggaagtgg gaatgaagat taagtcattt atattcatct tgttttgtca
8101 catatcttaa ttcttaaacc ctataaagta agacgaaccg gcgaagtagc tatgagaaaa
8161 tgtttttcac aattttcacc atcatcaatg ttcgctaaat atcacagtca atatttatgt
8221 ctgtctaatg gttaatcaaa tatatgaagc agtcttactg gaaagattgt tacttgataa
8281 gatttagagt cattttaaat tttacttgca tacacaacct caatggaatg tacaccgaac
8341 aaataaatgt tctattcatg gatcaataaa gcgaagaatt tgataaattc acatgaaatg
8401 aaaataaact aaaaattgtt tcgtctctac ctaggtatca gagctttgaa ggcaagtctg
8461 aagcttgttc ttttgttcca gtatagatgt gggtagggcg aataactttc atatctcgaa
8521 ctgttgtttt ggttaattaa tgaacaaata atgaattgct agaaagtatg acccactcat
8581 tgcatcgcgc cagtgaccta gtattcctgg aaggcgttgt gtcaacatca tgatgggaat
8641 gaactcggca tcccagcccc ttgataatat tagttcgact tccatatagt caactaaatt
8701 catatgaaaa ttcagaaaga gccgatacat acatagcaac gtaaaagaat actccataga
8761 gatcagcatt tgcgttcaat cccttggctt taaaccactc atccgtagaa gcctttcggt
8821 cgatctcctg tgctactgcg aggagagggt tggatgcggc gtccaattgt tcgagtagaa
8881 tctggatggg cttgattcga ggatcaactg tactgtatgt tcgatgcccg tatccaaaca
8941 atcttcgctc tcccctcttg acttcggcaa taagatcgtc gactttgtcc gggctaccaa
9001 gtttggtcat tgttttatat gctgtttcgg gagcaccaaa gtgcaacggg ccataggctg
9061 ctgctaaggc actgataaga ccagaaatcg gatccgccag agttgaagct gttaccagca
9121 tggcaaacgt ggaattgctt agtccgtgat caatactcaa tgctccgaag cggcggaaag
9181 aacgaagttt gattctgttt gggcgcccag tcacaggatc aacatggccc atcatattga
9241 ataaattctc ataaaacgtg ttttcgtggg ttgcagggac aaactcagtc ccgcggcggt
9301 gacttgcaac aagcccaaca actatagccc aggcagctat agatcgaaca acgcctttat
9361 cgatcatgtc ctcattgcgg tgataaatgt tgcctcccct taccgacggt attgtgtctg
9421 gatacgaggc aacgtacgct gataggccag caaggatcat agtgactggc ggagaagaac
9481 gactttgggt ggattgatta gctttcacaa actcacttca atatgtatat ggacttactc
9541 aaaagatctg atgactttca caacggtact cgggatatcg atcattgcag tcaccagggc
9601 tttacgtaaa gccgtttttt gcgaagagga tggaagcttt tcccagacca tgagatgcag
9661 aatatcctcg aaatcgcagt cccatagttg cccgatgctg tagtttctga ataaaagagt
9721 accggtctcg ccatctctaa tcacacatca atcccaagtg aatcccgaaa agtagaccac
9781 ctgtaaaaca ccgaggcgac ttacatgtgt atcaatttcg attcaatcgt ggccgtattg
9841 gcaagacccg gatcatagac cttcaatcca acacttccgc gatcaattcg agttgtgtct
9901 gtagcaggac ctttaatctg cttgaatgca gttgccggta ccgaatttcg cgaaatggga
9961 atctcatact ctcgctctgt gcgcgagtct ttgacgaata aagtgccttc ggacatcttt
10021 tgcctcccaa cactgtgttt cgagttttca ctttcggtcg tcttatgcga aatccaagaa
10081 gcaaaggagc ttctacactg agtcgaaaaa tacgtaggta ttggtcaacc ttggaagtag
10141 gttgcgatgg atctcgatat attcctagaa atgtggacgt ttgatccaaa ccgagtgtaa
10201 ttaaatctta ttagatctaa ggctatatac acatgaatgc agcttgcccg ccgaaataag
10261 tgacagttgg cttagccaac tgctttgggg aaaaaatgcc cccaactcgc tgtcgcctgg
10321 tgcgcagaaa tttctctgga atgaaaaagt gattgccgag gtgactaaaa caacagcaat
10381 aacaattaat gtcaaataac ttactccatc gcgccccaca ttgatggacg gctgctaata
10441 aaaccagcaa cctatgcata ctgcttaaag gtcaattctt gaataataca gcaccccaaa
10501 cagtaaaccg gctttgactg tgatctgatt cgaaatcaat ttcattttca tgtttcggtg

10561 atcgtgatcc cttccaagcc tatgggaaat ctcaggtttc agcctttcga ttgctggata
10621 gcttgaatgg agattctcat tcggtagccg aaaataatac tttcagggat aaaaggcatc
10681 atttctattc agctctagaa ctaagtttcc cgccaactat accggcttcc ttagtagcta
10741 catgaaattc actgcgtgga tcatctcttt gattgtcccg acaattccat tcacgttttt
10801 ggcgtctttg ggatctcat ggcccttatc atgcgtccat agcttcgtcc agcttttgga
10861 gcaaagagca aaaagcttga gagaatgttg atacacaaaa tccatttttc cagcaatatg
10921 aacagtcggt atccgaacca tgttatccag atcttggtcg taaacaaaag atttggtgtc
10981 attgatacga aatggaggca ttgcgccaat aaacacgccg catcgaaaaa gggtgtctgg
11041 cggatcgaaa ggatttcggc gtgcatgctg ggcaagaaac ccataagcaa gggtagcacc
11101 atgcgaaaat ccaattaccc catcaaatgg gccttcttcc tcgatgattt catacagtag
11161 acgataagcc tccaaaactg tttcttcgtc tgcagggttg aaagagcgtg gccatgtgta
11221 atagcttaag tatggccctt caaagaaacc ttggattccg ggacctggtg tggtagattc
11281 cagttcgccg tcaatgaatt caaatgaggc gatgccatca ccttcaagtt tccgtatgat
11341 cggacctaat atcaaaaata gtcagaattg gttgccagta atggggaact taaagcttga
11401 gtaccaagct gtgtccgaaa aatctggata tgcgttagcg aaatcaaatt ttcggtacct
11461 aggcatcacg ggacagtagg catacatcgg aattagtgcc tgagccatgc agacaaagaa
11521 atctaaaagt aggcataatg ataaagaaat ctaaggaaat gaactatatt tttccctggt
11581 gaatactggt gtaatagttt atagtggtat gtattatacg ctcaacacgg tgcttcttca
11641 agttcccttt atagaattcc tcgcttgtcc caatcagtat caccacccat catcggaatt
11701 tgcaatcaga gaagaattga cttgaatcta tcgggaaagt cagagaccgc ttatctgtaa
11761 agttgtatcc actcacagtt cgaagaagcg ttttatttgt aaccctccac ccgttggggg
11821 ttaaaatcaa gtcatcaagg taccttttag tcaatgagct cagacccagg gtcgtattcg
11881 gggagtataa tgactcaccg gccgtacgta gtagagactc gtatttcgta tccttcatcg
11941 ctgaaaaaag agccttctag ataggagacc gaagtggcac ttgtggggcc agtaatctca
12001 atcagctgtg tcgataaggc atgatgcgtt ttcctccctc tgagaccatc tgctagattt
12061 tctgtgatgt tctctagtcc ttgcaggatt cccttggcgt cgttgaaatt gcaaaaggca
12121 ttaggggcga agtgtaaact gagaagactg agattttgag tgtccacagc atggctaaag
12181 acgttcaacg tttgctctat gcttttgacg tgaaaaatcg agagatcgga cccgaagaaa
12241 cccaaaattc gaattagggt agacaattgc atctttgatc cttgacttcc gaaactcaat
12301 gatcggtggg cttgctgaga tattcaaata gcagtggtgg gcgaacaggg gcttttagat
12361 tccaggattc gcgcaaaaca agagacgaaa ttcttgggcc gattgaaaga aaatatttac
12421 cgatggtgtt taagactcag cgagtaccaa cacgcactac ttgtgtgctt gcttcttctg
12481 gaactactgc ctcctttagc agatcccaat cgtgacggcc cggggttact cgggaaagct
12541 aactagaaga gtacgatttt tcaacgtcac gaaaatatac caggagtaga tttcagagaa
12601 tctctgcgaa acaacgaagg aatttgaaaa tgacctcaat aagaactcta tttcacgaat
12661 ttagtaacgt atctgagtgc agcgtacgtc tttttggggt gcctgattag atatgattcg
12721 tatcctttgg aatataaatc aaaaccacga atcttgctcc aaacaacttg aatctaatat
12781 cagtcaatcg acagaaccag ctttcgcgcc tcggatagaa tgttaatcta ctgtttgcgc
12841 ggcgacagcc tgaccataag tggtttcatt ttccagaaca aatcagcagg gacatcagac
12901 cagttgtcat cgcgcgggta aagttccata tcaaaatata ggatcagccg actgagggta
12961 tgacgcatct gcatgtaggc cattctaaat tgcaaattgt gagactttga tagtagaatg
13021 aggagcaagg atacccacgg cttgccaata cagtttcttg ggccagttga aaaaggccag
13081 acagcatcca aagcatcatt ccggaattct tcgggtggat tctcgagcca tctttcaggt
13141 acaaagtcga gaggtcggtg gaaattctgg ggagaatgaa acgcagcgag ttgatggacc
13201 ccgacggaag tctaaaagct tgtttaattt tgcgccaaat tctgggaatc tgacatactt
13261 acatccgccg gaatgtagtg cccgtcaatt gttgcacctt gatgacctgt tcgcctagga
13321 aacttacagg gggcgggtgg gaaaaccctt aaaccttcct ggataactgc gttcatgtaa
13381 ggcatggcat tgcatttagc gttggatata tcagatgcag actgaaaatt cgcgtcaact
13441 tctgctttca atttattaaa actttccggg tgagtaagaa ggagaaatgt tatgccgctc
13501 aagccagttg ctgtaggctc agttcctcca agcagcatga gaataagtgt ctgagcaatt
13561 tcagacctcg tcatcccctt ttcgctattc ttcttgagaa tctgattctg tcagcttaaa
13621 gtagcgccga agtgacgggc tcaacatact ggattcacaa tgtcagctct gtcgagctgc
13681 atgtttagcc tcttctcagt cttctcgatc gagtattgca tatgtttctt tcttgctgtg
13741 ttgatttcag ttgttccgct ggtcaaatat ttggcagcct ttgccagaaa gtgtaagctc
13801 gggtaagctc ttagtaatcg aatagttccg ccccaactgg cgccctccaa aattgcatcc
13861 atccaaggat ggaatgagcc ggtctcaacg gcattgaaat tttcagcaaa acagagatca
13921 gccataatat cgaatgtcag acaagagtac cagtgtgtca tttcgttgac accgttcttt
13981 ttcggatcgg agatactttc tcgaagtttc gacatcagta gttcaacgta ttgcacaact
14041 aatccctctt gggcccgaag agctgcatct gagaaagagt gaatcacaag ttttcgaatc
14101 cgtacatgat cttcatggtt ggaaactaga cgcttgttag ctctctggtg tatctggtat
14161 atataacgaa aacattacct agaaggctgg gtgatccgtt ggtgttctgc gtgtagaagg
14221 taaagtcttt ttctaattct ggctttccag ctttggcagt atatatgtct aacccatatt
14281 tagcttttca taatggaaat tgactgagct cgtaaacact catacctttc catgcctgtg
14341 cagtgttgaa tgaaagaaca tctggggcaa gtcgtacaac tggaccatat ttctcatgga
14401 atcttttggc attctgatgt tgggtgtgtc gaatttcatc caggtaatta aatatggtaa
14461 aggcgcctcg gctgtagggt ccaggaaatt tgcgtagtgg atgcagataa atgttataaa
14521 aagctttgaa aaatatggaa catattgtct accgaaatgt tagtgacaaa cgattccaag
14581 gttcttgaga taaacagttt cgttgacgaa caagtccagc aacagcagca acaactgcaa
14641 atggattttc tttcgaaaag agtaacacgt tactcaggat gtctttaatt gagagggcca
14701 tgttacagga tgaaatctta atcgacaaaa gactcaacag actcgttttt gatggacaca
14761 agaagatttc atactttgag ccaagatact ttcttcaaag cttctcgtcg accttgccaa
14821 caaaattagt gaaacagtcc ggtgcatcat caaataagta tgacttgatt tacaaatgtg
14881 tcaatcatca ttgcgggcat agcaactggg cataatcggt ttccgaactc aaatcgcaag

14941 caaacggcct ccgaattgta ataacaatcc gccgtctttg tctcttcaag gcatggcccg
15001 agtctcactc ttaccttaaa atttggcaca ttacgttgaa tagtgaattg ggtctatcgc
15061 aaagttcgct atatcagaat cactgagcaa agcagccctt aaaatctgcc tagcgttttc
15121 aacgacttgc gtgaactttg gagctttctt tcattttaaa tttgattact tgacatctac
15181 agcatcatga cggggaatat tcagtacgat caagttttga ttgacatcaa agactatgtg
15241 tttcattaca aaatccagag ccctcacgcc tggacatgct cccgtaccgc acttcttgat
15301 tctattggtt gtgcaataga gtctatccac aagtccgctg aactccgcca gatgctaggg
15361 cccatagaca caggtgcgac tattccctgc ggatttcggc tacctgggac tgcataccag
15421 cttgatcctc tgaaaggggc tttcgatttt ggagctgcaa tacgcttcct ggaccacaac
15481 gataccatgg gaggtgccga ctggggccac ccatcaggta aatagtacat attatactta
15541 attactctgc gtctaatggg gttctagata acctgggcgc tctgctatcg gtgtccgatt
15601 ggctttgtcg atctactatt tctggaaagc cggctcgaac ttctggtcca ccgttgaccg
15661 tcagaacgtt tttggaagca atgatcaagg cttatgaaat tcaaggctgt tttctcatta
15721 aaaatgcttt caatgctttg ggattcgatc atgtgatcct agtgaaactc gcctcaacgg
15781 ctgtagtttc ttggcttatg ggattaagtg aactacaaac tctatctgct cttagtcacg
15841 tttggatgga tggtgctgct ttgagaatat atcgttctgg ctcaaatacg attccccgaa
15901 aagggtgggc ggctggagat gcttgtatgc gagctgttca cctggctctt ttagctcgtg
15961 ctggtcaagt tggcgccccg acagctctga ccatgccgcg atggggattc tatgacagca
16021 tgtggcaagg tctacaattc atattgcctc aaccatatgg gacatggtgc gtggagcatc
16081 tatttttcaa agtcatgcct gtagaaggac atagcatctc tggcattgaa gcggtattga
16141 tacagcgtgc caagatgctc gatcgaaaaa ttgatcctct gagagacatt gaaagaattg
16201 aggtccgtac caatggtgct gcaaatctga tcatcaataa aacaggaaaa ttacataatg
16261 cggcagatcg cgatcattgt atgcaatact cgcttgccgt gaccttgctg aaagggggcg
16321 ttccatcacc agaggactat atggatagta gcccttatgc atcaagtcca gccgtggaga
16381 gactaagggc ctgcattcac gtgagagagg acgaaaattt taccaaggat tatctggaca
16441 tcaataaaaa aagtgtccca tcggctgtta caattatctt gaaagatgga tctactatgg
16501 acgaggtgaa aatagaatat ccggttgggc actgtaagaa cagggccaca tctggggagg
16561 tagaaaagaa atttctgagg aacatgggtc tgatgtttag tgccgatgag ataagacgta
16621 ttgttgaggc tgtagaggga gaggacgatt tgctgatctc caaactggtg gatctgctgg
16681 cacgacctca gaatcattca ccgagattgt gatgtacgta aagcaaagaa caattggcta
16741 ttaacaattt ttaatggaag ttcatctgtc attcgaggca cttggaaatg tgaatcataa
16801 gatctgacaa tgacggcata tcttggaatg ttgaagaaga tgagataaga acagattagg
16861 ctcttttctt cttttgaata gccaatttcg ttgcctgcaa aacaattcct atcgtatatg
16921 ttgatatact agtatgcata ttccgtccat gaatggcatg tcgaataatc aaaattccta
16981 aagaaatata aatcagattt catctttggt gaattgttcg gttaccgaag gagaatcttt
17041 tgtatgcata agtttctaat gcttgggggc gataatacga gaggatttat ttctttggta
17101 aggaatgaat gattatgata ataaagtcag catctctgaa gtgataaaaa tgaatcggta
17161 gccttcggtc cttctaggtt caatctattt ctcaaataac tatatgcttc gaacaatatt
17221 tggttggaac ccgaatctaa agtatagagg ttgctaaaat cttcgaaatt atgggatctt
17281 acgctctttc aaacactgat ggagtagtca aaaatgaggt tatgcctata gccgtggtag
17341 gtataggatt ccgtgggcca ggcgatgcaa ccgacgtgga gaagttctgg aacatgatat
17401 gtgaagctcg tgaagctcga actgtggttc caaaggaaaa atggaataac gaagcgtttt
17461 atcatcctga ctctaaccga aatggaacgg taagcaaaga cactccggct cctgaaattc
17521 agttgactaa gtaatttgac aaaatgtagt cgaatgtctt ggcaggtcat tacttcaaag
17581 atgaccttgc gaaatttgac gcaccttttt ttaatatgac aaatgctgaa gccgaggttt
17641 gccaaatgat aaatcttatc atatgttatt ttaaactaat aagcatagtc actcgatccc
17701 cagcagcggt tgctgctcga gtgtacctat gaagctctcg agaatggtca gtattccaat
17761 tattccatgt acatttcaga tattaactca aaaaaaccag ctggagtacc catggataag
17821 gccacaggga gcaagacttc tgtgtttgta gggtctttct gtggtgatta tacggatatt
17881 attatgcgag atccagaaac tgttccgctc tatcaagcga ccagtagtgg tcactcaagg
17941 gctatcatct caaatcgtct ttcgtatttt tttgacttca gtgggcctag tgtcactatt
18001 gatactgcat gctcgtcaag tctagttgca ctccacctcg catgccagag cttgaggact
18061 ggtgaatctg aacaggcagt ggttgcaggt gcaaacgtga tcctgagcca tgaaatgact
18121 atatcaatgt ccatgatgag gtaaatattc tcaaaggtcg ccaaatttct tcattctgat
18181 tttatgcaga tttctttcgc ccgatggacg ctgctatacc tttgacgaca gagccaacgg
18241 atactcgcgt ggagaaggag tagcatcgct tattctgaaa ccactgcaca aagccttaga
18301 ggatggcgat accgtacgag ccgttatagt caactccggc gtcaatcagg acggccgaac
18361 caatgggata actctcccca gtcgtcaagc tcaagaatct ctgattgaga gcttgtacac
18421 ccaagcagga attgatcccg cagaaacttc ttttgttgag tgtcacggaa ccggtactcc
18481 agcaggcgat cctctagaaa caagagctat ctcagaggtc attggaagaa agagacccaa
18541 agaccagccc gtgcgaatag gatcggttaa aacaaatgtt gggcatctgg aaggtgcgag
18601 tggcgttgct ggtgtcatca aggccattct catgctagaa aatgaagtta tacttccgaa
18661 cagaaatttt gaaaggggaa atccgaatat tccattctct gaatggaatc ttcatgtacc
18721 aacggccact gaacgctggg aatccgcagg tccacgccga gtttccatta atagcttcgg
18781 ctatggtggt accaactcgc atgcaatttt ggagcaagcc actagctttc tccagtcacg
18841 aggactctcc ggaaaaacaa tgaaaacatt ttcactcaca gaccgtcgca ttagctctca
18901 ggagtacaac ggaataaacg gtcataacac cgaagaattg aatgcccaaa atggcacttt
18961 tcaatcgaat ggaactacaa atgggatgag caacggaact catcgggaaa acggtttaaa
19021 gagatctacg atcaaacaac cttctcgtgt ttttaccttg tcggcttttg atgaaggggc
19081 tggaaaatcc caagctcagc gattgagtca atacattact gatagattgg gccttgcaga
19141 cgaccagttc atggacgact tagcctacac cctcggagaa agacgctcaa aattgccata
19201 tcgattcgca attggaacat cctcggctga agggcttatc aagtctctaa gagacccaag
19261 tatctcgttc tcaaagtcaa aaggtgtccc tggattagca ttcgtcttta ctggtcaagg

19321 tgctcagtgg tatgccatgg gccgtgaact caatgacact gaacctgttt atcgcaattc
19381 actcgccaag attggaaaat atctacaaag ccttggagca gactggaata ttttcgagga
19441 attgtcgaga gatgagaaca cctcacaagt atctgttctg agtcaaccac tttgctccgc
19501 gatccagatt gcactggttg atctcctcgc ttcttggaac attaaaccga ttgccgtcac
19561 aggacattcg tccggtgaaa ttgccgccgc ttattgtgct ggtgctttgt ctgcagagga
19621 tgctatgtcc gctgcctatt ttcgaggtat atactccaat gagctaaaga gctcaggtaa
19681 ggtagaaggg ggtatggctg ccgtaggaat gaatcgcgag gtagcacttc ctattttacg
19741 tgacttgaaa caaggcaaag ccaccgttgc ttgcgaaaat agtccgcaaa gcattacagt
19801 atctggcgac gtcgcagcga ttgccgagct tgagactatc atgaaggatc aaggaacttt
19861 cttccgaaag cttccagtat cggttgctta tcattctttt catatggcgc acgtcgctga
19921 cgagtacctg gaagcaattt cccatatcaa ggtaaaggca gaagataaag gcgttgaatt
19981 tttctcctct gttacaggga gtcgggcgga actgtccgat ttaggaccct catattgggt
20041 tcgcaacttg cttggagaag ttaaattctc ggattctcta cgaaatcttt gtcttgaggt
20101 caacacaagc agaaagagtc gacgacgcaa agagaaatct gcgatcaata ctattattga
20161 gctgggaccg cattctgcac tggcaggacc aatcaagcaa attatccaag gagatcaacg
20221 tctgttggaa aaatccattc agtaccaatc tgctctcata cgtaagtcaa gcgccactga
20281 tacagtatac gctctggcag ccaagctttg ggcatccggt tatccagtgg atattgctgc
20341 gtgtaacaaa cgggcaatac agacgcattc gaaagtattg gttgatcttc ccccatactc
20401 atggaatcat ataaattcat actgggcaga gtccaggatc agcaaggctt tcagaaatag
20461 aaattatccc aggaccgaca ttctcggagc gcctgacaaa ggagcaaacc cacttgagcc
20521 gagatggcgc aatgttgtca gggcttctga gattccatgg gtgaaagacc acaaagtgca
20581 aacaaacgta gtctatcctg ccgctggata ccttgtcatg gcaattgaag ctgcacatca
20641 gcgagctatt gagcgaggat cggcggtcac gagctacaag ttgcgcgagg taacaattgg
20701 tcaagcacta gttgttccgg agcagtctgg ggaagttgaa acactcataa gtctcagacc
20761 tttctctgaa ggaacacgtc agtcatcaga tatctgggac gagttttacg tctactccgt
20821 cacggaagat gaacggtgga ctgagcattg tcacggttta gtcagtgtac aaaggggctt
20881 tcaacctaat gacgttacag gaaatgctca aatccttgct gaatctcaag agcatctcga
20941 agctatttct aaaatagatt caaactcgac atctgccgtt gagatcaaga ccttttatca
21001 aaatctgacg agcattgggc tggagtatgg cgctaccttc gcaaatttga ttgccgcgaa
21061 ggcaggtcca aatacgtgtg ttggaacaat agaaattcct gatacagccg catgtatgcc
21121 catgggattt caatacccgg ttattataca cccggcgact cttgacagca tgtttcacgg
21181 cctctttgcg gctcttgcag cagaccaggg tgacttgcaa gatcccatgg tgccaatatt
21241 ccttgacgaa cttttcgttg attcaaggat caccactaca cccaaagatc agctaaaggt
21301 gtacacatca acagaacgaa aagatggaag gcaggctatc gcaactgtcc gagtcacagg
21361 gcataccaca gaaaattctc gccccctcgt tacgatttct ggtctgactt gtatgaagtt
21421 ggccaatgat gcagtacaga atattgagag tgaagtcaag cctattgcat acaacatcaa
21481 atgggagcct gacgtcgact tgctttcctc ttcagatatc aacaacctgt gtgctgacat
21541 tgtaccacca cagcacgagg ccaagagaat acgtggacaa gaacaagctg gctactactt
21601 tatgaaggaa gctttagcaa acatgtcttc ggaaaaaatc aagaacatga tcccttatca
21661 caagagactg tgggcttgca tgcatgcgca catttcagct gtggaagagg gcaggctcgg
21721 tattccgaca gagtcgtgga caacgtgtag tctatccgag agagaaaatc tcataagtga
21781 aattagcaat tcaggtgctg aaggtgcgct actttgtcat gttgggaagt atttgccagc
21841 tatactatcc cgtgaagtag aggcgctgcc tcttatgatt gaggacggcc gtctcgatgg
21901 atactatagg gacaaccatc gatttgatcg caattatcaa gcagctgcaa ggtacatcaa
21961 cctcctaggg cacaaaaatc cgcatttgaa tattttggaa atagggggctg gtacaggtgg
22021 tgctaccttg ccactacttc aggccctagg aggaaccgaa ggcgaccttc caaggttcaa
22081 gaatttccat tttactgata tcagtagcgg atttttcgat gccgccaaag agaaactttc
22141 tgtctggtca aatctgataa cgtatggaaa gcttgacatt gaaaaggatc caggatcaca
22201 gggttacgag ctaggtacat acgacgtagt tgtcgcagca aatgttctac acgcaactaa
22261 atcaatgcac aatacaatga gcaatgtccg aaagcttctc aaacctggcg gcaagctaat
22321 cttggttgaa ttgacacgag aacggatgac cacatcaact attttcggta ctcttccggg
22381 atggtgggct ggagaagaag atggtagaac taaaggtcca acattgaccg aaacagaatg
22441 ggtttcagtt ttacagaacg ctggcttctc gggcgtcgat gcagctgtct gggactcgcc
22501 aaccgagttt gagcatcaag gatcaatgat ggtagccgaa gcagttgggt ccgaagatgc
22561 caaagaaagt gatgaggttc ttttgatcac ggggcaacgc agttcggacc taccaatagc
22621 catactttcc gaaaaattag tcgaagcgaa aattccagca aaggctgaag atctgatctc
22681 agtgaacccg gcaggcaagc tttgcatagt gtttgccgag gttatacaat ctattctggc
22741 tgatccaact tcaaatcaat tcgaggccgt caaaagaatt ttgacaaccg cgtcgggcat
22801 attatggatt gtccaaggag cttcatcgtc ccatcccgac aagaatctga ttactggctt
22861 tgccagaacg gttagatcag aatacggcag tatcagagcc gtcatcttag acatcgatga
22921 agaagaaggc ttcacacctt caacaatgtc gaagatattc gatcttttcc gcctccagtt
22981 tctggattct gcgccaccag agaaggtggt agatgtggaa tattcgctgc atgaaggaag
23041 attgcggata ccacgtctaa ttgaagacac gaaaacaaat agattcgtca attctgctgt
23101 ttccgaaaag gttcctgaac tccagccatt caatcaacat ggccgaccac tcgtgattga
23161 tgtggccaca cccggcctat tggatactat cagattcgtt gatgacactc gcttagttga
23221 accactagca gccgatcaag tggaggtaga agttaaagcc acaggactca atttcaaaga
23281 cgtcatgatg gcaatggggc aggttgaata cgaagctcca ggcttagaat gctctggagt
23341 agtaaaagcc gtcggagaac tggtaacaaa tgttactgtt ggagaccgtg tgtcattctt
23401 ctcatttggg gcttttgcaa atttcattcg ttcaaaggcg atttcggttc aaaaaatacc
23461 agacttcatg tcatgggagc tcgccgcagc gcttcctgtc acttacaaca cggcctacta
23521 ctcggtgttt cacgtggctc gtgtacaaaa gggtgaaact gtactgatac acgctgcatc
23581 aggtggttta ggtcaggcca tgattgagtt atgcaaggat attggcgccg agatctttgt
23641 tactgttgga actgcttcca agaaggagct tttgatgaaa caattcgcaa ttccccagga

23701 ccatattttc tccagcagag atggcagttt tgcacaggga atcaaaagca tgacaaaggg
23761 tagaggagtg gatgttatca tgaactctgt agcaggggaa atgcttcgaa tcacctggga
23821 gtgtattgca ccatttggac ggtttgtaga actaggtgct cgtgactata ccattaatac
23881 ccgccttgag atgcacaagt ttgaacgcaa tgtcacgttc tctcttgtca atttggtgag
23941 cttggtgaga gaacggcctg aagttgctgc ccaggtctgg tccgatgtga tgaatctgtt
24001 cagtgaacgt aaactgaagg gtccatcgcc tctcacagtt attggaattt cagaattaga
24061 gaagggtttt cgaacaatgc aatcgggaaa acataccggt aaaattgttg cggttcctca
24121 gcctgacgaa aaagtcatgg taagttataa tcttgccaat gttgagacgg ttctgacttt
24181 ctaggtccta ccccgagata atgacgcacc tctcttccgg gccgattcgt cgtatttgct
24241 tgttggaggg cttggagggc ttggtcgtgc tgttgctctc tggatggtca agcgcggtgc
24301 aactaatctc atattcgcct ccagaagcgg tctctcgaag cctgaggccc gagaactcgt
24361 tgaaattctt gagaatcaac atgtcacggt ctccgtccat gaatgcgata ttagtgactc
24421 tttatccctg ggcgtactac taaaacgaac cgagaacatg ccacctatcc gcggtgtcat
24481 ccaaggtgct atggttttac aggtaatttt cactaacctt ttctgaggca atgagctaac
24541 tttataggat acacttattg aaaacatgaa tttgcgagac taccaaacag tcatcaaacc
24601 aaaggttcaa ggaacttgga atcttcacca aatgctgccg aatgagctgg acttcttttt
24661 gatgctctct tccactagtg gtattattgg caacgcaagt caggctgcat atgcagctgc
24721 atctaccttc ctcgatgcaa tggctgctta cagaaattca aaaggccttg cagctgcgac
24781 acttgattta ggcgttatcc ttggcgtcgg ctatgtggct gaaaataaag aacttgccaa
24841 acatttggaa cgtcaaggat ttgaaggtac aacagaaaac gagctcatgg ctttaataca
24901 atccgcgatt atcaaacagc atgaattcga tgtcaagagc cagattgtga gtggtcttgg
24961 aacatggaac agtgccagtg gtgctgcata ctccggtgct ttgttcgccc atttccgccg
25021 cgcggcgctt aagtctgcgt ccaagtccgg tcaaagcggt gacggtaagg gccgaattca
25081 agatgaaata cgagatgcca cttctcttga agatgcagcc gctcgaatat gcgaagcaat
25141 aatctctaaa gtctcgtccc tatcgatgat tccggttgaa gatatcagcc aatcccggcc
25201 aatgtcggaa tacggaatgg actctttggt agcagtcgag atgcgcaatt ggttgttcag
25261 ggaactcgat gcaactgtgc ccattctaga gctgctgtcc aataactcat tggcggcgtt
25321 gtctctaaag attgcgaaga ggtcaaaatt ggtccgacca tctcttttgt ctgctatgga
25381 agactagatg agactgatat ttcgttgaaa tacaactctt gtgacttgat agcctgtttc
25441 tcaatcaatt taagtagata tttgttagca atttggtgtg tagaggcatt gtcattctgc
25501 tctagaggac caataattag attcgaaatc acatgcataa gtccccggcc tgagaccaat
25561 gtcgctcgct atgaaagtta ctcggtttgt gtactagtta tatctctatg tctgacaagt
25621 ccaataccaa aaattaatct aataattcat ttgtgaaaca gatgaataaa ggagacctat
25681 gagagtctat atttctaagg atggtatgaa gtaaacagtg tgatcaagcc gtttctcaat
25741 ctatcaagct tttcatttac catctaagta actcactttg agaaatggca aagtagagtg
25801 agaagaataa tacaggttac taattgttat aggtattcaa aatccgacaa aagtctctta
25861 atttctacga aggaaacagt gaaaatttgt tttaacccta cctaggtatc tggacggaat
25921 catattctgg tctgactagg tattctaatg ctctgatccc gttggtaaat ttagtatttt
25981 aggctcgaat aaaaggtcaa tgactaagaa actttgtgga gatcctcatg acctgatact
26041 ttttttcata gtttctgcta caagacgcct ttggattttt cctgtagcgg tcttgggcat
26101 ataactcgta atataaatct gagacacttc acattagtat atcaatgtct cattagactt
26161 gaatagagtt tacctctttt gggatcttga atttgacggc cctgtcggca aaccacgcaa
26221 gcaaatcctc ggtggacgtg acttttcctt cacgaaaagt gatggccaca gctagatttt
26281 gtccgtataa ttcatcatca attgcgaagg caacagcctc tgatacactg gggtgctgtg
26341 caaatatgtt gtcaatttca atcggactga ttttctctcc gcctttattg atcaactcct
26401 ttattcgccc tgtgaggatg agataacctt ctgggtcaag gaacccctga tcaccggttc
26461 tgaaatagcc atctgccgtg aacggagacg aaataccatt cccaatataa ccggaagtca
26521 catttaagcc caaaatgcaa acctcaccca cactgccctg ttgaagtgat tctccctcgt
26581 cgttttttat cctgacttcg accccttggg gaattccaac tgaaccaggt tttcgtgtca
26641 aaggtggtag gggattagaa gtaatttgat gagaagcctc tgtcattgca taagcctcaa
26701 tgacaggtgc acccatggct atttcaagtt gtttgtggac cgttggcgac agaggtgaag
26761 aacatgatct cacaaagcga attgcgggca tcggattagg tagctcactt ctaagaagaa
26821 tctgatgtat tgtgggaact gcagtatacc agtttgcccg atgatttata aagtcactcc
26881 agaaatcagt ggcggaaaac cgaactggta cgatgatacc tccgccactc attaatggag
26941 ctaggaatcc agctagtatc ccatgaacat gaaacaaggg cattactaaa taacctctgt
27001 cggaaactgt aagattgtat gtggcttgga tattagccat ggtgcggcac aggtttttat
27061 gtgtcaaagg gacctatgat atgttagcca agaagcttaa aaataccaag ataggtaggt
27121 gctctgctgc ggggagggaa tttacggctt ttggtcttcc agttgtcccc gacgtatgta
27181 agactaacgc gatatcttcc tcctctggcc gctgagaagg gactttggtg ttcccgttga
27241 cccaattaga atggcgaggc acaatctcaa tctcttgccc attccaaaac acttccaaga
27301 ctggaacacc aagcaattta gcggctccaa gcgcttcctc atttttactc tccgaccctc
27361 gggtcactac caaggccgct gttctcaagt cctccagata aaatttgaat tcgtcctgtt
27421 tgtatgccgg attcagtggg gctgcggtag ctcgttgaaa tgttgtggcc agaaagagaa
27481 tgacaaattc aatcgaattg ggcaatgaca tggccagcac gtttttggga ccgatgccaa
27541 aggacgcgag cttgtgttgg cacgacagaa tttgaatcaa gagctgactg tgagacaaga
27601 ttatctgagg ggaaggaatt atcactgcag tagcatcgct ccatgatgct aagcaattga
27661 ataacgttgc catcgcagct ttcacttttt gcctagagga cagctcaata ttcttctggc
27721 actgttatca tgttacataa ttatatatat atatatatat ttttggagct ttaatggcgg
27781 aacatattaa ataactatac ttttgagcct atagtggcct tttcatgttt ctacgtgcaa
27841 gcctaggtac ctgcccaggt aggtaagcac tcggttaccg aaaacaaata tatccattgt
27901 gtttggctcc ggcatgttcg tggatagcta cctaggtaag cttccataga ccaattatac
27961 ctttttttaa ttttgaaaaa ttgttctttg taattctaac cttcactagg tagatatctt
28021 gaagaagggg gaattgcaat cctacagagc tttcttgtag agagctcctc caaccagttc

28081 gcattgaaaa ccacctagat atttcattct gactctttta ttaaaccatc gtggcacggt
28141 gctaaagtta cgaaagtttt tcagcaaact gtttatgtaa tctaatcgct ttcaggatgt
28201 caatggcaaa acttgatgtg tccctgtcaa ggccggtgct cagcggatct gcacaaattg
28261 cgaaagttct tcacgatctg gatgtgaagg ttgtatttgg catcgttggt attcctatcg
28321 tcgagattgc agaagcagct cttgcacttg gaatacgatt tgtcgctttc cggaatgagc
28381 aggcagccag ctatgctgcg acagcttatg ggtatctcac aggacgtcca ggcgtgtgcc
28441 tagtagtcgg tggtcctggt gtgcttcacg caatggctgg aattggcaac tcctcaatta
28501 acaactggcc catgcttctt ttagcagggt cttgtgagac cagtcgcgtg ggaaagggtg
28561 gatttcagga actcgacgcc atatcattgc tcagtcctca cacaaaattc actgggcgtc
28621 catcattatt ctctttaagc gaaacgattt cgcaatctta tcggatggct tgctatggac
28681 gccctggtcc tacctttgtg gatcttcctg cagatgtgat tcagagtcga tctgaggatc
28741 gttctggtga tgtacgccag ataacacacg acaacacatc acacgcctct acaaagttgg
28801 ttgagcgtgc tgccgtcctt atcaagtcag ccaaagcccc cctcatagta ctcggaaagg
28861 gagcagcgta tgctagggcc gaagatgtga tccgtcaact gatacagcag acgaacctac
28921 ctttcctgcc cactccgatg ggtaaaggcg ttatgccaga ttcttcgcct ttgaacgccg
28981 cttcggctag atctgctgcg ctgcgaggag cggatgttgt tctacttctt ggtgctcgtc
29041 tcaattggat tcttcacttt ggtgaggagc caaaatgggg accttcggtg aaattcatac
29101 atatcgacat ttctgcagaa gaaataggac gcagcaacaa tgtggaaata ggcatcgtgg
29161 gagatttaaa atccgtagtt ccgcaactaa ctctagctgt ggaaagttgg agctacgacc
29221 tatcaacaga atatcttcgg ggccttttag cctcaaagca aaaaaacgaa gcaaaggcta
29281 gagagctagc taaaatcgac acagttcctc tgcgatatca ttctgtgttc aacataatca
29341 aagaaacctt tcaaagatat tctcctgcag aggatggaaa tatcgtctac atatctgaag
29401 gtgctaattc catggacatt tcgcgcagta tattctctgt tgaatatcct cgccttcgtc
29461 ttgatgcagg ttcacacggt acgatgggag tcggacttgg atacgccatt gcagcctatt
29521 gtgcatacaa ttcacccgat caagattcta catctattcc tcctacttca aggaaaaaaa
29581 tagtctgtat agaaggagac tcggcttttg gattctcttt ggcagaagta gagactatgg
29641 cccgatataa catggatata ctgattttcg tcttgaacaa tgggggaatc taccatggcg
29701 atagcgataa ggcggaggag tggctaaggc tgcaacaacg caccaaagct ggagaatcta
29761 atggcttgag aagcacctca ctgggctggg aaatcggata tgagaatatt gccacaatgt
29821 gcgggggcaa gggattcgtg gtacgcactc cggaagaatt agagaaggcc actgaagaag
29881 gcttcaaatc gacagtacca gttgttgtga acattatcat acatcctggt cacgagcaaa
29941 aattagtaag ttatggtttc atagccgtac aagtaatttt tactaaacta ttacaggaat
30001 ttgcatggca aacatcaatg aaaggcaaaa agtaagctta gaggtcagtt ttggatgggg
30061 aaaaaacggt cgtgcctaaa ccatcgtgcg tacgcacatg caattgatct tgagtaagat
30121 atttagagac atgagccagc aaattatgga acgttatttc gtgatgaaga gtagaaatta
30181 tctccatgtg ctttctcttc cttcaacaag ccgtggtgta cagcaactag attctttcgg
30241 aaggacagta taagcccccc gattccttct ttgggggata aaaaaaagat ttttccgtgg
30301 ctggcaaatc taaatgtttt ccagaacagt aagagaagcc tttggagtca gattgtggga
30361 cttcgtcgtt gaataggggt aatgacaaat ctatgaagtc cagaacatct acaatcaact
30421 cagactcgcg tccataactt ttggcaatct cagagattcg gataccgaag gaacagtata
30481 ggcttttgga tgaatctcat tcgctatatg attgggtttt ctagagaaac tccccggtgt
30541 cgtttgggct taggggtgta ctagttgact atttgttact ctggtgggta tcattggcaa
30601 ttggggtcgg attctcgagt cagcaggcaa gtaagcaagc ctatctcgtg cttaatgtta
30661 cttgttttat gtatttattt tatggctaga taaacttgtt tgtcatgctg gcaagagccg
30721 ttccataaca cagtttatgg atctatcaag cgttattccc taactcaatt ttgttttctg
30781 caccaagtaa cgagatttga ctgcgatgct tggagacaca gattctaata tgtctcctat
30841 tccggtgcca gagcatgcta gtgcctcgaa cggcaacact taccttccac ttacttttcg
30901 tgcagaggtt gtactgaaaa acgacaaccc ggagctgctc aacggctttg cagcgcttgc
30961 aaaagagaac gatccctcaa tgaaggtggc tgattctttc gaaggccaga atactcttgg
31021 ccaccatact agcaacaatg gagtcaatac gcctccaaac gaaattgaac ctcttgttac
31081 gaaacctatt aattcatctg agccgcttga aatttcgcaa aatttgaatg acagctttct
31141 tcctccaccc agatgccatt gtcatattgc agctccactc acatttgatg cccgtgctcc
31201 tgcttcaaca cgcttacgat acctcatcaa atactcagaa tcaataataa gctgcccggg
31261 tgtctatgat ggattatcag ctcgcctggc taccagtgtg ggatttccag ggctttacat
31321 gacaggcgct gggactactg cgtcaaggct tggagctgct gatttgggca ttgctcaatt
31381 acatgacatg cgtacaaatg ccgagatgat tgccaatttg aatcctgaag gtccacccct
31441 tattgcggac atggatactg gctatggtgg gcctctagta atctcaaagg ctgtcagaga
31501 atatattctg gctggcgttg ctggctttca tatcgaagac cagattatgc agaaaagatg
31561 cggacatctg gctggaaagg aggttgtcga agcagatgtc tttgtacaga ggatcaaagc
31621 gtgcaaacat gccgcgaaaa aaatgaggtc tgatattgtc ataattgcac gaaccgatgc
31681 cctgcagagt cgtggttaca acgagtgtat cagacgtttg aagcttgcaa aagatgcggg
31741 agcagatatg ggaattttag agggattcca aagcaaagaa caggctgctc aggcagtgcg
31801 agacttggca ccttggccat tgaccctgaa cagtgtcgaa aatggtatat caccattgat
31861 tactacacaa gaagcccaag atatgggatt ccgggctata atcttctctt ttgcaacaat
31921 ttctgctgca tatgtagaag tcaagagaac cctgcaatac ctaaaacaac atggttctac
31981 caacagccag attactccaa aagacatatt caatgcatgc gggcttcaag agtcaatcga
32041 aattgatgaa tttgttggtg gaactgcttt caaagctggt gtgtgatgtg tttcttttga
32101 atgtcgtttt tatgtactta gtttacgcgt aaaatgcgtt gtagaatgct agaagaattc
32161 aacatttctc ttcatgttct tcgttcaacc cgaaacttta agtttcatct tactattttc
32221 gttctgtaaa ttattcttac agatgtggga agtttggaaa tggatattga ttgaactctt
32281 tgcaagttag ataatgatga gaacaggcgc attttagata tgactgtcaa taacaggaac
32341 tgaatgctgt gaataatgac agtgctgttt tgactgtaga acgtatttta atttcattct
32401 tcagttactc tttttgagag ttggaagtcg aatagtatga actatgagga ccactattgc

32461 cgccacaatc acggacatca caaatgcaag ttttaatgcc tgctcatacg cactggtgac
32521 ctgcttttga gtcatcaaat ctaagtcccc aatggccctc actgattttc tgactctctc
32581 aatgatctgt atttcaattt cacattagac aagtacagaa ttcatgaaaa ttggatattt
32641 acctcttcct tgttgtctcc cgtgacaaat tgattaagat agacaaccaa tgcattctgt
32701 atgaccaaag tgctagaagc gattccgagt accataccaa tacttcggaa caatattaaa
32761 gtactactca ccaccgcttg ctctttttgt tggctgacgg tcaaaatgct cataaaagtg
32821 ccagggttct ggaagccctg tcccatgctt gggagggcca gaaaaaaaga atatgtccag
32881 tccggaaagt ccggacgcat gagtgcgaga catattgagc cagcaagcag caggcttgtt
32941 ccaagagtca atgaccattt cagcttctga gagtaggtta tgataaatcc cgttgccgtt
33001 ccgataatac tggtgcacaa gttgggggaga agaagtcgta atcctgaggt tgttgctgtc
33061 tctaggcgca ctgcttggaa gtataacggt gcgttgaata ttatgagata ggtgattaca
33121 gaaccgaaaa agttggctat taaaagcccg gctcgggggt ttcgtgtcac tatcgccgga
33181 ggaataattg gtcgcgaggc gtgatactcg acgtaaatga gcaaaggcgt gcaaattccg
33241 cataagacaa atgatgtaat gaccactgga tgatcccaat tgtaaacatt gccacctagg
33301 ttctagtgcc ttattaacac tattatcaat aaattgaaga acagggagaa gtgttcaact
33361 taccagggca agaatcaaga atgttattgt agaagacatc aaaatagatc ccttatagtc
33421 aaatgtcgca aacgcagccc gaattccttc gctggcaata tcagggtcaa tccccaagtc
33481 ttttggtgtt gtgaaacatg ctatcaaaaa cagtgtggcc agaataggga cctgtatgcc
33541 aaattcccat cgccagccca aatgatcggc aattacacca ccagttgcag cgcccagcgc
33601 agatccagta ccgaaggcaa tgtgaacata tgattgatat gctccacgga tttcgattgg
33661 gaccaagtca gacgttatta tagagctcat agacatcata cccccagccc cgagaccaca
33721 aatcgctctt ccgaagatga aaacgccaac gttgtctgca attgcacagc agatgttgcc
33781 gaggaaaaat atggccaaag agaatacgta aggctttttc cggcctatag tatctgaaag
33841 tcttccgaaa agcggttgaa aacttgtgga tgtcaaaagg aaagagacag taagccaaga
33901 ggcactatta gatgcctgga aatacgatgt aatcactgga tgtgtagaag ccatgagagt
33961 cgagtcaaac attgaaacaa atgaatttcc cagcactcca gcataaatta gccagaattg
34021 tgcaacactg actcctccga gataaggatt cttcagcttt gaaaatattg cttcttcctc
34081 ctcttcatct gcgtctccgt cttcgggcct gggacgcgca tcacgagacg tgtctggtaa
34141 ttcgtcatct tgaatatttc tttgtggtcc ctggcttttt gggactgctg cggttcccaa
34201 ataatccgga aatccatttt ctgttgcgtt tgtggtgaat agaggagtgc gctctgaggg
34261 cctttgggcc tcagagtcat gcgagtcggc ggtcattatg tgctgcgggt gtcgattcaa
34321 ttttacaacg atgatgtgga ctcaaatagc aaaagcttct aaggttggta aattgattct
34381 atagtctctg aggctttttt ggattaattt caaataagaa gaaaattttt agagcagtca
34441 tcttgcaagt cccgaagccg atggggccca tactttcgta agactcgtta cagggctgac
34501 aacagagttg taccccctatc tttcaacaat cagcattgct ttccagaata ggatacgaaa
34561 ggtttccagc catactgtat aaatcaaaaa tacgtctgtt cattggaata cgtgagtcgt
34621 aacataaacc tacatagtca aatttgtgag aaagtcacta gactgcaaca gaattaaata
34681 tctataattc attgagagcg agtcgagcct tgcattttag gaaattctca ttcaattcgc
34741 ttcgatataa tttgaatgca aactgacaag cgtacctggg tatctagaca gcggctgtat
34801 gttcgcggag cggatacgat catccgtagg cgaaattttg cgtttactcg tgctacgata
34861 ttcggtaggc tttttttggt tacgagcgaa aatccatcgt tgacttccag aatatggacc
34921 taatattggg cttgaagttt aattcttgaa tgaaacaaaa tacagtcaaa caggcgcgaa
34981 atagtacatt atgcaatggt aagggaaacg gtccgcgact attctcggtt tctgaatacg
35041 aataaaatta cgaactcaac attgagtaat gagtatacat gcttcggcaa tattttgtaa
35101 cagaacaaat tcactgactc tgtgatatct ggctggttgg cactgtaaga attatgccaa
35161 gttttcattt agatcgtgcc tcgttaatca ttcacgcagt ctttgtcact gccgttcgat
35221 caccgaagaa accaatgggt cacttctagc tgcgctgcta attttcgcga aatttgaccc
35281 ccaaattttc tatcatttcc agaaatcatt gattaatgta tcagcaaacg gtcgccgaaa
35341 tcctaccact caccacaggg caaaaagtgt cgaatgccta tctctgtatg tttcgggaca
35401 acgctacctg ggagatgggg gagatcagtc atcccatcat ttacccctga atgacatgta
35461 tacccaagta aagtaatgtt tagatcgttt caatccaaga tagcaatata tcagatgtgt
35521 acattaacgc gataacaaac aattattggt gttgtatgtc cattaacctt tcacagttac
35581 gtattgctgt agagaggcct caacagcccg ctatcatctt cagtcgttct gatatgacca
35641 aggctctact acactcggga ttggtaataa gttctatggg aactgtaatt attcaacttt
35701 ttgccgaggt agacgaaaag taagcttgtg ctcatccttt tcatgattgg tagtctgcac
35761 aagcaggtca tataggccat cttgacaagc aaatttcgaa ttaatttcat ttcgctcttc
35821 tttttcgtga acaaaactac ttgcgctatt ttattttgga atctatgtga acaacatctc
35881 caaccaacgt aatggcctca agatctatcc cttcctatag tcaatatcct gactatatcg
35941 tcggttcacc agcgccatac gttctcgaga tcctcatcga tcgaccttcg cgacacaatg
36001 cctttacaga ccgcatgtgg catgatctgg gaagactctt cgatcaagtt tccatagacc
36061 ccgaagtacg cgccgttatt ttagccgggt ctggtgccaa tttcagcgtg ggtctagaca
36121 tggttgaagc tgcgcagggg gagatcctga atggtcttca gccaggcgtt gagcctgctc
36181 gtcgcgcgca gactatccga agatacatat ttgcgttcca agactgtgtt tcagctgttc
36241 aacggtgcgc aaaaccagta gtatgcgtgt tgcacggtat atcgtatggc atctcaattg
36301 atatcgccag ctgtgcagat attcgcatat gtgcagaggg tactcgcttc tcggtcaaag
36361 aagttgatat tggcattgca gctgatctcg gctcgctctc aagactaccc aagattattg
36421 ggaatctagg ttgggtgaag gaagtttgtt tgactgcacg agaatttggt actgaggaag
36481 ccaggcgtgt tggtttggtt acagaaatcg ttgattcaaa agcgaatgcg atggaaaggg
36541 cgcttgagat tgccaccgtg ctagccgaga agagtccatt ggcggtccaa ggtaccaagg
36601 aaattattaa tcatgctgta gatcacacga taggagagag tctcatttac actggagtgt
36661 ggaattcaag tgccttacag accaaagatg tcagtgaagc tgttggtgcg tggaaggcga
36721 agggcagacc gacctttgaa aagctgtaga tgtttgagga tacgagtgtt cagttatgga
36781 tattttgtca ttgataaaaa acattgtccc agatttctgt atattgaaca actgagatcg

36841 caaataaaga gttgaggaac taaaatgcaa ccttgaaatg gagtcggggt ttaatcccat
36901 gtattggcgt aattccggta aactggcgga ttttcgctga aacccagcac tgggtcgagc
36961 tgcattaaag acagtacaac gacgtaaaaa ccaaaaacat tatctagact tgaaaaaacc
37021 aaagttattg acttgtgaaa agtgaattaa atacaaattt tagtatactt tctagaaagc
37081 ttgatgtaat aaacatggta tctgaatcat cggtaaacgc tcccatctca cacccactct
37141 gttcaatcgt cactctgtat ccaagtacga taattgtaat tggtgaatga ccaaattact
37201 tatggctaat tagctcaagt tttaccataa ctaggctaat tcgtcgcctt tttggctttt
37261 ctcgcctcct ctttcacacg tcgcttctct ctccatcccg ggccccattt catgattgct
37321 atcaggattg gtgagaaagc catgcaaaga aatccaaaga atgtagccgc ccatccacgg
37381 cccatggcgt tgatcatagg gatgatagct actgttgctc ctgcaccaag ccaacatctc
37441 acgagattgt tggcagcagc aactctgccc gccccttcag gataaagatc aaccaaaaga
37501 atggataaga ttgtgaagca caacgccgca gaggtcgtca aaatgaagag gaaaattaag
37561 ggaccggtga gatttgtgcc gtaatgaata caccatccgt atccgatagt tgatagtgcc
37621 tgaactgtaa acattggcag tgctacctcc aagcgcgcct tctcaatagg gaaatggcgc
37681 aagtcagttt gtttggtctt aacgagggga aagccaagct tctttgcgta tcgcctatag
37741 ttcatatcaa cggccttgcc accacccaga acagcaatca ttgcgccgag accgatgggt
37801 aggaaacaca gcgaaatctg aatatcgttg tatccataaa caatggtgaa ttgagagggg
37861 atagatgaaa tgacaccata gtatgaggca aaagagacgg ccacggcaaa cagtattaag
37921 ccaacctctg gttcaaagat gacggtcaga gtcgtaagcg ggttgacgat cttgatcttg
37981 ttcgccttag ggcgatcctt gcccgccgca tggtgcttct tgagatgata atacgtcaat
38041 atggacatgt tccaattagg tggaggaaca gtgccgtcgc caacgatctt ccgacaggtt
38101 tctgggaaga acaacagcat gataatgaaa accacgcccg atgaaattac cagaaaccag
38161 aaaactgagc gccatcccag atagtggcca agaagaccgc cgataatggg gccaatacca
38221 ggacctttga gagcattagt ttgtggaaac tcggtgcttt ggaataaggt tatttaccaa
38281 tgaattgacc cgacgaagcc cagcccgtaa agtaccctct ttcagatgaa gtagcaatgt
38341 cggcaacgac accactagca agagcaatag tgctgctact tcccgcactc tgaagcattc
38401 gcaatactaa aagggctgcg tagttgctct gaacggccag accaatgttg gctccaagat
38461 agatcacaaa acaaattatg aatgctggcc gtcggcccgc gccatctgag ataccggcga
38521 tgagagtagg ggtgattcct tggaggatct gtaccggtat cgatcagtgg atgatattcc
38581 taagaggatc tacaaaatca acgagccaaa gaggacgaac taaccatata tgtggtaaca
38641 gacaaattaa ttaatgtgtt gctcacgtta tagtaactgg cgagtgtgtt gaaaataggg
38701 tagtagatat tcgcacttag aggagagaaa aaactcgagg cggaaaccat aatgacgagg
38761 aacattttca aggatagct gaaggaagag aaatcggctt gaggaggtgg agctggtttc
38821 tccaccgcag gttcagcagg aaggtcccca ttttgcgtct tttctgcatt ttccactttc
38881 aagtcctctg tagcttgaga tccgttgccg tttgtttctg tggaagtaga agaactgacg
38941 aaagtcgcac tgttgggcac tgcaatggta gtcgaagtga gagctggctc ttctggaggt
39001 tgagcggcca tgcttagaag ctaacgtgga aaatgtgaac gcagaagaat aaacgaagac
39061 gatgtctatt cagtgaattg cactcgatac tgctcacgtc cctggcgtcg ttgtcaacgg
39121 gcaggacaag aattgtttct cgacaagaga cgagaaagaa aaaaaagttt gactcgtaaa
39181 aagcttatta gagctaagaa atatgaatga gactttagca gcttgtccag aacgagctaa
39241 aggggctact ctggctgata aggccggttc cggttttatg cccgggaccg ggaatagata
39301 ccgtatactt gaataagccc tacggagtac gtactcgaag gaaacaatga tctccggttc
39361 cgctttgacg ctttgatgat ttagggacgt actccctggc gttacagcgc atcctgtcca
39421 acatcggccg catgacaaga tagccaaggc caatcatcca ccaacacaaa agcttgtatt
39481 ggcctcattt ggtgcaatag accccactag cactagtctt tgatcctaat cttcagctgt
39541 acttctccgc ccaaggaact agcaagttac agctacggta accgaacata cgcacaccct
39601 ccagtgcttt cgtattgggg tattatgcca aaacaagtgg atcgccaggg agattaccat
39661 gcagtacaag tcaagctgag aaattggagt ctatccttct cttcttgggg ccattttgca
39721 cagggattgc cactgactag acaagacttc gatattaacc gtgtgtaatt tcttcgtaat
39781 tttgtctccg tcaaacttcg gggagttacg gtaagtggaa atcgtaccgt aactaactga
39841 atacagtaat tttactccac tagcctaatc ttacctcggc cgatctaaca actagttact
39901 ttcaaggttc aacctcctct aactatgagg acggtaaagt tccgcctgga tttgaagaat
39961 gcaactcagc ccccccaact gctgctagat tttagacgac atttagaaag tgaatggtgt
40021 gaccagttcc cttgccaggc atgaatctga tcgacaccgg acgggtcttg actatagaga
40081 ttcgggaaat aacacaatgg tcacatccat ggtacccata ttgaccggtc aagctgtgca
40141 gccatgatgt ccggagagct tcatgagccc gctaccggcc gtgaatacat cataatatcc
40201 tatcagactt gacaatgatt ttggatcaat tcgattgatt gcctgtcttc tcggacgctt
40261 cattgtctgt gattcattta tcttgtgttc aagtttgagt ccactttggg acaaatagct
40321 aaaaccgcac ttacttgatt ttttttcct atctgctacc atgtatatct tttcgccagc
40381 tgttttgtta actctacctt cgttgatttt ctcatcccca tcaactttag cctcgagcaa
40441 ttcggtgaaa cctcaccctt caaaccccag tctggacgtc acctgtgctt acgccgcgag
40501 agaccacata caatctttgt tcaccgcgct gacgaccgga aactctacaa tattttatga
40561 tcacgtggta gatgatgtgg actggaatgt ccaaggcact catcctctcg caggtcgcta
40621 tcacaacaag acggtattct tgattaatgc cgtcaatcgc attggcaagc tgcaggatgc
40681 cagtcgtcct cactcgtctg agctacttaa catcgttggg ggttgcaatg aagaatggag
40741 cgcgcaagaa atcagagtta cagcttatct aaacaatggt atgccctagt cttcattttg
40801 tccgactatg ctggtccttc atttcatcga ttaacatttc tctaggcgcc ttgttcgata
40861 acacatatgc ctggctcacc cgttggaatc ctcgcggcca aatcgttcag gttcgtgcgt
40921 atctagattc tgctctagtg gccaaaagtg tctttgaaaa cgaggcatca acaaattcta
40981 cgtttaccac tgtcagggat actcccgagc ctggacccgg tggatatgga atcttgcctt
41041 agaccggttg cttgaaataa atagagttct ttcttagaca atcaatatcg attcaaaaaa
41101 aaaaaaaaaa atggtgtcaa tttttctaga atgtactaag atctggcaag gatcgttcct
41161 aagccagtaa tacttacgta ttataaactg ctcagtgaat tgtcgttgaa cttaagaata

41221 ctgataaaca ataattatgg ccccgaatag tagactcaaa tattctcatt gtaaagatca
41281 ctccaccgat gtgtaccgag tccaaatctc attcttcttc agttcatcaa tcacgtcctt
41341 gtccattgtc attgcaactt cccagctccc atcttctaga cttggcagaa tcacaggaca
41401 gccatcaatg agacctttat gcggaagacg aaatcgttca aactttccaa ggccgccaaa
41461 atcgataggg atatcgttga actttttcca actagtaagc agcacctggg aacctgtaat
41521 atcatctata ttggcgtaca tgccacgtgg attcttggcc tggccgatga ctttcacaaa
41581 agcttgggca ccaccctcca cgacattgct gataccttcg cggattgcat gggttgccaa
41641 cgtcagtcca tttggaccaa tcactttcga gactggaagc gaagccgaca tctgcatgaa
41701 ggcgttacca atgtaatcgg ccgcaagagg aggcattgtc agagctcgaa attctacggg
41761 cattttgaag ttactttctt cctcctgagt aactttccca attgcaacac gagccataat
41821 cacacaattc cagatcagtc cacagacggc atcgtgcgag ctaataaacg gttgcaaaac
41881 ggcttcgtct tgggaatcag ctgattcttt atgaagaaaa gacgaagcat cgtttttgag
41941 tgctttaatc ttggctgcag atagacggaa tgtctcgcta gctaccggct ttgtcgattc
42001 tttcatccat gctggttttg gaggattggt atcaaacaag gtataaccac gagcgctaga
42061 gatatcccca ggcacatcaa aattgaacaa gggcgacttg tcgaactttc tttcagacag
42121 gtgataatac ttttcgacac ccagttgctt cccttcgcga atgtcaatga gccgcttgca
42181 atggctagca agaactcggg cgacgaaaga gtccccggac gcgtccattg caaagtggtg
42241 tataccgata gcaagaagca taccaccgtc caccaagtta gcttgaatgc cacaagtagg
42301 gaccgcagct gctccgtagg agaatcggtg tacaggcagt aaaagctcag catccaaagc
42361 ctcctgtttg aaattgctgt tcttgagtcg atcgtagttt aactcgggga tattgcccag
42421 attcttgacg ttgagcggga catgatgcgc ctgaagctca cgaagacctc cgacagtatc
42481 gacaaccttc gtagccagta tagggatctc gttcactgta gcctgaacgg cttcccgcaa
42541 gatattgacg gtcgatgcat tgtcttgggg ttgtggaagt tcaatgcaaa ggatttgttg
42601 tagataggta ccagatgatc tatgatccac ggcagaggga gaaacatcgt agcctggacc
42661 tgagatctca aacgttctgg tagacccaat ctcgatcttg tcgagttcat tttcaataga
42721 tcccattttg aatgtgcgag aattggtttt tcgaagacta ataggggttga cacaaaagtg
42781 gattctagtg cgctggaagt attttaatat atcaaatatc aaaaatgaac ctgggaaata
42841 ggatgttggg tatgcaatct tctttgtcca caattttaag tggatttctg caggtggtcg
42901 taaaatatta atcgacaatg tcaaactttt tacctttaat tgaaattagg atgtacctaa
42961 cagtcatttg ctaagggtta tggatggggg acaattctgt aagtaatcat tttacttaat
43021 tttattcgcg cctcctcttg tcacagccct ggaggtattt tagttgcgat tacttgatgg
43081 taacccaaat tttcaaaagg cacatttcac ttcggttccc gatctcatgt cttgggggga
43141 ctgaagatct ctctcaatac ctacaggatg taaaacagtc taacagctta gggtggccaa
43201 ttagttctaa cggtaaccga gtatgaacaa gtctataatc ttcattcatt cacatttcgg
43261 ttcgctcccc actagtatgt acacattact atgccaaaaa ataggagacg cctgttacgg
43321 cacagtagga aaagacatca agaatcgaac catcccagtc gatagaaact gccattacgt
43381 aggttgtgtc atcatatacc tagtaaaagt gttcaatcga tatttaatca tcccttagtg
43441 ttccatgggt ataccgtcaa gcttctctcg tagaccaagc tctcgggact ttcgtagaac
43501 gctaaactac ccatactttc aattcaacag cccctactta tcgcaaaaat gcagccaagc
43561 aaggtaaaat ctagcacacc tatgattttc tttcaacacg aaaaagctaa tgatacgtag
43621 gcatgctgca gtgtccctcc cgttgaggct gactacaaac ctaaaggcga ttttgaagac
43681 tttgccaggc taagaacatg tgaagttttc ctctcgaaaa actttttcta aacgcaatcc
43741 taatctcaat tagatcatac cggcccttca tcggcatcca cggccattct ccttgtcggg
43801 gatatttttg gtccaagcgg ccaagtactt caggtatgat tgtttcgcta aacgacagag
43861 gtggtccatt tatcttacct actctaacca ttatactaca gggtgccgat ataattgcct
43921 acggcggtga aacgaagtac caagtcttcc atccggattt tctgcgcggt gagtatgccg
43981 agcactcatg gttccctcca gatactcctg agaagggcgc cgctattgga gtgtattttg
44041 gtggccccgc aaatcctggg aaagctttag aaagtattcc gtccattatc aaagcgatag
44101 agtccaagag taacggaacc attacgaaat ggggtgcatt agggctttgc tggggtggta
44161 aagtaagtgg tggcgtcgaa caatcacatg agaaactcat actcgagact aatttccgcg
44221 aatagatcgt gacacttaat tcaggctcgg gaactccatt cttggcgatt gtatctgctc
44281 atcctgccat ggttgatcct caagacgcac cgaacgtctc agtccctttt gctctacttg
44341 cctcgaaaga cgaggatcct acagcagtga agaatttcat aaacgatctt caagtcgata
44401 actttgttga aacctatccg gacatggtac atgtaagttc caagcgatac tgaatttatt
44461 ggaatgttta ctaatatcac ttagggcttc atggcagctc ggtaagtctc cttcaatatg
44521 atacctccaa tggcacgatg gaaaactgac ttcgaccatg tagaggtgac ttgagtgacg
44581 agaaagtcaa agcaggatat aagcgtgctt acgaacagtg cttggagttt ttgtaagttt
44641 tcttgggtaa tgctccgtct tctattcctt actaattctt taatttgtag ccatcagcat
44701 ttataatcaa ctaaacaatt tgatcaagaa agttgttagc tgagacattc gagggagctg
44761 caggcgagta cagaattcta aaaactggca aatgcctttg ctatgtcctg cgcaggtgta
44821 tgctttagta tcggacacgc atttatgttt gccacgaaat cgacccgatt tcgtaaacag
44881 gaggaccttc ataaatacga tttgaaatta caaaagtctc agttctttgt aagttgatta
44941 tcagtgacta tggtagtcag gccgcggctc cttattccag attaacctgg aagagaactt
45001 tgcgcgttgc ataggaatta tgtacattga tattgggtct tgggtttccc aacaaccaaa
45061 tcttgaatcc agcagttgat taaaaggtct gttatattat cttaaaatca tctagtataa
45121 ttagagaaaa tgaaatgtgc cgtacgtgtc tggaaaacct ctgtagttaa gctgaagcat
45181 gttcatatag gtaataaata tcgatgcatt ccgataggta gatccaagtt cctcgattta
45241 agcaaggaat catttgaatg aactccaaga gcttcatcta ggttcagcac gatactttga
45301 ccacttttcc gggattggtg aagttcttcg tcacaaattg agctgcacct ccgttggtag
45361 taacataaag actccactcg tcgctttgcg cgcggccaaa ctcgagggcc gttgatcccg
45421 ccagcaaagt attggtactc aaggtgatag agttcgagct tccagctgca gcaaatcgaa
45481 gctcattgtc gcctgcaatg aacacatctc ccagcggatc aaaaacgaaa tcatcgactc
45541 cagcgagacc tgaaactact gtagacgcag agcctgtcgg cgccccattc ccgttcacgg

45601 gaacctttcc gagcagactt tgatcggtac tggaaaaata caccgtgcca tttttgactt
45661 tcaaaccgtt gattccaata tcttctccgg ttgtctcgcc agccatgagc gagctagtta
45721 tgagcttttc ggttgcacca gtgttgacat tcaacgacca cacgtcaccg ccgcgagagt
45781 cggcaatcaa aagaatgtca ctggtagagc tcagtgaagt cattccattg aaaaaaatgc
45841 tttgagggaa atcggcaaca tgggtgacag atggagacgc gcctccagga acgtagccat
45901 tcaagttaag tctccatacc gagtaggagc cgggacctgg tgtgatcgta acggtagagg
45961 tgttgcctgc aatcacatag aaaatatctt tcccaagttc agtgattcca gccacagcta
46021 gataatcatc gaaagtatga ataagcgtgg ctggtgtagc aagggttgga tcgacctggt
46081 agatttcagg ggtattgagc agggtcacaa cgatctgatc attctgtctc acggccaaat
46141 tttccaccca tgttccagtg gggaattccc atacggtgga tacagtcaca tctgagagat
46201 ctaactggga tcggcggaca aggggtgaag ccatggtgac actgactgtg ctgaacagcc
46261 agaaaatgat tggagcgcgg agatgcatgg tgcaggagat aattgatgga aatataatca
46321 attttggaac cgaaagtgca acctgcagat tacaaagaag gaagacacag cgtcgatctt
46381 atttataggc atttttggga ggcatgaatg acaagtcttc aagattactc ggtgttgaat
46441 tgcttttttc gatcccttac tccagaagaa ttagagttcc tcgtctgcgg aactttaagt
46501 tcgaatagta ctgtagcttg cgctgtcgat tccagcatga tcgactaagt cagagtaagt
46561 ctgacatgta gttatgggct ggatctcgta aaattgcgcc tacaatctac gttcgagtgt
46621 gtggctatga attaggaatc cttccttggc cccaaaatat tgactgaact caactagaaa
46681 acatctggac tgggcttttg aacaattggc atgactcgta ttgaagtaac tagaaatgcc
46741 ggaacgtggc atggcgagaa tatctaaaag tgaactctct aaagtgctcg taattatagg
46801 gaggacgtgt ggttttcggg ttaccattaa attacgcatc tgatacacta tttcaattag
46861 ccctttgggt ggatgctcac tcgggttacc atcacaatac gacgcgtcgt atttgtattt
46921 caatctgcat agttatgcgg ctgagttgtt gaacattgtg cggaaatttc caacaatatg
46981 cggaactttt ctcgtaatgt tggctgaaag actgtccttt ttcaccatga ttgattaatt
47041 atctcgtatt cgggatgtta ttgcaggaag taatggccct ctatcgtgac atgggtaaaa
47101 tcgaaggtgg attgacttca gccgcatctg ctggtaacaa accatacaga tgaagtactt
47161 tctagcatca aaattactcc atgcatgtca ggatggtcac ttcggaaagc cattcatgca
47221 ttgaaatcga gatacctagg catattagtg agctaagtga aattgaagca atatttctga
47281 tctgagaccc gtcacgacat agtagtaaaa accttggaca cttcgtactc taaagactca
47341 ttgatcagcc aacttcaaat tttactattc agagccaaat ctcagatata cttaataaat
47401 cttagaagtc tacctaagta ctcagtctat tcaaaagcac ttcttgctca ctggtgaagg
47461 cttcttaatt tgttccaatt caaagttgat aatgcctttc agaaattcgt atttatggtc
47521 atcgagaaca gccctgtgct gcaaaaacta tagttacaca accagcaact ttaagcaata
47581 tctatgatca atctatgctt tgatcaaaag ggaataaatg gcaggccgtc cacggctatc
47641 aaaatagctc caagtcacaa atcatactgt atacaaaaat aatggataaa cgcaaaagga
47701 tcgagctatc ggccgatcga gtaatatatg gttgctctag cctatcgcat tcaggggggt
47761 agacggaagg gtagacacga ccaatatact cttattaaat cgctagatgt catatatata
47821 cggattgaaa aatgtgaagt ggaaggcaaa aactgcatag aaaacgcttt tttagcctgc
47881 cataggatcc gtcacaagat ctccgaatcc cccattaata ccaaagaatt ggaacctaaa
47941 tcagaacggt catgggtcag tgaaaaagcc acaatggatt attttaatgg tggattcatt
48001 tgggtatagg taaaagaaga atcttaccag tcttcgacat tcaaagtcgt cggggctgta
48061 ggattgccac gtgagctatt gctccaattg tcgtcctgac tagtggaatc cacaggattg
48121 ccagcaagag ccgcatccag accagcccag ttctgcattc cttctattcc aaatatcgat
48181 tgcggtgcag ctgcgccagg caagttgttc atgtccgagt gaacaccact cgcggacgca
48241 attgacgggc tcccctggct tcctgcctga ttccgacccg gcatttgctg ctgtgattgc
48301 tgatggtggt tcatgaagac attaccgtga ccacccatat tattgacgga agattggatt
48361 tggggatcta tcatgctgtt tgtttgtctg ggaaagaagt tggtgccgtc tgggaacaat
48421 tggtccggct ggaagttctc ccaaagcgtg ggagatagat ttggcgaggt ccgggtcacc
48481 aagaagaggt ccgggggagt ggcaggaagt gaaaaggacg gattaaacgg agtgtttgct
48541 cgagaaggtc cagtattttg gataccatcc tgatgcttag taatatgagg cgaaatttga
48601 gacagcgaac ctgccaggcc tggttgacca gacatgggat tgatttcacg cgaaggcgtc
48661 gcagctggag tttgcggacg agatcgctca tacgaaactg gtggtgtcgg gttcccattc
48721 ggtaaaccga ggtccatatc gtcgaacttt ctcttgggag gctctgtttt tcgatggttg
48781 gttccagcgc tctgttcagg ttccgagggc ctattcttct ggtgcctttt accaggtgct
48841 ttttgcagac gctcctccaa catcttattc cccagaatag attcgaacag cgtatgaacc
48901 atttttgcga ctaaccaaac tttggaaacg tccttcaacg catgcatgca ggtgttcatt
48961 cgctcctgac acgcagccac gaccgacggc actgacgacc gcatctgata tacatgcata
49021 attaatgctg agaaaagact atacacaatg aaagccggtg tataccggat ttcgttgtgg
49081 ttttgtagct tttccacaat agaggtgatc attccggcag cttggaaagc aattgtccgt
49141 gagggatacg atatttcctc gcggtagctg ctcgcactgg ccgaagctgg aggcatgtgt
49201 gctcgatgca ataggcaaag agttgtgctg catccgcgtc agtcatacat ctctcaattc
49261 aagatgtcaa tcaagagact cactaatagt tcgaatgcag caaagccgcc cagaaatgat
49321 ggttttttct ttcccagtac actattttgg gacagttttg gagccagtcg gccaacgcca
49381 tatcggaatg cgtcagatca atagcattgg tccgtcggga tttcgaagca acagagtact
49441 gttgggataa gacaagtccc ataatctcgc acagcttgac gtactgtaag aaaaattgca
49501 cgtggacagg gtcaggtggg tactcggctg gttggtcgcc ttcgtcttca ataaagtcat
49561 cctcagtaag catctcgacg tcagaatcgt cgatgttgat attaatcggc cttccgaggg
49621 caacagcaac tgatcggtcc ctcgtgaaaa gcgtccacca tattcgcttc cataaacgct
49681 tgtcggatct actaagctgt gacgtttcga cgctaagaat ggaaagtcag caatgcctgc
49741 gtagccccag attcgggttt attcgactgt tctatactaa cctgcgatgc ataccggaac
49801 cctgtgcgac tagaatcgct actcgactcc aataaaacac attcttggta acatctaaat
49861 ttattgtcag tttcatggct cgcattcttt aggatgaact acgcattgag gctgaccttc
49921 ggggccttcc caataccacc ccattagcac caatgcttga acaagagtaa cacgatcgtc

```
    49981 ttcgtagttg gcgtcataaa gtgccttggc acgtttatag aacgttgttg aggctgggat
    50041 tgtagagcca ttttgatcca tcaattttga gttgttgcac actcgagatc cagctaataa
    50101 gatagcttgc agcagcagca gtgacggtgg attttgcggg tcatggtatt gacgcatgaa
    50161 gcggttgcga ttgacgattg ggaccactgg agccacccac ttgaaataag cctcgacaag
    50221 ttcatcgcaa agatcccgtg gaggaagcaa gaacgcgcca cgtcggtgga gaatctcaat
    50281 ttcaagactg tctaactcgg tcaagcggcc ttgactaccg cgtatagtgt cgggaagagg
    50341 gtaatgcaca acatccgtaa agcctcgatc gtggaccaga agcgataggt tggaagattc
    50401 acccaaatac gcaacgcgcc cgggttcctt gatgggcgct cttgtgaact ttggcttcat
    50461 gatttgagca ctgaatggtc cagaatgggc tgcttctgga tttggcctga cagaaggcgc
    50521 caatgtattc gccggcatcc cgtcgataga tacgggagga ttcgctgaga aagtcgggtt
    50581 ctcgtgcgtc cgcggtgtac tagattccct gtcatcgtcc tcatctttat tgacatcttg
    50641 gttcttagtt tcgctgtatc aggtaaccaa gcagttagca taaccgacgc ttaagtagag
    50701 ggagagggtt tttcttctct ctttggcaac atacccgtta tcatccttgt ttctcggctg
    50761 ttgattcttc ttcctctttg gctggggtat ccgacattca ataccaaaag ccacacaatt
    50821 ggtgcaaggt acacctaagc tggcagcgtc gcaccggacc tgtcatatgc gagttagatc
    50881 catttgctcg tgatagtccg agaaaggctg gattcgccga cagacgggcc gagatcctgg
    50941 caacactgtc tagagtaggg gagagaacgt accttgcgag catgacaagt ctgttgtaga
    51001 acgaaaaaaa aaaagtcagc atataagttt tcctttttg tcgatttaaa catcatcgtt
    51061 taaaacccct cgtcctcctc cagtgacgag acggtctggt tatttgatgt ataaaagcga
    51121 caatgcagag aacatagggt ttccaaatta ggcatggtgg acggatagtg ggggcatcca
    51181 agttttgccg aggcacaaac agggagctgt gagaaacgaa atgcgcgcgc aacggcactg
    51241 gagagcacaa gattagtgtg gaaaaacgta cctcgcatgc tcgggacgcg cgttgtctgt
    51301 aaaatttcgg ttaatactta ttagggtatt tatatgtaaa cgacagataa agacattaaa
    51361 aagggatgat caacttgggg cttggatttt ccggtcttgc gtccatccca aatttgcgat
    51421 tggcagggga taaagcgcat cgactccatt cacttgctgg gaagatagtg ctaatggtcc
    51481 ctgtaggatg accatataag agggaagacg atacttgtgt gatcacgtac cttcggaaac
    51541 ttgcgatgcc agcattgggc accgcggatg atgatgaaga cattgaccca gactgatgtc
    51601 cccctgcttt agatcggtca ttgctcgggg tgggactcgc ttcgttgtgc tgttgctgct
    51661 gctgctgctg ctgctcttgc ttttccgtgg ggttatccat gttggcttta ctgtctgtat
    51721 ccataattta ttgcagcaga acgtagctga tttccatcga ttcgtgagaa agcgctggtt
    51781 ttcggtgaga ttgaaacgtt tcgtttgtgg cggcgaaata taattgatac gcaggtttgt
    51841 gggaaattca acgcagtagt tgatataagc gaaaggcgcg aattttgtcg caagaaaaca
    51901 aggggatctc aagatccgag aaatgtcggg tagagggcgg ttcttttaaa taggagcacg
    51961 agttcacgcg atccggagaa tgagggttgc cctctgtctc tgctcgtggc ggcaacgagc
    52021 aaaatgggat tgagataaaa cccccgtcca agctgacctg taaagcttct tgtccagaaa
    52081 agcgatgcct aggaggtcta tggtgggacg aggaaaatac ggatgaaagc gggagaatct
    52141 gggacgaagt cgattgtgca aggacaa
//
```

#### *Talaromyces funiculosus* putative maleidride BGC

```
LOCUS       CP036239               41008 bp    DNA     linear   PLN 26-FEB-2019
DEFINITION  Talaromyces funiculosus strain X33 chromosome 17.
ACCESSION   CP036239 REGION: 406294..447301
VERSION     CP036239.1
DBLINK      BioProject: PRJNA508439
            BioSample: SAMN10522600
KEYWORDS    .
SOURCE      Talaromyces funiculosus (anamorph: Penicillium funiculosum)
  ORGANISM  Talaromyces funiculosus
            Eukaryota; Fungi; Dikarya; Ascomycota; Pezizomycotina;
            Eurotiomycetes; Eurotiomycetidae; Eurotiales; Trichocomaceae;
            Talaromyces; Talaromyces sect. Talaromyces.
REFERENCE   1  (bases 1 to 41008)
  AUTHORS   Li,D.C. and Chen,J.Y.
  TITLE     Talaromyces funiculosus under extreme acidic stress
  JOURNAL   Unpublished
REFERENCE   2  (bases 1 to 41008)
  AUTHORS   Li,D.C. and Chen,J.Y.
  TITLE     Direct Submission
  JOURNAL   Submitted (13-FEB-2019) Department of Plant Pathology, Shandong
            Agricultural University, Daizong Street No. 61, Taian, Shandong
            271018, China
COMMENT     ##Genome-Assembly-Data-START##
            Assembly Method       :: HGAP v. 2.3
            Genome Representation :: Full
            Expected Final Version :: No
            Genome Coverage       :: 120.0x
            Sequencing Technology :: PacBio
            ##Genome-Assembly-Data-END##
FEATURES             Location/Qualifiers
     CDS             complement(396..1271)
                     /gene="tfL12"
                     /note="TauD-like alpha ketoglutarate dependent
```

```
                     dioxygenase"
     CDS
complement(join(2538..3117,3172..3697,3984..4099,4290..4401,4452..4486,4538..4787,4843..5005,5300..
5838,5895..6128,6178..6457,6514..6740,6793..7335,7383..7502,7605..7623))
                     /gene="tfL11"
                     /note="ABC transporter"
     CDS             8096..8674
                     /gene="tfL10"
                     /note="Thioesterase"
     CDS             complement(join(9585..10196,10254..10304))
                     /gene="tfL9"
                     /note="Hydrolase"
     CDS             join(11126..11585,11648..11976)
                     /gene="tfL8"
                     /note="Maleidride dimerising cyclase"
     CDS             complement(12662..13216)
                     /gene="tfL7"
                     /note="Isochorismatase-like"
     CDS             join(13802..14137,14193..15452)
                     /gene="tfL6"
                     /note="MFS transporter"
     CDS             join(15903..16086,16137..16639)
                     /gene="tfL5"
                     /note="Phosphatidylethanolamine binding protein-like"
     CDS
complement(join(16890..17228,17286..17792,17852..18103,18165..18545,18605..18886))
                     /gene="tfL4"
                     /note="AMP CoA ligase"
     CDS             join(20267..20384,20443..20559,20611..20900)
                     /gene="tfL3"
                     /note="Phosphatidylethanolamine binding protein-like"
     CDS             join(21442..21653,21722..21919,21996..22745,22808..22890,22959..23044)
                     /gene="tfL2"
                     /note="Alkylcitrate synthase"
     CDS             join(23385..23712,23767..24911)
                     /gene="tfL1"
                     /note="Alkylcitrate dehydratase"
     CDS
complement(join(25299..25912,25957..26064,26132..26204,26258..28863,28929..30159,30213..30952,31009
..31455,31512..32174,32243..32324,32376..32474,32541..32672,32728..32737,32801..33085,33142..33573,
33645..33697,33751..33951))
                     /gene="tfpks1"
                     /note="Highly reducing polyketide synthase"
     CDS             join(35138..36275,36336..36552,36603..36672)
                     /gene="tfR1"
                     /note="Cytochrome P450"
     CDS
complement(join(38187..39081,39137..39281,39350..39494,39599..39991,40064..40153,40218..40235))
                     /gene="tfR2"
                     /note="Transcription factor"
BASE COUNT    11223 a   9268 c   9201 g  11316 t
ORIGIN
        1 tactcgatca gtccctcaga tcacgtcaaa tgagccttgc gaccgagaaa ggtagtgcat
       61 aattgtaagg aatctagata aattcaaaac aacgtaaata ggagacaggc caaacaggcc
      121 cttttgggaa tggtcgaatt ggcctgatga tactttgtga gcaaatcgac gttatttctg
      181 tctcatactg tagtaggaca aatgaaaaca aggcgtcaac accagtgtca gagccatttc
      241 acttagaatc cgccaacgca aagtgggaaa tggctggatg ttatgctgct actggcatga
      301 cgcgccgttt taaggcccta atccggataa tccctaaaat aaaaagatgt acaacttcaa
      361 actgaccgtc gaaagagaaa agcacacagc gtctagcata cataaatcgg tgtctccata
      421 ccagtgcccg tacgatgcat caccctcgcg tcgttgccct cccatccatc tctcctatgc
      481 atcgtgcaac gattatccca catgacaata tcatgcggtt gccacacctg cttgaacacg
      541 tatttgccgc tgttaatgta gctgaaaatc tcgttcaaga tcgcagtgct cgtctccagc
      601 gacaggccca caatccaatt gcgctcggca ttgaggtatc ccacatacac ggctttcttt
      661 ccagagtatg ggtttgtgcg cacaatgggg tgacgaacgt gtttccagag acggaaatct
      721 tcctcttcag gcttttcctg gccgggacgc agacggccgt agccgtcgta cacgatatcg
      781 aactggataa gacgcccctc gatgatcttg cgggtggatt ccgggagcgc gtcgtacacg
      841 gcatacatgt ttacccagta cgtgtttccg ccgatgcgtg gcacttgcag tgcgtgcagg
      901 atttggccgc agggaggctt gtcaaagtac cagctatcac tgtgccattc gaggtcgaca
      961 ctgccgagag tgccgatggg cttaccgtcg actttgacat tgctaatgac ggtgatttct
     1021 tcttgctctt cgtgcccgat cgaagcgctc ttgacgggga catggcggcc aaagaggttg
     1081 gagaagttta catgctgctg ggtagtgatg tcgtagccac ggaaacgcag cacgccgtag
     1141 tcacgccatg cctgcttaac ggtctctact tggcctgggg gcagatgctc aaagtcgaag
     1201 ccgacgatat cggcgccgca gctggcttcg atgggaacca cgctgactgc gctttcgccg
```

1261 ttttggacca tggtgtcttt gcgagaaggt ctcgaccgat ctgtacccgg gacgttggct
1321 tttcttgctt taggctttga aagggttcac ttagtccata ccaggtaata aggttgagaa
1381 tatgcgtctc aaacgactaa ataccactgc tgtaagaaaa ggagttagaa tggtggttcg
1441 gtagggctaa tcaggcagct ctacggactt ttgaagataa ttagaattct ctggatggat
1501 aggatttcca aatacgatga agaggtacgc agtatcttac ctaggaaacc agatagctct
1561 tataccattc tcatcaacag agagatcatt cagccccatg gaaaaagagg tatctgcttg
1621 aaaaatagcc cgattcagga gcccaagctg tggattttgc ggatttaaca atataccaac
1681 acccttaagt ccgatataac tctgattatg attctgcata aatagctgat ccggaagtgt
1741 atgtctatgc atggccgaga ccgacttgtg cttgtccaga aacgagccgg tcatatgaag
1801 ctctaaagcc ggtgatcctt accgaaaaac acaaagcgta caatgtgtct agataattaa
1861 aagggcctat attcaagaga catgtattca aaaaagcaat attgtagaaa atgagtagtc
1921 tattgatgtt tctaaatatc aaaggtagac gctgttcatg accagcatcg gataatttcc
1981 aggagccagc caggagctgc tgttcaacag ggaaaagaca atgtatattt tcagctgcac
2041 caaagacttc ttggttcccg tatgatgaag acagtatcaa gattgtatag gtcccgttaa
2101 cggctttatc accgatatag acgtgtttgc tacttcttga tttgcgtggc attgacctcg
2161 actccgttga tttaagttca agtaagcttg gcttacgaag aatgcataat gtaatatata
2221 cacttcgtac agtccaacat gagcgtagga gtgttctgtc gacgttccaa gcttgcttac
2281 ttagcaacag cgtcctggat cggatctatg agctcgttac gaggcatgac gcgagaagaa
2341 aatttgacga catccaacat cttcccatcc atcatgatta ctaggaaatt atcattaatc
2401 ctgagactca taagcatggc aaactttatc caagttaagt agaagttgaa tgtgcaagtg
2461 gtgtatactt tataaatcaa gtctatgttt cgcgcaaaag ccccaattat tccgcattcg
2521 atggtcatta aagtttatct tccatgcata ctcttgaaca atgattctgt cccaagtagt
2581 tcctggggtg gaccaaactc tacaagccgc ccgctatcca agacagcaac catatccgaa
2641 tccatgatag tgtccagtcg atgcgcgact gtaataatag tatgctcttt gaactcagtg
2701 cggatgatct tttgcataag ctggtccgta gcaccgtcaa cattactcgt cgcctcatct
2761 agaattagta tcttgctttt ccgcatcaag gctctcgcaa gacagaaaag ttgttgttca
2821 ccatgagaca gtggttgagt tttcatttga gcgtcaagtc cacctcgaga gttgatggtt
2881 tctagtagct ccactttga tagagcttcg ataataacag cgtctgagac agactttgat
2941 ggatctgcgt tgacgcggac agtctcatta atgatgaacg gatcctgagg aatggtgacg
3001 aggcgggtcc gaatctcttc tcttttaact gtctggagat ctaagccgtc gatgagaatg
3061 gtccccgagt ctaagtccag aagacggagc agagctgata ggagagtgct tttaccactg
3121 tgcaaacaaa cttagcgtca ctgaactcat tatttgagca tgtatacata ccttcccgtt
3181 ctaccacaga tgccaacctt ctgcccaggt cgtatcgaca tggatatatc ttgaagggtc
3241 ggggtagccg atggtcctcc gtatgaagcc gttacattct tgaattccac tgcaccctga
3301 gatggccatt caggcggagg aaccactgtt tcttgtggtt tgttctcaga tgccaccgat
3361 gattcaaagt ttttcagtcg cgcaatagaa cccagagacg tctcaagctg tgtccagctg
3421 gtaatgagta ctgtaagtga ctgagtaaac cctaaaacat tgttcaaggc aataccaatg
3481 gaagcaccgc ttgtagtgga gtttagcttg accgccaatg aaacaacaac gactgccatg
3541 accgagacaa ctaggttgag aaccagggtc aaccaccgtt gaatgcaata aaggagatag
3601 taagggcgtt gactgatatc cataagttgg gtacttgttg tgagcgatgg cctctgccag
3661 ccaaaggctc taatcgaagc cagtccctct agggtctcta agaaatgtgt atagactggg
3721 cttctagcct ccagatctaa gaaacgaagt tgccctgaat atcatacgtt agcattgttt
3781 gcctaaccaa atgaaaggaa gtttccgtac gagatgtgcg gagatagatg agctgcaaga
3841 agtaaatgat aaccaaaaca aatggtattg ttattgccat gaaagctgag ccttgtgcaa
3901 taagtcccgc ttgtgtgatg cttgagaaga tctctgatcc aattagtgga aaaacacaga
3961 ccaactcgca attaatagct tacgaataat acaagttgcg acggcgagtg gcaggttgtg
4021 atctatcaaa ccgatatcct ggctgaatct gttcaatgtc accccgacgt ctgtctttga
4081 aaaataggat tgaggtgcac taataacagt tagctttcga taaagcgttt aaagggaaag
4141 tatcctactt tattgttgtc tgaagaagag aactgtgtaa tctcgcggat gatcttggag
4201 aaatgaatat gaacgctagc ctatcttcaa attagccatc ttctccttta ttctggtgac
4261 tgaattgcaa tgttgctaat tatacgaacc atatagtcga aactctgaaa gcgacagcag
4321 caaatgcgag aatgacatag acactcatat acttggcgat gtcaccgcca ttatcattac
4381 tccaccattc aagccatacc tctgcttcaa ttagcatctg attcttatag atccttgggg
4441 aattcgctta cgcggaaagt acgtggcaaa ggcaagcaat gcagcgctgc agaaaaagaa
4501 agtagcactt ataccgctaa tagacctgaa atagtaccta tatactgcaa ggtcgccggt
4561 tttacgggtt aggtccgtaa tatcatctag tgtgacgcct ttgatctttg gccttttctt
4621 tgtagatgga tcggaagtgg tcagttgagt tttttcttct gttatggaag gaataacagt
4681 gctgatgtat ccgtccttgg ttcttagttc atcgaaagtt ccttgctcag cgactcttcc
4741 atctttgccg aggacaacaa ttttatccgc cagctgaaaa taacgtgcta tcaatgaccg
4801 tcagcaagga gcccgtcatc ctggattatg ggctagactt acttgaatgg gtgacaagta
4861 taacggttgc ctttaacttt ttgagaatgc cgtttggtcc cagtaatcta tcaacaacag
4921 ctttctctgt cttggagtct agcgcgctca atatgtcatc gagaattaca atatctttgc
4981 gtgcatatat agccctggcc agggcctaag acgaaaagtg ttaggtcatg ttgtctatgt
5041 tgttgaaaat aggagtactt gccagtctct gtctttggcc gccacttaaa gttaggcctc
5101 gattgccaac gaagctctca tcaccttgcg ggaattgcat gatatcctcg cccaatgcac
5161 aggaatgaag gactgtgtca taccattcct cgtcttttgc atcccccccg tctaagccgc
5221 aaatgctttg cctgatactt gcattgatga tccacggagt ctgagggcaa taagacattg
5281 caattgagga cacagttacg ttaccgctat caaatggtag ttcacctatg attgccctca
5341 tcattgtcgt tttcccagat cctacgggtc ccgtgaccac attcaacgtg ccgaagttaa
5401 agtcaatgct gatgtcttgg attgcaatgt ctgcttttgg tgcaggtcgc acgcttaatt
5461 gcttgattga tacagccgcc gaagggttag taactgaggg aattcctgtt ttgttcgaaa
5521 tatcaggaag tgcgatccca tctttagaga tagaactgcg acttgaacaa tcagttctat
5581 ctgatgaatc caatatcttt ctttgatctg ttcgatttgc agcgagaaga tatttctgga

5641 tgcgttcaaa gcagccaata caagcaatag cattcggaat tgctgaaatc aactgggcag
5701 ccggctgggt gactaatgta ataattgaaa gagaagtgaa agccgtgttc gtgtcaagag
5761 agcccgagcc accggatctt gcctgaatga caaaggctat aaaggtcaga acgggagcaa
5821 agtatgccgg acagtatgct acgcaatttt tggtcagtcg taagttgcaa aggcagggat
5881 ttcatctgac taacaggcaa cattggaacc caaaaccgtc catctgtacc ctgcagcttt
5941 tttgagctca tgaatgcgtt gctcttggat gttatgagcc ataaacccgg acaggcccat
6001 catcttgaca tttttcatgg agcccaacat cgttgaggtc ataccaatcc gtcgctgaat
6061 cgcaccgttc cagatctttt gtctgttgct cataactccc gccattcgag agtttgctac
6121 tccgcagact ggtcgagcgt tagtttatag aaccaagctc agttcaaaga tactcactgg
6181 ctacgataat gattggtatt atacagacag cgccgagctt tacctccagc aggtacatgc
6241 caataccaac ttcgataagt ctactccaag tctcgttaat gctctgcatg gcaaaagcta
6301 ttctatcaat atccgtgctc atgagagtta cagctgcgga atcatcgtat agcccgtctg
6361 gtgaggtgag agttttgttg tatatgattc cgactagtgc gcctcgcagc atcgtgattg
6421 ctcggtaaag ttgctgatta taaccaacgg tacaaatctg atgtaatcta ttcaatgttt
6481 gtaatgagac tgctgaacta tggtaatact tacggcaata ccagtataca cgaagaatgc
6541 agcggcaatg agacccaggc cgttattgct attctgtctc catgtcggtt cttcaacata
6601 tgagattgtc gtactgatca gaaaaggctg agcatacgta aatccaatta acgcgagtcg
6661 gggaggtact attcgaagaa taggccacaa aagacatcga gcgcacgcca acgggaggct
6721 atagcgatgt tctggcttgg ctgttgatag tcagcacgaa aatacaatgg atgtgaaaat
6781 gaaaggacta acaacgcgta tcccatgtcg cttgcatctt ttctcggagc atttgagaag
6841 ataatgcttc gtccgtagga tagaggtcat ccaaggtcaa cagtttacgg aaacccttca
6901 agaagagcgg attgagccac caaaggacgc ttctatccag gatgcctcgc gttgcctccg
6961 gcggatatcc cttgtactgt tctttcaaat ttttggactt ttcttggctt tccaacgcca
7021 gcatcgcagt tttcacagcc attgtcagag tgaaaatgac cgcaatcgcc aaagggccct
7081 ggcggaggta cagcgttctg gcctgaggaa tgtcgaaaat gatagaagta agtagatata
7141 gacatactgg cgtcgagggt cgaacagatc gttgttgctc cagatatgac agcagaacga
7201 gaccaaaggt ggcgaccaag gacaacacag tggctgcaat ggtcgcaact gttcgtaaag
7261 ctgggttaac aagccatagg atcagctgcg ccaactgcag cccagcaagg atattggccg
7321 tgacctgcgc catgtctctt aagcatgtta ttcatataaa cagagatagc tttgatactt
7381 acccctttcc aagcatagct tgacagtatg gatgtgcttt tcacgctcga cctccatagc
7441 cagaacagcc tcaaaggaac gacgagaaga aaaatagaag ttactgatat tgataagatg
7501 ctctgttcaa ataatagcgt aaaatcccat ccttggcgac acgatcctgc gacaataggt
7561 ccgaagacct gatctgcgct aagcgggcag gcagttgcgc gcacgtcgaa agtcatctgg
7621 catgagtgaa attccgataa taaatactca ataaaaatca cgctgcacga ggttgtttca
7681 agaaatagga gatcggagaa agaatcgcga caaacgtcgg tcaataagcg tctgtgcctg
7741 tgacgaggtt gtcgttgttt tgatgatttt tgctagagaa taattatttc cggggactta
7801 agtaggggga atggcagcta attagggcgg cgatcccggc acaaaaaaac atttcccagc
7861 acctaattgg atgcactcca atgatctata tgctacgagg ccgagcatat gcgtataaat
7921 gtgcgtataa atgttgtaca taacctatgc cattcgcagt aactcacgct ttgaactgat
7981 atgttgatta tatgcacgga actcggcaag tcattacgct ctcttgtcgt actacgagga
8041 atcatatcat aacacctcta caatcgatat catttaaatc gcaggcaccg aggaaatgcc
8101 taacctgcca cggcaatttg tcccagcaaa tctcgacttg actgcgcgag agcactttcg
8161 ccggtatgct tggtgtgata caatatacga aaacccaagc ctccgaccag tcatcacagt
8221 gaaccagcat tcatggtcgg atgtgccctc gacgttcatg tggctgagtc taggggcgcc
8281 ggacgcactc ctcgcggccc agtccttctg gaaagtgggc gacacatccc cagagagttc
8341 acccgagaac agaacggagc tatggacttt gtactctttt ggaagggcag tagagagttt
8401 cttgcatgtt gcccacggcg gcttccttgc cagtcttctc gatcaacaga ccggctccat
8461 tgttatcaca caccctgtac cgacgaatcc tcgcactgtg tcaagcacta tcaaatacca
8521 caaggcattg catacccctg gtgctgtgtt atgtcgatcg tggattagca aagtcgaagg
8581 acgtaaggtg tgggccaagg ctgtgttgga agatggtact ggagcacttg tggcagagat
8641 gggagcgttg tggattttct tgcgaccgag tctatagatc aagacacaac tactaccgga
8701 tgatagtgta gaacgcagtc gggtacctat ctccagacta gctaggcacc ccttgtatca
8761 agctcacaaa gctagatcta tattacatat attccatagg tttccaaagt agatacccaa
8821 cattcgtgga caccgtaaag agtgacaaga agcattttcc tttgctcgag tatcaaaaat
8881 ctttaacatg cacctaatac acttaacgct agacatgaat ggctttaatt tgcatgcatc
8941 gtcactgttg taatacaaaa caattggaga atggactgac agttggattg gtcccttgaa
9001 taatctcaag ctaaaattga actaaatttg cgaaaacctg caggcatcaa agagaaaaga
9061 gaagcaaagg ggggttgcat gtcaagcaca acattttgta cagaccgggc cggaaccggc
9121 caaagatgct ccttagagtt ctgcagtcag gaataggcca attgcaagtt aagttagtaa
9181 caccggacac tttctactca tcctctttaa gcccggtcag aaacaacatg catcgtatcc
9241 actatgcagt atccaggacg tattttttca ctagtccttt tgaccatgca agctatccgg
9301 taccggccat tgggtacgtg gaccagtttc gatttattct agactctacg tcttacttac
9361 cgcagcaggc cggaagattg gctttgcatg tacggtcttt ttctgctcat tagaggatcg
9421 cgctattgag ttatccacag cttatcccat ctgactcatt gaggtgtagt tttttttttt
9481 ttttggcccc tttcttacag cgcagatgct cagctccagt ctccataact tgcacaacgt
9541 agatttcaaa taatattcta tgtccaaggc tcaaaaaaaa tctaaagaaa catggatctg
9601 cggctaagat ctctaatcgc cgccgccatc tttgctacca acttcgcatc cgtggggatt
9661 tcatgaccct tgtcatgtgt caaaatcgcc gatgattccg acttgcacaa attgtaaagg
9721 tcgagagaat acttgtaaac aaaatcttta ctacccatca catggagcgt cggaactgta
9781 attttcccct ctagcccttc attcaggatc ggttcattgt tgtcattcat acggaacgga
9841 gggaacgagc tgaagaatat ggcacaccga aataaaggtg agtcgtaggg gttctgctcg
9901 gcgtggtgga ccatcaatcc agcggctact gctcccccat gcgaaaagcc tagtacacag
9961 tcaaatggcc cctcttcttc gatgatttcg tagagcatgt cgtatgctct cgtgacggag

10021 tcgccgtcat cgtgtactgt ttgcggccaa ttgtagtaac tgaagaaagg tccctcgtag
10081 aagccttcaa ccccggggcc aggagcgtct tctacctcgc cttcaactga gtggaaagta
10141 gcagtgctat cttttcggag ctctctcaca aggggcgcta gctgagtgtc aagaatctgc
10201 ggaaggcctg gtaagtagtc tgcgtcaaca aaaaggtacc tcgtagctct tacttctgca
10261 tttgttcctg ctccatgcag acacagaatt ttgaggccag gcattttgtc gttggaggaa
10321 gctaggctat ttgcgctgta caccttagag tgatggatac agcagttaga ttcgctacga
10381 gagcaaagat tgaacaagtg caccggataa ttctgataga gtccactata tatcaactcg
10441 atattttaaa tacaccaaca tgtttgtaat caactttagc ggtcttgcat tttccaaaga
10501 taatgcgcca gctaatcttt tggttaatct cttgcttctt gctaggtaga agcacactaa
10561 aataaatgcc ctagattcgc tgtatgtagt ctctgttatc tgaaagagcc aatggtctaa
10621 tcccggctgt attgaatcaa atcgagaatg gataaattgc gagagaagag gctgataaag
10681 gctgacggta tcatttggac caatgaggaa agccccatgc taaaaatgat cttaccgttc
10741 acatttactg gactgcatgg agctattatt acataaaact caagtacgcc gaacggtaaa
10801 ccgaaaaaag aaatcttagt catgacaggg gctcttttct tccaatcaat cccggtgtaa
10861 atgtatggcg gcaagttagc agagtatcct acggagtcat tctagcaaga tcgtcttaag
10921 gacatatccc gatatccggt cggttcaatc tgttcgcaaa gcgcaactta gcttcaatca
10981 cattattgaa agtaaacaat ttgtattcta gctcccataa acatatcgat cttggagtat
11041 attataatta ccagagctac ctccagtctc gtctaccttt tcaccacaat caacccaacc
11101 tggaaacttc cacaacaccg ccaccatgat tgccttgaac actctcctcc ttagcctgcc
11161 attggcattc tccaaaatat tcacatccgc cacccctttt actactcaac agcccgtcgg
11221 ttgggtgcat cccgatattg agtttgccga gcgagatggc ttcttctttg agaacccaga
11281 gacagactgc aaattcgtct cgcagccata catctacaag aaattcaaga ccctggagac
11341 cgatggtagc gtcttgttct ccatgattaa caaagatgtt cactttacca tcgtcggaag
11401 tcatcccggt gccggtgtgt accatgactt gatgcacttt tacgtcaatg ccttgagacg
11461 agttgccctt gttgctggta ctgagcaccc agaggctttc cgtgtgtacc ccaaggctat
11521 ccatggcggc tgtgacaccg agtggtctgt gcaggaaatg aactttcagg gtatctcgaa
11581 tgctggttag tttccttgtt actgtgtgat tatagagatg gctctgatgc taacaatcca
11641 acgcaaggaa ctccttttga catcatcaac gtctgggtta ctcgctggga tgcggagaag
11701 aagcaaatgg ttgaaatccg tacttacatt gatgccatga aagtcaccca actcatccac
11761 gagagcgaag gctggtggaa cggttcccgc cacctccatc actacgaatg gatgcccggt
11821 ccatacggta tgcccaactt gaccgagctc tatgctctta tgcctgaaga ggaccgaccc
11881 aagggacgcg gccctggtga aggccttact ggtcaggtcg tcggctacct gcttcccgag
11941 caggaggaag ccgctcgtga gggctatgtc catgcttaag acaactatag ccgaatttca
12001 cggttttcaa ctacggaaac agcgaagcat gtcatgattg cgatcgacta acttgcgtga
12061 gatggttatt agtgatgaac tcattaaatt cttgggccaa aactgaatca aatttactgg
12121 aatgtgtttt agatctgagc tttgttggaa gaatgttgca aaccgattta ggtttttggg
12181 gtggaaaagg gggtttttgg gactaccaag attttgaaat tagcaacgag accgcatcga
12241 taaatgtttt tgagatttac tttgtgattg aacaatataa gatatggagt cgaggaaatg
12301 gttgcagatt tatcttcatt aaaactgagt agtcagtacc tacatagata gtccccttca
12361 atgagaacat ttaatgtcat aaccgcgccg atgttgctac aaaccttgat tcttccaagc
12421 ttgtcatact atatctcact cagcatgcgt actgctaatt ggtctgctgt tcccagcata
12481 tatcgtatct ttctaggtag gtactcgata caaaatgctc tccgaaacac tagccacagc
12541 acatgtcaac aggattatgt atctctaggc aagcactcaa agcgcgaaga tttacagtgg
12601 tatcagctct attaagacaa attatgtaca ctagagaggc tcaaagatat taaacgtctc
12661 aagcggccaa cttctgcaac cactcctctg catctagaac ctgtcctctc ttgctgaata
12721 ttttggtgac aagagtttcg tgtagtgccg gatcaccatc ggcacacaaa ttcttgagaa
12781 ccacaagtcc aaagtcctta tcggatgcct cgcaaacagt tgccagaact acccctccag
12841 tgctgatgcc cgctagcacg agcgtctcaa tgcccaaccc tttcaacaca aggtccagac
12901 cactgccggt aaaggcactc acgcgcttct tctcgataag tatatcgccc tctttaggtg
12961 caattgctgg atcgatttga gtttctggcg aaccgctgac aaaggagttg gttttgacag
13021 cggcggcaaa ggtagcattt gaagccacaa cctcgggatg tccagggcgg aaagcgaccg
13081 tcacgtagat gactttgacg tgaggacgag ctgcatcaat tgtcttggcg aggcgttcaa
13141 ggtgatctga ggccaggggc atgcgaccaa caatgccggc ctggtagtcc attacgagga
13201 gggctgtttt ggtcatcttg gcggctgtcg gtgttcaata gtctgtttct ggaattaggt
13261 ttaaaataag ccggtagata ttgggggcga caaatgtgga tgtgctgcca gcaggttaga
13321 ataatgagct gaacctgagc ttcgatttaa gactgtccat ttgtgagggg aagccctgaa
13381 ccgggatgct tttacggccg gtaacggagc aaccggtccc tgtcccggcc gggcagtttt
13441 gtatctctgt ttcttccacg gcaatcaacg accgggtccg aaccgttagt tatgatcatc
13501 ttctatcacc gaaaacattc tgccatatcc tagaagtcga aatattacct cgctcaggcc
13561 ggtttcttat ttcggtaaaa taaagttggc aacggccatg aagtgagaaa aacattcact
13621 tggtccatct agttagatag ctacgagtga atcgacatct agattcttct agctagtgtg
13681 aattagacta tagagttcat aatcaaacag tggcccagaa gttcatcacg tatagcaatc
13741 tatagactga ttcaagggtg agccaagtgt acccccatat tggatcaaca ctccgagcaa
13801 catgtctcgc cgactcaaga gcaatggaag tcttgccaat gaccctcatt tctctggtaa
13861 caatggcata gctgttcaag cagcaggcga agtccagccg ggtcccaaac caatagcagc
13921 tgtgaccgcg gaggagacaa cacctttgtt gccgcagaat actgagcagt attcaagctt
13981 ctcgacagct cagaaaacct tcattatttt cactgctgca tttgcgtcta cattttcgcc
14041 attctcggcg aacatttatt atccagcgat aaattcgatt gcccaagatt tgcatgtgac
14101 acctgctatg atgaattata ccatcactgc ttacatggta agtcaattct gcagatgaca
14161 tgaaggtaca gccctcatcc ttagaatcac agatctttca aggagtggca ccaacattta
14221 tgggaaattt atcggacacc gtgggtcgaa gacccgtgta tgtactctgt ttcggcatct
14281 acttgtgcgc aaatatcggc ctcgcccttc agcgcaacta ctgggcctta ctagggcttc
14341 gtgccctgca aagcactgga atcagtgcca caattgctct ctcaaacgca gtggccgcag

14401 acacggtcac ctctgccgaa agaggcacat atctgggaat tgcttcgcta gggggtatac
14461 ttggcccagc gttaggtcct acgttaggcg gattgatcag taaattttgg gcttggtatg
14521 gaatcttctg ggttttggcg gttctctccg gctccgtgtt tctcttgatg cttctcttct
14581 ttccagaaac atgtcggcat attgttggaa atggctctgt tcctcctccg ccatggaatc
14641 ggtccttaat caacatcata tcagactacc gcaagcaaaa ggcgggtatg gacatggagg
14701 acggccatct tcggcgtcaa caattagctc agaaaaggcg tattcgattt cctaatccac
14761 tctccacttt tcgcctctta ttccagctgc ccacaagttt ggtcctattt gtaaatggaa
14821 tgttattcgg cgcatactat gctattacat ctagtatacc ggccgaattt gatgccattt
14881 accatttgaa tgatctacag atgggcctca catatatacc aattggccta ggaacaatac
14941 tctcttcttt tacaaatggg tgggcagttg attggaactt ccgaagaatc gctgctagga
15001 ccggtggctt gcctcccatt aaaaatggca aacaagatct cactgagttt ccgattgagc
15061 gttcaagact ccaaatcgct attccatcag caatcgcagg tgctctctgt attggaacat
15121 atggatggtt tctgcactac gaaatgccgt tatgggttgc gatattgttg ttgtttttga
15181 tcggctattt tatgactgca agctacaatg tcatgaattt gctgattgtt gatctgaact
15241 atgaggctcc agctacggca acagccgcaa ataactttgt gcgatgttcc attggtgctg
15301 gggccactgc aggtatcata cccctgctgg attacatggg tcgaggccca agttacacga
15361 tactggctgc aatatgtatc agtgttaccc cattgtcgat ggtggtttac aagtacggac
15421 tgcactggag gcactcaaag gatcacgggt ggtagcttag aaagaaggat agtacacgga
15481 aggggacatc ttaacttgcg atgttcatag cgcatctatt ccttgttctt tctcctatac
15541 tatcagaggg gttatgattt tatttatata gtgtgctttc aacgatgaaa agttttagat
15601 agcagtagca agggtaaaat cgtggaaata gtgcatttta cccggaaaaa tatatatcgt
15661 atattttcaa tgctaatgtt taaagttaat agaatttgaa agtttacatc ccttcttctg
15721 gaatccacat tctaatatgc cgagtctcag tgttgaatac cgttaatgca tccaatctaa
15781 ccgacaacgc ttttagaaaa tgtgttatat tccacccgcc gattttccgt tctgcgacat
15841 actcaatctt cttctcaacg ataggtatag ctcttggttt cagtgtcttt gaaaattcca
15901 ccatgatccc acaaagttgt gtacgaaaaa tcggggcaat tccgtttctc cttcatatgg
15961 cattgttttc aagctatgct tacgcccaga ctccccccgag ctataccccta tctacgtcaa
16021 acagtctgaa tgttacattc aatggaagag tttctattta cgctggccag tctctgaatc
16081 catatggtaa gcaagccatg aatcttggtt atacagcagt gctgacattt ctgtagacgc
16141 gatgttcatg cctactctgg cctataccgg cttagatccc ttcgagccat atatggcatt
16201 catgatagac gtcgaagtcg ttcactccgg actcgcgtac cctctgctac attggtacca
16261 gccagactta tgggcggata catcaaccga cgcgtttatt ctacgcaact tgaccaacaa
16321 tgcagcagca tacgttggcc cgcaaccaaa ttctggccct agccattcat atgtggttct
16381 gttattccga cagccgttga actataagtt cccggattgt tttcaatata tgctacctct
16441 gagcatggag gctagagcag gatttgatct tcaccttttc atggagatgg ctggcttgaa
16501 agaattggta gctgcaaact attttacatc ccagaatccc gagagtcggc caaccacaac
16561 ctcgctaatg aagcctccat gtgccacagg gaaatttgcc ggaaatagcg agactcggat
16621 cttgaagtct gttctgtaag gcaatgggag acgaataaaa ctaacgacat tgattacgca
16681 gaatttcttt gagaaagact gttatttggc aaatatagac aggcgtactt cattgttttt
16741 atcaagtatg gatttaacct ttaaagtcta gatactagac tcggttatct atcgtcaatt
16801 tatgcccatt cacacagctt cataactgac aaaaataaaa ataaaatacc acaaaatatg
16861 tgatcaaata catgcccaag gtctatctac agtctagctt cactctgtct gcttctctgt
16921 ctctcacggt ctctcagaat ccttcgcaaa atcttcccgc tcgggctttt tggaattata
16981 tctataaact caatacctcc tttgagccac ttgtaatgcg ccttctcttg cttaaagtgg
17041 tcgtgaattt cttccatcaa ctcggcgtca ttttggaaga gttggcctct atcatgaggt
17101 gcgagcacca caaaagcttt tgggacctct ccagcggcct catctggcac tggaatgact
17161 gctgcatcag caacaagagg gtggctgatc agacaagcct cgagctcggc tggtgcaact
17221 tggtgtccct aaaggttggt ttataggtta agacaaagac cacgaggacg gacggaaata
17281 cgtacattga ctttgatcaa ctccttgatt ctgtctatga taaataggtg ctcgtttccg
17341 tcgccgttcg gattcttgac gaacatggcc tcatctccgg ttcttagcca ggcgtctttt
17401 ccgctaccaa aagtttctgc agtggctttc tcgttgttca agtatccaag attggtaaca
17461 ttgggtgctc gaatgagtaa ctctccagtc tttccatatt cgtctatatc tgagccatca
17521 gaaagtgaga tgatgcgtgc ctcaacgcct ggggcaaggc ttcctgaaga tcccgggaag
17581 atatcaagag ccgaagtcat ggataccgct gaagtcgtct ccgtcatccc gtatccttga
17641 cagatatacc aagaagggta tttttgaga agtgcgcgat atgtctcgac tcccaagggt
17701 gctgcgccac tgaagacctc actgacagaa ctgagatcaa attgatccaa acactttttg
17761 ttattcagta tctgaataat aatgggagga acctaggaaa aatttagatt agtaaatgga
17821 ctgaagtaac tctaccttct gataaactta caataaagag gacatttatt ttgaatcttt
17881 caattgcagc cgctagaacc ggcaactcaa accttgggag aacaagcaca gagtctcctc
17941 ggtagatact gatatggcaa atagagacga taccgtatat gtggctttga gggagtagtc
18001 caaggaccac tttggtctgg ccgcgtcgac gtgctggtga ctcatacgca gccacttgga
18061 taacattaac tatgacattg taatgagaca ccatcactcc tttctaaagt atgggttagt
18121 atattccata tttaaatgat caatggcttc aagaagtaat ataccggtaa tccagttgtt
18181 ccactggaat acataaggaa tgcacattgc cgagcaccct gccccttctc ccatcggagt
18241 ggctccagtt ctggcaaacg agatccctct gaaatgagtt gatcaaccgt cttcaagtca
18301 gacggcgacg tgaaaccaag ctgcttttca taattcattg acaatatata tatatgtttc
18361 ctcgcaatgc cactggcatt cgcggccttg agtgccgttt ccagtaatga agtacaggtg
18421 aaaatcgctt ttgatcctga agacttcact tgatgcgtca attctgaggc actgtatgcc
18481 gagttggcag gcgatgagac cccggagaga ttgtgtacag cccatgagag cggcacactg
18541 tccaactgta tttcgtcgtg tcaggtacta gcatcttgaa acatttccct acttcaagac
18601 tcacagtgtt atgcgcaaaa attgtaatta gtttgtccca ttccgatcct ttattcggag
18661 accagttgaa ttccttggct aaagctctag caagatactc cactcgtcgt tttacctcca
18721 gagacgtata atgagtacca gacagtccac atacaaatgg agatctgctg tcgcgtagct

18781 ttgttctgcc atacttctcg tcaagcatga aatgactgat gggaattgag tctggtacag
18841 tctccaagtc gacatttggt acccatttgg gttgtttgaa caccattgcg acaaaaaatt
18901 tggggctgca aaatttccta ttgacgttct gggcttaatg gtgaccagga acacaatgtt
18961 tatcagtcta gtggagatcc tttcttcggt attagagcaa caatcaatct gctgcatgac
19021 aggctaggtt agtaaaattt tcccctttac tttcgttgtg ctttacctta cataaagtca
19081 gaaaggaatc gtcgaattgg gtcaatgtgt gctcgcgggc attgatcgga tattcctaag
19141 cctcggtggt tgaatgaaca ccgggagaaa tccgaaaatc tgctattcac ctattaataa
19201 aaatgggtaa ctactcggcg ggttatttac aaacaagcac gctggggctg aaagaatgtt
19261 tatgtgtgtc tatttttctt tcagcaagcc ttccaggaaa gttactatgc cctatcgatt
19321 caatgcagaa ccatctctga acatgagcag acctagacct atagcttctt agtacaaata
19381 tttgatagta aatatcactc agtgtgagct cattgggtgc aagtatatgg atacggccga
19441 cctggacgag aaataccgcg ggctgattga gcgctactac aattcccaga aaagatatac
19501 agccattaca tagcttcaac tgcataagtt ctcattgaga ggccatgctc cacctgtctc
19561 taaaattgaa gtgcttctgc catgaactgc gcatcacgtg acagacagaa accctaattc
19621 gtatacttac atagaggagt tcgtattaga aacccacatc acgatcacat cacgatgtca
19681 ttgaaattgc tgtgactaga agataattct gttttcaagc taagagaaac aagacaaatt
19741 acactgtgct ccgcggcctt catctccggt gcaaatcgcg ctgggggcca gataactcaa
19801 tatgcaattt tcgggtatcg gctgcttttc ggccaatttt tatgtgctgc cagccccagt
19861 aagatttacg aacaaagtag ctatctaatt agtaacgatt gtctaataca tggaaatcaa
19921 agcttggaaa tacgtgtcta tattaaaatt ttagttgcca tatgtcccct taattttcgt
19981 cataaaattt cagggtcaaa cagtacgtaa cactcccatt cctgatctcg cctgttgacc
20041 tataggagta tgtaaatata aagtaactat agtccaatcg tgtagagttt ctattgatag
20101 gaagaagtga atgctaatta tcaagcaaag ttccccttac tcaacacttt cggatgtgaa
20161 aagtaacctc gataccgccc attgagatac cgaggaccta tatcacaacg tagaatccga
20221 aaaatcgaaa cagcactctt tcttcacttc ataagataag tgcaggatgt cttggtttga
20281 taacgtgaaa actgctctaa atcttcccaa gaaagattcc gagaccttgg gattgacatt
20341 tgatgaacgg caagtcatcc cgggagaata catccctaag agtggtaagg gcctctgcaa
20401 tgtgtcaata aacgatatta agttaatgca atatctgaat agaggcgcaa ttggctccgg
20461 atctgagttt caatcaattc acaggtacat atcttgcaat ctgtatcgat cttgatgctc
20521 cattcccttg tttcagcttt ttgggaccta ttcttcattg taagagtctc taatttcgtc
20581 atcttaacag agtatactga tatattacag ggattcaatc ggatttgaag ccatggactg
20641 ccgccgatgg caccataaag ctaaaagcaa gcacgccttt catatcagac tacgttggcc
20701 ccgcaccacc gccgcctagc cgctcgcacc ggtatgtctt catgttatat gagcaaccgg
20761 aaggatttga ctataccaag tacgcacccc caaacggcca gaagatggga atgtggccgc
20821 ggatcagata cgatctaaaa gcatttgaga aggaagcgaa gctgggtccg attgtggcaa
20881 gcaatttctt tcgaagcaat tgaggtcaat gatttttcct atttctacta agaaaactat
20941 tgatgatata aagttcaaaa cggatagctg gcaatcatat aatacccta gcttccaagg
21001 tgtatatgat catattttta gcctgagcct tgggtgactg tgggtggttt tcggccatct
21061 gatacattcg catagatgac ttctccagaa atcgaaaaac cctgagatag tttgtaaaat
21121 tagtcagcct cggtccaaat agccaataat atgtctcaaa ataagaggaa aagagtgtgg
21181 ttgggtggtc caatgtttac tcgtagtaca tgcacaaaag ttgccggacc tttttgcccc
21241 agcaaataat tcgccgtcaa catagggtaat aaatacattt gtttctgagc tgctctcctt
21301 cctcagtagg cacagtagtc tttgtcagct ctgattgcta atgagctact cggttattta
21361 ttatgactct ctttgctgta tatgaagtgg aagatttgct ctttgatttc tggtatctca
21421 gtactgtaag caccattcat catgtctgac ggaaccttat tcatccaaga ttctcggtct
21481 gggaagaaat atgagattcc cattcgacac aatacggtgc ttgcaaccga tctcaagaag
21541 atcaaagctt cgtcagtagg agcaaaccgg gccgacaaag tcgcagatgg tctccgtttg
21601 tacgatcccg gccttgaaaa tacgactgtt attgagacca gtatgacata cgcgtaaggc
21661 ctgacttccg gtgaacgaaa atgagaaatg aaaaatttct taagattgat ttttcaatta
21721 gggattcaga tagaggatta cttatgtttc gcggctatgc tctggagcaa ctctgggaga
21781 gtgatttcga ggacatgctt catttgatgg tctggggcaa atatccaact ccaagtcaga
21841 gcgaatcgct tcgcaaagac ctagcttcgt tgatgtcaaa tatccccagc actgtatttg
21901 aggtcattga agactttccg tacgtaaccc gaaaattttg tgatggtatt actaaggctg
21961 tgaggagtat gaaatgctga taacccgtgg ggcagacgcg actgccctcc aatgcccatg
22021 ttagtggccg gtcttgcagc ttatctatca aatgatttgg attccattcc cgctttcaac
22081 gggggtaata tctatcacgg aaacgttgag aaaacggatg aagctattct caaaactgtt
22141 gcggcatttg catcagttgt gggcatcgcc agtagtcacc gaagggagat taaatttacg
22201 ccgccctcat tagataaggg ctaccttgat aatcttttca caatgatggg agtagttgag
22261 ccgacaactg gatcgccttc tcctgacaaa ctagattgct tccgccggtt taccattatc
22321 aacacagatc acgggatggc gttgtctgca ttctcacatc ttgtcgcaac ttcagcattg
22381 gcagacccta tctctggcct cattggctct ctcgttgctg cttatggccc tttacatttc
22441 ggcgccccag aagcggcata caagaccatc agaaacatcg gaggtcccga aaacgttcca
22501 gcatttttag aggaagtcaa aagtgggaaa aagaggctct ttggttatgg acaccgcaca
22561 tacaaaaccg tggacccaag gcttgccccg atcaagtccg cgctacagac gctggatgtc
22621 ccagacgatg tacctttaaa gacggcctac gagattgatc gtcttgcagc aaatgacgag
22681 tatttcctaa agcgaggtct ccatgctaac gccgatttct acactcccta ttgttttatt
22741 atgatgtgag ttacttacaa tcttcagtct cattaaccca tgacactcta ctcacattta
22801 ctcgtagagg atttgagccg gaggagtttc cgattgcaat gttcgcacaa cgaattatcg
22861 gtataatggc tcattggaga gaagcgatgc gtatgtatac ctatttcatt taattaagtc
22921 atggatttac tacagtgcta attcctctca tgatatagtt cgcaaggtga aattgttcag
22981 gcctacgcac gtctacactg gagaaacaga gccagtggag catactaggg tttcatccaa
23041 actgtaaaat gataacctca gtcgaagaag tgtaaaagct gtaaataatt ttcattacac
23101 tgtttgatat tatacgaggg taattcaatt gttgcgatct ccgatccagc acaccggcaa

23161 cttttgactt gtacctacat cgataaagaa cgcaaactgg ccgattggtc attttgattg
23221 gtaagtactg ggagacggga atgctaagtt actttctata gagatcggta ataaattgaa
23281 acgtgggtat atcatggata tattatgaac agacatcttg aaatattgct tttcacttcc
23341 tctttttttgc aagtttaaaa aaaaaaagga tcgaaacggt aacaatggct tctttgcagt
23401 atgaccagat tattgttgat atcaaagatt acgtttttca ccacaaagtt gactcggaga
23461 aagcatggaa gaatgctcgt atagctcttc ttgatgcaat cggctgcgcg attgaaacag
23521 ttttcaaaag tgaggactgt agacggatgt ttggacccat agtgccagga tcaacaattc
23581 caaatggatt ccgactgccc gggacatcat acatcatgga ccctttgaaa ggctcatttg
23641 atatgggaac ggcaattaga taccttgacc acaacgatgc gatagctgga gctgattggg
23701 ggcacccttc cggtagatca catatcattt cccaactgac tacggtgcta acgtatacac
23761 caccagataa cctaggagca attcttgctg tttctgattg gctgtgtcgc tcttcaaaag
23821 agggcgtaat ttcgcataat gggcctcctt tgaccatgaa aacgattctg gaagctttga
23881 tcaaagcgta tgaaattcaa ggatgcatgt tgctacgcaa tgctttcaac gcgtacgggc
23941 tagaccacgt cattcttgtg aagttggctt caactgcggt cgttagctgg ctgatgggat
24001 cgacggaggg ccagacaatg gcagcgatat cacaagtctg gatggatgga caagctcttc
24061 gagtttatag acaaaagggt aacaccatac ctcgcaaagg ttgggcagct ggagatgctt
24121 gcatgaaagc aactcagctt gcacttttga cgaaagccgg gcagcccgga tctccgacac
24181 cattgacaat gcctcgatgg ggtttctatg ccaattcatt tggtaacaac tcttttgatc
24241 tcccgaagaa gtactcaagt tgggtgatcg aaaacattat attcaaagtc atgccagtgg
24301 agggtcatgg tgtaccatca gtcgaagctg ccatgatcca cctcaggacg ttgaaggcta
24361 gacaacttag tgcccaaaag gatatctccc aaatcgtcat tagaaccaat gcagcaaccg
24421 acatgataat caataaaaca ggggaactta gcaatgcagc agaccgagat cattgcttac
24481 aatatctcat tgctcttact tttctgaaag gcgacctacc agaagctgaa gattttcttg
24541 acagcaatat gtggtccaaa agctcggatc tagacaatct cagagacaag atcaagattt
24601 tcgtggatga gcgcctgaca aaagattata tggacctgga tgtgaagagt gttgctactg
24661 gaatgacact tatgctgtcc gatggcacac atctaagcga agtactcgta gaattcccaa
24721 ttggccacgg aaagaatcca aagacgcaag atgttgtcga gcaaaagttc cagaaaaaca
24781 tgaaactcat gttctcgcca gaagaaatcg acaatataat tgaaatggtt gaacataagg
24841 acaatgagga gaaacctgtc tcggagtttc tggatttgtt ggtacgcgag agctctttat
24901 cttctagact atgataatta gactattagc caagggttct aatattagaa attttatata
24961 cctagttagt gactgtagca ttgaattcaa tctgcatata acggaaatta gtgctcaaga
25021 tgtcgctaaa ataattcccg aagttcttta tctatgattt gcgtgagctg taaagtgtct
25081 aagtatttgg tgaatggctg ccggattcca gataatgtat ttcaagtaaa tctccggtaa
25141 cagatatcta ttatattcta ttcatgtacc aagtcatggg atttagagag tgagatttgt
25201 atagctgacg gtatttccag gtttataaag caaaagctga ggcatcaata ttcaaagcca
25261 aaaagacgaa ataaaataaa tcaaaaacaa tcaatctaga gcttctcccc ctcgttcaca
25321 tgaaccaacc tcgatcgctg agcaatcttt gtagccaact gactaagggg cgtgttggcc
25381 aaaagttcta aaataggaat tgtcgagttc atctcttttg aaatccaatt acggatttcg
25441 acagcaacaa gtgagtcaat gccgtagtct gaaatggggt tgccaacatt gacatgttca
25501 acgcctacgc cagcacgaga tgcgatcttg ctaactagag cagtgcaaat gatgtcggac
25561 gcatcttcca aggttgttgc tttgcgcaag ttctttcgca attgaccggc agatcctgcg
25621 cctccagaat cagtggagcc ttgagatgtc agactgcgga agtgcgcaaa taaaggtaga
25681 ttgagagagt cgagagaatt gccctcattc catactccga gaccagtcac acactgagag
25741 agaagcgctg gtcggcgcgg attctcgata gcaaagtgaa gaaggctcat gagcgtcttc
25801 ttgtcagtga gttcgaagcc ttgtctttgc atagcctctc taagttcgga gtttgtagca
25861 agataaccga catcagaaat tgcaccaaga tcaatagttg tcgcaggaag gccttgggaa
25921 tttcgatagc tcgcaaatga atccaaaagg tgttaccagc tgcgtaagca gcttgtgttg
25981 cattgccaat aattccgctt accgaggact ccatgacgaa gaaatccatg tctttgggta
26041 gagcgttgtg gagattccat gttcctgata caatcgatca gtaaaaagat cttaacttaa
26101 gcacagtatg agttttgaat ccattacgca ccttgcacct tcgggttcaa aactgcttta
26161 tagtcgccga gagacatatt ttggaataag atatccttgt aaaacttggt cagtatcacg
26221 taaaaagaat tacatgaaag aggatcaaaa gacttacacg cagtaccata gctccttgga
26281 taactccttt aattgggggc atctcttgac tagcctgttc gacagttgac tgcatttgag
26341 cagggtccgt aatatcacaa ctgtaaacgg caactttgac tccattggac tccagagctt
26401 gaacagtatc tttagcatcg gcattgtttc gaccactacg actaaccagg atgagattag
26461 aggcgccatg ctcagccatc caaagagctg tagcacgtcc gattcctcct aatccaccaa
26521 ctaacatgta agatgcatct gagcggagta aagacttggt attgtctcca ggtatcactt
26581 tgaccatatc ttcggggcca ttgatggcca caagttttcc catatgacca ccggtctgca
26641 ttgtgcgtag agccttttcg acatctgaga tgggataact gtgaacttgg gctggtccac
26701 gaattgcttc agtcctgaaa agttctatta cctcggcaat gaccttgctg gcgacttgag
26761 gtctctgttc aaatagatca tcaagcaaga atcctgagaa agttgtattt ctgctcagtg
26821 gcttcatctc aaggcgagta ttgatcatga tatctctttg gccaagctca acaaagcgac
26881 cccagggagc aatacagtcc caggtaagcc gaaggttctc tcctgcaagc gagttcataa
26941 taacgtccac gcccctacct tgtgtcattt tcttcacacc ctttgcaaaa ctaccgtttc
27001 tgctgaagaa aatatgatct tctggtatct tgaagtaatc catcaagaac tgtttcttat
27061 ctgctgtacc gacagttgca taaacctctg cgcccactag ctggcatagc tcaataatag
27121 cctgaccaag accgccactg gcggcatgca ccagaacagt ttctcccttt tgtaactttg
27181 cgatataatg gcatgaataa tatgcggtcg cgtacgttac cgggagtgac gcagctcttt
27241 caaaggacaa atgatcggga atcttgtgga aacgatcaaa cggacctcga taatcggtgc
27301 taaagagtcc ggcaccgtag gaaagaaccc tgtctccgac ctgcagatga gtgatgtttt
27361 tgccaactgc ggtgacaacg cctcccgcct cccatccgag acttttgagt tggatctgac
27421 ctaaagcagt catgacatcc ttaaagttga atccagtggc tttaacttcc atctggaccc
27481 aatccgctgg cagatcttcc tcgaatctgt cgtcgtccac aaaatagatg ctgtctagca

27541 gccctggagt gccgacatgc atgcgcaatg gtcttccagg ttgtctgaac ttttggagct
27601 ctggcttggg ctcagctgtc gaggtagata caaatttcac aagttcagta tcttcgataa
27661 gccttggtat catgagaata ccgtttctct cagcatattc aggctcaata ccttttggtg
27721 ccgtaccaat tgaaaaatga cgttcgaaca gagagatgat aagctctgct gttcgtttat
27781 cactcagcat cttttcggcg tcaagatcca atgttaaaac tggcctttca ccgctttctg
27841 agcgtactgt tcttgcgata cctgagatca ggttggtact agggtccgaa gacaaggcag
27901 tagctccacg agtaacccat agaacaccac cactttcgag gaatagggtt ttcaaaacat
27961 tccactgctt ttcagaagga ttaatcaagg tttgtgaggt tgaatcacta agcaaaatgc
28021 aaacttggtc cttcagtggc gtagcagcct caaatgaagt gatatcagca tctatttgat
28081 gagctataaa ttgttccttt agaatctgag cggctgggtt tccgtcgccg ccatccaaaa
28141 tcacaagagt attgggatac tttggagttg ttggcttgga agttagagct gtcaaaaaga
28201 cggcagcatg tttggtacta tttgctgcat tgggaatggt gacgtcctga gaataatgag
28261 tttctgacaa aacctttgtc cagttttcat cagaaaggat gatatcccca tcctctgaat
28321 gtgcttgccc tggtagcata ctgggtagaa atccgaacat cgttgaaaca gcgagagact
28381 tcataggtcg tccaaccaac agcaatttac ctccaggctt gaggagtttg tgggcattct
28441 ccagagtcct gcggactgct tcttcacaac caagtggctg gaagacaatc aaaacatcgt
28501 aggactctag agaaacatct tgctttgcag gatccacctc gatgtctagt tctttgcgtg
28561 taaccaagtc gttccacttg gcacatttcg ccttcactaa gttagcaagg tcaaagtcgt
28621 catctgcata ttcgaatttc gagaagtaag gtagctgttg cttcttagat tccaatgctg
28681 caagaattgg aatggaaaga aatccagaga gcttgccaca tgccaagata gagaggttag
28741 gttgtttgaa agccatcatt tcaaggtatt tagacaccat cgtgctgtcg gtgaacatat
28801 cagagggact ataaaatcca gataggtcaa cttcacttaa taacgaagat gcttcgattt
28861 cgcctaattt atttttgtca gctttccaaa tcacgtaaga gaagatagca agtgatggct
28921 tatcatacct ttcagtaagc taggcaatcg ctcgccaatg tgataaaggg tacggccttc
28981 atgaccgtat gattgtactt tgtcgagaag atcattttgt tcatcttcac tggctgagat
29041 ccagatttca gatagactac ccttattatc gtcgatagtc ttttttacca taccatttaa
29101 cagagcccag aggctctgct ggtatggcgt tttgatggca gacagactgt tttgcatgtt
29161 atcaatgact gacttgagaa gataaaacgc cgccacttca aatttatgac gaatgtttat
29221 gtgacgcact accggtgggg accctagtaa cttcgcgaga ccgtcagcag agagcatgtc
29281 aatatccgct ttccaatcga agttataagc acggctttga ctctcggacg acccgtctgt
29341 ggcttctgct actccaagtg tggtacaatt gattccccga atggtgataa caggctcact
29401 ttccgcttct tggtgagcac ttagaacagt gagagacgct gaaatagtgc gtggatcttt
29461 gatctgggtg gaagtgtagc aaacgagctt gtctccaggc gtagaggtga tgctattgga
29521 aacataaatt tcctccacag acaccggcac cgcaggatct ttcatttcgt cggcatcaag
29581 agagacgaaa atggtatgga aaaggctatc cagtgtagca ggatggataa taaatgagaa
29641 ttgaaacttc tgcggcatta cagcagcagt gtcaggtatc gtgatatttc caacacaagt
29701 attggggcct gctcgtgcct gtgacatgca agcaaatgtt tcgccgtact caagaccaag
29761 cctggcgatc gtttggtaga attgagcgac gtcgatgttc ttctggcatt tctcgtcata
29821 ctcattaatc attgaaacaa gctttcttct gtcgtccgcc ttttgggatt ctccgtcgat
29881 cacgtttgct gattttggag gtgtttggat ggctatgagt ccacgacagt gctctgtcca
29941 gcggtcctca ccgctaacag atgacacgat aaactcgtcc catagatctg acggagaagt
30001 catgctatct gaataggact tcaaggtcac agacacttca acctcacctg gattctcagg
30061 aatgacgagg gctgctccga tgacaacttc tcgaatgttg tagccggcga ttgtcggtac
30121 agacctctga atggcccgct gagtcgctgc ttcaatagcc ttataaattt attagtatcc
30181 gtaatttgaa gcagagaata gaaaagacct accatgacaa tgtagcctgc tgcagggtag
30241 acaatattgg cttgaataac atgatcacga acccagggaa tctcagaaag ccggatatgg
30301 ttcgcccaac gaggttccat tgggctacat gttctctcca aagcaccaag cagatctgtt
30361 cttggaaatg gcctattctt atacgctttg cttttccatg attcatgcca gtaagtgttg
30421 ctgtgattcc aagcatatgg cgggaggtca actagtaccg atagtttctc tgtgctagaa
30481 atgcgattga cgcgcgacag attaaccggg tagccaagag taaaaaggga agaaactagt
30541 tttaggacag tatcgactgc gttcttgttt cgcaggagaa cagaatggta tgcgatggag
30601 ccggagctga gtttctcgtt tgcctgaatg atctgcttaa ttggaccagc gagtgccgaa
30661 tgaggcccaa cttcgatgat atcatcaatt ttagattgca ggttccgctt ctgtgtcttc
30721 tttccagatg atgtatccaa gcagagctgt tgcaatgact ggctaaactt gacttcattg
30781 agcatattgg aaacccaata atccggcccc agatcgcatg cagcagcctt ttgtccagta
30841 acggacgaaa agaaatcccc ttcagtattg gactgaactt tgatctcaga tatagcttcc
30901 ctgtactctt tctcgacctc agccatatga tgtgagtgat atgcaacctc aactggcgaa
30961 gaattagtat taagaagctt aatcaatttg tcatccaaga aaacatacca ataagctttc
31021 gtgcaaatat cttctcttct tcaagaatca atagtaattc atcaattgca gtctcatcgc
31081 cagaaatggt aacgcttgat ggactgtttg aacaggccac cactgctttg cctttaacca
31141 atttggcaag gtatggaagg gcgtcctctt ttgaaagtcc aacggccatc attgatcctt
31201 tgaaaatgcc tttttgcgca aggctcgttg atgcaacacc acgaaagtat gcaacacgca
31261 tggcgtcttc taggcttagt gctccaatag cgtaggctgc agcaatctca ccactggaat
31321 gtccggtaac ggagtccggt cgaataccaa aagacgcaag taggtctacc agagctatct
31381 gtacagcgga acaaagaggt tggctgtata aagcacggtt aatgctagag cccatgggat
31441 ccctcgtaag ctcatctaac agatcgaagt taacgtatta tcgcctgaga tttacgggaa
31501 taatagcata ccaacaacat cgaatggtgc cccagcagct ttgagacaat cggaagtctt
31561 ctcaacagta gcccgaaaaa gaggatacgt tgagatcaac tccttgccca tgccacacca
31621 ctgtgcacct tggcctgtga agacgaatcc caactttctc ttctttgtcc caccgctgct
31681 gaaggtcacc ccgctaccta gagactgggc caagtccttt gttgaccttc caactactgc
31741 tgtcttccaa atatggtggg ttcttcgctc attgagggtg taggccaaat tcttaaggaa
31801 ttcctggtcc acatgatctg cgcggtcttc gaggtatcgc tgaaggttct ctacctgtag
31861 tttgcttgtg tgctcgtcga agctagagag cacgaagagg cgcgcagagt cctcttcagg

31921 taagtttcga gctccgttct gcacttcctc tatgtggcca ttaccattag agtgaccgtt
31981 tgcagtcccg ttggagtgcc cattgctgag cccatttgcg tgtccattgg tgataccatt
32041 ggagtaacca ctggagtgtc cgttcgtgta ttctaacttc ggcgccaatg aagttaggga
32101 tttggttgtt ctataatatc cagagtatcc ccgtagtttc aaataatcct ctgcactctc
32161 aaggataaca tgggctgttt cgatttagtt aataagaata tcaaagtatg gacaagcaaa
32221 tcggtgtatg tgcaccacat accaattgct ccaccgtatc caaagctgtt aacagaggcc
32281 cgacggggta ctggtccttc ccaaggctca atggccaatg gaacctgcgg cataatttag
32341 cgatatgcgt ctttcctata tcattcagcc catacсttga gcttccactc ctcgagcgga
32401 attctggggt tagctttctt gaagtttcta ttcggtagat agaccttgtt ttcaagcata
32461 agaacagatt ttatctaggt tttcttgtat cagcaatctg cataagtaga aggctaccga
32521 ggaaatagcc aattacttac cacacctgcc acaccgcttg tggcttcaag gtggccaacg
32581 tttgttttca ctgatccgat acgtataggg ctacttgagt cccggccttc acagaaaacc
32641 ctggagactg ctcgcgtttc aaggggatca ccctacgaag atcgtgtcag tatttaccaa
32701 taggtaatca tagcatttta tacgtacggc ttgagttctg ttcagtatgt cagcaatgat
32761 ctttcatcct caaaaacaaa taacagaagt cagaactcac ccagttccat gacactcaac
32821 atagctagtg cctaggggat cgagcccaga cctttcgtag actcctctta tcaatttttc
32881 ttgggccaca ccatttggca tggttattcc gggagttctt ccatcttgac tggcaccagt
32941 gccgcgaatc actgcacgta ctgtatcacc atcgcgcagt gcatcactaa gaggtttgag
33001 tagcaaaacg ccaactcctt ctcctcgcgc ataaccgttt gcgcgctcgt caaaagagta
33061 gcatcgtcca tcgggagaaa gaaatctgaa ggaattcagt caatttccta ttccaggaaa
33121 gaaaagcatg tataaaccca cctcatcatg ctcatggtga cccaggcctc atggctcaga
33181 atgacattca cgcctgctac tatagacatt tttgcttctc cagtccgcag agtttgacag
33241 ccgaggtgta aacctatcaa actggttgag catgcagtgt ccacatccac actctggcct
33301 tgcagatcga agaagtagga caatctgtta gcaaagtttg atcggcactg tcctgaattt
33361 gtgcactgat acattggggt tgattctgga tctctccaaa gaatatcagt gtagtcggta
33421 gtaaaggctc cagcgaacac tgaggtatca ctaccacaga attgctccat ggtaatacca
33481 gagttctcca tggcttcgta agtactctcg aggagaaatc tctgttgagg atcgagagct
33541 gccgcttcag cagccgacat ctgaaagaag ggactgtatt ttgaaagtca gaatcatcta
33601 gcgacttggt cttatcgctc aactaggtct gaaggctgac ttacgcatcc cattgtgata
33661 catcctgctg caggaaatgt ccgccagtga cgttgtactt caaacgttta gattatttct
33721 caaaaagctt gggcaacgta gttaactcac agttccatgt ctattagcat cagggtggta
33781 gaacgcgtca tggttccacc tattcttggg aatcttggac caactctcgc gaccctcgga
33841 tatcatcttc cataaattct ctacatttgt tgcatcacca gggccccgga atcccatgcc
33901 aacaattgca ataggcatcg aggcatcttc ggttaattct tgcctggtca tgatgaacgg
33961 aatagaacca aacagacgta gagagagttt attgaagaag aaagaaagta aaacaagaaa
34021 gaagaaattt gggtgagttt atggggtatt ataaagatca aatataaggt atggtgttgt
34081 tgctcgtcca ggcaggaaga cacagctttc acaccaagtg ttagaaacct tacgacgaaa
34141 ctacaaagat tgctcttaga gaagttcttc ttggaataaa atatattcaa tgttctaagc
34201 tgacttatcg tactagaagt gccgaaatcg ccggtttgga tgctgggaaa gctctagctg
34261 agttcgcgaa gactttcact cagtcaaaca aagtcatact agctttaaag tcgccgatag
34321 actccaatat cttgtgtatc gaggggcttg ttctattata gtcagcagct attagggtaa
34381 ataagtgacc tataattcta aaataactgt agcacggcca gaaatatctt tctggggctc
34441 atattgagga ccaaaacagg tatacagctt cggccaattg cggacaattc ctatacttgt
34501 agatacacgt tagatagctg cttagtttta ctttgcgtcg aaatataggt actttatctg
34561 aatagtatgc gaaaatcggc acggctcaaa gtgaccatgg cattaacaat ctagacgtac
34621 gtctccagaa gggataacat tacaagaatt gttaataatt gttccacaaa ggaatattta
34681 cggccacagg ctttgatatt ctcctaagca actgtaatta gtaatgcgga ttcgttcacc
34741 tgataatcag cccggtctac catacatctt cggtcaattt ttgtttctgc aaggcaatcc
34801 gtgcttcaaa accgaaagct ccctgcccat caggccagtt cactacctag cctcaactcg
34861 atacaagtat ttgtataact ttatgttgtt cctcggtatt ttattactgt cttgtgacac
34921 taaaatttgc caaacttagt aatagtcctg gatgtgtgta gttgaagttg tccgggttat
34981 ttaaattgta ctagagaatc tgtttagttg gacggatcag atcatatcgc aatgaaatga
35041 aatcttctat aagtcaaaaa tatcttaatt ggcagaataa tcgctgattt taccattttc
35101 tctatttcaa cttgggggcg tacccatcat atccaatatg gcactactcg acgcggtctc
35161 agctcatgta gccatatccc cccatctcct ggttttacta tgctgcggca tggtatggct
35221 catttttcaa tccatctcat atgcgacaag agtgcgaaag tacaacaaga ttatttgtgc
35281 aaaaggttgc cttccttgca agacctatcc tcacaaagat cccatcctag gactcgatct
35341 cttcattgag aaccttcggc ttttgaacaa gggcgggatt ttagagaaat tttccgagcg
35401 ttactaccaa caaaatacat ggacgtacac ccagcttttg ttcagggaaa aagtcatcaa
35461 tacagctgat cccgagaata taaaggcaat tctcgcaacc cagtttacag actttcagca
35521 ttctccccgc cgtaaagcag cgttttatcc cactttcggc catggcatct tcacaactga
35581 cggagcagaa tgggaatttt ctcgagccct cctacgtccg aactttgtga gaagccaggt
35641 aggcgatcta gatattttcg aggcacacat tgcgcgtctc atcgatcgaa tacctaaaga
35701 tggatcgatt gtggatatac agcatctatt ctttgcactc accatggata cagcgacaga
35761 atttctattt ggtcagagtg ccgatgtgct tgtggaaggg gaatcaagcg tgaggggaga
35821 aaagtttgcc gaggcctatg actatgtgac agaaatcgtt gggattcaag ccaagcttgg
35881 tcagatcgtg gccaagattc cgaacaagcg ttacaccgat tctatcaaat atatccacga
35941 atatgtggaa atgtatgtac agaaggctct tgatattgca aagtctggtc aggacggagt
36001 aaagaacggc agggaaccgc accagaaata cgtattcttg gacgagctgg cgaaaacagg
36061 agttgacaaa aagaagatcc gggacgaact gttgaatgtt cttttcgcag gtcgcgatac
36121 aacggcaggc ttgctgtcat tcacattcta tatccttgca cgccgtcccg atgtgttcga
36181 gaaactaaga gctgaagtga tgaccttagg ctctgaacgg ccgaatttcg aagaaataaa
36241 gaatatgaag tatcttcagt acactttaaa agaaggtccg ttcatttggt ctctcctcta

36301 tagacaattg atgctaatac tctccggtta aatagtcaat cgcctctgtc caattgtccc
36361 cgttaatgct cgtgccgctg ttcgcgatac cactctgcct gtcggaggcg gtccagatgg
36421 gaaatctcca atcttcgtca agacaggaca aacagtgaac tatcaaatat atacgatgca
36481 cagacgaaaa gatttgtatg ggaggacgc cctagagttt gtgccggaga gatgggagca
36541 catcaggcca acgtaagctt ttttgattct gatcttcctt tagttggcta atatgacaat
36601 agatggcaat atcttccatt caatgccggt ccacgtattt gcattggtca acagtttgcc
36661 ttgactgagg cgtgagtgta ctccttccaa atctctttat ggtagattaa ctcatcggta
36721 gttcatacac cataattcgt ttgctacaag cattcaagtc tataagaccg agagaaggag
36781 aaggatcttt gtcggaatta cttactctca ccacggcggt tcgtggaggc gttcacgttg
36841 gactaacacc agcatagacg cagttttctt atcgtaaggt agttttcacg accatccacc
36901 tggtttatag ccagtaatct aaaacaagcc taagtgtgtt tctgttattc tcgatgactg
36961 tgatgctgaa cttgtttttg tctatataaa ttgcaattga tgctcgatgt tattgttcca
37021 ccttccaagg ctgaaaaaat atcagtatac aaaagtgcac tagatgcgca ggaaatttgc
37081 agttgtatcg acatgtacag cagagacata attgttctca tctaactagc gtttctcttc
37141 caagttgctg atctcttatt gcagtttcat ggtaacatcc taaaatggtt gaccagccct
37201 gggcattcga cgttttttaa aatgctgact aatggtttag agatcaccac aactttcaca
37261 ataggtagtt gggcttggca aaagattcgt gtttgatgtt gcatgtggcc ggcaagtttc
37321 tgcgagtatg aggaagcgtg cgaatgcaat aattcaccat aaaatatgtc gtctctaagc
37381 ccggtcccgt tggggtccaa aatcctcctg aaggattctt tgcggatgtt ggggcggtcg
37441 caatcggtct atatgtaaga tcataagatg cattagtaat ctatttaatg ctaactaact
37501 aataagttac tagactacat gatgtagacc aacattaacc ggagggcacc ctgttgtaac
37561 gtcccaatct tccaaatggg caaatcatga cgtgtgttgg ccctagtcct cgtgtcgaag
37621 tcgcttattt tacgtacagc aagactacaa tttgagatca ctcatccacc ttgccacttg
37681 ccactcatga tcggcaggct gaatcggctg ttttctaaca gaaatcttag ctacgtatat
37741 ccggaggctc ggaggtagtc aaccgccggt gctaactaca ggatctgtcg atgttgacaa
37801 cgacgtttgg aggacaccac gttagtagac attgctgtag tgtccctaag aggataccag
37861 tcactatcgg ccgtaccggg tgcatgcagc ggtgactccg aagcgaagct ccggggaccc
37921 gttagttcct atatatgaca tcttatacat acgtagcagc cactcagctg agatcagaaa
37981 gcatatcata atcacgtttt atccgttgag agtcttatct ccgatgcaga gctagtttct
38041 ttatgcagcg aatctgttct tctagttacg taaatttcat ttgtcttcgt tgagttatgg
38101 acttgtactc gattgcggta tggacactgc cctcccttgc ttgaagtaaa tagaaagttc
38161 gatatctcaa aagccagtaa cagtttctaa atccgtattc cttccaaatg gtcgataatg
38221 tcatataact tggaagccgt tttcatcgac ttcctcatat ctatttctga cgcagcattc
38281 gaaacatcac tttcaaaggg cacgaatgac agttcgcgtc tattggccat gcgcaaatct
38341 tgcatggccc gaatcaaatc ttctaaaacg ttcgacagcg caaagagacg acgccgcaac
38401 aaatgcatca acacaaattc tctatcgtca ccaacaagtt gatgccggcc aaacagaata
38461 cgccaattgc ccctttctgg accaccagtc gcgctcatta ctggctgatt ctgttcaccc
38521 acgactgatt ttggtggaga aggtaatttt ttctgggttc catcgacgct ggaaccatta
38581 gacccattac gttcatggga catagactgt cgagaactgg gcatgaattg tccgtttggt
38641 tgtgagggtt gaagccattc cagacggtcc agcatcgaga atgccccttg ttgtttcgag
38701 gatggtggca aatcattgta atctgagaat tcaacattgc ataattgctg aagcaagtct
38761 accgcgtcgg ccattgcatt tgaaagcaat tttaaggttt ctccctgctg gaccttgcat
38821 gcagtacact gcacggcaat tgggatggtc tctcgcaact gcgcctcaat attaagtgca
38881 agatccatcg ctccagaata ggtgaatgta tttcctttcc tgatgcttaa agtgatgcta
38941 agagcagact caagacaacg acatgcattt tcctgggggg gcggcgcagg atcgtttgga
39001 aactcccatc caaatattcc ggtcatatcc ccatcgtggt cagtgtgggt gaatgttaat
39061 tggccatgtt cgatttgttg gctgagggtt ttgtcagcat cactttctag gattctctag
39121 agagcaggga atataccttc gctgcttgtc ttcatctgag tcatccaaac tttgccgagc
39181 agatgagtct gttcggtgtc taacggtatt tgtcattgaa gatgacaatg atgagtgcat
39241 tgtagggaag tcgatatttt caaggcatga gttaaagaag tctaaagaac cgtcaatttg
39301 taagatcaac tctcgagata tccatgaatg tcaaggcaag acaacctacc tcgatcattc
39361 aagtctgtca ctactccacg aactggagaa aatacagaac tctctatgtt catactttgc
39421 gtacgtgcaa taataggtgt catcgtatta tttgttcggt ttctttcttg gttcacaaca
39481 ccaggcgcgg ggtccttatt aacaagccag atcataagtt agtaaactgt attggcaagg
39541 tgcaagggca aatctttcaa catgcatcgg tgatcattgt atcattttcg acgtcaacat
39601 caacaccgcc ccagtccaca gcatagaagt gacttggcgg ttgtccaaca acactagcgg
39661 tttcactgtt tggctgtgat actatcgaga tacgttcctt gggtcgcctg atatgtgaag
39721 aggaccccgg tgaagttggt tgcgcaaggc gtcgtggtct gcctatgcgt cgagaaacgc
39781 tataatggca ggggatattt tgcgccaggc atctcgcaca tgagggtttc tcacgagaac
39841 atttggtctt gattgcttga cacgcatcgc aggtttgtcg cagttttgat gtgttgttcg
39901 tcgtggaggt gttatttgtg gtagaagctg acatcttgga tttttgcact atcctacata
39961 gaccgtggga ggaagtgaca cagctagaga gctaggatcg agattgataa ctggataatc
40021 agaatatata gatggagggg ccgtcaaggt gatctacacc tacgtgctcg atatctgatc
40081 tgcatggagc tccaacaata ccactctgag tcctccagta atcaggagca cacggcttat
40141 cggacgctaa aagctatatt gtagtcagca cggtgcagtg gtatttttct gcaatcgggt
40201 tggtgaatgt caattacaac atttaagggg gccatttttta tcactctatg tcctttgttc
40261 ccgtctacac catactcttt tcacgcattc cccactggca tccagcatta ggctcacgag
40321 gcgttgatga gttattcaat cagatatgac agcaggttat agtcacatgg gtgtttaccg
40381 gaaggtctcg gatggaagag cttgatgaag ttggcattct tcggtcatta gcctaccaac
40441 ctcatgattg ctttgatcta cttggaacgc ttctattaat ttactttgac cgaagaaaag
40501 cttagcaatc tcgcacgctg aatacgcccc aaatgtatgc acagcactgg tggttcgtct
40561 gtccgccaag catatttcac gtaagtgaag atgtggatcc cgaaggatgc aaagtcagtt
40621 tcgaatagtt agtgagattc ttagcgcttc tactagaaaa aaaaagtttg tgatctacta

```
    40681 gtacgaggct tagaagctgg ctattatctg ttccactctg cacggtcgtg tcgaagagct
    40741 gcaaccccct cttttatgcc actttgggaa ttgcatgcac gtaccaagta tcgcatagtg
    40801 tattgtagcc agtcggagtt cctctgtgtt ttattccgat catagatttc agctagatct
    40861 gcatagttcg taacagtaag aaatgcctag gccttgtcga acatcctgac agccaaccgg
    40921 aaagtttacc accaagggcg atcgctagcc ttccattatc gtcgcttact ttctgttctg
    40981 cactaatgaa taagctttac gccggaca
//
```

## Ts1R1

```
LOCUS       ABAS01000011            3691 bp    DNA     linear   PLN 22-DEC-2008
DEFINITION  Talaromyces stipitatus ATCC 10500 gcontig_1105507293323, whole
            genome shotgun sequence.
ACCESSION   ABAS01000011 REGION: 762106..765796
VERSION     ABAS01000011.1
DBLINK      BioProject: PRJNA19557
            BioSample: SAMN02953686
KEYWORDS    WGS.
SOURCE      Talaromyces stipitatus ATCC 10500
  ORGANISM  Talaromyces stipitatus ATCC 10500
            Eukaryota; Fungi; Dikarya; Ascomycota; Pezizomycotina;
            Eurotiomycetes; Eurotiomycetidae; Eurotiales; Trichocomaceae;
            Talaromyces.
REFERENCE   1  (bases 1 to 3691)
  AUTHORS   Nierman,W.C., Fedorova-Abrams,N.D. and Andrianopoulos,A.
  TITLE     Genome Sequence of the AIDS-Associated Pathogen Penicillium
            marneffei (ATCC18224) and Its Near Taxonomic Relative Talaromyces
            stipitatus (ATCC10500)
  JOURNAL   Genome Announc 3 (1) (2015)
   PUBMED   25676766
  REMARK    Publication Status: Online-Only
REFERENCE   2  (bases 1 to 3691)
  AUTHORS   Nierman,W.C.
  TITLE     Direct Submission
  JOURNAL   Submitted (02-MAY-2007) The Institute for Genomic Research, 9712
            Medical Center Drive, Rockville, MD 20850, USA
REFERENCE   3  (bases 1 to 3691)
  AUTHORS   Fedorova,N.D., Joardar,V., Maiti,R., Schobel,S., Amedeo,P.,
            Galens,K., Inman,J.M., Galinsky,K.J., White,O.R., Whitty,B.R.,
            Wortman,J.R. and Nierman,W.C.
  TITLE     Direct Submission
  JOURNAL   Submitted (01-OCT-2007) J. Craig Ventor Institue, 9704 Medical
            Center Drive, Rockville, MD 20850, USA
COMMENT     Assembly name: JCVI-TSTA1-3.0
            Genome coverage: 8.09x
            Annotated scaffolds were added in December 2008.
FEATURES             Location/Qualifiers
     CDS             join(562..901,988..1532)
                     /note="putative enoyl CoA isomerase"
                     /gene="ts1R1"
BASE COUNT     1039 a    823 c    764 g   1065 t
ORIGIN
        1 gggggtatct caaggccaat atcacaggtg gccaagtatg tccttgagat aggagacaaa
       61 gtacagccat taactgccag gaatatcgtt tgccttatga ctactaatgg gatcggagct
      121 tctattgaaa ctccaatcaa aatttcacag ctacccaccc attgctgcat accttgctga
      181 actgactaga aaatattccg atagcatatt acgaactccc caacgtggtt gtggatcctg
      241 accccagccg tcgatataga agtcagacta gatacagaaa tgcccgaagc tgacggtctc
      301 acaatatact acgctccatg cccagacaat tttcttgaga agggagataa aagcccgatt
      361 ctttggttga gcctggcatc tagtcaaagt atgtaggcag aaataccaca tagcagaggt
      421 tatttccatg tgttgcttcc gacggctgct ctaacctcga aaagtatatc ctgtgtagat
      481 tgaacgtagc cagtatttaa ttcattttgc ttaacccttc tgacctttca tattgtcttt
      541 tcttcaaaaa cccactccat aatggctgat agattgaatt atttacccac aaacgtgcat
      601 ataaccttcc cagatataca ctgtatacag atcgaattct catacggacc agagaatgac
      661 ttatctgcgg ccgtgttatc atggaagcgt gatcttttcg atactctcgc agagtcttct
      721 atcattcgaa ccgtgatcat cattgtggga aactccgatt tacagagttc tcatcaggaa
      781 aaaattgtat tagagggact ggccgcatta tcgaacacat caggcaaggg ggactctaga
      841 gaggaatgct caagtgaatc aattcagaaa tggcttgata gtattcagag gctgtccaaa
      901 tgtatgtgtt tcaaccttgg aataagacaa agcccatctg caatgcaaag aaccaactta
      961 ttgactgact gtttggataa cccgcagtct gcgtatgcat tttgcaaggt ttctgttcta
     1021 ccttcgagtt gcaagttgca catgtctgcc acgtgcgaaa ctgtactgct gatctccaac
     1081 tgtcgtccac tgttactaca gtaaacccta cttttgaaga gcaggcaaag aacgttctta
     1141 ctccgcccca gttaatcaca tcagaactca acgccaaaca aattctctcg ggtctcaaca
```

```
     1201 tagtctggga cgtccggcac tccaaattcg agactatatc tcaagcattg aacctatgcc
     1261 gccaagtatc ttccagaccg ggaccagtcc ttttttcccg tgaaagaaga agcatccggt
     1321 caagtccttg tgtcatcact ccaacacagg ctctagaagc ttcggatctt cgtctcaaaa
     1381 acgtctatga tcgtgccact actacttctt ctactgctag tagttttact tcagcccca
     1441 catcatacgg tgcgtacgag gaggcatcaa ttgcggctga tctggatcaa attttcggtg
     1501 ttggaagtgg gatcaggcag tcgaagcttt agagcttgag gtgattatca gggtttcgtc
     1561 ggatgtaacg tgatgaatct atccttttga aagctacaat aagagaatct tctagtattt
     1621 ccttgttacg tatgtgggtc atgatgtaga aagagtgtat ctccatatta cggcactttg
     1681 agatgaaggt tagttagtaa atgtaagatg atgtcgagga cttcaaatgt ggaatccggc
     1741 aacaaatgtt ttctacaaat gagaaagaac ataattcaca cacagttcta gcttcatata
     1801 taggtagtag aacaagtcta aaccccagac ataagccaaa gcaaccagta attgttgact
     1861 ggttactgag taccgttatg attccctgtc tagatttggt cattgtatag cctggtacta
     1921 ttcctttcgt acggtgcttc atgattgcat gcgtccgaac acgagtattt cgagtcagta
     1981 gcattggtca caaattactt ttcccgttca tccgacgggt cattctgcaa agtcaccacc
     2041 aacctctaac cataaacgtt gcaatccagt cttgtcgggg aatgctgaac ttcccggata
     2101 actctctatg aaatccatct gccaaggagc tggaatatgg gggcgaaacc ttctttttat
     2161 acggatttcc tacttcgcaa ctgactaact atctctgatc ataaagttag tcgatcagct
     2221 ccaccatttc ttcaatcaac catctaggtc cttccaacgc gggtactttt tccttcttcc
     2281 tgcaggattc tatacaagta gacggggtga agacgggacc gaacaagtaa tcggtgtctt
     2341 ttgtctcagc ttcccattca agttttgtac aacacctatg tgactacgta gaacaatgac
     2401 ccgaagctga tccgtcgttc aatcggcaca tttaagcata actccacttg atttcagttc
     2461 ttacaagcta caaacaacaa gaggccatgg ctcagagcta gaaatgattt aatcaaggat
     2521 gttacttagc ctgaagcggt ctccaccatc tccatcctgt gtatatagtg tcgaatgtct
     2581 gttaatccct attcaacctt gtattgtatc aggtatccta ttattacgag atcaatgtta
     2641 atgtagctta caactatatt aggatttcat gtaagaattg tcgcgaatcc ttggtaaaac
     2701 cagccttttc tctgcttatc atactggtac ccacggaaat ctactaccgg taacatacga
     2761 tatgcgaaag gattcggatc ctagctctga tcctaaacaa acgatctgag gcctggctag
     2821 ctagtacagt acgcacatat caattcgaac ttaaagtagt aggtactgta tagagatctg
     2881 tactttaatg tttagtagta ttcaactgga ttaaggtcaa gctgggtcag gatcatcgcc
     2941 cgcttttggt gcggtataaa agaaagctta taaatattgg cttaactata aaaactggcc
     3001 cattagacat cataatgctt atcacttcat tttcttaaaa cgttcataga atgacagcct
     3061 tggccgtgga atctggctga ccgaagacct acaggagatc tcagatgctc gtatccttcg
     3121 caaccaaaac caaaggcgta aatatcgagg aagtttccgc acctaatctt taagtaaaag
     3181 acatttgatt ggtggaaatg agcaaggaga gccaattatc attatctttc aaagttaact
     3241 atattgagcc gggtttgctt gcattctctg gcgaatagat tacgccagct cggtccccta
     3301 tattagtaga atgctggggc cgctgattag tcgtccttat cgttccgact cgctgtacac
     3361 gtgggaagac attgattggg ttgcttctag ttcatggtcc atacactagt caattggccc
     3421 taagcatctc tccagttgtg gtctttagtg tacgacgggg ggccaaatcc aagtttcctt
     3481 gactcgtcaa taaggatcga gctaggcgca gtagtttaat gttggcggac agggattatc
     3541 ttgtttttgt gtctgtaagc tcaattaaaa tgggtactcg tagcttcccg tcccattttg
     3601 aatgatttct tcagcgagag aaacctcagt caattgagat cgacaggaga tgcggatatc
     3661 tatatggtca attggaatgg ataatacagg a
//
```

#### ZopS

```
LOCUS       LC516887                2278 bp    DNA     linear   PLN 19-MAR-2021
DEFINITION  Diffractella curvata No. 37-3 DNA, zopfiellin biosynthetic gene
            cluster, complete sequence.
ACCESSION   LC516887 REGION: 36649..38926
VERSION     LC516887.1
KEYWORDS    .
SOURCE      Diffractella curvata
  ORGANISM  Diffractella curvata
            Eukaryota; Fungi; Dikarya; Ascomycota; Pezizomycotina;
            Sordariomycetes; Sordariomycetidae; Sordariales; Lasiosphaeriaceae;
            Diffractella.
REFERENCE   1
  AUTHORS   Shiina,T., Matsu,Y., Ozaki,T., Nagamine,S., Liu,C., Hashimoto,M.,
            Minami,A. and Oikawa,H.
  TITLE     Biosynthesis of Fungicide Zopfiellin
  JOURNAL   Unpublished
REFERENCE   2  (bases 1 to 2278)
  AUTHORS   Ozaki,T., Minami,A. and Oikawa,H.
  TITLE     Direct Submission
  JOURNAL   Submitted (08-JAN-2020) Contact:Taro Ozaki Hokkaido University,
            Department of Chemistry, Faculty of Science; Kita 10 Nishi 8,
            Sapporo, Hokkaido 060-0810, Japan
FEATURES             Location/Qualifiers
     CDS             complement(join(973..1013,1074..1439,1488..1785))
                     /gene="zopS"
                     /note="Enoyl CoA isomerase"
BASE COUNT      592 a    504 c    519 g    663 t
ORIGIN
```

```
        1 ttattctgtt cgttgagatc tgtgggtcgc agtgtctcaa aaataaatgt gtccaaagag
       61 gagtttataa aatgctacgc cggatgttta taagctcccg attataaggt gaatgaatat
      121 gcttgtctag gaagctgaga tgctcagatg atatctgccg gctcctaatg cttccgtgca
      181 tcataagcca atgagacctg ataccgtatg caagtacatg caagggtagg tatgaaaata
      241 tctggaggct tatctcgggg tgaaggctgc gtgcctgttc aaacactcag ttcaatgcta
      301 ttatgtgcat gagtcaaatg gcatggcagc tatctactct actcgggcat cttgcaaaca
      361 atcctttcca tgtcgttttt aggaaggttg cattcttctc cctacatacc ccacccagtc
      421 actcctgttt aagccgctgt caacttgatt tgataggctt gaaatggggc ttcatacccc
      481 tttttaaaat gcagccggaa aaaggagccg ctaaatgggt ttgccaagct ttgataggac
      541 gtggaaatat atgaaaagtg tttctcggac tccggcaatg accctggagt agaaaattac
      601 gggcggatta ttactaatgt ccgtagatta ccttaaacat ctaccttcga attggttgcc
      661 cgggctggtt ccgaagttta atatcctaag aatataagtt tttcgtatta tactcttatg
      721 aggccatgct gttttattat tgggtacttg cgctctcggg tgacaattgc tactctcaaa
      781 ctatatgcac gctaagtatc tatgatcggc ctggtcaacc atcgatctgt tgcatctgct
      841 cccacggtat attgcatgca taggtcaagt gttatgttaa agttttaggc tcgtccgctg
      901 ttgttattct actattccca gatccctcca tgccgcgagc caagccccaa attcgacggt
      961 ctcatttatt ctctacagtt tattgaagct cggttgccta acttggccgt ggcctatgta
     1021 atgaagttag tgatatcgtt tcgagcggac ggaaagctag tgcgatgact tactggtgta
     1081 attcaccggt cggactttct gtatacgtgc agtagactgt gcaatagatt ccgcgagtct
     1141 tagggcgtca aaaatggcct cgtccattgt ttctgataca ctggaaacaa attccattct
     1201 gtgtgcgtct tgggcaacga atgctcctcc agacataagc atttcccgga atacactagc
     1261 gccgtccata gtcaacactg caggtggcat ctccgtcatt gctgcctcaa gagaggtgtt
     1321 tttctgcgtc gcgtctccaa gcaaaaacct cgcgtttgat gtgcagatgc gtatgtctga
     1381 gccacaggca atatccaacg ccagttttgt gcagatgccg tgtaagactg atatgatagc
     1441 tttttgatgt gttaacggcc atccaaatcc atggtggcag aactcaccag tattgcattt
     1501 caacggggca atcatgtatg acctgaactt ttcttttgtc gtctccgtat atacctctgt
     1561 gtttgagtcc gtgttgaaag caagggcgct atcatcttta ccaacggcaa ttataataac
     1621 ccgcagcttg gaatcctggg agaggtattc aagcaatctc ttcaacgcca aatctaattc
     1681 tgcatcaggt gttcgagaac acttcacctc gacaatagca acgtaattgc cccggaaatc
     1741 aacacccagt cttgtatgta cagctcttga ctggtcttga gacataatgg aaagagtggc
     1801 cgctgaaaag tccaaactaa tagagtgatg gtgagctagc taccctatga gcaggatacg
     1861 actacttaaa aatggtagaa tacgtgttac tattagatgc tggttattgg catctaggtg
     1921 ggggaaggta tttgcctaaa aggtagccga taagtggtgc cttgcgccga aatgttaccg
     1981 ataagttccc gccataatct tcggtgggct gttgattagt tcccgcaccg ggtagatcgt
     2041 gtcgatccca tttgagaggt catcggatgg ttatacaaag cgagggttac ttacataagc
     2101 ctgaagtaac attctcatac aattctttct ttgccgagac attttagtcc tagctggacc
     2161 ttttggtaaa ggttgcccaa aagcttgaaa attgaagatc caaaagattt ttgaatacat
     2221 tcaaacgaac cgaaaactcc cgaattgacc accttgggtg cgacaagaga cactccct
//
```
